# Rational Design of Minimal Synthetic Promoters for Plants

**DOI:** 10.1101/2020.05.14.095406

**Authors:** Yaomin Cai, Kalyani Kallam, Henry Tidd, Giovanni Gendarini, Amanda Salzman, Nicola J. Patron

**Affiliations:** Engineering Biology, Earlham Institute, Norwich Research Park, Norfolk, NR4 7UZ, UK

**Keywords:** gene regulation, promoters, transcription factors, cis-regulatory elements, plant biotechnology, synthetic biology

## Abstract

Promoters serve a critical role in establishing baseline transcriptional capacity through the recruitment of proteins, including transcription factors (TFs). Previously, a paucity of data for *cis*-regulatory elements in plants meant that it was challenging to determine which sequence elements in plant promoter sequences contributed to transcriptional function. In this study, we have identified functional elements in the promoters of plant genes and plant pathogens that utilise plant transcriptional machinery for gene expression. We have established a quantitative experimental system to investigate transcriptional function, investigating how identity, density and position contribute to regulatory function. We then identified permissive architectures for minimal synthetic plant promoters enabling computational design of a suite of synthetic promoters of different strengths. These have been used to regulate the relative expression of output genes in simple genetic devices.

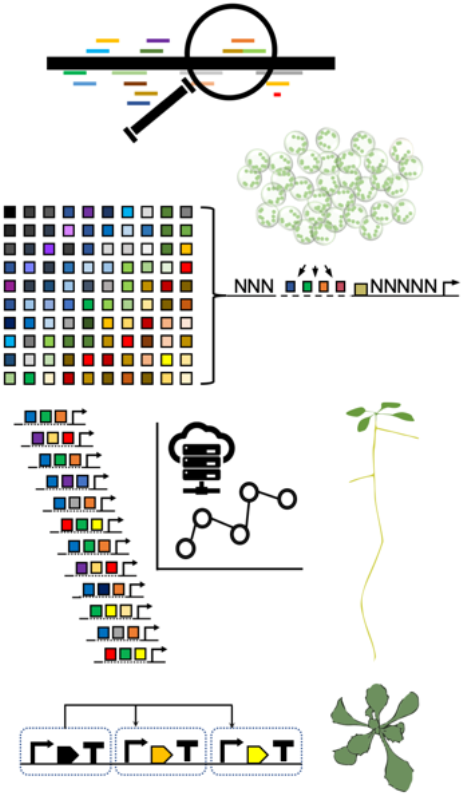

## INTRODUCTION

Transgenic techniques are used to investigate the function of plant genes and to develop new products for agriculture and industry. Biotech crops, typically containing at least one transgene, are now planted on over 190 million hectares each year (1), and plants are finding new roles as platforms for biomanufacturing pharmaceuticals (2, 3). For many years, the majority of transgenic events involved only a single gene of interest and a selectable marker gene. However, recent advances in DNA assembly techniques pioneered by the nascent field of synthetic biology have enabled the facile construction of multigene constructs for plants (4). Researchers are now able to apply these tools to design and deploy synthetic genetic circuits and reconstruct heterologous biochemical pathways in plant systems (5, 6). However, the realisation of synthetic genetic circuits that function as expected requires the ability to precisely and predictably regulate gene expression. Spatiotemporal quantities of endogenous gene products are regulated through numerous mechanisms including transcript elongation (7), antisense transcription (8), and several post-transcriptional and translational processes (9, 10). Although these mechanisms could be leveraged to fine-tune the expression of transgenes, information flow from synthetic genetic circuits is initiated by transcription and, therefore, control of transcription is considered the simplest way to balance the expression of transgenes within a synthetic genetic circuit (11). To achieve this, regulatory elements with predicable characteristics are highly desirable. However, suites of promoters with different levels of expression for plants are not widely available and the promoters that are used are often several kilobases in length and their functional elements have only rarely been identified and characterised. Many plant scientists and biotechnologists still rely on a small set of natural promoters first isolated in the 1980s. In particular, constitutive promoters from plant-infecting DNA viruses or the opine biosynthetic genes found on tumour-inducing (Ti) plasmids of *Agrobacterium tumefaciens* that recruit the host cell’s transcriptional machinery (12). These include the 35s promoter from the double-stranded DNA virus, Cauliflower Mosaic Virus (CaMV), which is reported to have at least partial function in numerous plant species as well as in bacteria (13), fungi (14, 15), and vertebrates (16, 17). Eukaryotic promoter sequences are comprised of complex arrangements of motifs and elements. Early deletion and rearrangement studies identified several key functional elements in the promoters of plant-infecting viruses and bacteria, revealing synergistic interactions between *cis*-elements (18–20). Engineered variants were made by swapping domains to achieve promoters of similar strengths with reduced sequence homologies (21). However, progress towards rational-design of synthetic promoters was limited by a lack of comprehensive data for plant transcription factor binding sites (TFBSs) as well as by the technical limitations of rearranging and rewriting DNA sequences before chemical gene synthesis was widely available. Consequently, although some functional elements were characterised and the possibility of designing synthetic plant promoters was discussed (22), these goals have yet to be fully realised. Progress has been made by engineering variations of natural promoters for tissue-specific expression (23). In addition, a number of synthetic regulatory elements comprised of binding sites for orthogonal TFs fused to a minimal core promoter have been used to enable inducible constitutive expression (24–26). However, to initiate the flow of transcription, these require a non-orthogonal promoter to drive expression of the orthogonal TF, for which natural are typically used (27).

In recent years, significant progress has been made in the design of synthetic regulatory elements for microorganisms, initially with the rational design of ribosome binding sites (28), transcription factors (29) and enhancers (30) and, subsequently, promoters (31–34). Such studies were substantially enabled by comprehensive datasets of TFBSs as well as by the ability to deliver sizable and complex libraries of sequences to populations of cells, sorting, selecting and sequencing cells with desired expression profiles. Equivalent experiments are challenging in plants due to the limitations of DNA-delivery technologies and a paucity of cell-lines. However, genome sequencing technologies have recently shed light on epigenetic states and chromatin accessibility in plant genomes (35) and have identified candidate binding-motifs for many plant transcription factors (36–38). However, genomic datasets alone cannot be used to predict the intrinsic regulatory functions of DNA sequences and assessing the contribution of sequence motifs to regulatory activity is considered essential for characterising function (39). Genome engineering technologies are enabling the functions of specific *cis*-regulatory elements to be dissected (40). Rationally engineered suites of synthetic plant promoters of different strengths have yet to be reported. Here we describe a series of investigations to identify and functionally characterise plant *cis*-regulatory elements, revealing how complexity and relative positions contribute to regulatory functions. We use these data to predict the performance of computationally-designed minimal synthetic constitutive promoters and demonstrate predictable behaviour in dicotyledonous plants in transient expression and when integrated as stable transgenes. Thus, we present suites of minimal synthetic plant promoters of varied strengths, activated by either endogenous or orthogonal transcription factors and demonstrate how these can be used to control the relative expression of output genes in simple genetic circuits.

## MATERIAL AND METHODS

### Identification of candidate transcription factor binding sites (TFBSs)

The Position Weight Matrices (PWMs) from the *Arabidopsis thaliana* (Arabidopsis) cistrome dataset (http://neomorph.salk.edu/dev/pages/shhuang/dap_web/pages/browse_table_aj.php) dataset (36) and the Plant Transcription Factor Database (41) were used to create a motif file for the command line version of FIMO (MEME suite) (42). This was used to scan FASTA files of promoter sequences with a threshold p-value of 0.0001. Candidate TFBSs were mapped back to the promoter sequences. Expression data for TF-encoding genes across multiple Arabidopsis tissues was obtained from the Expression Atlas (http://www.ebi.ac.uk/gxa) (43).

### Construction of plasmids

All constructs were designed in Benchling (San Francisco, CA, USA), synthesised as double-stranded DNA fragments (Twist Biosciences, San Francisco, CA, USA) and cloned into a universal acceptor plasmid (pUAP1 (44) or pUPD2 (45)), to produce standardised Level 0 Phytobricks, conforming to the plant common syntax standard (44). Expression cassettes and multigene constructs were assembled using the Type IIS DNA assembly protocol described in (46). Synthetic and control promoter parts were assembled with the omega 5’ untranslated region from tobacco mosaic virus (5UTR-ΩTMV; pICH41402, Addgene #50285), the coding sequence of firefly luciferase (LucF; pEPAS0CM0008, Addgene T.B.C.), a C-terminal FLAG tag (pICSL50007, Addgene #50308), and a 3’ untranslated region and terminator sequence (3UTR) from *Agrobacterium tumefaciens* octopine synthase (*AtuOCS*) (pICH41432, Addgene #50343). A calibrator construct (pEPYC1CB0197, Addgene T.B.C.) for ratiometric quantification was assembled from *A. tumefaciens* nopaline synthase (*AtuNOS*) promoter (pICH42211, Addgene #50255), 5UTR-ΩTMV, the coding sequence of NanoLuc luciferase (LucN, pEPYC0CM0133, Addgene T.B.C.) and *AtuOCS* terminator. For stable plant transformation, synthetic and control promoters were assembled with the 5’UTR from cowpea mosaic virus (CPMV), a chimeric coding sequence consisting of an N’-terminal HiBit (pEPYC0CM0258, Addgene T.B.C.) the *uidA* coding sequence, encoding β-glucuronidase (GUS; pICSL80016, Addgene #50332) and a C’-terminal yellow fluorescent protein (YFP; pICSL50005, Addgene #117536), and *AtuOCS* terminator. This reporter cassette was assembled with a plant selectable marker cassette conferring resistance to kanamycin (pEPYC1CB0308, Addgene T.B.C.). Synthetic and control promoters were additionally fused to either a transcription activator like effector (TALE) or a synthetic transcription factor comprised of a Gal4 activation domain (GB0900, received from the Orzaez laboratory) and a PhiC3 binding domain (GB_UD_32AB, received from the Orzaez laboratory). A table of all synthetic promoters used in this study is provided in Supplementary Data 1 and tables with the details of all plasmids used and constructed for this study are provided in Supplementary Data 2. All plasmids and their complete sequences have been submitted to the Addgene repository.

### Growth of plant material

*Arabidopsis thaliana* (Col-0), *Nicotiana benthamiana* and *Brassica rapa* plants were germinated and grown in potting medium (two-parts sieved compost to one-part sand) within controlled environment chambers with a 16 h photoperiod at 22 °C with 120–180 μmol/m^2^/s^2^ light intensity. For the two days before leaves were harvested for the preparation of protoplasts, the photoperiod was reduced to 8 h.

### Protoplast preparation and transfection

Protoplasts were prepared from the leaf tissues of *A. thaliana*, *N. benthamiana* and *B. rapa* as previously described (47) and diluted to 10^4^ – 10^5^/mL for transfection. A total of 4.5 μg purified plasmid DNA, comprising equal molar ratios of the plasmid containing the expression cassette for which expression was measured (*test-p*:ΩTMV*:LucF*:*AtuNOSt*) and a calibrating plasmid (pEPYC1CB0197; *AtuNOSp*:ΩTMV*:LucN*:*AtuNOSt*) were added to each designated well of a 2.2 mL 96 deep-well plate containing 200 μL protoplasts (10^4^ – 10^5^ /mL) and mixed gently by shaking. PEG solution was freshly prepared by mixing 2 g PEG (poly(ethylene glycol), MW 4,000 Da) with 2 mL 500 mM mannitol and 0.5 mL 1M CaCl_2_ and 220 μL was added to each well. After 5 mins at room temperature, 1.2 mL W5 (154 mM NaCl, 125 mM CaCl_2_, 5 mM KCl, 2 mM MES pH5.6) was added and protoplasts were collected by centrifugation at 100 g for 2 min and resuspended in 100 μL W5 solution. Resuspended protoplasts were transferred to a round bottom 96-well plate (pre-prepared by incubation with 0.1% bovine serum albumin for 10 mins). Transfected protoplasts were incubated at 22 °C with 100 μmol/m^2^/s^2^ light intensity for at least 16 hrs. For each batch of protoplasts, a control plasmid, pEPYC1CB0199 (*AtuMASp*: ΩTMV*:LucF*:*AtuNOSt)* and the calibrator (pEPYC1CB0197; *AtuNOSp*: ΩTMV*:LucN*:*AtuNOSt*) were used to transfect three aliquots of protoplasts.

### Production of stable transformants

Transgenic Arabidopsis lines were produced by agrobacterium-mediated transformation of floral tissues. Assembled plasmids were transformed into *A. tumefaciens* (GV3101) and liquid cultures were grown from single colonies in growth medium supplemented with 50 μg/mL rifampicin, 25 μg/mL gentamycin and 50 μg/mL kanamycin at 28 °C. *A. tumefaciens* cells were collected by centrifugation and resuspended to OD_600_ 0.8 in 5% sucrose, 0.05% Silvet L-77 and sprayed onto Arabidopsis floral tissues. Plants were sealed in black plastic bags for 24 hours. Seeds were collected from mature siliques and surface sterilised with 70% EtOH for 10 min followed by 3-5% sodium hypochlorite for 10 min. For selection of transgenics, sterilised seeds were germinated and grown on Murashige and Skoog medium supplemented with 75 μg/mL Kanamycin with 16 h light 22 °C.

### Determination of transgene copy number by digital droplet PCR (ddPCR)

Samples of leaf tissue (0.1 g) were ground in liquid nitrogen. DNA was extracted using the cetyltrimethylammonium bromide (CTAB) extraction protocol described in (48) and 2 μg genomic DNA was digested with 20 Units EcoRV for 2 h at 37 °C. 400ng of digested genomic DNA was used in digital droplet PCR (ddPCR) reactions with QX200™ddPCR™EvaGreen®Supermix (Bio-Rad, Hercules, California, USA) and oligonucleotide primers to the *UidA* transgene sequence (5’-CGGCGAAATTCCATACCTGTT and 5’-TCAGCCGATTATCATCACCGA) or a homozygous single-copy reference gene, *AtADH1* (AT1G77120; 5’-ACTTCTCTCTGTCACACCGA and 5’-GGCCGAAGATACGTGGAAAC). Droplets were generated using the QX200™Droplet Generator (Bio-Rad), PCR reactions were run on the C1000 Touch™Thermal Cycler (Bio-Rad) and analysed on the QX200™Droplet Reader (Bio-Rad). Absolute transgene copy number was calculated using the QuantaSoft™software (Bio-rad) to analyse the ratio of droplets in which the target (UidA) was amplified to those in which the reference (*AtADH1)* was amplified.

### Quantification of gene expression

Luciferase expression was detected using the Nano-Glo^®^ Dual-Luciferase^®^ reporter assay system (Promega, Madison, WI, USA). Protoplasts were homogenised in 30 μL passive lysis buffer (Promega) containing protease inhibitor (P9599, Sigma-Aldrich, Dorset, UK). Following incubation on ice for 15 min and centrifugation (100 × g, 2 min, 4 °C), 30 μL supernatant was removed and mixed with 30 μL ONE-Glo™ EX Luciferase Assay Reagent (Promega) and incubated at room temperature for 10 min. LucF luminescence was detected using a Clariostar microplate reader (BMG Labtech, Aylesbury, UK) with a 10 s read time and 1 s settling time. Gain was set at 3,600. LucN luminescence was detected from the same sample by adding 30 μL NanoDLR™ Stop & Glo^®^ Reagent (Promega). After incubation for 10 min at room temperature, luminescence was detected as above. Normalised expression is reported throughout this manuscript as the ratio of luminescence from the test construct (LucF) to the calibrator (LucN; pEPYC1CB0197), normalised to the luminescence of the experiment control (LucF; pEPYC1CB0199/ LucN; pEPYC1CB0197).

Expression from stably-integrated HiBit:GUS:YFP transgenes was quantified using the Nano-Glo® HiBiT Extracellular Detection System (Promega). 10 mg leaf tissue was homogenised in 50 μL passive lysis buffer (Promega) containing protease inhibitor (P9599, Sigma-Aldrich). Homogenised leaf tissues were centrifuged at 18,000 g 10 min 4 °C and 2 μL supernatant mixed with 48 μL Bradford reagent (ThermoFisher Scientific, Waltham, Massachusetts, USA). Protein concentration was estimated by absorbance at 595 nm and concentrations were normalised. 5 μL normalised extract were diluted to 30 μL in passive lysis buffer and mixed with 30 μL Nano-Glo® HiBiT Extracellular Detection Reagent (Promega) and luminescence was detected as above. GUS expression was visualised by submerging 10-day-old seedlings in 0.5 mM K_3_Fe(CN)_6_, 0.5 mM K_4_Fe(CN)_6_, 100 mM sodium phosphate buffer pH7.0, 10 mM EDTA 0.5 mg/mL X-Gluc) for 24 h at room temperature. To remove chlorophyll, this was replaced with 70% EtOH followed by 100% EtOH for 8 h each. Images were taken using a Leica M205FA stereo microscope (Leica, Wetzlar, Germany). YFP expression was visualised using a SP5 (II) confocal microscope (Leica) with a 20x air objective, excitation 514 nm, emission 530 nm. Final images were prepared using Fiji ImageJ (49) (https://imagej.net/Fiji).

## RESULTS

### Constitutive promoters are comprised of multiple functional elements with the potential to bind numerous transcription factors

To identify candidate *cis*-regulatory elements (CREs) for use in minimal synthetic constitutive promoters, we analysed promoters widely used for exogenous expression for the presence of candidate TFBSs. These included promoters from vascular plants as well as those from plant-infecting pathogens that recruit the plant’s transcriptional machinery, including CaMV35S, *Agrobacterium tumefaciens* nopaline synthase (*AtuNOS*) and Mirabilis Mosaic Virus (MMV). The data indicated that constitutive promoters have, in principle, the ability to bind multiple classes of TFs (Figure 1A). Analysis of Arabidopsis gene expression data indicated that few of the TFs predicted to bind to constitutive promoters show constitutive expression themselves (Supplementary Data 3). We also performed *de novo* motif identification using MEME. This analysis identified the presence of a *cis*-regulatory element common to all 14 pathogen promoters (common-CRE or C-CRE) (Supplementary Data 4). In six of these promoters, the C-CRE contained a predicted binding site for a basic-leucine-zipper (bZIP) TF. These C-CREs can therefore be considered to be equivalent to the previously described Activation Sequence 1 (As-1), shown to directly bind members of the TGACG-motif binding (TGA) family of basic-leucine-zipper (bZIP) TFs (50–52). The other eight pathogen promoters were not predicted to bind bZIP TFs. However, it was previously shown that this region of *AtuNOS* is able to bind TGA4 in the presence of a cofactor, OBP5 (53). Consistent with early studies in which regions of promoters were sequentially deleted, specifically deleting individual C-CREs significantly reduced transcriptional output (Figure 1B). This was in contrast to the majority of candidate CREs, of which deletion did not significantly change expression (Supplementary Data 5). To investigate whether the position of the C-CREs within the promoter was essential, the element was relocated varying its proximity to the TSS. While expression reduced when the C-CREs in CaMV35S and MMV were moved further from the TSS, relocating the C-CRE in *AtuNOS,* which is already located distally to the TSS, had a negligible impact (Figure 1C).

**Figure 1.**
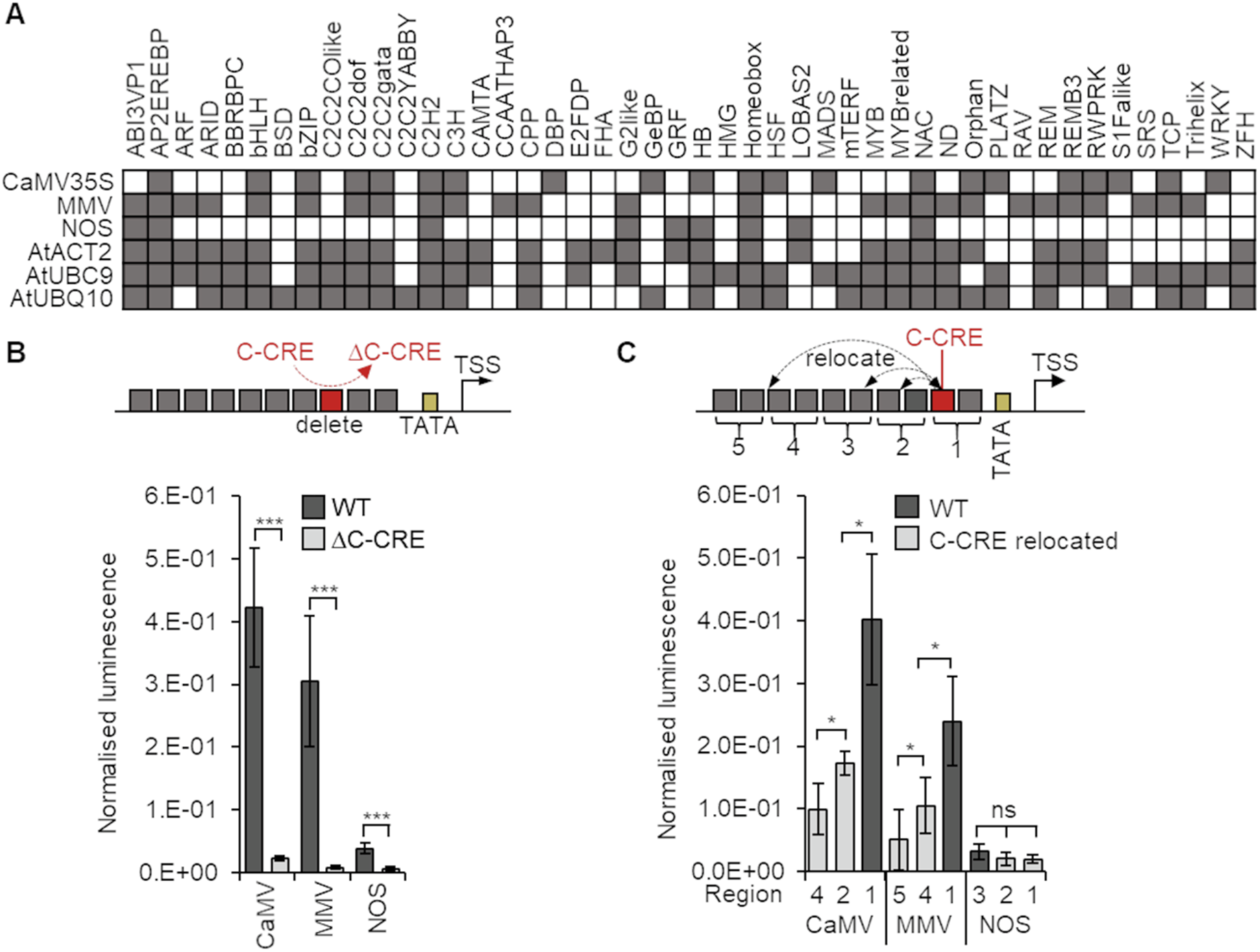
Identification and characterisation of plant *cis*-regulatory elements (CREs). (A) Identification of candidate CREs in constitutive promoters (B) Deletion, or, (C) Relocation of a CRE common to all pathogen promoters (C-CRE) significantly reduces expression. Error bars = 2 × standard error; n=3; P-values were calculated using unpaired two-tailed Student’s t-test; *P < 0.05, ***P < 0.001, ns = not significant.

### Orthogonal tools with a range of expression levels

To identify a functional basic design for minimal synthetic plant promoters (MinSyn-P), we first built and tested synthetic promoters with a range of expression levels that respond to orthogonal TFs. The initial design was based on previously reported synthetic Transcriptional Activator Like Effector (TALE)-responsive synthetic elements to which single binding sites for TALES were added (26). These synthetic promoters consist of 19 bps of random sequence, followed by a second region of random sequence of flexible length (to which CREs are added), a TATA box sequence (TATATAA) and a 43 bp minimal core including transcriptional start site (TSS) (Figure 2A). We successfully verified that this general design could be used to build promoters with a range of expression levels by adding different numbers of binding sites for either TALES or recently described synthetic Gal4:ΦC31 TFs (Vazquez-Vilar *et al.,* 2017) (Figure 2B).

**Figure 2.**
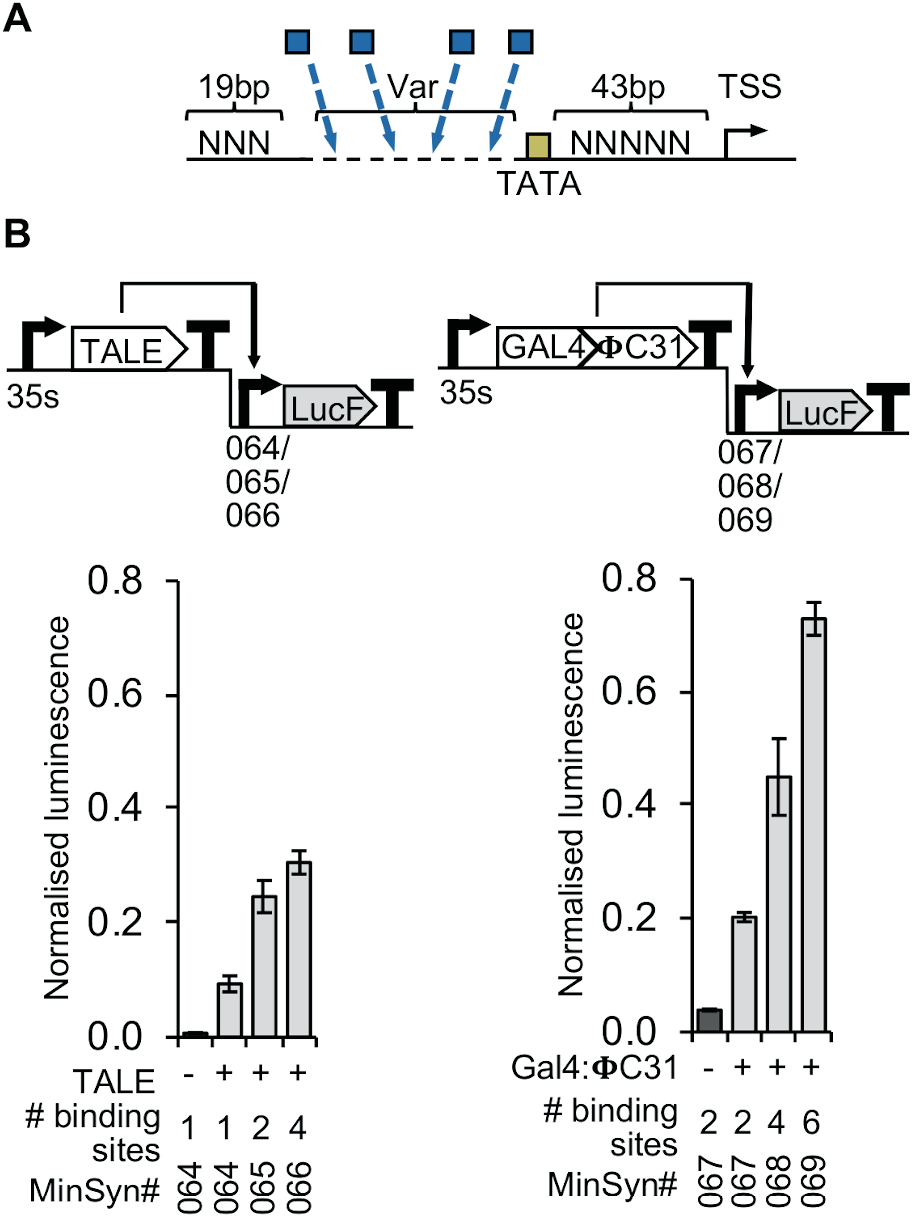
Minimal synthetic promoters (MinSyns) of different strengths regulated by orthogonal transcription factors. (A) MinSyns regulated by Transcriptional Activation Like Effectors (TALES). (B) MinSyns regulated by GAL4:ΦC31 transcription factors; n=3.

### Expression from minimal synthetic regulatory elements by passive cooperativity

To define rules for the design of constitutive MinSyns that respond to endogenous TFs, experiments were progressed to test the function of candidate CREs identified from constitutive promoters (Figure 1). To do this, we first inserted three copies of the same CRE into the variable region of the MinSyn. While it was expected that some candidate CREs might be false-positives, would recruit repressors of transcription (and therefore expression would not be observed), or might need a specific local sequence context, no expression was observed from any MinSyns with only one type of CRE, with the exception of MinSyns containing only multiple copies of C-CREs (Figure 3B). To further investigate the other CREs, we combined random combinations into the variable regions on MinSyns. In the majority of cases, this resulted in significant expression (Figure 3A). In a few cases, combinations of CREs did not result in significant expression (Supplementary Data 6). This is consistent with expectations that some CREs will mainly recruit transcriptional repressors, and that other CREs need to be correctly co-located to enable TFs to form functional heterocomplexes. To test if expression from MinSyns with multiple CREs was dependent on specific TF-TF interactions, the relative positions and spacing of CREs within the variable region of the MinSyns were altered (Figure 3B). In one set of variants, we added up to 20 bp of additional sequence between the CREs. To control for the effect of local sequence context, we made three variants for each set of CREs, two with random sequence and one with the native flanking sequence from the natural promoter from which the CRE was identified. In a second set of variants, the relative positions of the CREs were permutated. Neither changes to the relative position nor moderate increases in spacing had any significant effect on expression. To determine if the relative location of the CREs to the TATA box was critical and to assess if the minimal length of the MinSyns was limiting function, random sequence was inserted between the variable regions containing the CREs and the TATA box. In chromosomal DNA, DNA looping allows distal enhancer elements to interact with the proximal regions, however, as our design goal was minimal constitutive promoters, further extensions to accommodate such interactions were undesirable. Expression was significantly impacted when more than 50 bps of sequence was inserted between the first CRE and the TATA box (Figure 3C).

**Figure 3.**
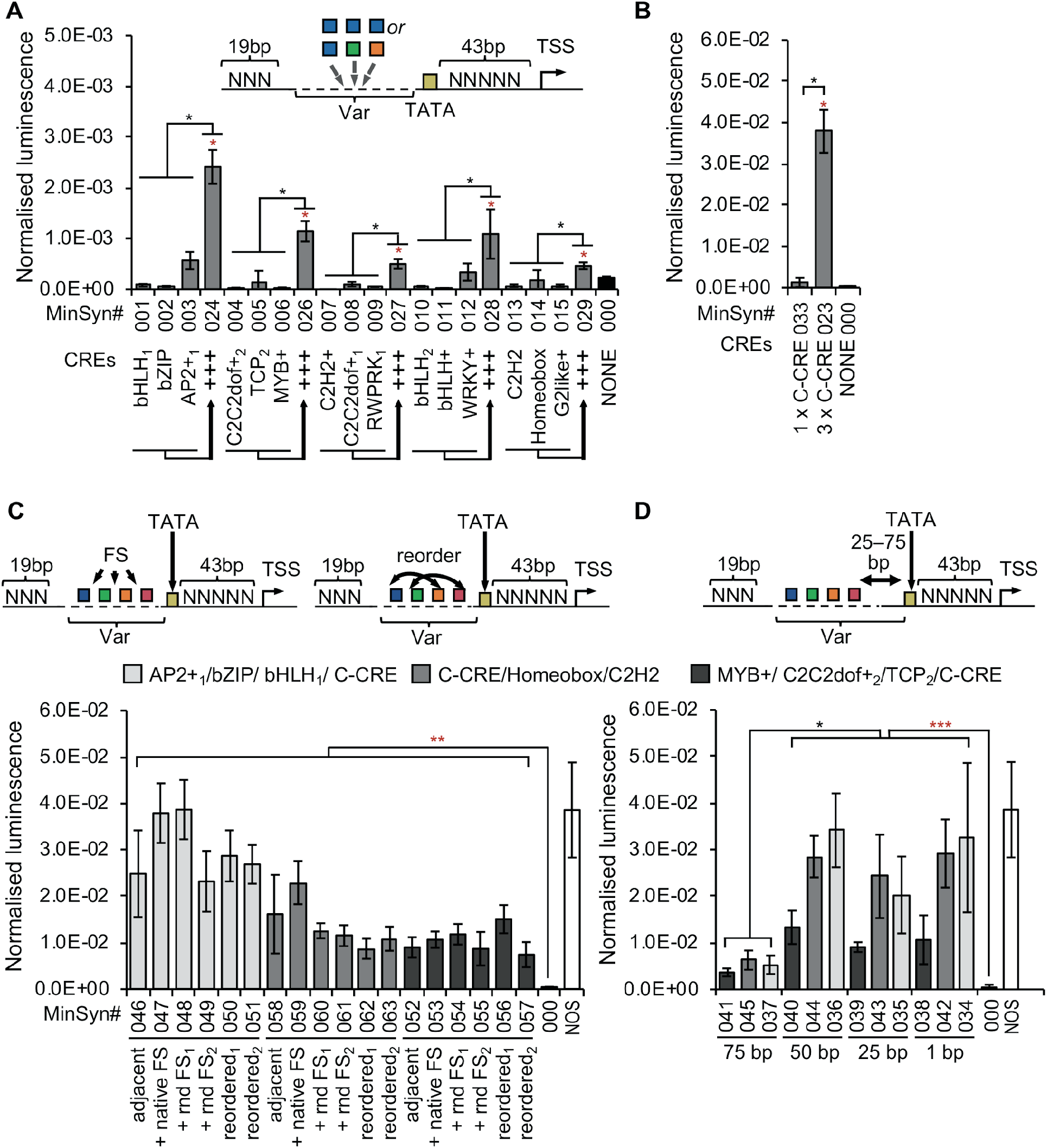
Defining design features of minimal synthetic promoters. (A) Combinations of multiple *cis*-regulatory elements (CREs) enables significant expression compared to repeating copies of the same CRE. (B) The C-CREs enables expression in the absence of other CREs. (C) Rearranging the relative positions of CREs by either inserting native or random flanking sequence (FS) or by reordering does not significantly change expression levels. (D) Relocating CREs greater than 50 base pairs (bp) from the TATA box significantly reduces expression. Error bars = 2 × standard error; n=3; P-values were calculated using unpaired two-tailed Student’s t-test; *P<0.05, **P<0.01, ***P<0.001; Red asterisks indicate a significant difference from MinSyn000, which has no CREs. Black asterisks indicate a significant difference from MinSyns indicated by solid black brackets.

### Computational design of Minimal Synthetic promoters (MinSyn) with predictable strengths

We applied the knowledge gained from these experiments to develop a script to create a library of 1000 constitutive MinSyn (Supplementary Data 7) for which was predicted. For each MinSyn, the script selects a random number (N) between three and ten that defines the number of CREs in the variable region and creates a random DNA sequence of 5 to 30 bases to comprise the sequence of the variable region. It then selects a single CRE sequences from the pool of previously identified CREs. This pool includes two C-CREs predicted to directly bind TGA TFs and one C-CRE for which direct TGA-binding was not predicted. The first CRE is added to the random DNA sequence and the process repeated N times without replacement. Thus, each MinSyn contains between three and ten different CREs, each added to the variable region in the randomly selected order. From our initial experiments, we observed that the strength of the promoters was most affected by the inclusion of multiple CREs of which the C-CREs had the most significant impact. C-CREs predicted to directly bind TGA TFs have the strongest effect when proximal to the TATA box. In contrast, the relative position is less important for C-CREs not predicted to directly bind TGA TFs, with strength only decreasing when located more than 150 bps from the TATA box. These observations were used to assign scores to each nucleotide base as follows: bases within CREs were each assigned a specific score and bases within or flanking C-CREs assigned a higher score adjusted by a numerator reflecting the proximity of that base to the TATA box, with different numerators used for TGA-binding and non-TGA binding C-CREs. Finally, scores were summed and the total was divided by the total number of bases. To convert the score into a predicted strength, we applied the prediction to the existing set of tested MinSyns for which strength had been experimentally determined, thus defining a numerator.

We were therefore able to formulate a predicted expression level for each promoter in the library. As expected for the profile of CREs in the pool, the majority of computational-designed promoters were predicted to have relatively weak expression. Twenty-four MinSyn sequences were selected from the library for synthesis and testing and the predicted and actual levels of expression were compared (Figure 4). For the whole population of 24, the predicted and actual values showed reasonable correlation (Figure 4, red dashed line, R^2^=0.6407), however, there were some outliers. We reanalysed the sequences for the presence of known TFBSs that were not present in the pool used to create the library of MinSyns (e.g. those created unintentionally at sequence junctions of CREs). In 17 cases, additional known TFBS were identified but in most cases, there was insufficient data to determine how the TFs predicted to bind might affect expression (if they were activators or repressors). In three cases, the new motifs were predicted to bind additional TGA, NAC or cytokinin-response factor (CRF) or transcriptional activators that would explain the deviation from the predicted activity (Figure 4, red data points). Four MinSyns were selected for further analysis (Figure 4, blue data points).

**Figure 4.**
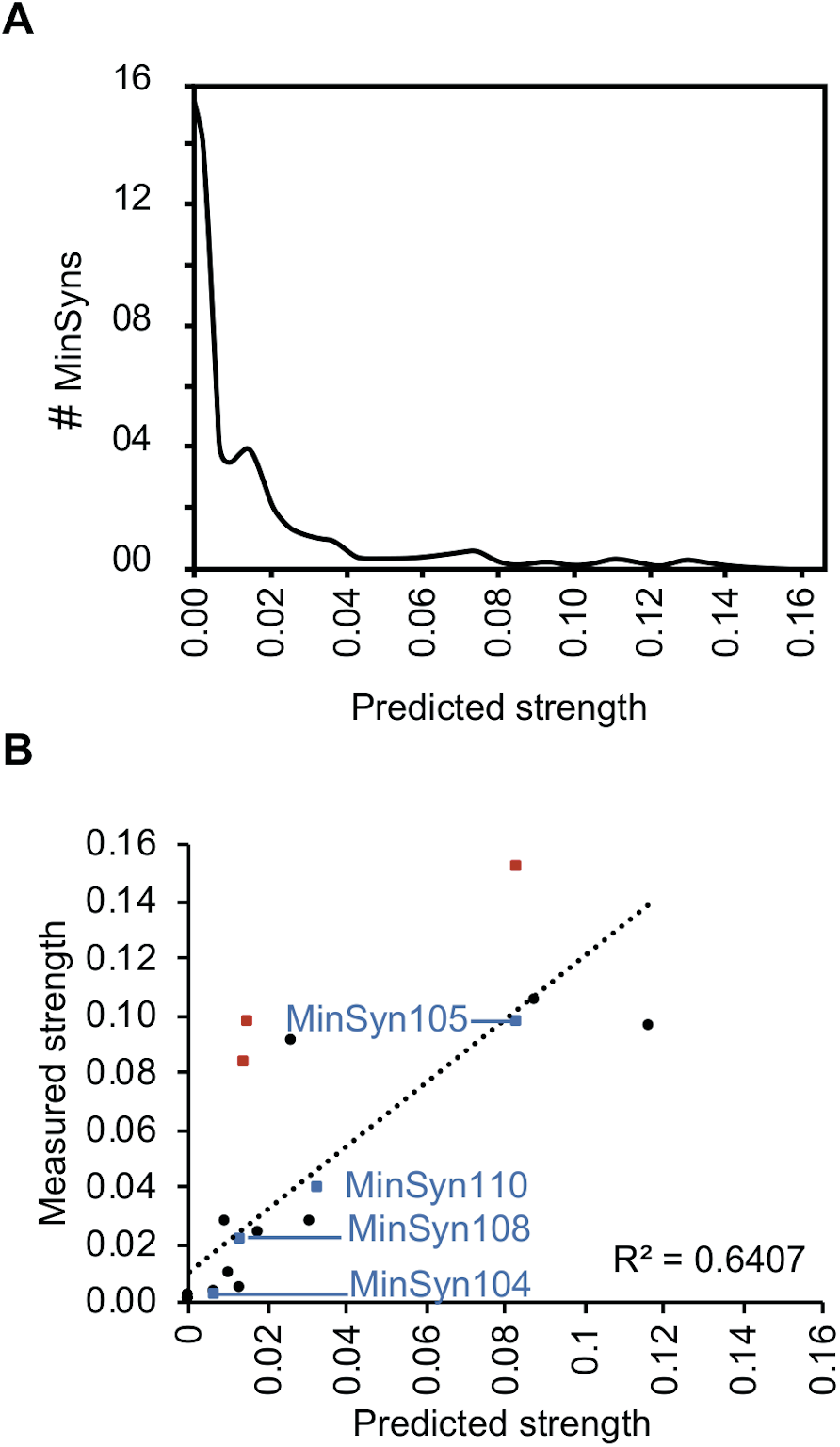
Computational design of Minimal Synthetic Promoters (MinSyns). (A) Of a population of 1000 MinSyns, the majority were predicted to have relatively weak expression (B) Comparison of predicted and measured strengths of 24 computational-designed MinSyns. Red squares indicate MinSyns with unintended *cis*-regulatory elements formed at sequence junction that may explain deviance from predicted strength. Blue squares indicate MinSyns selected for further characterisation (Figure 5).

### MinSyns function in multiple species and as stable transgenes

Transient expression is used both for rapid experimentation and for production-scale protein expression in plants (3, 54). Therefore, minimal promoters that perform as expected in transient expression are useful for several applications. However, for other applications, the ability to maintain expected levels of expression when stably integrated into the genome is desirable. Other studies have reported a strong correlation between the performance of transiently-expressed and stably-integrated transgenes (24). To investigate the performance of MinSyns in stably-integrated transgenes, MinSyns of varying strengths were fused to multifunctional synthetic reporter protein-fusion enabling qualitative and quantitative detection of expression by luminescence, fluorescence and histochemical staining. Patterns of expression were assessed in five independent transgenic lines by GUS-staining and fluorescence microscopy (Figure 5A). Additionally, protein was extracted and expression quantified by detection of luminescence with data normalised to transgene copy number as determined by digital droplet PCR (Figure 5B). As expected, expression levels varied somewhat between independent lines (most likely the effect of local genomic context), however, the MinSyns expressed in most leaf and root tissues and the trends of expression levels observed in transient assays were maintained in stable lines (Figure 5B). We then compared the performance of MinSyns in two additional plant species, *Brassica rapa* and *Nicotiana benthamiana,* in transient protoplast assays. The overall expression trend observed in these species was maintained, with expression levels in *B. oleracea* being comparable to Arabidopsis, but expression levels in *N. benthamiana* being slightly higher (Figure 5b).

**Figure 5.**
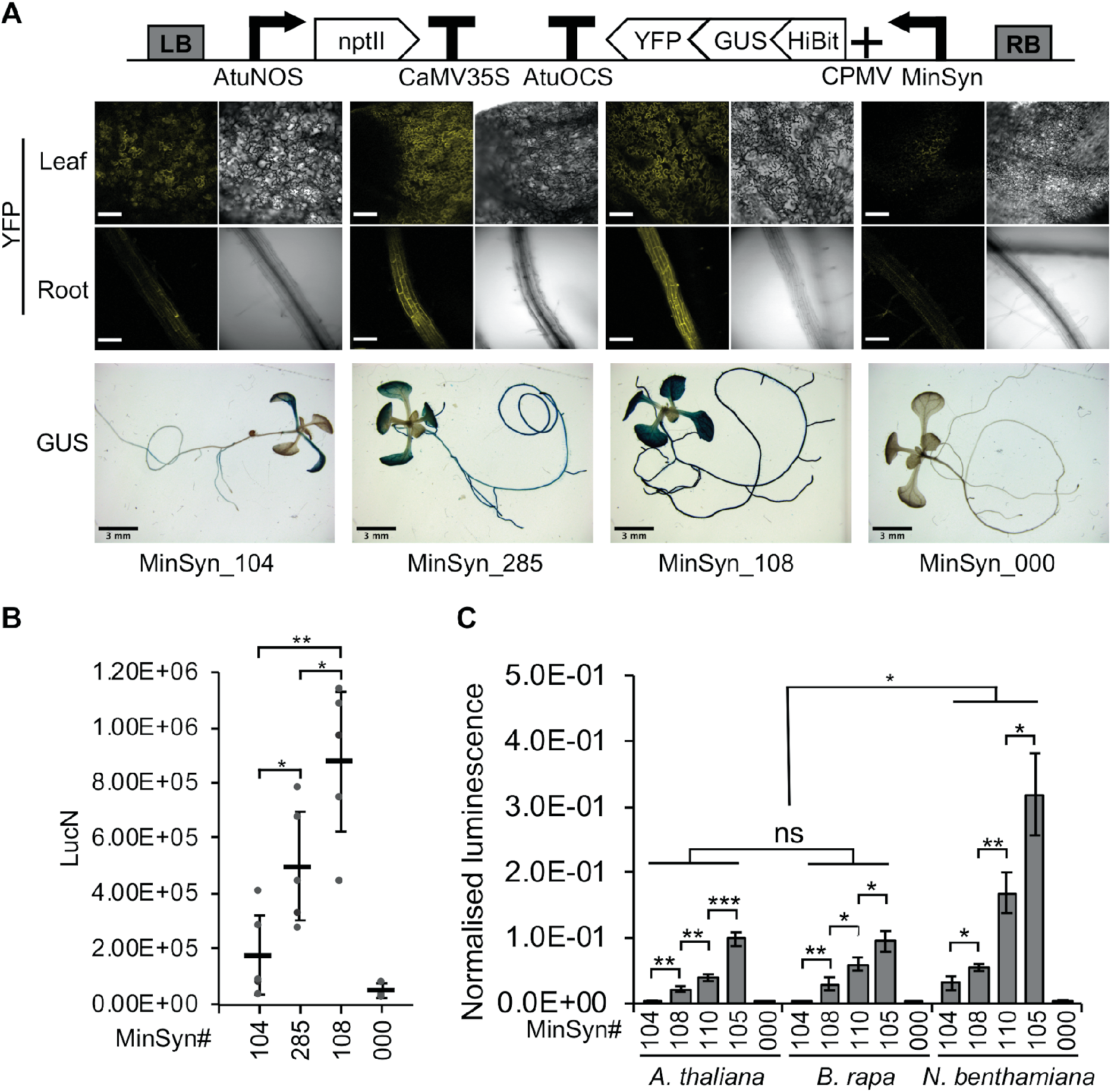
Characterisation of Minimal Synthetic Promoters (MinSyns). (A) Expression from stably integrated transgenes detected by florescence microscopy (yellow fluorescent protein; YFP), scale bar = 150 μm and histochemical staining (β-glucuronidase; GUS), scale bar = 3 mm. (B) Expression levels from stably integrated transgenes quantified by detection of luminescence via Hi-Bit tag and normalised to transgene copy number. (C) Transient expression levels in three plant species. Error bars = 2 × standard error; n=5; P-values were calculated using unpaired two-tailed Student’s t-test; *P ≤0.05, **P≤0.01, ***P≤0.001; ns= not significant.

### Minimal synthetic elements for plants enable relative control of gene expression in synthetic genetic circuits

To demonstrate the utility of MinSyns in synthetic genetic circuits, we constructed simple multigene constructs in which all promoter elements were synthetic. Initially, we simply used a MinSyn to initiate transcriptional flow by controlling expression of an orthogonal TF, which activated expression of reporter (Figure 6A). Similar levels of expression were detected to circuits in which the TF was controlled by CaMV35s, which is widely used to initiate transcription in transgenic plants. We then demonstrated the ability to control the relative ratio of expression of two genes using two MinSyns with different numbers of cognate binding sites for an orthogonal TF to control expression of two reporters and a third MinSyn to control expression of the TF (Figure 6B).

**Figure 6.**
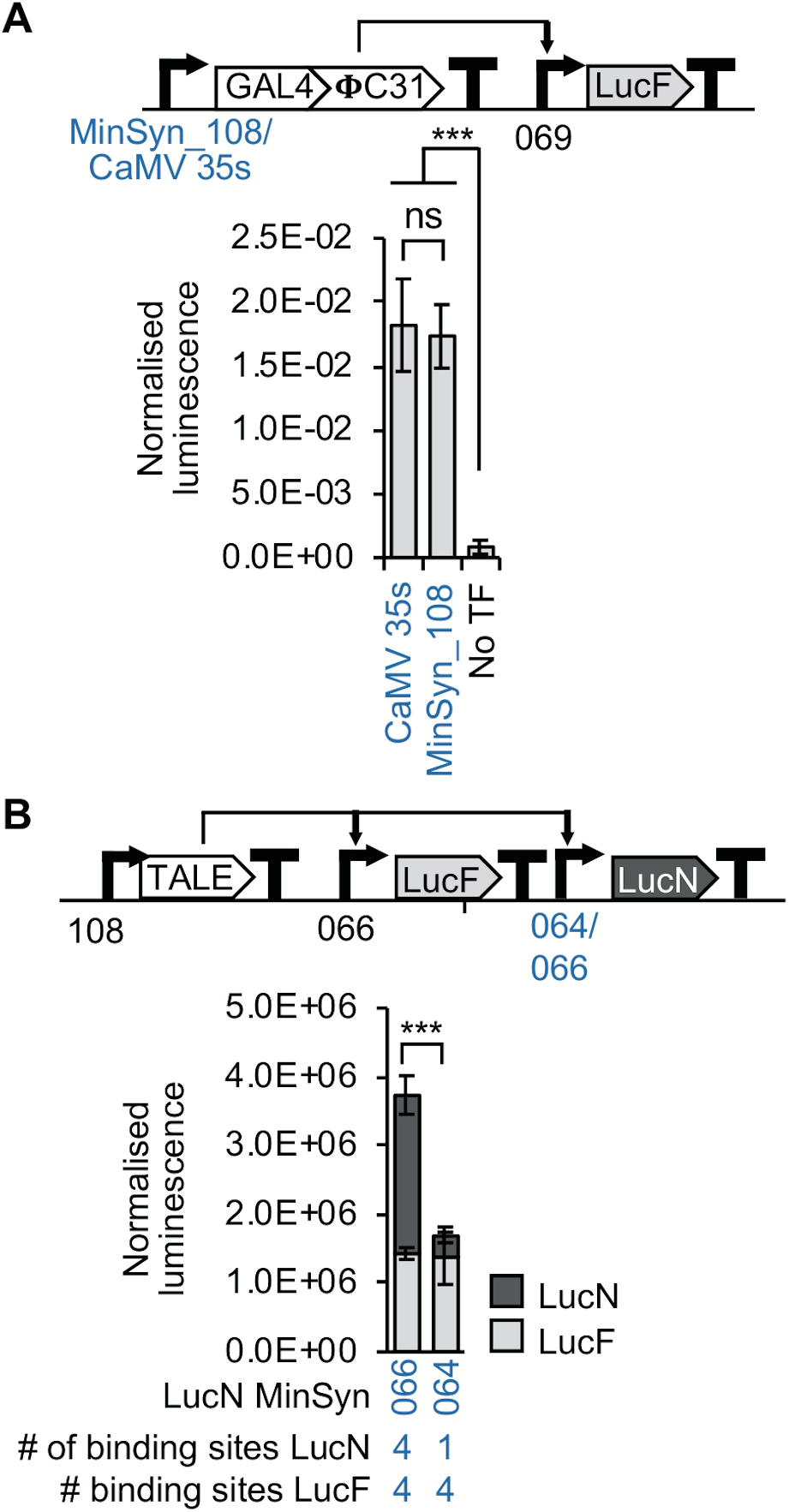
Initiation transcription from simple genetic devices with Minimal Synthetic Promoters (MinSyns). (A) Constitutively expressed MinSyns drive expression of the orthogonal transcriptional factor GAL4:ΦC31, which regulates expression of a reporter. (B) The relative expression of two reporters is regulated using MinSyns with different numbers of binding sites for Transcriptional Activation Like Effectors (TALES). Error bars = 2 × standard error; n=3; P-values were calculated using unpaired two-tailed Student’s t-test; ***P≤0.001; ns = not significant.

## DISCUSSION

Despite their dominance in plant research and biotechnology, comprehensive sequence analyses of even the most widely used constitutive promoters have not previously been reported. Analysis of expression levels of TF predicted to bind to these promoters, indicate that constitutive expression is unlikely to depend on steady-state presence of specific TFs across multiple cell types, but rather on the ability to utilise a wider range of TFs present in different cell types (Figure 1A and Supplementary Data 1). This is consistent with data obtained from early experiments in which the use of specific subdomains of CaMV35S resulted in tissue-specific expression (55). Promoters from numerous plant pathogens that have evolved to utilise the plants transcriptional machinery contain a common regulatory (C-CRE), likely to either directly or indirectly bind the TGA sub-class of bZIP TFs (50–53). This C-CRE has significant effect on the expression levels of both natural and synthetic promoters (Figure 1B and C, Figure 3B) and was the only CRE able to promote detectable levels of expression without the presence of additional functional elements (Figure 3A and B). Several bZIP TFs are known to have a role in different disease and stress response pathways (56–58), which could therefore explain their dominance in pathogen regulatory elements. However, these promoters are known to confer broadly constitutive expression of stably-integrated transgenes, including in healthy, non-stressed plants. Several bZIP TFs have been shown to function as pioneer TFs, able to displace nucleosomes in chromatin inaccessible to other TFs, thus enabling the assembly of other TFs (59, 60). It has recently been hypothesised that some bZIP proteins inhibit chromatin compaction, initiating the formation of enhanceosomes (higher-order multicomponent transcription factor–enhancer complexes) (61). Optimal positioning of pioneer TFs, in particular, has been suggested to be necessary for gene expression (62), which could explain the significant impact of relocation (Figure 1C). However, such roles have yet to be determined for plant TGA TFs. Further, although Transfer-DNAs (T-DNAs) integrated into plant nuclear genomes via *Agrobacterium*-mediated transformation might be packaged into chromatin, thus supporting a role for pioneer TFs in the promoters of the *NOS*, *MAS* and *OCS* opine biosynthetic genes from *A. tumefaciens,* the structures of Caulimovirus DNA in plant nuclei are unknown.

Other than bZIP-binding C-CREs, multiple CREs needed to be combined to obtain significant expression from MinSyns (Figure 3A). These data indicate that, in the absence of the bZIP binding motif, multiple TFs are required for the proper recruitment of the transcriptional machinery. Previous studies have presented evidence that TF-complexes enable transcription either through direct protein-protein interactions or through the formation of enhanceosome complexes, but also without direct protein-protein interactions via a synergistic or collaborative binding process sometimes called passive cooperativity (63, 64). Varying the relative positions and combinations of CREs within the MinSyns variable region revealed that direct protein-protein-interactions were unlikely (Figure 3C), therefore passive cooperativity is a reasonable hypothesis. This is consistent with experiments demonstrating that TFs can be substituted within enhancer complexes, enabling enhancer re-engineering by exchanging TF motifs (65). However, passive cooperativity is proposed to enable the displacement of nucleosomes. While all core histones and the linker histone, H1, have been shown to associate with transiently delivered exogenous DNA in mammalian cells (albeit with aberrant stoichiometry) (66, 67), this has not been investigated in plant cells. Several synthetic promoters and cognate orthogonal TFs for plants, including those that can be induced by chemical signals, have been engineered for plant systems (24–26). In this study we aimed to expand on those efforts, creating regulatory elements of different strengths for use in the construction of larger genetic circuits, particularly biosynthetic pathways, in which it is desirable to control the relative expression levels of different proteins. We provide two options for such constructs: MinSyns of different strengths regulated by endogenous TFs (Figure 5C) or MinSyns of different strengths regulated by synthetic orthogonal TFs (Figure 2 and Figure 6). While the strength of MinSyns that bind orthogonal TFs correlates directly with the number of TF binding sites (Figure 2A, Figure 6A), predicting the strength of constitutive MinSyns that utilise endogenous plant TFs was more challenging. The strength of the computationally designed MinSyns were broadly predictable (Figure 4) but predictability was undermined by the inadvertent introduction of additional TFBSs at sequence junctions. Similar issues were encountered during the creation of synthetic promoters for yeast (68, 69), however the availability of complete datasets of yeast TFBSs allowed programming scripts to be modified to exclude these sequence motifs (68). We considered modifying the script for plant MinSyns to discard sequences in which additional elements were formed, however, as the dataset is incomplete the results would be unpredictable. A second option would be to include any newly created TFBSs in the prediction of strength. This also proved challenging, as relatively few plant TF-DNA interactions have been functionally characterised. In addition, the local context of binding sites has been shown to alter the activity of some TFs from repressors to activators (65, 70), making it difficult to predict the impact on overall expression levels. Indeed, this phenomenon could also contribute to the difference between predicted and observed strengths of some MinSyns.

These investigations have enabled us to design a suite of minimal synthetic plant promoters of varied strengths, activated by either endogenous or orthogonal transcription factors, that provide numerous options for the construction of large and complex genetic circuits for dicotyledonous plants. We have characterised their performance as stable transgenes, finding that transient assays were broadly predictive of behaviour. In previous work, we have observed that permutations of other components such untranslated regions and terminator sequences also impacts the final expression levels of a synthetic transcriptional unit (71). In this work we have controlled for variance by maintaining the same sequences, allowing us to measure the intrinsic properties of the promoters. Further work will be required to determine if and how the properties of MinSyns are modulated when used in combination with different sequence elements.

## ACKNOWLEDGEMENTS

YC, KK and NP conceived the study. YC performed and analysed all experiments with plant cis-regulatory elements. KK, HT and YC performed and analysed experiments with orthogonal transcription factors. YC and GG analysed and visualised gene expression data. AS and YC designed and optimised the ratiometric transient protoplast assay. YC and NP wrote the manuscript and all authors commented and approved. NP was responsible for fundraising and supervision. Plasmids containing Level 0 DNA parts: GB0900 (Gal4-AD) and GB_UD_32AB (PhiC31), GB0036 (35s terminator) were a kind gift from the Orzaez laboratory.

