## Supplementary Data for "Rational Design of Minimal Synthetic Promoters for Plants"

**Supplementary Data 1.** Table of all Minimal Synthetic Promoters (MinSyns) tested in this study. Sequence-variants of cis-regulatory elements (CREs) predicted to bind the same class of transcription factors (TFs) are identified by a subscript numerator (e.g. 3 x $bHLH_1$ and 3 x $bHLH_2$). Some CREs consist of multiple, overlapping TF binding sites indicated by a "+" followed by a list of all TFs predicted to bind to that CRE in parentheses (e.g. $C2C2dof+_2$(C2C2dof+MADS+Orphan+REM). Multiple CREs within in the same MinSyn are separated by "/". Asterisks (*) indicate that the MinSyn sequence is spilt into multiple Level 0 plasmids (e.g. distal, proximal and core regions), which are co-assembled into the expression construct.

| Name | CREs in Var region | Description | Link to sequence | Plasmid Code (Level 0 Phytobrick) | Plasmid Code (Expression cassette) |
|---|---|---|---|---|---|
| MinSyn_000 | NONE | Control MinSyn (no CREs in variable region) | https://benchling.com/s/seq-0qsQE70PiNhTiNi5bHiB | pEPYC0CM0035 | pEPYC1CB0007 |
| MinSyn_001 | 3 x $bHLH_1$ | three copies of a single CRE | https://benchling.com/s/seq-wZFNbyhVnAZN2L4t976N | pEPYC0CM0366 | pEPYC1CB0426 |
| MinSyn_002 | 3 x bZIP | three copies of a single CRE | https://benchling.com/s/seq-1EFNTdKRRpjTkhYw2sDh | pEPYC0CM0367 | pEPYC1CB0427 |
| MinSyn_003 | 3 x $AP2+_1$ (TCP) | three copies of a single CRE | https://benchling.com/s/seq-FgX1DQI9OQ9jr6nHEfuo | pEPYC0CM0368 | pEPYC1CB0428 |
| MinSyn_004 | 3 x $C2C2dof+_2$ (C2C2dof+MADS+Orphan+REM) | three copies of a single CRE | https://benchling.com/s/seq-Ie8CjiZ1fgHjECHX2DEb | pEPYC0CM0379 | pEPYC1CB0439 |
| MinSyn_005 | 3 x $TCP_2$ | three copies of a single CRE | https://benchling.com/s/seq-k6tmMv8g91T31lMmRcyu | pEPYC0CM0380 | pEPYC1CB0440 |
| MinSyn_006 | 3 x MYB+ (MYB+c2c2gata+C3H) | three copies of a single CRE | https://benchling.com/s/seq-YiWhJ0E1fSpbn0D6zcfl | pEPYC0CM0381 | pEPYC1CB0441 |
| MinSyn_007 | 3 x C2H2+ (C2H2+C3H+GeBP) | three copies of a single CRE | https://benchling.com/s/seq-CqGe4PaJqf5EVxmf63YM | pEPYC0CM0372 | pEPYC1CB0432 |
| MinSyn_008 | 3 x $C2C2dof+_1$ (C2C2dof+TCP+NAC) | three copies of a single CRE | https://benchling.com/s/seq-Gu0h6lV0fgko9wq4Ooic | pEPYC0CM0373 | pEPYC1CB0433 |
| MinSyn_009 | 3 x $RWPRK_1$ | three copies of a single CRE | https://benchling.com/s/seq-4DiYhtKMS4jTrO9z8O4t | pEPYC0CM0374 | pEPYC1CB0434 |
| MinSyn_010 | 3 x $bHLH_2$ | three copies of a single CRE | https://benchling.com/s/seq-F3OqlBKFww0p5Ue40ZF1 | pEPYC0CM0385 | pEPYC1CB0445 |
| MinSyn_011 | 3 x bHLH+ | three copies of a single CRE | https://benchling.com/s/seq-Lf2OvoC3FHuWrlDtdO3z | pEPYC0CM0386 | pEPYC1CB0446 |
| MinSyn_012 | 3 x WRKY+ (C2C2dof+WRKY+ABI3VP1+RWPRK+CPP+C2H2+Orphan+REM+ARID) | three copies of a single CRE | https://benchling.com/s/seq-PSxlZv6z9gCLbq50TCnl | pEPYC0CM0387 | pEPYC1CB0447 |
| MinSyn_013 | 3 x C2H2 | three copies of a single CRE | https://benchling.com/s/seq-9Qmk9BuaKbdFtHq2BtW6 | pEPYC0CM0390 | pEPYC1CB0450 |
| MinSyn_014 | 3 x Homeobox | three copies of a single CRE | https://benchling.com/s/seq-wHkZrqd5FzLVbSi82twd | pEPYC0CM0391 | pEPYC1CB0451 |
| MinSyn_015 | 3 x G2like+ (G2like+ABI3VP1+NAC) | three copies of a single CRE | https://benchling.com/s/seq-de11VZwrJhuE9NySaMs7 | pEPYC0CM0392 | pEPYC1CB0452 |
| MinSyn_016 | 3 x $RWPRK_2$ | three copies of a single CRE | https://benchling.com/s/seq-OGYnTcMdmkvb2WEWIhSq | pEPYC0CM0375 | pEPYC1CB0435 |
| MinSyn_017 | 3 x WRKY | three copies of a single CRE | https://benchling.com/s/seq-Ptw5qtT6l7qAcuVQWg7q | pEPYC0CM0376 | pEPYC1CB0436 |
| MinSyn_018 | 3 x DBP | three copies of a single CRE | https://benchling.com/s/seq-PSPzNNkut5USu9SK6J0O | pEPYC0CM0377 | pEPYC1CB0437 |
| MinSyn_019 | 3 x $MADS+_2$ (MADS+C2C2gata) | three copies of a single CRE | https://benchling.com/s/seq-B9obPiQwht5tHuhS89yN | pEPYC0CM0382 | pEPYC1CB0442 |
| MinSyn_020 | 3 x $TCP_3$ | three copies of a single CRE | https://benchling.com/s/seq-V34zUQX5ySxNzGN0qLOb | pEPYC0CM0383 | pEPYC1CB0443 |
| MinSyn_021 | 3 x G2like | three copies of a single CRE | https://benchling.com/s/seq-N6u8ZjGf94wXLTeNUbyn | pEPYC0CM0384 | pEPYC1CB0444 |
| MinSyn_022 | 3 x AP2+ (AP2+LOBAS2) | three copies of a single CRE | https://benchling.com/s/seq-IbRApeJR28SUd6pQg99w | pEPYC0CM0393 | pEPYC1CB0453 |
| MinSyn_023 | 3 x C-CRE | three copies of a single CRE | https://benchling.com/s/seq-PH818sWZDkwfe5LkP1MS | pEPYC0CM0365 | pEPYC1CB0425 |
| MinSyn_024 | $bHLH_1$/bZIP/$AP2+_1$ | three different CREs | https://benchling.com/s/seq-Pdin3HP2AviGfrRDnB60 | pEPYC0CM0431 | pEPYC1CB0493 |
| MinSyn_026 | $C2C2dof+_2$/$TCP_2$/MYB+ | three different CREs | https://benchling.com/s/seq-HT8QBHITlc0aUofyzNtH | pEPYC0CM0435 | pEPYC1CB0497 |
| MinSyn_027 | C2H2+/$C2C2dof+_1$/$RWPRK_1$ | three different CREs | https://benchling.com/s/seq-SUOKjZKcdhbo042o130i | pEPYC0CM0433 | pEPYC1CB0495 |
| MinSyn_028 | $bHLH_2$/bHLH+/WRKY+ | three different CREs | https://benchling.com/s/seq-3JYJRKIjujcE0QF8fe36 | pEPYC0CM0437 | pEPYC1CB0499 |
| MinSyn_029 | C2H2/Homeobox/G2like+ | three different CREs | https://benchling.com/s/seq-0tR8wRGTga3dWw2ojdUw | pEPYC0CM0439 | pEPYC1CB0501 |
| MinSyn_030 | $RWPRK_2$/WRKY+/DBP | three different CREs | https://benchling.com/s/seq-QzeRW1REYLnxpLKTQ8K5 | pEPYC0CM0434 | pEPYC1CB0496 |
| MinSyn_031 | $MADS+_2$/$TCP_3$/G2like | three different CREs | https://benchling.com/s/seq-cjnH9arWrL87HofRr8p2 | pEPYC0CM0388 | pEPYC1CB0498 |
| MinSyn_032 | Homeobox/$G2like_2$/$AP2+_2$ | three different CREs | https://benchling.com/s/seq-qOrS1fNh1I3sknYW7YZE | pEPYC0CM0440 | pEPYC1CB0502 |
| MinSyn_033 | 1 x C-CRE | single copy of the C-CRE | https://benchling.com/s/seq-e1nuSZ2sZzYsudP557Jh | pEPYC0CM0041 | pEPYC1CB0079 |
| MinSyn_034 | $AP2+_1$/bZIP/$bHLH_1$/C-CRE | three different CREs + C-CRE | https://benchling.com/s/seq-esQkM6GttTF3qfAQAQND | pEPYC0CM0310 | pEPYC1CB0352 |
| MinSyn_035 | $AP2+_1$/bZIP/$bHLH_1$/C-CRE | 25 bp between CREs and TATA | https://benchling.com/s/seq-AvdnsDcbLsTGoydoVsOf | pEPYC0CM0312 | pEPYC1CB0354 |
| MinSyn_036 | $AP2+_1$/bZIP/$bHLH_1$/C-CRE | 50 bp between CREs and TATA | https://benchling.com/s/seq-i6vT6WeKtnr5SauKJDR2 | pEPYC0CM0313 | pEPYC1CB0355 |
| MinSyn_037 | $AP2+_1$/bZIP/$bHLH_1$/C-CRE | 75 bp between CREs and TATA | https://benchling.com/s/seq-gskfrYVYdC3Fi8vEOL8z | pEPYC0CM0314 | pEPYC1CB0356 |
| MinSyn_038 | MYB+/$C2C2dof+_2$/$TCP_2$/C-CRE | three different CREs + C-CRE | https://benchling.com/s/seq-apzE3SCGRK74bHdhJlvS | pEPYC0CM0225 | pEPYC1CB0269 |
| MinSyn_039 | MYB+/$C2C2dof+_2$/$TCP_2$/C-CRE | 25 bp between CREs and TATA | https://benchling.com/s/seq-Eu3N4i4XhQDJbSHC3Ovb | pEPYC0CM0209 | pEPYC1CB0253 |
| MinSyn_040 | MYB+/$C2C2dof+_2$/$TCP_2$/C-CRE | 50 bp between CREs and TATA | https://benchling.com/s/seq-3ghoZvi65XsWYie06FCb | pEPYC0CM0210 | pEPYC1CB0254 |
| MinSyn_041 | MYB+/$C2C2dof+_2$/$TCP_2$/C-CRE | 75 bp between CREs and TATA | https://benchling.com/s/seq-LGRCpftG1qJMzZj67PFb | pEPYC0CM0211 | pEPYC1CB0255 |
| MinSyn_042 | C-CRE/Homeobox/C2H2 | two different CREs + C-CRE | https://benchling.com/s/seq-Mjgku0iWBVrSI1v132qv | pEPYC0CM0311 | pEPYC1CB0353 |
| MinSyn_043 | C-CRE/Homeobox/C2H2 | 25 bp between CREs and TATA | https://benchling.com/s/seq-llYzHiCBO6XzLOE1zowj | pEPYC0CM0212 | pEPYC1CB0256 |
| MinSyn_044 | C-CRE/Homeobox/C2H2 | 50 bp between CREs and TATA | https://benchling.com/s/seq-qu6hTPHmly538zhUWBUe | pEPYC0CM0213 | pEPYC1CB0257 |
| MinSyn_045 | C-CRE/Homeobox/C2H2 | 75 bp between CREs and TATA | https://benchling.com/s/seq-apzE3SCGRK74bHdhJlvS | pEPYC0CM0214 | pEPYC1CB0258 |
| MinSyn_046 | $AP2+_1$/bZIP/$bHLH_1$/C-CRE | three different CREs + C-CRE | https://benchling.com/s/seq-TBv49IqInULaRzZrA6z7 | pEPYC0CM0182 | pEPYC1CB0187 |

| Name | CREs in Var region | Description | Link to sequence | Plasmid Code (Level 0 Phytobrick) | Plasmid Code (Expression cassette) |
|---|---|---|---|---|---|
| MinSyn_047 | AP2+$_{1}$/bZIP/bHLH$_{1}$/C-CRE | Native flanking sequences for each CREs, up to 10 bp for each CRE | https://benchling.com/s/seq-esQkM6GttTF3qfAQAQND | pEPYC0CM0310 | pEPYC1CB0352 |
| MinSyn_048 | AP2+$_{1}$/bZIP/bHLH$_{1}$/C-CRE | Random flanking sequence between CREs | https://benchling.com/s/seq-m70LnNmMxibPARIVXdTq | pEPYC0CM0395 | pEPYC1CB0455 |
| MinSyn_049 | AP2+$_{1}$/bZIP/bHLH$_{1}$/C-CRE | Random flanking sequence between CREs | https://benchling.com/s/seq-hKF7cEnrzRI7ZvmT8TQH | pEPYC0CM0396 | pEPYC1CB0456 |
| MinSyn_050 | bZIP/C-CRE/bHLH$_{1}$/AP2+$_{1}$ | three different CREs + C-CRE | https://benchling.com/s/seq-JYXCbEXuLujUG4vRS6Sa | pEPYC0CM0451 | pEPYC1CB0509 |
| MinSyn_051 | bHLH$_{1}$/AP2+$_{1}$/C-CRE/bZIP | three different CREs + C-CRE | https://benchling.com/s/seq-4ysOthO8IwMi4Gza4KvA | pEPYC0CM0452 | pEPYC1CB0510 |
| MinSyn_052 | MYB+/C2C2dof+$_{2}$/TCP$_{2}$/C-CRE | three different CREs + C-CRE | https://benchling.com/s/seq-84vUPFSvAkNf4HlawIf2 | pEPYC0CM0183 | pEPYC1CB0188 |
| MinSyn_053 | MYB+/C2C2dof+$_{2}$/TCP$_{2}$/C-CRE | Native flanking sequences for each CREs, up to 10 bp for each CRE | https://benchling.com/s/seq-x2PKDjg86Cj55qiL0XGt | pEPYC0CM0225 | pEPYC1CB0269 |
| MinSyn_054 | MYB+/C2C2dof+$_{2}$/TCP$_{2}$/C-CRE | Random flanking sequence between CREs | https://benchling.com/s/seq-16dbTNGncrGBQ98PC2fq | pEPYC0CM0397 | pEPYC1CB0457 |
| MinSyn_055 | MYB+/C2C2dof+$_{2}$/TCP$_{2}$/C-CRE | Random flanking sequence between CREs | https://benchling.com/s/seq-SLE1so0KdCvy4Ceg2GoO | pEPYC0CM0398 | pEPYC1CB0458 |
| MinSyn_056 | TCP$_{2}$/MYB+/C-CRE/C2C2dof+$_{2}$ | three different CREs + C-CRE | https://benchling.com/s/seq-p5YhEx6oyxBxS2jEiHpq | pEPYC0CM0453 | pEPYC1CB0511 |
| MinSyn_057 | C2C2dof+$_{2}$/TCP$_{2}$/MYB+/C-CRE | three different CREs + C-CRE | https://benchling.com/s/seq-cTjP9K65UEBrXAVDv6pp | pEPYC0CM0454 | pEPYC1CB0512 |
| MinSyn_058 | C-CRE/Homeobox/C2H2 | two different CREs + C-CRE | https://benchling.com/s/seq-QSCATVwH0gJ3UiGp2JCX | pEPYC0CM0184 | pEPYC1CB0189 |
| MinSyn_059 | C-CRE/Homeobox/C2H2 | Native flanking sequences for each CREs, up to 10 bp for each CRE | https://benchling.com/s/seq-Mjgku0iWBVrSI1v132qv | pEPYC0CM0311 | pEPYC1CB0353 |
| MinSyn_060 | C-CRE/Homeobox/C2H2 | Random flanking sequence between CREs | https://benchling.com/s/seq-mGTHb9piclAMIRJRPBpW | pEPYC0CM0401 | pEPYC1CB0461 |
| MinSyn_061 | C-CRE/Homeobox/C2H2 | Random flanking sequence between CREs | https://benchling.com/s/seq-ojbsHEj0CNtVv71B8JPE | pEPYC0CM0402 | pEPYC1CB0462 |
| MinSyn_062 | C2H2/Homeobox/C-CRE | two different CREs + C-CRE | https://benchling.com/s/seq-i7GRR4HRQxNjR00gv1Ty | pEPYC0CM0455 | pEPYC1CB0513 |
| MinSyn_063 | Homeobox/C-CRE/C2H2 | two different CREs + C-CRE | https://benchling.com/s/seq-qyco6lU97tfWXfMmb9Gp | pEPYC0CM0456 | pEPYC1CB0514 |
| MinSyn_064 | 1 x TALE AvrXa27 binding site | Binding site for synthetic TF | https://benchling.com/s/seq-MkgA2YoAajmr7xhfPyBJ | pEPAS0CMR0015 | pEPKKα2KN0091/<br>pEPYCα2rKN0010 |
| MinSyn_065 | 2 x TALE AvrXa27 binding sites | Binding site for synthetic TF | https://benchling.com/s/seq-slokOMWiDRAxfmqqNsjN | pEPAS0CMR0016 | pEPKKα2KN0092 |
| MinSyn_066 | 4 x TALE AvrXa27 binding sites | Binding site for synthetic TF | https://benchling.com/s/seq-gDZylOe6miWzsyrgor0n | pEPKK0CM0107 | pEPKKα2KN0093/<br>pEPYCα2rKN0009/<br>pEPYCα1KN0008 |
| MinSyn_067 | 2 x ϕC3 binding sites | Binding site for synthetic TF | https://benchling.com/s/seq-fgzjQewCH5vr3GfCEkiG | pEPKK0CM0065 | pEPKKα2KN0100 |
| MinSyn_068 | 4 x ϕC3 binding sites | Binding site for synthetic TF | https://benchling.com/s/seq-t0a3QQRnQ7uywEow0mdL<br>https://benchling.com/s/seq-JioHeeBtqPHTH6h1YVs4 | pEPKK0CM0062 +<br>pEPKK0CM0111* | pEPKKα2KN0101 |
| MinSyn_069 | 6 x ϕC3 binding sites | Binding site for synthetic TF | https://benchling.com/s/seq-NpXSAnIIWwDqevLWFfSt<br>https://benchling.com/s/seq-aoVmB0iw0bGZJpfb4zu4<br>https://benchling.com/s/seq-JioHeeBtqPHTH6h1YVs4 | pEPKK0cm0063 +<br>pEPKK0CM0064 +<br>pEPKK0CM0111* | pEPKKα2KN0102/<br>pEPYCα1KN0003 |
| MinSyn_101 | bHLH$_{2}$/bHLH$_{1}$/AP2+$_{1}$/Homeobox/WRKY | computationally designed | https://benchling.com/s/seq-4DGAEuNIAqGYKpgxFpuE | pEPYC0CM0407 | pEPYC1CB0467 |
| MinSyn_102 | C2C2dof+/AP2+$_{1}$/WRKY | computationally designed | https://benchling.com/s/seq-8fiNn13Fn41xS0R1NwAa | pEPYC0CM0408 | pEPYC1CB0468 |
| MinSyn_103 | bHLH$_{1}$/MYB+ | computationally designed | https://benchling.com/s/seq-viwbvfSENd3wupQtCuh8 | pEPYC0CM0409 | pEPYC1CB0469 |
| MinSyn_104 | RWPRK$_{1}$/CCAAT+/HSF+/MADS+/AP2+$_{2}$/G2like/DBP/C-CRE/MYB+ | computationally designed | https://benchling.com/s/seq-UkP4FqWF5RxWyr0d4wNf | pEPYC0CM0410 | pEPYC1CB0470 |
| MinSyn_105 | bZIP/C-CRE/Homeobox/MYB+/C-CRE | computationally designed | https://benchling.com/s/seq-OeJ5B32kVIqfCwcM449w | pEPYC0CM0411 | pEPYC1CB0471 |
| MinSyn_106 | C-CRE/AP2+$_{1}$/HSF+/C2C2dof+$_{2}$/CCAAT+/C-CRE | computationally designed | https://benchling.com/s/seq-MzLsjqJF4Wk33AQyMV5N | pEPYC0CM0412 | pEPYC1CB0472 |
| MinSyn_107 | RWPRK$_{2}$/C-CRE/WRKY/G2like+/C-CRE | computationally designed | https://benchling.com/s/seq-kbeGUgNxVrqNfb80nFY2 | pEPYC0CM0413 | pEPYC1CB0473 |
| MinSyn_108 | bHLH$_{1}$/C2H2/AP2+$_{2}$/AP2+$_{1}$/Homeobox/C-CRE | computationally designed | https://benchling.com/s/seq-PLJyq4WT8G03wG3gQY8o | pEPYC0CM0414 | pEPYC1CB0474/pEPYCα1KN0002/pEPYCα1KN0007 |
| MinSyn_109 | bHLH$_{1}$/C-CRE/MYB+/C-CRE | computationally designed | https://benchling.com/s/seq-vH4V5K7vNg2X7tWDlGy4 | pEPYC0CM0415 | pEPYC1CB0475 |
| MinSyn_110 | TCP$_{2}$/RWPRK$_{1}$/bHLH$_{1}$/MYB+/C-CRE/C-CRE | computationally designed | https://benchling.com/s/seq-sX2AR6aULzb2lyuoMLYX | pEPYC0CM0416 | pEPYC1CB0476 |
| MinSyn_111 | MADS+$_{1}$/MYB+/C-CRE/WRKY+/MADS+$_{2}$/C2H2+ | computationally designed | https://benchling.com/s/seq-pBG4wmcNc3SwSlXKuhWS | pEPYC0CM0417 | pEPYC1CB0479 |
| MinSyn_112 | TCP$_{3}$/C2H2+/Homeobox | computationally designed | https://benchling.com/s/seq-VkwMb37FrE9KbWSkpxI5 | pEPYC0CM0418 | pEPYC1CB0480 |
| MinSyn_113 | C-CRE/Homeobox/AP2+$_{2}$/MADS+$_{1}$/WRKY/bHLH$_{2}$ | computationally designed | https://benchling.com/s/seq-odcECHuFtf3378P8NuD1 | pEPYC0CM0419 | pEPYC1CB0481 |
| MinSyn_114 | C-CRE/C2H2+/G2like/MADS+$_{1}$/AP2+$_{2}$/bHLH$_{1}$/RWPRK$_{2}$ | computationally designed | https://benchling.com/s/seq-1plZVxeHhgxrTP8IH7Fz | pEPYC0CM0420 | pEPYC1CB0482 |
| MinSyn_115 | CCAAT+/C2H2+/DBP | computationally designed | https://benchling.com/s/seq-gtv2NB02AKAKqMo9z4M3 | pEPYC0CM0421 | pEPYC1CB0483 |
| MinSyn_116 | RWPRK$_{2}$/bHLH+/C-CRE/TCP$_{1}$/bHLH$_{2}$/bZIP/bHLH$_{1}$ | computationally designed | https://benchling.com/s/seq-NTaiQg01dsWOX8fZ2nik | pEPYC0CM0422 | pEPYC1CB0484 |
| MinSyn_117 | MADS+$_{1}$/bHLH$_{1}$/C2H2+/bHLH+/bZIP/WRKY/G2like+/TCP$_{3}$/AP2+$_{1}$ | computationally designed | https://benchling.com/s/seq-LPnjtxz4pTd5Clrc5IAC | pEPYC0CM0423 | pEPYC1CB0485 |
| MinSyn_118 | Homeobox/C-CRE | computationally designed | https://benchling.com/s/seq-917cevKHSyFgOWBg3jra | pEPYC0CM0424 | pEPYC1CB0486 |

| Name | CREs in Var region | Description | Link to sequence | Plasmid Code (Level 0 Phytobrick) | Plasmid Code (Expression cassette) |
|---|---|---|---|---|---|
| MinSyn_119 | $TCP_2$/$TCP_3$/C2H2+/$AP2+_2$/DBP/C2H2/C-CRE | computationally designed | https://benchling.com/s/seq-1WZttWPPAJjzSQqTEmkS | pEPYC0CM0425 | pEPYC1CB0487 |
| MinSyn_120 | C-CRE/HSF+ | computationally designed | https://benchling.com/s/seq-0ngHrUjGdIIUWOUsjwBz | pEPYC0CM0426 | pEPYC1CB0488 |
| MinSyn_121 | $MADS+_1$/$C2C2dof+_1$/bHLH+/$TCP_1$/$MADS+_2$/$AP2+_2$/DBP/C2H2/$AP2+_1$ | computationally designed | https://benchling.com/s/seq-HFTwco2h8HqVVgcKtsoT | pEPYC0CM0427 | pEPYC1CB0489 |
| MinSyn_122 | C2H2+/Homeobox/$AP2+_2$/$AP2+_1$/$RWPRK_2$ | computationally designed | https://benchling.com/s/seq-bnSRl1g6bO8GMnIJlThr | pEPYC0CM0428 | pEPYC1CB0490 |
| MinSyn_123 | WRKY+/Homeobox/$RWPRK_2$/CCAAT+/C2C2dof+/G2like+/DBP/C-CRE/bZIP | computationally designed | https://benchling.com/s/seq-n1wTXpDjDpO1EEmi2CES | pEPYC0CM0429 | pEPYC1CB0491 |
| MinSyn_124 | MYB+/$MADS+_2$/$AP2+_2$/$AP2+_1$/C-CRE | computationally designed | https://benchling.com/s/seq-7EzWHcgHI3lvJTQBBtuB | pEPYC0CM0430 | pEPYC1CB0492 |
| MinSyn_285 | bZIP/Homeobox/$C2C2dof+_2$/C-CRE | computationally designed | https://benchling.com/s/seq-EouughPWqfDqXnyTEFR9 | pEPYC0CM0244 | pEPYC1CB0286 |

**Supplementary Data 2** List of Plasmids

**1. Level 0 Phytobricks**

| Addgene # | Plasmid code | Part type | Description/Origin | Compatibility with Assembly Systems | Cloning overhang (top strand) | | Source of plasmid |
|---|---|---|---|---|---|---|---|
| | | | | | 5' | 3' | |
| #50270 | pICSL12006 | PROM+5UTR | CsVMV (Cassava Vein Mosaic Virus) | MoClo, Loop, GB | GGAG | AATG | Engler et al(2014) |
| #50272 | pICH85281 | PROM | MAS (Agrobacterium tumefaciens) | MoClo, Loop, GB | GGAG | AATG | Engler et al(2014) |
| #50255 | pICH42211 | PROM | NOS (Agrobacterium tumefaciens) | MoClo, Loop, GB | GGAG | TACT | Engler et al(2014) |
| - | GB0552 | PROM+5UTR | 35s (Cauliflower Mosaic Virus) | GB, Loop | GGAG | CCAT | Vazquez-Vilar et al (2017) |
| T.B.C | pEPYC0CM0071 | PROM | 35s (Cauliflower Mosaic Virus) | MoClo, Loop, GB | GGAG | TACT | This study |
| T.B.C | pEPYC0CM0095 | PROM | MMV (Mirabalis Mosaic Virus) | MoClo, Loop, GB | GGAG | TACT | This study |
| T.B.C | pEPYC0CM0084 | PROM | CaMV35S (C-CRE relocated) | MoClo, Loop, GB | GGAG | TACT | This study |
| T.B.C | pEPYC0CM0089 | PROM | CaMV35S (C-CRE relocated) | MoClo, Loop, GB | GGAG | TACT | This study |
| T.B.C | pEPYC0CM0099 | PROM | MMV (C-CRE relocated) | MoClo, Loop, GB | GGAG | TACT | This study |
| T.B.C | pEPYC0CM0115 | PROM | MMV (C-CRE relocated) | MoClo, Loop, GB | GGAG | TACT | This study |
| T.B.C | pEPYC0CM0120 | PROM | NOS (C-CRE relocated) | MoClo, Loop, GB | GGAG | TACT | This study |
| T.B.C | pEPYC0CM0119 | PROM | NOS (C-CRE relocated) | MoClo, Loop, GB | GGAG | TACT | This study |
| T.B.C | pEPYC0CM0168 | PROM | 35s (ΔAP2+$_1$) | MoClo, Loop, GB | GGAG | TACT | This study |
| T.B.C | pEPYC0CM0170 | PROM | 35s(ΔbHLH$_1$) | MoClo, Loop, GB | GGAG | TACT | This study |
| T.B.C | pEPYC0CM0169 | PROM | 35s(ΔbZIP) | MoClo, Loop, GB | GGAG | TACT | This study |
| T.B.C | pEPYC0CM0171 | PROM | 35s(ΔC_CRE) | MoClo, Loop, GB | GGAG | TACT | This study |
| T.B.C | pEPYC0CM0281 | PROM | 35s(ΔC2C2dof+$_1$) | MoClo, Loop, GB | GGAG | TACT | This study |
| T.B.C | pEPYC0CM0280 | PROM | 35s(ΔC2H2+) | MoClo, Loop, GB | GGAG | TACT | This study |
| T.B.C | pEPYC0CM0285 | PROM | 35s(ΔDBP) | MoClo, Loop, GB | GGAG | TACT | This study |
| T.B.C | pEPYC0CM0279 | PROM | 35s(ΔHSF+) | MoClo, Loop, GB | GGAG | TACT | This study |
| T.B.C | pEPYC0CM0277 | PROM | 35s(ΔMADS+$_1$) | MoClo, Loop, GB | GGAG | TACT | This study |
| T.B.C | pEPYC0CM0282 | PROM | 35s(ΔRWPRK$_1$) | MoClo, Loop, GB | GGAG | TACT | This study |
| T.B.C | pEPYC0CM0283 | PROM | 35s(ΔRWPRK$_2$) | MoClo, Loop, GB | GGAG | TACT | This study |
| T.B.C | pEPYC0CM0278 | PROM | 35s(ΔTCP$_1$) | MoClo, Loop, GB | GGAG | TACT | This study |
| T.B.C | pEPYC0CM0284 | PROM | 35s(ΔWRKY) | MoClo, Loop, GB | GGAG | TACT | This study |
| T.B.C | pEPYC0CM0288 | PROM | MMV(ΔG2like) | MoClo, Loop, GB | GGAG | TACT | This study |
| T.B.C | pEPYC0CM0292 | PROM | MMV(ΔCCAAT+) | MoClo, Loop, GB | GGAG | TACT | This study |
| T.B.C | pEPYC0CM0174 | PROM | MMV(ΔC2C2dof+$_2$) | MoClo, Loop, GB | GGAG | TACT | This study |
| T.B.C | pEPYC0CM0290 | PROM | MMV(ΔbHLH+) | MoClo, Loop, GB | GGAG | TACT | This study |
| T.B.C | pEPYC0CM0289 | PROM | MMV(ΔbHLH$_2$) | MoClo, Loop, GB | GGAG | TACT | This study |
| T.B.C | pEPYC0CM0175 | PROM | MMV(ΔC_CRE) | MoClo, Loop, GB | GGAG | TACT | This study |
| T.B.C | pEPYC0CM0286 | PROM | MMV(ΔMADS+$_2$) | MoClo, Loop, GB | GGAG | TACT | This study |
| T.B.C | pEPYC0CM0172 | PROM | MMV(ΔMYB+) | MoClo, Loop, GB | GGAG | TACT | This study |
| T.B.C | pEPYC0CM0291 | PROM | MMV(ΔWRKY+) | MoClo, Loop, GB | GGAG | TACT | This study |
| T.B.C | pEPYC0CM0294 | PROM | NOS(ΔAP2+$_2$) | MoClo, Loop, GB | GGAG | TACT | This study |
| T.B.C | pEPYC0CM0178 | PROM | NOS(ΔC_CRE) | MoClo, Loop, GB | GGAG | TACT | This study |
| T.B.C | pEPYC0CM0177 | PROM | NOS(ΔC2H2) | MoClo, Loop, GB | GGAG | TACT | This study |
| T.B.C | pEPYC0CM0293 | PROM | NOS(ΔG2like+) | MoClo, Loop, GB | GGAG | TACT | This study |
| T.B.C | pEPYC0CM0176 | PROM | NOS(Δhomeobox) | MoClo, Loop, GB | GGAG | TACT | This study |
| T.B.C | pEPYC0CM0173 | PROM | MMV(ΔTCP$_2$) | MoClo, Loop, GB | GGAG | TACT | This study |
| T.B.C | pEPYC0CM0287 | PROM | MMV(ΔTCP$_{3)}$ | MoClo, Loop, GB | GGAG | TACT | This study |
| T.B.C | pEPYC0CM0035 | PROM | MinSyn_000 | MoClo, Loop, GB | GGAG | TACT | This study |
| T.B.C | pEPYC0CM0366 | PROM | MinSyn_001 | MoClo, Loop, GB | GGAG | TACT | This study |
| T.B.C | pEPYC0CM0367 | PROM | MinSyn_002 | MoClo, Loop, GB | GGAG | TACT | This study |
| T.B.C | pEPYC0CM0368 | PROM | MinSyn_003 | MoClo, Loop, GB | GGAG | TACT | This study |
| T.B.C | pEPYC0CM0379 | PROM | MinSyn_004 | MoClo, Loop, GB | GGAG | TACT | This study |
| T.B.C | pEPYC0CM0380 | PROM | MinSyn_005 | MoClo, Loop, GB | GGAG | TACT | This study |
| T.B.C | pEPYC0CM0381 | PROM | MinSyn_006 | MoClo, Loop, GB | GGAG | TACT | This study |
| T.B.C | pEPYC0CM0372 | PROM | MinSyn_007 | MoClo, Loop, GB | GGAG | TACT | This study |
| T.B.C | pEPYC0CM0373 | PROM | MinSyn_008 | MoClo, Loop, GB | GGAG | TACT | This study |
| T.B.C | pEPYC0CM0374 | PROM | MinSyn_009 | MoClo, Loop, GB | GGAG | TACT | This study |
| T.B.C | pEPYC0CM0385 | PROM | MinSyn_010 | MoClo, Loop, GB | GGAG | TACT | This study |
| T.B.C | pEPYC0CM0386 | PROM | MinSyn_011 | MoClo, Loop, GB | GGAG | TACT | This study |
| T.B.C | pEPYC0CM0387 | PROM | MinSyn_012 | MoClo, Loop, GB | GGAG | TACT | This study |
| T.B.C | pEPYC0CM0390 | PROM | MinSyn_013 | MoClo, Loop, GB | GGAG | TACT | This study |
| T.B.C | pEPYC0CM0391 | PROM | MinSyn_014 | MoClo, Loop, GB | GGAG | TACT | This study |
| T.B.C | pEPYC0CM0392 | PROM | MinSyn_015 | MoClo, Loop, GB | GGAG | TACT | This study |
| T.B.C | pEPYC0CM0375 | PROM | MinSyn_016 | MoClo, Loop, GB | GGAG | TACT | This study |
| T.B.C | pEPYC0CM0376 | PROM | MinSyn_017 | MoClo, Loop, GB | GGAG | TACT | This study |
| T.B.C | pEPYC0CM0377 | PROM | MinSyn_018 | MoClo, Loop, GB | GGAG | TACT | This study |
| T.B.C | pEPYC0CM0382 | PROM | MinSyn_019 | MoClo, Loop, GB | GGAG | TACT | This study |
| T.B.C | pEPYC0CM0383 | PROM | MinSyn_020 | MoClo, Loop, GB | GGAG | TACT | This study |
| T.B.C | pEPYC0CM0384 | PROM | MinSyn_021 | MoClo, Loop, GB | GGAG | TACT | This study |
| T.B.C | pEPYC0CM0393 | PROM | MinSyn_022 | MoClo, Loop, GB | GGAG | TACT | This study |
| T.B.C | pEPYC0CM0365 | PROM | MinSyn_023 | MoClo, Loop, GB | GGAG | TACT | This study |
| T.B.C | pEPYC0CM0431 | PROM | MinSyn_024 | MoClo, Loop, GB | GGAG | TACT | This study |
| T.B.C | pEPYC0CM0435 | PROM | MinSyn_026 | MoClo, Loop, GB | GGAG | TACT | This study |
| T.B.C | pEPYC0CM0433 | PROM | MinSyn_027 | MoClo, Loop, GB | GGAG | TACT | This study |
| T.B.C | pEPYC0CM0437 | PROM | MinSyn_028 | MoClo, Loop, GB | GGAG | TACT | This study |
| T.B.C | pEPYC0CM0439 | PROM | MinSyn_029 | MoClo, Loop, GB | GGAG | TACT | This study |
| T.B.C | pEPYC0CM0434 | PROM | MinSyn_030 | MoClo, Loop, GB | GGAG | TACT | This study |
| T.B.C | pEPYC0CM0388 | PROM | MinSyn_031 | MoClo, Loop, GB | GGAG | TACT | This study |
| T.B.C | pEPYC0CM0440 | PROM | MinSyn_032 | MoClo, Loop, GB | GGAG | TACT | This study |
| T.B.C | pEPYC0CM0041 | PROM | MinSyn_033 | MoClo, Loop, GB | GGAG | TACT | This study |
| T.B.C | pEPYC0CM0310 | PROM | MinSyn_034 | MoClo, Loop, GB | GGAG | TACT | This study |
| T.B.C | pEPYC0CM0312 | PROM | MinSyn_035 | MoClo, Loop, GB | GGAG | TACT | This study |
| T.B.C | pEPYC0CM0313 | PROM | MinSyn_036 | MoClo, Loop, GB | GGAG | TACT | This study |
| T.B.C | pEPYC0CM0314 | PROM | MinSyn_037 | MoClo, Loop, GB | GGAG | TACT | This study |
| T.B.C | pEPYC0CM0225 | PROM | MinSyn_038 | MoClo, Loop, GB | GGAG | TACT | This study |
| T.B.C | pEPYC0CM0209 | PROM | MinSyn_039 | MoClo, Loop, GB | GGAG | TACT | This study |

| 1. Level 0 Phytobricks | | | | | | | |
|---|---|---|---|---|---|---|---|
| Addgene # | Plasmid code | Part type | Description/Origin | Compatibility with Assembly Systems | Cloning overhang (top strand) | | Source of plasmid |
| | | | | | 5' | 3' | |
| T.B.C | pEPYC0CM0210 | PROM | MinSyn_040 | MoClo, Loop, GB | GGAG | TACT | This study |
| T.B.C | pEPYC0CM0211 | PROM | MinSyn_041 | MoClo, Loop, GB | GGAG | TACT | This study |
| T.B.C | pEPYC0CM0311 | PROM | MinSyn_042 | MoClo, Loop, GB | GGAG | TACT | This study |
| T.B.C | pEPYC0CM0212 | PROM | MinSyn_043 | MoClo, Loop, GB | GGAG | TACT | This study |
| T.B.C | pEPYC0CM0213 | PROM | MinSyn_044 | MoClo, Loop, GB | GGAG | TACT | This study |
| T.B.C | pEPYC0CM0214 | PROM | MinSyn_045 | MoClo, Loop, GB | GGAG | TACT | This study |
| T.B.C | pEPYC0CM0182 | PROM | MinSyn_046 | MoClo, Loop, GB | GGAG | TACT | This study |
| T.B.C | pEPYC0CM0310 | PROM | MinSyn_047 | MoClo, Loop, GB | GGAG | TACT | This study |
| T.B.C | pEPYC0CM0395 | PROM | MinSyn_048 | MoClo, Loop, GB | GGAG | TACT | This study |
| T.B.C | pEPYC0CM0396 | PROM | MinSyn_049 | MoClo, Loop, GB | GGAG | TACT | This study |
| T.B.C | pEPYC0CM0451 | PROM | MinSyn_050 | MoClo, Loop, GB | GGAG | TACT | This study |
| T.B.C | pEPYC0CM0452 | PROM | MinSyn_051 | MoClo, Loop, GB | GGAG | TACT | This study |
| T.B.C | pEPYC0CM0183 | PROM | MinSyn_052 | MoClo, Loop, GB | GGAG | TACT | This study |
| T.B.C | pEPYC0CM0225 | PROM | MinSyn_053 | MoClo, Loop, GB | GGAG | TACT | This study |
| T.B.C | pEPYC0CM0397 | PROM | MinSyn_054 | MoClo, Loop, GB | GGAG | TACT | This study |
| T.B.C | pEPYC0CM0398 | PROM | MinSyn_055 | MoClo, Loop, GB | GGAG | TACT | This study |
| T.B.C | pEPYC0CM0453 | PROM | MinSyn_056 | MoClo, Loop, GB | GGAG | TACT | This study |
| T.B.C | pEPYC0CM0454 | PROM | MinSyn_057 | MoClo, Loop, GB | GGAG | TACT | This study |
| T.B.C | pEPYC0CM0184 | PROM | MinSyn_058 | MoClo, Loop, GB | GGAG | TACT | This study |
| T.B.C | pEPYC0CM0311 | PROM | MinSyn_059 | MoClo, Loop, GB | GGAG | TACT | This study |
| T.B.C | pEPYC0CM0401 | PROM | MinSyn_060 | MoClo, Loop, GB | GGAG | TACT | This study |
| T.B.C | pEPYC0CM0402 | PROM | MinSyn_061 | MoClo, Loop, GB | GGAG | TACT | This study |
| T.B.C | pEPYC0CM0455 | PROM | MinSyn_062 | MoClo, Loop, GB | GGAG | TACT | This study |
| T.B.C | pEPYC0CM0456 | PROM | MinSyn_063 | MoClo, Loop, GB | GGAG | TACT | This study |
| T.B.C | pEPAS0CMR0015 | PROM | MinSyn_064 | MoClo, Loop, GB | GGAG | AATG | This study |
| T.B.C | pEPAS0CMR0016 | PROM | MinSyn_065 | MoClo, Loop, GB | GGAG | AATG | This study |
| T.B.C | pEPKK0CM0107 | PROM | MinSyn_066 | MoClo, Loop, GB | GGAG | AATG | This study |
| T.B.C | pEPKK0CM0065 | PROM | MinSyn_067 | MoClo, Loop, GB | GGAG | AATG | This study |
| T.B.C | pEPYC0CM0407 | PROM | MinSyn_101 | MoClo, Loop, GB | GGAG | TACT | This study |
| T.B.C | pEPYC0CM0408 | PROM | MinSyn_102 | MoClo, Loop, GB | GGAG | TACT | This study |
| T.B.C | pEPYC0CM0409 | PROM | MinSyn_103 | MoClo, Loop, GB | GGAG | TACT | This study |
| T.B.C | pEPYC0CM0410 | PROM | MinSyn_104 | MoClo, Loop, GB | GGAG | TACT | This study |
| T.B.C | pEPYC0CM0411 | PROM | MinSyn_105 | MoClo, Loop, GB | GGAG | TACT | This study |
| T.B.C | pEPYC0CM0412 | PROM | MinSyn_106 | MoClo, Loop, GB | GGAG | TACT | This study |
| T.B.C | pEPYC0CM0413 | PROM | MinSyn_107 | MoClo, Loop, GB | GGAG | TACT | This study |
| T.B.C | pEPYC0CM0414 | PROM | MinSyn_108 | MoClo, Loop, GB | GGAG | TACT | This study |
| T.B.C | pEPYC0CM0415 | PROM | MinSyn_109 | MoClo, Loop, GB | GGAG | TACT | This study |
| T.B.C | pEPYC0CM0416 | PROM | MinSyn_110 | MoClo, Loop, GB | GGAG | TACT | This study |
| T.B.C | pEPYC0CM0417 | PROM | MinSyn_111 | MoClo, Loop, GB | GGAG | TACT | This study |
| T.B.C | pEPYC0CM0418 | PROM | MinSyn_112 | MoClo, Loop, GB | GGAG | TACT | This study |
| T.B.C | pEPYC0CM0419 | PROM | MinSyn_113 | MoClo, Loop, GB | GGAG | TACT | This study |
| T.B.C | pEPYC0CM0420 | PROM | MinSyn_114 | MoClo, Loop, GB | GGAG | TACT | This study |
| T.B.C | pEPYC0CM0421 | PROM | MinSyn_115 | MoClo, Loop, GB | GGAG | TACT | This study |
| T.B.C | pEPYC0CM0422 | PROM | MinSyn_116 | MoClo, Loop, GB | GGAG | TACT | This study |
| T.B.C | pEPYC0CM0423 | PROM | MinSyn_117 | MoClo, Loop, GB | GGAG | TACT | This study |
| T.B.C | pEPYC0CM0424 | PROM | MinSyn_118 | MoClo, Loop, GB | GGAG | TACT | This study |
| T.B.C | pEPYC0CM0425 | PROM | MinSyn_119 | MoClo, Loop, GB | GGAG | TACT | This study |
| T.B.C | pEPYC0CM0426 | PROM | MinSyn_120 | MoClo, Loop, GB | GGAG | TACT | This study |
| T.B.C | pEPYC0CM0427 | PROM | MinSyn_121 | MoClo, Loop, GB | GGAG | TACT | This study |
| T.B.C | pEPYC0CM0428 | PROM | MinSyn_122 | MoClo, Loop, GB | GGAG | TACT | This study |
| T.B.C | pEPYC0CM0429 | PROM | MinSyn_123 | MoClo, Loop, GB | GGAG | TACT | This study |
| T.B.C | pEPYC0CM0430 | PROM | MinSyn_124 | MoClo, Loop, GB | GGAG | TACT | This study |
| T.B.C | pEPYC0CM0244 | PROM | MinSyn_285 | MoClo, Loop, GB | GGAG | TACT | This study |
| T.B.C | pEPKK0CM0063 | DIST | MinSyn_069 | MoClo, Loop, GB | GGAG | TGAC | This study |
| T.B.C | pEPKK0CM0062 | DIST+PROX+CORE | MinSyn_068 | MoClo, Loop, GB | GGAG | CCAT | This study |
| T.B.C | pEPKK0CM0064 | PROX+CORE | MinSyn_069 | MoClo, Loop, GB | TGAC | CCAT | This study |
| T.B.C | pEPKK0CM0111 | NTAG | MinSyn_068 and MinSyn_069 | MoClo, Loop, GB | CCAT | AATG | This study |
| T.B.C | pICSL20002 | 5UTR | CPMV (CowPea Mosaic Virus) | MoClo, Loop, GB | TACT | CCAT | This study |
| #50285 | pICH41402 | 5UTR | TMVΩ (Tobacco Mosaic Virus) | MoClo, Loop, GB | TACT | AATG | Engler et al(2014) |
| T.B.C | pEPYC0CM0258 | NTAG | HiBit | MoClo, Loop, GB | CCAT | AATG | This study |
| - | GB0900 | NTAG | GAL4 activation domain | MoClo, Loop, GB | CCAT | AATG | Vazquez-Vilar et al (2017) |
| T.B.C | pEPAS0CM0008 | CDS | LucF | MoClo, Loop, GB | AATG | TTCG | This study |
| T.B.C | pEPYC0CM0133 | CDS | LucN | MoClo, Loop, GB | AATG | TTCG | This study |
| #50332 | pICSL80016 | CDS | uidA (GUS) | MoClo, Loop, GB | AATG | TTCG | Engler et al(2014) |
| - | GB_UD_32AB | CDS | PhiC3 binding domain | GB | AATG | GCTT | Vazquez-Vilar et al (2017) |
| T.B.C | pEPKK0CM0068 | CDS | TALE | MoClo, Loop, GB | AATG | GCTT | This study |
| #68260 | pICSL80037 | CDS | NptII | MoClo, Loop, GB | AATG | GCTT | Lawrenson et al (2015) |
| #50308 | pICSL50007 | CTAG | FLAG tag | MoClo, Loop, GB | TTCG | GCTT | Engler et al(2014) |
| #117536 | pICSL50005 | CTAG | YFP | MoClo, Loop, GB | TTCG | GCTT | Raitskin et al (2019) |
| #50343 | pICH41432 | 3UTR+TERM | OCS (Agrobacterium tumefaciens) | MoClo, Loop, GB | GCTT | CGCT | Engler et al(2014) |
| #50339 | pICH41421 | 3UTR+TERM | NOS (Agrobacterium tumefaciens) | MoClo, Loop, GB | GCTT | CGCT | Engler et al(2014) |
| #50338 | pICH72400 | 3UTR+TERM | g7 (Agrobacterium tumefaciens) | MoClo, Loop, GB | GCTT | CGCT | Engler et al(2014) |
| #50337 | pICH41414 | 3UTR+TERM | 35s (Cauliflower Mosaic Virus) | MoClo, Loop, GB | GCTT | CGCT | Engler et al(2014) |
| #68187 | GB0036 | 3UTR+TERM | 35s (Cauliflower Mosaic Virus) | MoClo, Loop, GB | GCTT | CGCT | Vazquez-Vilar et al (2017) |

| 2. Expression cassettes | | | | | | | | | | | | | |
|---|---|---|---|---|---|---|---|---|---|---|---|---|---|
| | | Level 0 Parts | | | | | | | | | Cloning overhang | | |
| | | PROM | | | | | | | | | | | |
| Addgene # | Plasmid code | DIST | PROX | CORE | 5UTR | NTAG | CDS | CTAG | 3'UTR/TERM | Acceptor | 5' | 3' | Source |
| T.B.C. | pEPYC1CB0308 | | pICSL12006 | | - | - | pUAP80037 | - | pICH41414 | pICH47732 | TGCC | GCAA | This study |
| T.B.C. | pEPYC1CB0305 | | pEPYC0CM0244 | | pICSL20002 | pEPYC0CM0258 | pICSL80016 | pICSL50005 | pICH41432 | pICH47811 | TAGT | TTGC | This study |
| T.B.C. | pEPYC1CB0477 | | pEPYC0CM0410 | | pICSL20002 | pEPYC0CM0258 | pICSL80016 | pICSL50005 | pICH41432 | pICH47811 | TAGT | TTGC | This study |
| T.B.C. | pEPYC1CB0478 | | pEPYC0CM0414 | | pICSL20002 | pEPYC0CM0258 | pICSL80016 | pICSL50005 | pICH41432 | pICH47811 | TAGT | TTGC | This study |
| T.B.C. | pEPYC1CB0503 | | pEPYC0CM0035 | | pICSL20002 | pEPYC0CM0258 | pICSL80016 | pICSL50005 | pICH41432 | pICH47811 | TAGT | TTGC | This study |
| T.B.C. | pEPYCα1KN0002 | | pEPYC0CM0414 | | pICSL20002 | GB0900 | GB_UD_32AB | - | pICH41432 | pDGB3_α1 | GGAG | GTCA | This study |
| T.B.C. | pEPYCα1KN0007 | | pEPYC0CM0414 | | pICH41402 | - | pEPKK0CM0068 | - | pICH41432 | pDGB3_α1 | GGAG | GTCA | This study |
| T.B.C. | GB_UA_114A | | GB0552 | | - | GB0900 | GB_UD_32AB | - | GB0036 | pDGB3_α1 | GGAG | GTCA | This study |
| T.B.C. | pEPYCα1KN0008 | | pEPKK0CM0107 | | - | - | pEPAS0CM0008 | pICSL50007 | pICH72400 | pDGB3_α1 | GGAG | GTCA | This study |
| T.B.C. | pEPYCα2rKN0009 | | pEPKK0CM0107 | | - | - | pEPYC0CM0133 | pICSL50007 | pICH41421 | pDGB3_α2R | GTCA | GGAG | This study |
| T.B.C. | pEPYCα2rKN0010 | | pEPAS0CMR0015 | | - | - | pEPYC0CM0133 | pICSL50007 | pICH41421 | pDGB3_α2R | GTCA | GGAG | This study |
| T.B.C. | pEPKKα2KN0091 | | pEPAS0CMR0015 | | - | - | pEPYC0CM0133 | pICSL50007 | pICH72400 | pDGB3_α1 | GGAG | GTCA | This study |
| T.B.C. | pEPKKα2KN0092 | | pEPAS0CMR0016 | | - | - | pEPYC0CM0133 | pICSL50007 | pICH72400 | pDGB3_α1 | GGAG | GTCA | This study |
| T.B.C. | pEPKKα2KN0093 | | pEPKK0CM0107 | | - | - | pEPYC0CM0133 | pICSL50007 | pICH72400 | pDGB3_α1 | GGAG | GTCA | This study |
| T.B.C. | pEPKKα2KN0100 | | pEPKK0CM0065 | | - | - | pEPYC0CM0133 | pICSL50007 | pICH72400 | pDGB3_α1 | GGAG | GTCA | This study |
| T.B.C. | pEPKKα2KN0101 | | pEPKK0CM0062 | | | pEPKK0CM0111 | pEPYC0CM0133 | pICSL50007 | pICH72400 | pDGB3_α1 | GGAG | GTCA | This study |
| T.B.C. | pEPKKα2KN0102 | pEPKK0CM0063 | pEPKK0CM0064 | | | pEPKK0CM0111 | pEPYC0CM0133 | pICSL50007 | pICH72400 | pDGB3_α1 | GGAG | GTCA | This study |
| T.B.C. | pEPYC1CB0003 | | pICH42211 | | pICH41402 | - | pEPAS0CM0008 | pICSL50007 | pICH41432 | pICH47732 | TGCC | GCAA | This study |
| T.B.C. | pEPYC1CB0007 | | pEPYC0CM0035 | | pICH41402 | - | pEPAS0CM0008 | pICSL50007 | pICH41432 | pICH47732 | TGCC | GCAA | This study |
| T.B.C. | pEPYC1CB0079 | | pEPYC0CM0041 | | pICH41402 | - | pEPAS0CM0008 | pICSL50007 | pICH41432 | pICH47732 | TGCC | GCAA | This study |
| T.B.C. | pEPYC1CB0109 | | pEPYC0CM0071 | | pICH41402 | - | pEPAS0CM0008 | pICSL50007 | pICH41432 | pICH47732 | TGCC | GCAA | This study |
| T.B.C. | pEPYC1CB0122 | | pEPYC0CM0084 | | pICH41402 | - | pEPAS0CM0008 | pICSL50007 | pICH41432 | pICH47732 | TGCC | GCAA | This study |
| T.B.C. | pEPYC1CB0127 | | pEPYC0CM0089 | | pICH41402 | - | pEPAS0CM0008 | pICSL50007 | pICH41432 | pICH47732 | TGCC | GCAA | This study |
| T.B.C. | pEPYC1CB0133 | | pEPYC0CM0095 | | pICH41402 | - | pEPAS0CM0008 | pICSL50007 | pICH41432 | pICH47732 | TGCC | GCAA | This study |
| T.B.C. | pEPYC1CB0137 | | pEPYC0CM0099 | | pICH41402 | - | pEPAS0CM0008 | pICSL50007 | pICH41432 | pICH47732 | TGCC | GCAA | This study |
| T.B.C. | pEPYC1CB0153 | | pEPYC0CM0115 | | pICH41402 | - | pEPAS0CM0008 | pICSL50007 | pICH41432 | pICH47732 | TGCC | GCAA | This study |
| T.B.C. | pEPYC1CB0157 | | pEPYC0CM0119 | | pICH41402 | - | pEPAS0CM0008 | pICSL50007 | pICH41432 | pICH47732 | TGCC | GCAA | This study |
| T.B.C. | pEPYC1CB0158 | | pEPYC0CM0120 | | pICH41402 | - | pEPAS0CM0008 | pICSL50007 | pICH41432 | pICH47732 | TGCC | GCAA | This study |
| T.B.C. | pEPYC1CB0173 | | pEPYC0CM0168 | | pICH41402 | - | pEPAS0CM0008 | pICSL50007 | pICH41432 | pICH47732 | TGCC | GCAA | This study |
| T.B.C. | pEPYC1CB0174 | | pEPYC0CM0169 | | pICH41402 | - | pEPAS0CM0008 | pICSL50007 | pICH41432 | pICH47732 | TGCC | GCAA | This study |
| T.B.C. | pEPYC1CB0175 | | pEPYC0CM0170 | | pICH41402 | - | pEPAS0CM0008 | pICSL50007 | pICH41432 | pICH47732 | TGCC | GCAA | This study |
| T.B.C. | pEPYC1CB0176 | | pEPYC0CM0171 | | pICH41402 | - | pEPAS0CM0008 | pICSL50007 | pICH41432 | pICH47732 | TGCC | GCAA | This study |
| T.B.C. | pEPYC1CB0177 | | pEPYC0CM0172 | | pICH41402 | - | pEPAS0CM0008 | pICSL50007 | pICH41432 | pICH47732 | TGCC | GCAA | This study |
| T.B.C. | pEPYC1CB0178 | | pEPYC0CM0173 | | pICH41402 | - | pEPAS0CM0008 | pICSL50007 | pICH41432 | pICH47732 | TGCC | GCAA | This study |
| T.B.C. | pEPYC1CB0179 | | pEPYC0CM0174 | | pICH41402 | - | pEPAS0CM0008 | pICSL50007 | pICH41432 | pICH47732 | TGCC | GCAA | This study |
| T.B.C. | pEPYC1CB0180 | | pEPYC0CM0175 | | pICH41402 | - | pEPAS0CM0008 | pICSL50007 | pICH41432 | pICH47732 | TGCC | GCAA | This study |
| T.B.C. | pEPYC1CB0181 | | pEPYC0CM0176 | | pICH41402 | - | pEPAS0CM0008 | pICSL50007 | pICH41432 | pICH47732 | TGCC | GCAA | This study |
| T.B.C. | pEPYC1CB0182 | | pEPYC0CM0177 | | pICH41402 | - | pEPAS0CM0008 | pICSL50007 | pICH41432 | pICH47732 | TGCC | GCAA | This study |
| T.B.C. | pEPYC1CB0183 | | pEPYC0CM0178 | | pICH41402 | - | pEPAS0CM0008 | pICSL50007 | pICH41432 | pICH47732 | TGCC | GCAA | This study |
| T.B.C. | pEPYC1CB0187 | | pEPYC0CM0182 | | pICH41402 | - | pEPAS0CM0008 | pICSL50007 | pICH41432 | pICH47732 | TGCC | GCAA | This study |
| T.B.C. | pEPYC1CB0188 | | pEPYC0CM0183 | | pICH41402 | - | pEPAS0CM0008 | pICSL50007 | pICH41432 | pICH47732 | TGCC | GCAA | This study |
| T.B.C. | pEPYC1CB0189 | | pEPYC0CM0184 | | pICH41402 | - | pEPAS0CM0008 | pICSL50007 | pICH41432 | pICH47732 | TGCC | GCAA | This study |
| T.B.C. | pEPYC1CB0197 | | pICH42211 | | pICH41402 | - | pEPYC0CM0133 | pICSL50007 | pICH41421 | pICH47732 | TGCC | GCAA | This study |
| T.B.C. | pEPYC1CB0199 | | pICH85281 | | pICH41402 | - | pEPAS0CM0008 | pICSL50007 | pICH41432 | pICH47732 | TGCC | GCAA | This study |
| T.B.C. | pEPYC1CB0253 | | pEPYC0CM0209 | | pICH41402 | - | pEPAS0CM0008 | pICSL50007 | pICH41432 | pICH47732 | TGCC | GCAA | This study |
| T.B.C. | pEPYC1CB0254 | | pEPYC0CM0210 | | pICH41402 | - | pEPAS0CM0008 | pICSL50007 | pICH41432 | pICH47732 | TGCC | GCAA | This study |
| T.B.C. | pEPYC1CB0255 | | pEPYC0CM0211 | | pICH41402 | - | pEPAS0CM0008 | pICSL50007 | pICH41432 | pICH47732 | TGCC | GCAA | This study |
| T.B.C. | pEPYC1CB0256 | | pEPYC0CM0212 | | pICH41402 | - | pEPAS0CM0008 | pICSL50007 | pICH41432 | pICH47732 | TGCC | GCAA | This study |
| T.B.C. | pEPYC1CB0257 | | pEPYC0CM0213 | | pICH41402 | - | pEPAS0CM0008 | pICSL50007 | pICH41432 | pICH47732 | TGCC | GCAA | This study |
| T.B.C. | pEPYC1CB0258 | | pEPYC0CM0214 | | pICH41402 | - | pEPAS0CM0008 | pICSL50007 | pICH41432 | pICH47732 | TGCC | GCAA | This study |
| T.B.C. | pEPYC1CB0269 | | pEPYC0CM0225 | | pICH41402 | - | pEPAS0CM0008 | pICSL50007 | pICH41432 | pICH47732 | TGCC | GCAA | This study |
| T.B.C. | pEPYC1CB0286 | | pEPYC0CM0244 | | pICH41402 | - | pEPYC0CM0133 | pICSL50007 | pICH41421 | pICH47732 | TGCC | GCAA | This study |
| T.B.C. | pEPYC1CB0319 | | pEPYC0CM0277 | | pICH41402 | - | pEPAS0CM0008 | pICSL50007 | pICH41432 | pICH47732 | TGCC | GCAA | This study |
| T.B.C. | pEPYC1CB0320 | | pEPYC0CM0278 | | pICH41402 | - | pEPAS0CM0008 | pICSL50007 | pICH41432 | pICH47732 | TGCC | GCAA | This study |
| T.B.C. | pEPYC1CB0321 | | pEPYC0CM0279 | | pICH41402 | - | pEPAS0CM0008 | pICSL50007 | pICH41432 | pICH47732 | TGCC | GCAA | This study |
| T.B.C. | pEPYC1CB0322 | | pEPYC0CM0280 | | pICH41402 | - | pEPAS0CM0008 | pICSL50007 | pICH41432 | pICH47732 | TGCC | GCAA | This study |
| T.B.C. | pEPYC1CB0323 | | pEPYC0CM0281 | | pICH41402 | - | pEPAS0CM0008 | pICSL50007 | pICH41432 | pICH47732 | TGCC | GCAA | This study |
| T.B.C. | pEPYC1CB0324 | | pEPYC0CM0282 | | pICH41402 | - | pEPAS0CM0008 | pICSL50007 | pICH41432 | pICH47732 | TGCC | GCAA | This study |
| T.B.C. | pEPYC1CB0325 | | pEPYC0CM0283 | | pICH41402 | - | pEPAS0CM0008 | pICSL50007 | pICH41432 | pICH47732 | TGCC | GCAA | This study |
| T.B.C. | pEPYC1CB0326 | | pEPYC0CM0284 | | pICH41402 | - | pEPAS0CM0008 | pICSL50007 | pICH41432 | pICH47732 | TGCC | GCAA | This study |
| T.B.C. | pEPYC1CB0327 | | pEPYC0CM0285 | | pICH41402 | - | pEPAS0CM0008 | pICSL50007 | pICH41432 | pICH47732 | TGCC | GCAA | This study |
| T.B.C. | pEPYC1CB0328 | | pEPYC0CM0286 | | pICH41402 | - | pEPAS0CM0008 | pICSL50007 | pICH41432 | pICH47732 | TGCC | GCAA | This study |
| T.B.C. | pEPYC1CB0329 | | pEPYC0CM0287 | | pICH41402 | - | pEPAS0CM0008 | pICSL50007 | pICH41432 | pICH47732 | TGCC | GCAA | This study |
| T.B.C. | pEPYC1CB0330 | | pEPYC0CM0288 | | pICH41402 | - | pEPAS0CM0008 | pICSL50007 | pICH41432 | pICH47732 | TGCC | GCAA | This study |
| T.B.C. | pEPYC1CB0331 | | pEPYC0CM0289 | | pICH41402 | - | pEPAS0CM0008 | pICSL50007 | pICH41432 | pICH47732 | TGCC | GCAA | This study |
| T.B.C. | pEPYC1CB0332 | | pEPYC0CM0290 | | pICH41402 | - | pEPAS0CM0008 | pICSL50007 | pICH41432 | pICH47732 | TGCC | GCAA | This study |
| T.B.C. | pEPYC1CB0333 | | pEPYC0CM0291 | | pICH41402 | - | pEPAS0CM0008 | pICSL50007 | pICH41432 | pICH47732 | TGCC | GCAA | This study |
| T.B.C. | pEPYC1CB0334 | | pEPYC0CM0292 | | pICH41402 | - | pEPAS0CM0008 | pICSL50007 | pICH41432 | pICH47732 | TGCC | GCAA | This study |
| T.B.C. | pEPYC1CB0335 | | pEPYC0CM0293 | | pICH41402 | - | pEPAS0CM0008 | pICSL50007 | pICH41432 | pICH47732 | TGCC | GCAA | This study |
| T.B.C. | pEPYC1CB0336 | | pEPYC0CM0294 | | pICH41402 | - | pEPAS0CM0008 | pICSL50007 | pICH41432 | pICH47732 | TGCC | GCAA | This study |
| T.B.C. | pEPYC1CB0352 | | pEPYC0CM0310 | | pICH41402 | - | pEPAS0CM0008 | pICSL50007 | pICH41432 | pICH47732 | TGCC | GCAA | This study |
| T.B.C. | pEPYC1CB0353 | | pEPYC0CM0311 | | pICH41402 | - | pEPAS0CM0008 | pICSL50007 | pICH41432 | pICH47732 | TGCC | GCAA | This study |
| T.B.C. | pEPYC1CB0354 | | pEPYC0CM0312 | | pICH41402 | - | pEPAS0CM0008 | pICSL50007 | pICH41432 | pICH47732 | TGCC | GCAA | This study |
| T.B.C. | pEPYC1CB0355 | | pEPYC0CM0313 | | pICH41402 | - | pEPAS0CM0008 | pICSL50007 | pICH41432 | pICH47732 | TGCC | GCAA | This study |
| T.B.C. | pEPYC1CB0356 | | pEPYC0CM0314 | | pICH41402 | - | pEPAS0CM0008 | pICSL50007 | pICH41432 | pICH47732 | TGCC | GCAA | This study |
| T.B.C. | pEPYC1CB0425 | | pEPYC0CM0365 | | pICH41402 | - | pEPAS0CM0008 | pICSL50007 | pICH41432 | pICH47732 | TGCC | GCAA | This study |
| T.B.C. | pEPYC1CB0426 | | pEPYC0CM0366 | | pICH41402 | - | pEPAS0CM0008 | pICSL50007 | pICH41432 | pICH47732 | TGCC | GCAA | This study |
| T.B.C. | pEPYC1CB0427 | | pEPYC0CM0367 | | pICH41402 | - | pEPAS0CM0008 | pICSL50007 | pICH41432 | pICH47732 | TGCC | GCAA | This study |
| T.B.C. | pEPYC1CB0428 | | pEPYC0CM0368 | | pICH41402 | - | pEPAS0CM0008 | pICSL50007 | pICH41432 | pICH47732 | TGCC | GCAA | This study |
| T.B.C. | pEPYC1CB0432 | | pEPYC0CM0372 | | pICH41402 | - | pEPAS0CM0008 | pICSL50007 | pICH41432 | pICH47732 | TGCC | GCAA | This study |
| T.B.C. | pEPYC1CB0433 | | pEPYC0CM0373 | | pICH41402 | - | pEPAS0CM0008 | pICSL50007 | pICH41432 | pICH47732 | TGCC | GCAA | This study |
| T.B.C. | pEPYC1CB0434 | | pEPYC0CM0374 | | pICH41402 | - | pEPAS0CM0008 | pICSL50007 | pICH41432 | pICH47732 | TGCC | GCAA | This study |
| T.B.C. | pEPYC1CB0435 | | pEPYC0CM0375 | | pICH41402 | - | pEPAS0CM0008 | pICSL50007 | pICH41432 | pICH47732 | TGCC | GCAA | This study |
| T.B.C. | pEPYC1CB0436 | | pEPYC0CM0376 | | pICH41402 | - | pEPAS0CM0008 | pICSL50007 | pICH41432 | pICH47732 | TGCC | GCAA | This study |
| T.B.C. | pEPYC1CB0437 | | pEPYC0CM0377 | | pICH41402 | - | pEPAS0CM0008 | pICSL50007 | pICH41432 | pICH47732 | TGCC | GCAA | This study |
| T.B.C. | pEPYC1CB0439 | | pEPYC0CM0379 | | pICH41402 | - | pEPAS0CM0008 | pICSL50007 | pICH41432 | pICH47732 | TGCC | GCAA | This study |
| T.B.C. | pEPYC1CB0440 | | pEPYC0CM0380 | | pICH41402 | - | pEPAS0CM0008 | pICSL50007 | pICH41432 | pICH47732 | TGCC | GCAA | This study |
| T.B.C. | pEPYC1CB0441 | | pEPYC0CM0381 | | pICH41402 | - | pEPAS0CM0008 | pICSL50007 | pICH41432 | pICH47732 | TGCC | GCAA | This study |
| T.B.C. | pEPYC1CB0442 | | pEPYC0CM0382 | | pICH41402 | - | pEPAS0CM0008 | pICSL50007 | pICH41432 | pICH47732 | TGCC | GCAA | This study |
| T.B.C. | pEPYC1CB0443 | | pEPYC0CM0383 | | pICH41402 | - | pEPAS0CM0008 | pICSL50007 | pICH41432 | pICH47732 | TGCC | GCAA | This study |
| T.B.C. | pEPYC1CB0444 | | pEPYC0CM0384 | | pICH41402 | - | pEPAS0CM0008 | pICSL50007 | pICH41432 | pICH47732 | TGCC | GCAA | This study |
| T.B.C. | pEPYC1CB0445 | | pEPYC0CM0385 | | pICH41402 | - | pEPAS0CM0008 | pICSL50007 | pICH41432 | pICH47732 | TGCC | GCAA | This study |
| T.B.C. | pEPYC1CB0446 | | pEPYC0CM0386 | | pICH41402 | - | pEPAS0CM0008 | pICSL50007 | pICH41432 | pICH47732 | TGCC | GCAA | This study |
| T.B.C. | pEPYC1CB0447 | | pEPYC0CM0387 | | pICH41402 | - | pEPAS0CM0008 | pICSL50007 | pICH41432 | pICH47732 | TGCC | GCAA | This study |
| T.B.C. | pEPYC1CB0448 | | pEPYC0CM0388 | | pICH41402 | - | pEPAS0CM0008 | pICSL50007 | pICH41432 | pICH47732 | TGCC | GCAA | This study |
| T.B.C. | pEPYC1CB0450 | | pEPYC0CM0390 | | pICH41402 | - | pEPAS0CM0008 | pICSL50007 | pICH41432 | pICH47732 | TGCC | GCAA | This study |
| T.B.C. | pEPYC1CB0451 | | pEPYC0CM0391 | | pICH41402 | - | pEPAS0CM0008 | pICSL50007 | pICH41432 | pICH47732 | TGCC | GCAA | This study |
| T.B.C. | pEPYC1CB0452 | | pEPYC0CM0392 | | pICH41402 | - | pEPAS0CM0008 | pICSL50007 | pICH41432 | pICH47732 | TGCC | GCAA | This study |
| T.B.C. | pEPYC1CB0453 | | pEPYC0CM0393 | | pICH41402 | - | pEPAS0CM0008 | pICSL50007 | pICH41432 | pICH47732 | TGCC | GCAA | This study |
| T.B.C. | pEPYC1CB0455 | | pEPYC0CM0395 | | pICH41402 | - | pEPAS0CM0008 | pICSL50007 | pICH41432 | pICH47732 | TGCC | GCAA | This study |

| Addgene # | Plasmid code | Level 0 Parts | | | | | | | | Acceptor | Cloning overhang | | Source |
|---|---|---|---|---|---|---|---|---|---|---|---|---|---|
| | | PROM | | | 5UTR | NTAG | CDS | CTAG | 3'UTR/TERM | | 5' | 3' | |
| | | DIST | PROX | CORE | | | | | | | | | |
| T.B.C. | pEPYC1CB0456 | | pEPYC0CM0396 | | pICH41402 | - | pEPAS0CM0008 | pICSL50007 | pICH41432 | pICH47732 | TGCC | GCAA | This study |
| T.B.C. | pEPYC1CB0457 | | pEPYC0CM0397 | | pICH41402 | - | pEPAS0CM0008 | pICSL50007 | pICH41432 | pICH47732 | TGCC | GCAA | This study |
| T.B.C. | pEPYC1CB0458 | | pEPYC0CM0398 | | pICH41402 | - | pEPAS0CM0008 | pICSL50007 | pICH41432 | pICH47732 | TGCC | GCAA | This study |
| T.B.C. | pEPYC1CB0461 | | pEPYC0CM0401 | | pICH41402 | - | pEPAS0CM0008 | pICSL50007 | pICH41432 | pICH47732 | TGCC | GCAA | This study |
| T.B.C. | pEPYC1CB0462 | | pEPYC0CM0402 | | pICH41402 | - | pEPAS0CM0008 | pICSL50007 | pICH41432 | pICH47732 | TGCC | GCAA | This study |
| T.B.C. | pEPYC1CB0467 | | pEPYC0CM0407 | | pICH41402 | - | pEPAS0CM0008 | pICSL50007 | pICH41432 | pICH47732 | TGCC | GCAA | This study |
| T.B.C. | pEPYC1CB0468 | | pEPYC0CM0408 | | pICH41402 | - | pEPAS0CM0008 | pICSL50007 | pICH41432 | pICH47732 | TGCC | GCAA | This study |
| T.B.C. | pEPYC1CB0469 | | pEPYC0CM0409 | | pICH41402 | - | pEPAS0CM0008 | pICSL50007 | pICH41432 | pICH47732 | TGCC | GCAA | This study |
| T.B.C. | pEPYC1CB0470 | | pEPYC0CM0410 | | pICH41402 | - | pEPAS0CM0008 | pICSL50007 | pICH41432 | pICH47732 | TGCC | GCAA | This study |
| T.B.C. | pEPYC1CB0471 | | pEPYC0CM0411 | | pICH41402 | - | pEPAS0CM0008 | pICSL50007 | pICH41432 | pICH47732 | TGCC | GCAA | This study |
| T.B.C. | pEPYC1CB0472 | | pEPYC0CM0412 | | pICH41402 | - | pEPAS0CM0008 | pICSL50007 | pICH41432 | pICH47732 | TGCC | GCAA | This study |
| T.B.C. | pEPYC1CB0473 | | pEPYC0CM0413 | | pICH41402 | - | pEPAS0CM0008 | pICSL50007 | pICH41432 | pICH47732 | TGCC | GCAA | This study |
| T.B.C. | pEPYC1CB0474 | | pEPYC0CM0414 | | pICH41402 | - | pEPAS0CM0008 | pICSL50007 | pICH41432 | pICH47732 | TGCC | GCAA | This study |
| T.B.C. | pEPYC1CB0475 | | pEPYC0CM0415 | | pICH41402 | - | pEPAS0CM0008 | pICSL50007 | pICH41432 | pICH47732 | TGCC | GCAA | This study |
| T.B.C. | pEPYC1CB0476 | | pEPYC0CM0416 | | pICH41402 | - | pEPAS0CM0008 | pICSL50007 | pICH41432 | pICH47732 | TGCC | GCAA | This study |
| T.B.C. | pEPYC1CB0479 | | pEPYC0CM0417 | | pICH41402 | - | pEPAS0CM0008 | pICSL50007 | pICH41432 | pICH47732 | TGCC | GCAA | This study |
| T.B.C. | pEPYC1CB0480 | | pEPYC0CM0418 | | pICH41402 | - | pEPAS0CM0008 | pICSL50007 | pICH41432 | pICH47732 | TGCC | GCAA | This study |
| T.B.C. | pEPYC1CB0481 | | pEPYC0CM0419 | | pICH41402 | - | pEPAS0CM0008 | pICSL50007 | pICH41432 | pICH47732 | TGCC | GCAA | This study |
| T.B.C. | pEPYC1CB0482 | | pEPYC0CM0420 | | pICH41402 | - | pEPAS0CM0008 | pICSL50007 | pICH41432 | pICH47732 | TGCC | GCAA | This study |
| T.B.C. | pEPYC1CB0483 | | pEPYC0CM0421 | | pICH41402 | - | pEPAS0CM0008 | pICSL50007 | pICH41432 | pICH47732 | TGCC | GCAA | This study |
| T.B.C. | pEPYC1CB0484 | | pEPYC0CM0422 | | pICH41402 | - | pEPAS0CM0008 | pICSL50007 | pICH41432 | pICH47732 | TGCC | GCAA | This study |
| T.B.C. | pEPYC1CB0485 | | pEPYC0CM0423 | | pICH41402 | - | pEPAS0CM0008 | pICSL50007 | pICH41432 | pICH47732 | TGCC | GCAA | This study |
| T.B.C. | pEPYC1CB0486 | | pEPYC0CM0424 | | pICH41402 | - | pEPAS0CM0008 | pICSL50007 | pICH41432 | pICH47732 | TGCC | GCAA | This study |
| T.B.C. | pEPYC1CB0487 | | pEPYC0CM0425 | | pICH41402 | - | pEPAS0CM0008 | pICSL50007 | pICH41432 | pICH47732 | TGCC | GCAA | This study |
| T.B.C. | pEPYC1CB0488 | | pEPYC0CM0426 | | pICH41402 | - | pEPAS0CM0008 | pICSL50007 | pICH41432 | pICH47732 | TGCC | GCAA | This study |
| T.B.C. | pEPYC1CB0489 | | pEPYC0CM0427 | | pICH41402 | - | pEPAS0CM0008 | pICSL50007 | pICH41432 | pICH47732 | TGCC | GCAA | This study |
| T.B.C. | pEPYC1CB0490 | | pEPYC0CM0428 | | pICH41402 | - | pEPAS0CM0008 | pICSL50007 | pICH41432 | pICH47732 | TGCC | GCAA | This study |
| T.B.C. | pEPYC1CB0491 | | pEPYC0CM0429 | | pICH41402 | - | pEPAS0CM0008 | pICSL50007 | pICH41432 | pICH47732 | TGCC | GCAA | This study |
| T.B.C. | pEPYC1CB0492 | | pEPYC0CM0430 | | pICH41402 | - | pEPAS0CM0008 | pICSL50007 | pICH41432 | pICH47732 | TGCC | GCAA | This study |
| T.B.C. | pEPYC1CB0493 | | pEPYC0CM0431 | | pICH41402 | - | pEPAS0CM0008 | pICSL50007 | pICH41432 | pICH47732 | TGCC | GCAA | This study |
| T.B.C. | pEPYC1CB0495 | | pEPYC0CM0433 | | pICH41402 | - | pEPAS0CM0008 | pICSL50007 | pICH41432 | pICH47732 | TGCC | GCAA | This study |
| T.B.C. | pEPYC1CB0496 | | pEPYC0CM0434 | | pICH41402 | - | pEPAS0CM0008 | pICSL50007 | pICH41432 | pICH47732 | TGCC | GCAA | This study |
| T.B.C. | pEPYC1CB0497 | | pEPYC0CM0435 | | pICH41402 | - | pEPAS0CM0008 | pICSL50007 | pICH41432 | pICH47732 | TGCC | GCAA | This study |
| T.B.C. | pEPYC1CB0498 | | pEPYC0CM0436 | | pICH41402 | - | pEPAS0CM0008 | pICSL50007 | pICH41432 | pICH47732 | TGCC | GCAA | This study |
| T.B.C. | pEPYC1CB0499 | | pEPYC0CM0437 | | pICH41402 | - | pEPAS0CM0008 | pICSL50007 | pICH41432 | pICH47732 | TGCC | GCAA | This study |
| T.B.C. | pEPYC1CB0501 | | pEPYC0CM0439 | | pICH41402 | - | pEPAS0CM0008 | pICSL50007 | pICH41432 | pICH47732 | TGCC | GCAA | This study |
| T.B.C. | pEPYC1CB0502 | | pEPYC0CM0440 | | pICH41402 | - | pEPAS0CM0008 | pICSL50007 | pICH41432 | pICH47732 | TGCC | GCAA | This study |
| T.B.C. | pEPYC1CB0509 | | pEPYC0CM0451 | | pICH41402 | - | pEPAS0CM0008 | pICSL50007 | pICH41432 | pICH47732 | TGCC | GCAA | This study |
| T.B.C. | pEPYC1CB0510 | | pEPYC0CM0452 | | pICH41402 | - | pEPAS0CM0008 | pICSL50007 | pICH41432 | pICH47732 | TGCC | GCAA | This study |
| T.B.C. | pEPYC1CB0511 | | pEPYC0CM0453 | | pICH41402 | - | pEPAS0CM0008 | pICSL50007 | pICH41432 | pICH47732 | TGCC | GCAA | This study |
| T.B.C. | pEPYC1CB0512 | | pEPYC0CM0454 | | pICH41402 | - | pEPAS0CM0008 | pICSL50007 | pICH41432 | pICH47732 | TGCC | GCAA | This study |
| T.B.C. | pEPYC1CB0513 | | pEPYC0CM0455 | | pICH41402 | - | pEPAS0CM0008 | pICSL50007 | pICH41432 | pICH47732 | TGCC | GCAA | This study |
| T.B.C. | pEPYC1CB0514 | | pEPYC0CM0456 | | pICH41402 | - | pEPAS0CM0008 | pICSL50007 | pICH41432 | pICH47732 | TGCC | GCAA | This study |

**Supplementary Data 3**
Expression profiles of *Arabidopsis thaliana* genes encoding transcription factors predicted to bind to the following constitutive promoters:
**(1)** Cauliflower Mosaic Virus 35s (*CaMV35s*)
**(2)** Mirabilis Mosaic Virus (*MMV*)
**(3)** Agrobacterium tumefaciens *nopaline synthase* (*AtuNOS*)
**(4)** *A. thaliana* actin (*AtACT2*)
**(5)** *A. thaliana* ubiquitin-conjugating enzyme 9 (*AtUBC9*)
**(6)** *A. thaliana* polyubiquitin 10 (*AtUBI10*)
TPM = Transcripts per million

(1)

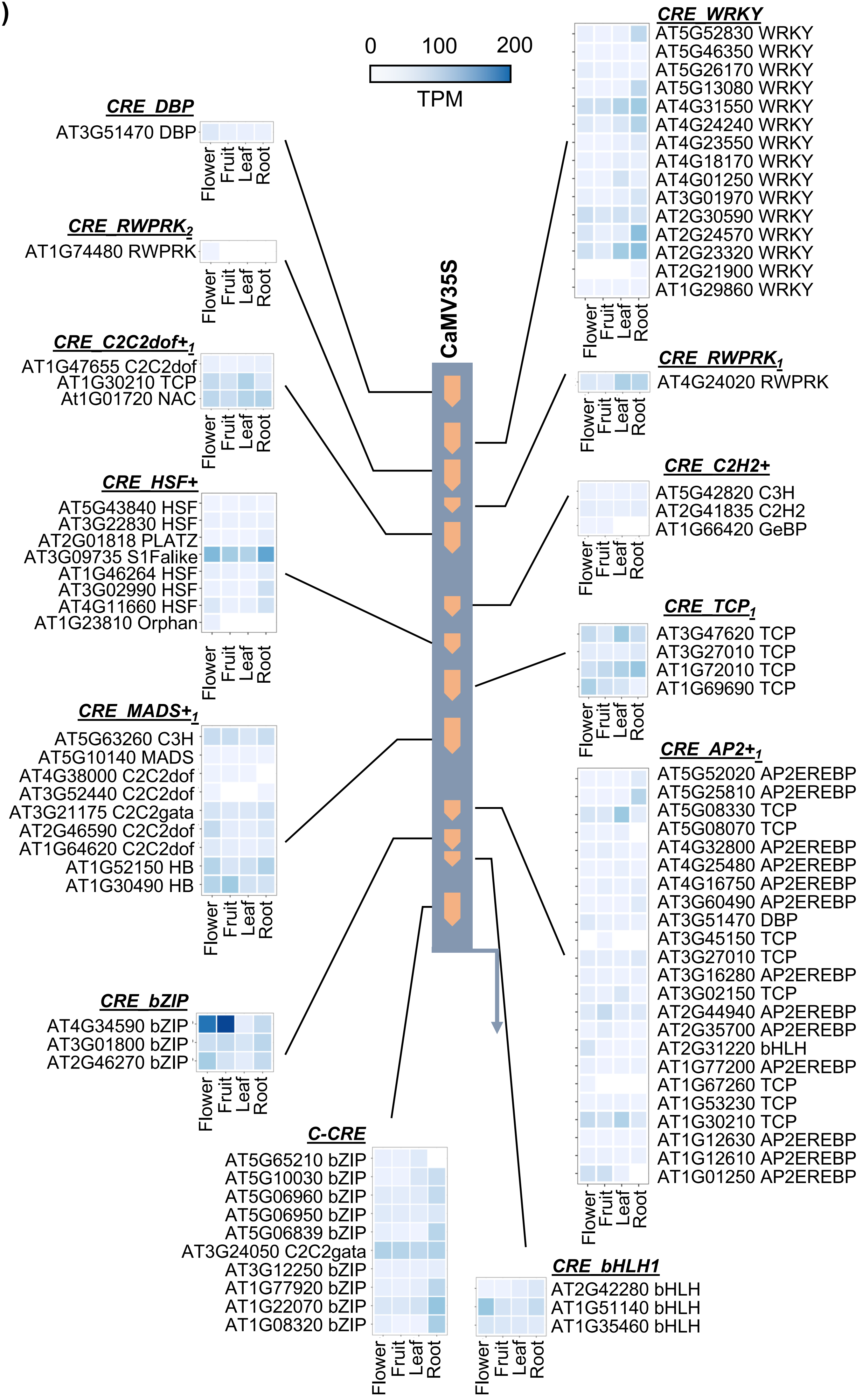

0 100 200
TPM
CaMV35S
CRE_DBP
AT3G51470 DBP
CRE_RWPRK2
AT1G74480 RWPRK
CRE_C2C2dof+1
AT1G47655 C2C2dof
AT1G30210 TCP
At1G01720 NAC
CRE_HSF+
AT5G43840 HSF
AT3G22830 HSF
AT2G01818 PLATZ
AT3G09735 S1Falike
AT1G46264 HSF
AT3G02990 HSF
AT4G11660 HSF
AT1G23810 Orphan
CRE_MADS+1
AT5G63260 C3H
AT5G10140 MADS
AT4G38000 C2C2dof
AT3G52440 C2C2dof
AT3G21175 C2C2gata
AT2G46590 C2C2dof
AT1G64620 C2C2dof
AT1G52150 HB
AT1G30490 HB
CRE_bZIP
AT4G34590 bZIP
AT3G01800 bZIP
AT2G46270 bZIP
C-CRE
AT5G65210 bZIP
AT5G10030 bZIP
AT5G06960 bZIP
AT5G06950 bZIP
AT5G06839 bZIP
AT3G24050 C2C2gata
AT3G12250 bZIP
AT1G77920 bZIP
AT1G22070 bZIP
AT1G08320 bZIP
CRE_WRKY
AT5G52830 WRKY
AT5G46350 WRKY
AT5G26170 WRKY
AT5G13080 WRKY
AT4G31550 WRKY
AT4G24240 WRKY
AT4G23550 WRKY
AT4G18170 WRKY
AT4G01250 WRKY
AT3G01970 WRKY
AT2G30590 WRKY
AT2G24570 WRKY
AT2G23320 WRKY
AT2G21900 WRKY
AT1G29860 WRKY
CRE_RWPRK1
AT4G24020 RWPRK
CRE_C2H2+
AT5G42820 C3H
AT2G41835 C2H2
AT1G66420 GeBP
CRE_TCP1
AT3G47620 TCP
AT3G27010 TCP
AT1G72010 TCP
AT1G69690 TCP
CRE_AP2+1
AT5G52020 AP2EREBP
AT5G25810 AP2EREBP
AT5G08330 TCP
AT5G08070 TCP
AT4G32800 AP2EREBP
AT4G25480 AP2EREBP
AT4G16750 AP2EREBP
AT3G60490 AP2EREBP
AT3G51470 DBP
AT3G45150 TCP
AT3G27010 TCP
AT3G16280 AP2EREBP
AT3G02150 TCP
AT2G44940 AP2EREBP
AT2G35700 AP2EREBP
AT2G31220 bHLH
AT1G77200 AP2EREBP
AT1G67260 TCP
AT1G53230 TCP
AT1G30210 TCP
AT1G12630 AP2EREBP
AT1G12610 AP2EREBP
AT1G01250 AP2EREBP
CRE_bHLH1
AT2G42280 bHLH
AT1G51140 bHLH
AT1G35460 bHLH
Flower
Fruit
Leaf
Root

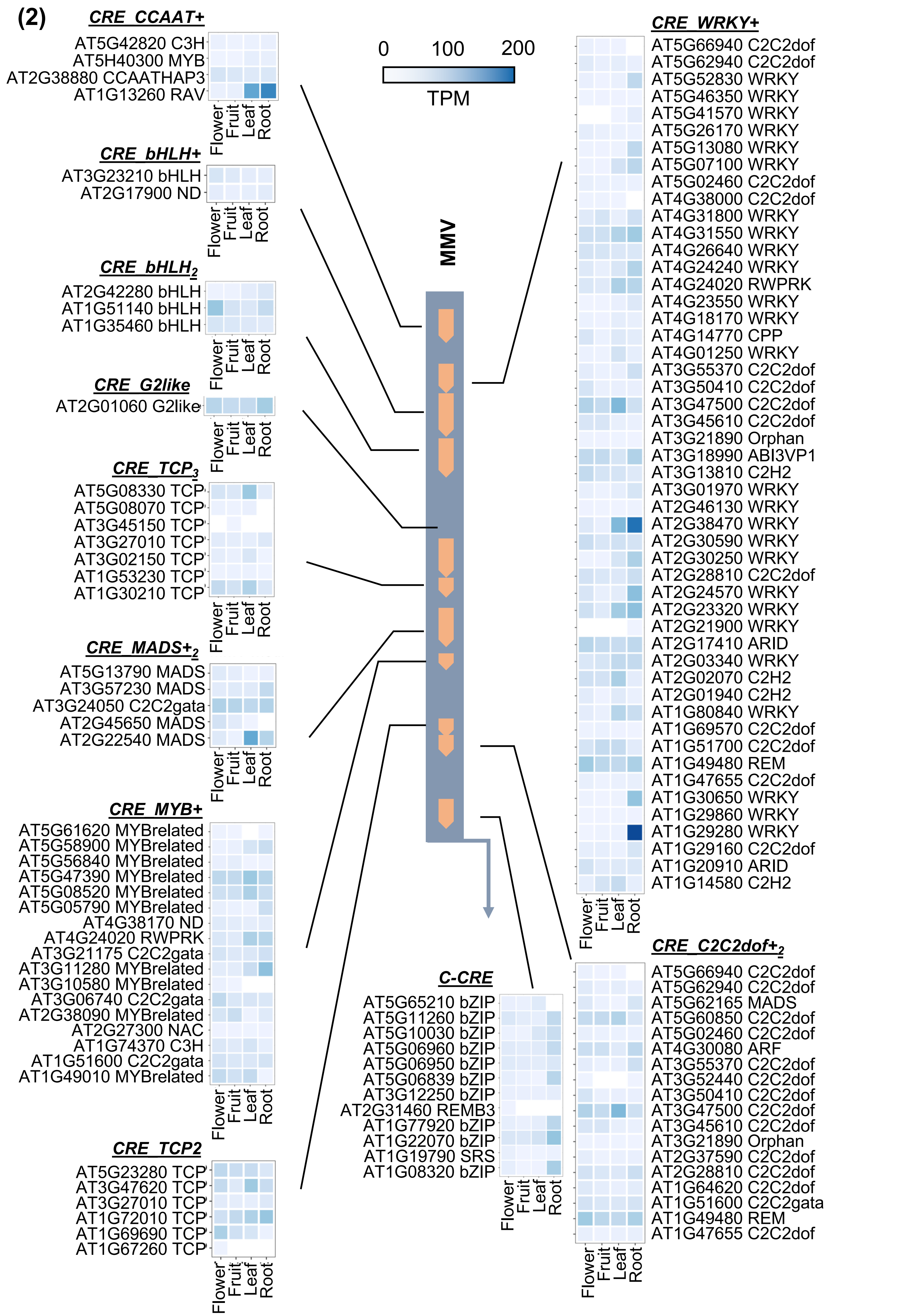

(2)
CRE_CCAAT+
AT5G42820 C3H
AT5H40300 MYB
AT2G38880 CCAATHAP3
AT1G13260 RAV
CRE_bHLH+
AT3G23210 bHLH
AT2G17900 ND
CRE_bHLH2
AT2G42280 bHLH
AT1G51140 bHLH
AT1G35460 bHLH
CRE_G2like
AT2G01060 G2like
CRE_TCP3
AT5G08330 TCP
AT5G08070 TCP
AT3G45150 TCP
AT3G27010 TCP
AT3G02150 TCP
AT1G53230 TCP
AT1G30210 TCP
CRE_MADS+2
AT5G13790 MADS
AT3G57230 MADS
AT3G24050 C2C2gata
AT2G45650 MADS
AT2G22540 MADS
CRE_MYB+
AT5G61620 MYBrelated
AT5G58900 MYBrelated
AT5G56840 MYBrelated
AT5G47390 MYBrelated
AT5G08520 MYBrelated
AT5G05790 MYBrelated
AT4G38170 ND
AT4G24020 RWPRK
AT3G21175 C2C2gata
AT3G11280 MYBrelated
AT3G10580 MYBrelated
AT3G06740 C2C2gata
AT2G38090 MYBrelated
AT2G27300 NAC
AT1G74370 C3H
AT1G51600 C2C2gata
AT1G49010 MYBrelated
CRE_TCP2
AT5G23280 TCP
AT3G47620 TCP
AT3G27010 TCP
AT1G72010 TCP
AT1G69690 TCP
AT1G67260 TCP
0 100 200
TPM
MMV
C-CRE
AT5G65210 bZIP
AT5G11260 bZIP
AT5G10030 bZIP
AT5G06960 bZIP
AT5G06950 bZIP
AT5G06839 bZIP
AT3G12250 bZIP
AT2G31460 REMB3
AT1G77920 bZIP
AT1G22070 bZIP
AT1G19790 SRS
AT1G08320 bZIP
CRE_WRKY+
AT5G66940 C2C2dof
AT5G62940 C2C2dof
AT5G52830 WRKY
AT5G46350 WRKY
AT5G41570 WRKY
AT5G26170 WRKY
AT5G13080 WRKY
AT5G07100 WRKY
AT5G02460 C2C2dof
AT4G38000 C2C2dof
AT4G31800 WRKY
AT4G31550 WRKY
AT4G26640 WRKY
AT4G24240 WRKY
AT4G24020 RWPRK
AT4G23550 WRKY
AT4G18170 WRKY
AT4G14770 CPP
AT4G01250 WRKY
AT3G55370 C2C2dof
AT3G50410 C2C2dof
AT3G47500 C2C2dof
AT3G45610 C2C2dof
AT3G21890 Orphan
AT3G18990 ABI3VP1
AT3G13810 C2H2
AT3G01970 WRKY
AT2G46130 WRKY
AT2G38470 WRKY
AT2G30590 WRKY
AT2G30250 WRKY
AT2G28810 C2C2dof
AT2G24570 WRKY
AT2G23320 WRKY
AT2G21900 WRKY
AT2G17410 ARID
AT2G03340 WRKY
AT2G02070 C2H2
AT2G01940 C2H2
AT1G80840 WRKY
AT1G69570 C2C2dof
AT1G51700 C2C2dof
AT1G49480 REM
AT1G47655 C2C2dof
AT1G30650 WRKY
AT1G29860 WRKY
AT1G29280 WRKY
AT1G29160 C2C2dof
AT1G20910 ARID
AT1G14580 C2H2
CRE_C2C2dof+2
AT5G66940 C2C2dof
AT5G62940 C2C2dof
AT5G62165 MADS
AT5G60850 C2C2dof
AT5G02460 C2C2dof
AT4G30080 ARF
AT3G55370 C2C2dof
AT3G52440 C2C2dof
AT3G50410 C2C2dof
AT3G47500 C2C2dof
AT3G45610 C2C2dof
AT3G21890 Orphan
AT2G37590 C2C2dof
AT2G28810 C2C2dof
AT1G64620 C2C2dof
AT1G51600 C2C2gata
AT1G49480 REM
AT1G47655 C2C2dof
Flower
Fruit
Leaf
Root

(3)

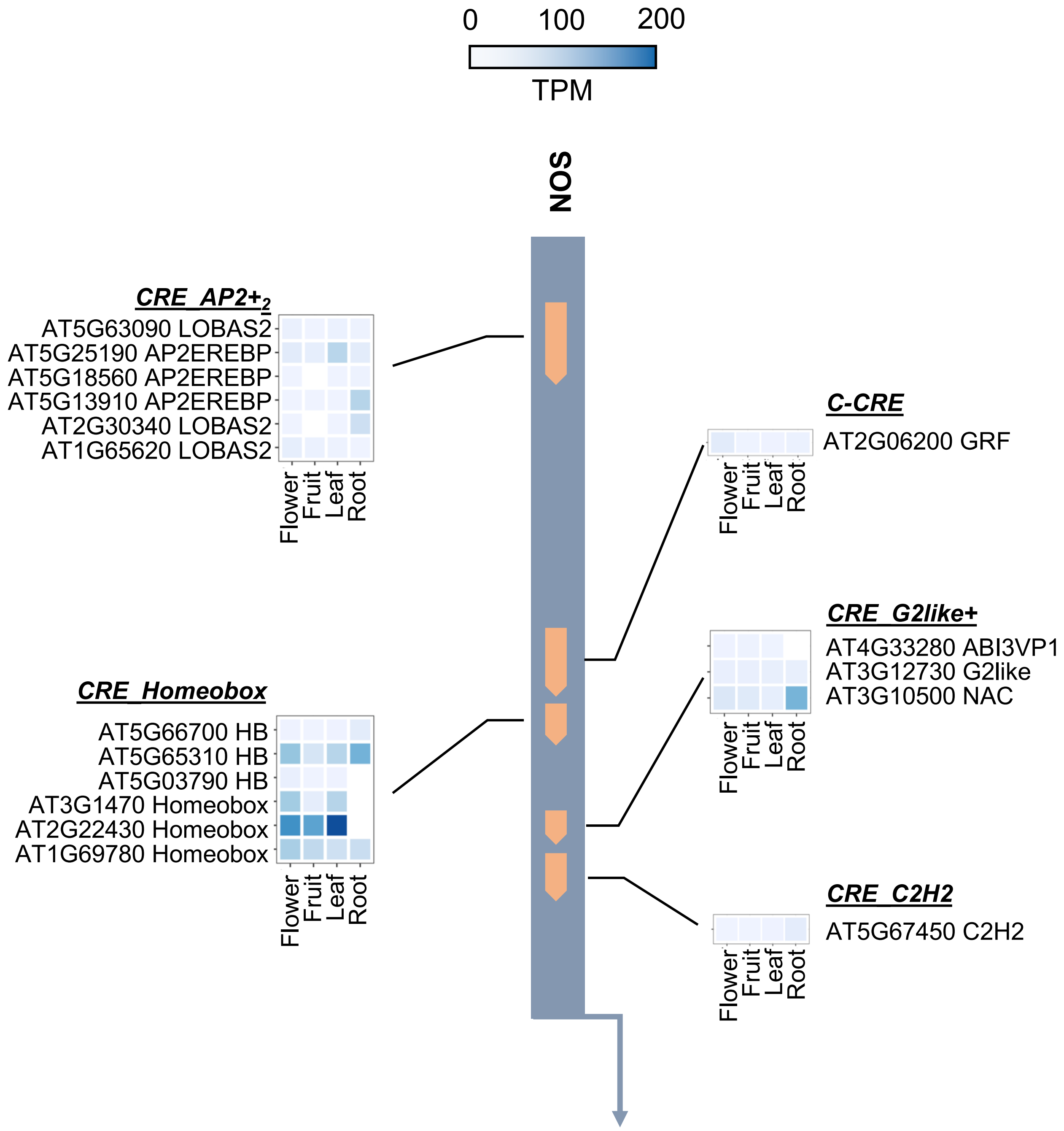

0
100
200
TPM
NOS
CRE_AP2+2
AT5G63090 LOBAS2
AT5G25190 AP2EREBP
AT5G18560 AP2EREBP
AT5G13910 AP2EREBP
AT2G30340 LOBAS2
AT1G65620 LOBAS2
Flower
Fruit
Leaf
Root
C-CRE
AT2G06200 GRF
Flower
Fruit
Leaf
Root
CRE_Homeobox
AT5G66700 HB
AT5G65310 HB
AT5G03790 HB
AT3G1470 Homeobox
AT2G22430 Homeobox
AT1G69780 Homeobox
Flower
Fruit
Leaf
Root
CRE_G2like+
AT4G33280 ABI3VP1
AT3G12730 G2like
AT3G10500 NAC
Flower
Fruit
Leaf
Root
CRE_C2H2
AT5G67450 C2H2
Flower
Fruit
Leaf
Root

(4)

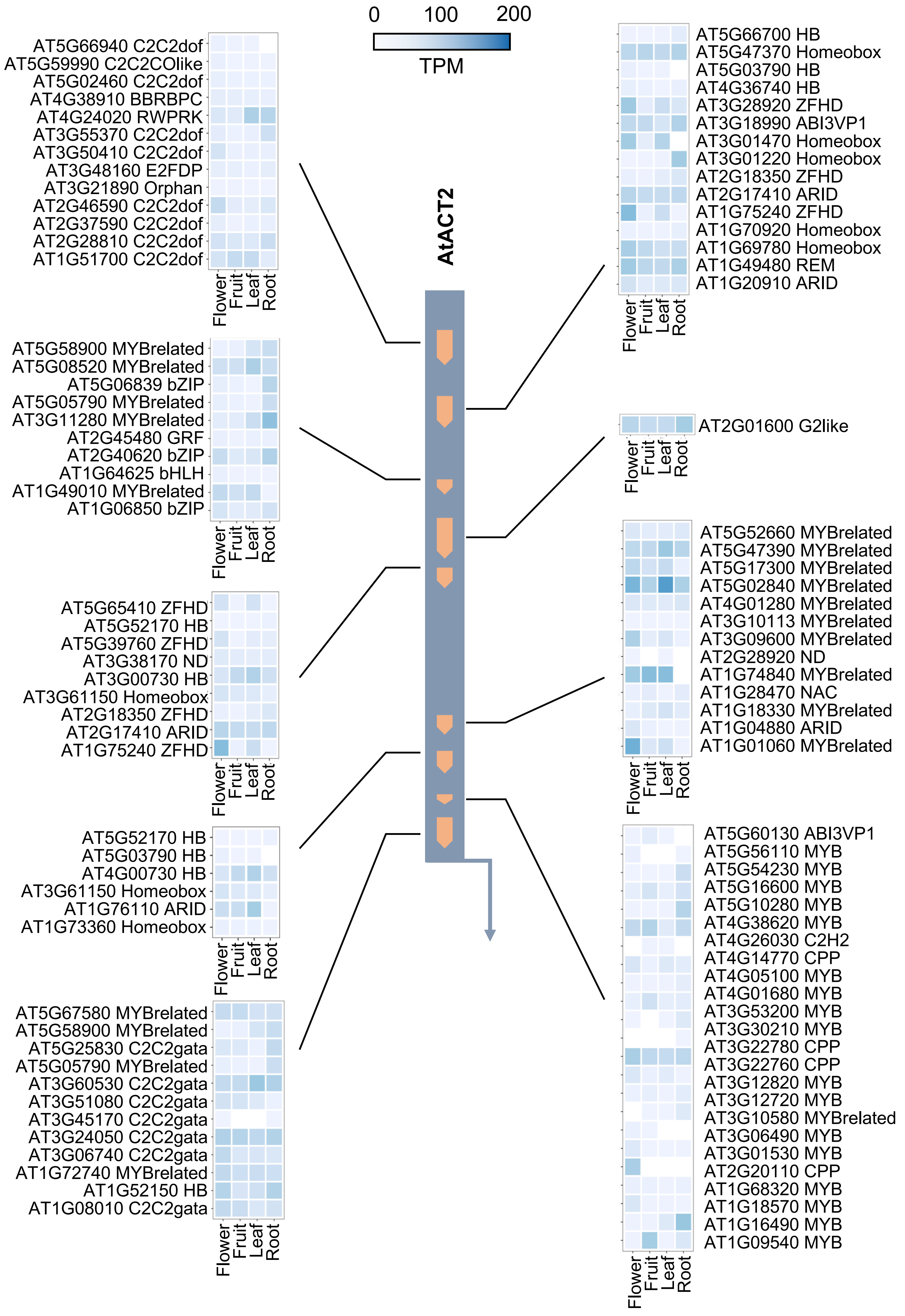
0
100
200
TPM
AtACT2
AT5G66940 C2C2dof
AT5G59990 C2C2COlike
AT5G02460 C2C2dof
AT4G38910 BBRBPC
AT4G24020 RWPRK
AT3G55370 C2C2dof
AT3G50410 C2C2dof
AT3G48160 E2FDP
AT3G21890 Orphan
AT2G46590 C2C2dof
AT2G37590 C2C2dof
AT2G28810 C2C2dof
AT1G51700 C2C2dof
Flower
Fruit
Leaf
Root
AT5G58900 MYBrelated
AT5G08520 MYBrelated
AT5G06839 bZIP
AT5G05790 MYBrelated
AT3G11280 MYBrelated
AT2G45480 GRF
AT2G40620 bZIP
AT1G64625 bHLH
AT1G49010 MYBrelated
AT1G06850 bZIP
Flower
Fruit
Leaf
Root
AT5G65410 ZFHD
AT5G52170 HB
AT5G39760 ZFHD
AT3G38170 ND
AT3G00730 HB
AT3G61150 Homeobox
AT2G18350 ZFHD
AT2G17410 ARID
AT1G75240 ZFHD
Flower
Fruit
Leaf
Root
AT5G52170 HB
AT5G03790 HB
AT4G00730 HB
AT3G61150 Homeobox
AT1G76110 ARID
AT1G73360 Homeobox
Flower
Fruit
Leaf
Root
AT5G67580 MYBrelated
AT5G58900 MYBrelated
AT5G25830 C2C2gata
AT5G05790 MYBrelated
AT3G60530 C2C2gata
AT3G51080 C2C2gata
AT3G45170 C2C2gata
AT3G24050 C2C2gata
AT3G06740 C2C2gata
AT1G72740 MYBrelated
AT1G52150 HB
AT1G08010 C2C2gata
Flower
Fruit
Leaf
Root
AT5G66700 HB
AT5G47370 Homeobox
AT5G03790 HB
AT4G36740 HB
AT3G28920 ZFHD
AT3G18990 ABI3VP1
AT3G01470 Homeobox
AT3G01220 Homeobox
AT2G18350 ZFHD
AT2G17410 ARID
AT1G75240 ZFHD
AT1G70920 Homeobox
AT1G69780 Homeobox
AT1G49480 REM
AT1G20910 ARID
Flower
Fruit
Leaf
Root
AT2G01600 G2like
Flower
Fruit
Leaf
Root
AT5G52660 MYBrelated
AT5G47390 MYBrelated
AT5G17300 MYBrelated
AT5G02840 MYBrelated
AT4G01280 MYBrelated
AT3G10113 MYBrelated
AT3G09600 MYBrelated
AT2G28920 ND
AT1G74840 MYBrelated
AT1G28470 NAC
AT1G18330 MYBrelated
AT1G04880 ARID
AT1G01060 MYBrelated
Flower
Fruit
Leaf
Root
AT5G60130 ABI3VP1
AT5G56110 MYB
AT5G54230 MYB
AT5G16600 MYB
AT5G10280 MYB
AT4G38620 MYB
AT4G26030 C2H2
AT4G14770 CPP
AT4G05100 MYB
AT4G01680 MYB
AT3G53200 MYB
AT3G30210 MYB
AT3G22780 CPP
AT3G22760 CPP
AT3G12820 MYB
AT3G12720 MYB
AT3G10580 MYBrelated
AT3G06490 MYB
AT3G01530 MYB
AT2G20110 CPP
AT1G68320 MYB
AT1G18570 MYB
AT1G16490 MYB
AT1G09540 MYB
Flower
Fruit
Leaf
Root

(5)

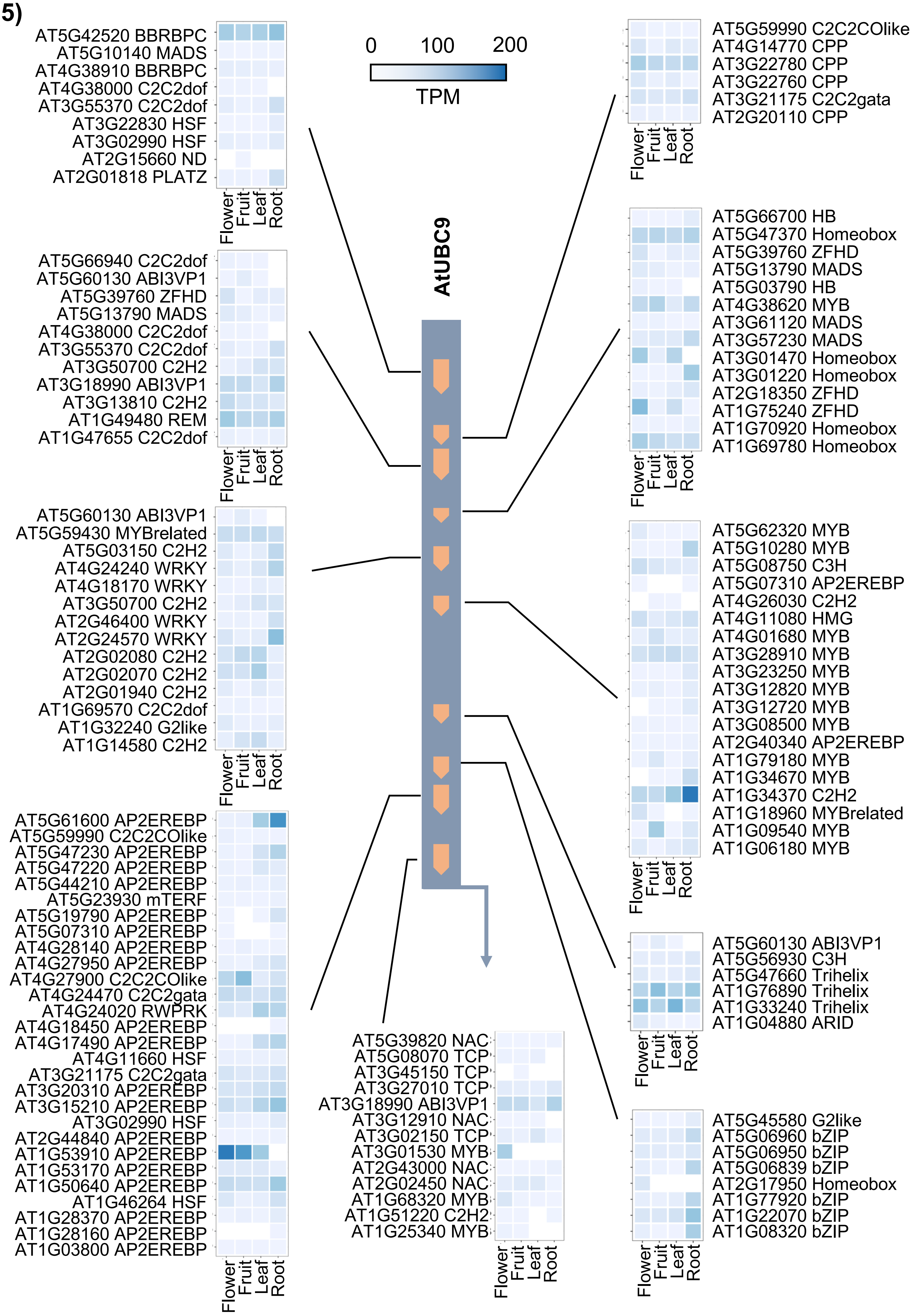

AtUBC9
0
100
200
TPM
Flower
Fruit
Leaf
Root
AT5G42520 BBRBPC
AT5G10140 MADS
AT4G38910 BBRBPC
AT4G38000 C2C2dof
AT3G55370 C2C2dof
AT3G22830 HSF
AT3G02990 HSF
AT2G15660 ND
AT2G01818 PLATZ
AT5G66940 C2C2dof
AT5G60130 ABI3VP1
AT5G39760 ZFHD
AT5G13790 MADS
AT4G38000 C2C2dof
AT3G55370 C2C2dof
AT3G50700 C2H2
AT3G18990 ABI3VP1
AT3G13810 C2H2
AT1G49480 REM
AT1G47655 C2C2dof
AT5G60130 ABI3VP1
AT5G59430 MYBrelated
AT5G03150 C2H2
AT4G24240 WRKY
AT4G18170 WRKY
AT3G50700 C2H2
AT2G46400 WRKY
AT2G24570 WRKY
AT2G02080 C2H2
AT2G02070 C2H2
AT2G01940 C2H2
AT1G69570 C2C2dof
AT1G32240 G2like
AT1G14580 C2H2
AT5G61600 AP2EREBP
AT5G59990 C2C2COlike
AT5G47230 AP2EREBP
AT5G47220 AP2EREBP
AT5G44210 AP2EREBP
AT5G23930 mTERF
AT5G19790 AP2EREBP
AT5G07310 AP2EREBP
AT4G28140 AP2EREBP
AT4G27950 AP2EREBP
AT4G27900 C2C2COlike
AT4G24470 C2C2gata
AT4G24020 RWPRK
AT4G18450 AP2EREBP
AT4G17490 AP2EREBP
AT4G11660 HSF
AT3G21175 C2C2gata
AT3G20310 AP2EREBP
AT3G15210 AP2EREBP
AT3G02990 HSF
AT2G44840 AP2EREBP
AT1G53910 AP2EREBP
AT1G53170 AP2EREBP
AT1G50640 AP2EREBP
AT1G46264 HSF
AT1G28370 AP2EREBP
AT1G28160 AP2EREBP
AT1G03800 AP2EREBP
AT5G39820 NAC
AT5G08070 TCP
AT3G45150 TCP
AT3G27010 TCP
AT3G18990 ABI3VP1
AT3G12910 NAC
AT3G02150 TCP
AT3G01530 MYB
AT2G43000 NAC
AT2G02450 NAC
AT1G68320 MYB
AT1G51220 C2H2
AT1G25340 MYB
AT5G59990 C2C2COlike
AT4G14770 CPP
AT3G22780 CPP
AT3G22760 CPP
AT3G21175 C2C2gata
AT2G20110 CPP
AT5G66700 HB
AT5G47370 Homeobox
AT5G39760 ZFHD
AT5G13790 MADS
AT5G03790 HB
AT4G38620 MYB
AT3G61120 MADS
AT3G57230 MADS
AT3G01470 Homeobox
AT3G01220 Homeobox
AT2G18350 ZFHD
AT1G75240 ZFHD
AT1G70920 Homeobox
AT1G69780 Homeobox
AT5G62320 MYB
AT5G10280 MYB
AT5G08750 C3H
AT5G07310 AP2EREBP
AT4G26030 C2H2
AT4G11080 HMG
AT4G01680 MYB
AT3G28910 MYB
AT3G23250 MYB
AT3G12820 MYB
AT3G12720 MYB
AT3G08500 MYB
AT2G40340 AP2EREBP
AT1G79180 MYB
AT1G34670 MYB
AT1G34370 C2H2
AT1G18960 MYBrelated
AT1G09540 MYB
AT1G06180 MYB
AT5G60130 ABI3VP1
AT5G56930 C3H
AT5G47660 Trihelix
AT1G76890 Trihelix
AT1G33240 Trihelix
AT1G04880 ARID
AT5G45580 G2like
AT5G06960 bZIP
AT5G06950 bZIP
AT5G06839 bZIP
AT2G17950 Homeobox
AT1G77920 bZIP
AT1G22070 bZIP
AT1G08320 bZIP

(6)

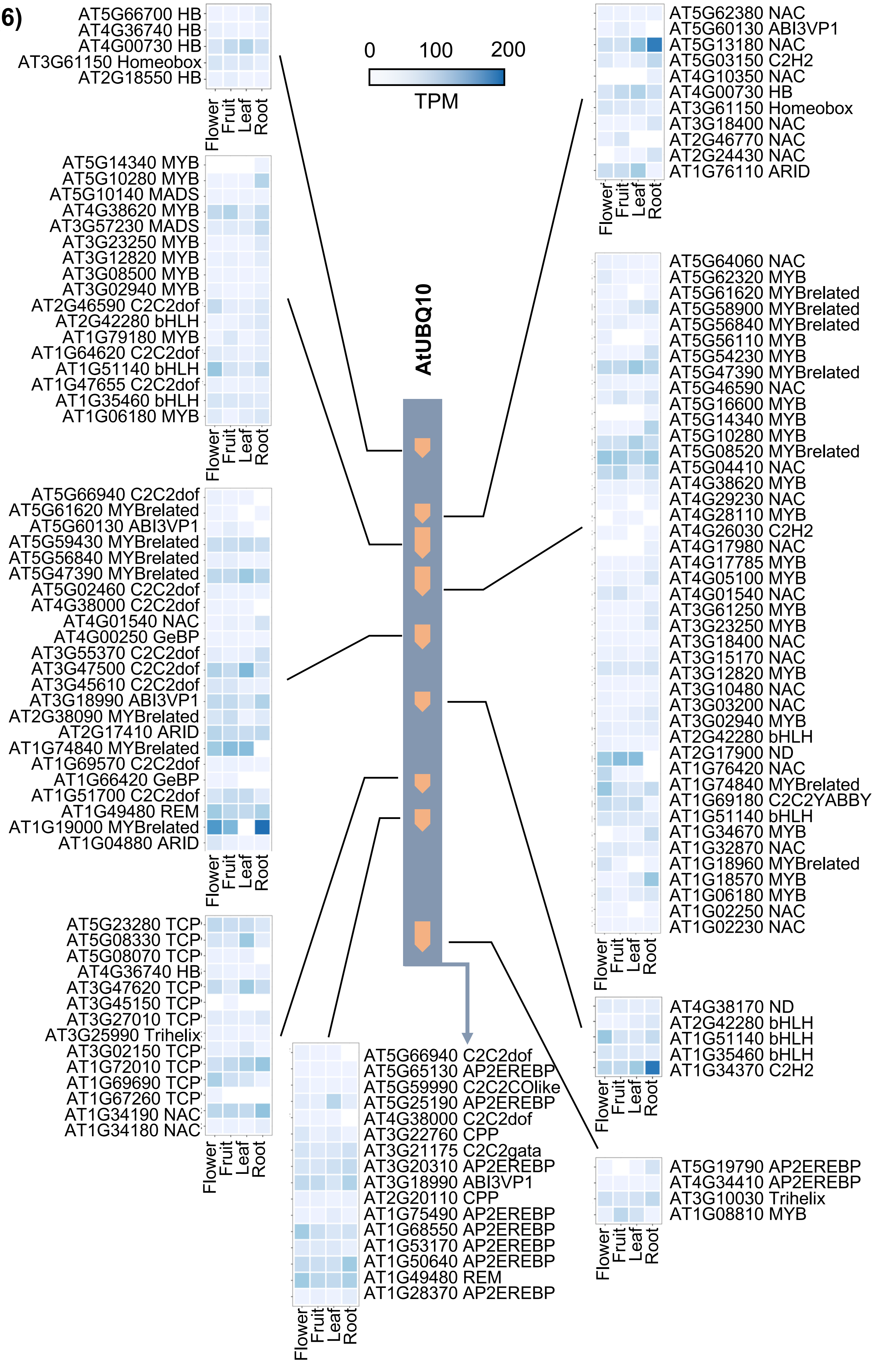
AtUBQ10
0
100
200
TPM
Flower
Fruit
Leaf
Root
AT5G66700 HB
AT4G36740 HB
AT4G00730 HB
AT3G61150 Homeobox
AT2G18550 HB
AT5G14340 MYB
AT5G10280 MYB
AT5G10140 MADS
AT4G38620 MYB
AT3G57230 MADS
AT3G23250 MYB
AT3G12820 MYB
AT3G08500 MYB
AT3G02940 MYB
AT2G46590 C2C2dof
AT2G42280 bHLH
AT1G79180 MYB
AT1G64620 C2C2dof
AT1G51140 bHLH
AT1G47655 C2C2dof
AT1G35460 bHLH
AT1G06180 MYB
AT5G66940 C2C2dof
AT5G61620 MYBrelated
AT5G60130 ABI3VP1
AT5G59430 MYBrelated
AT5G56840 MYBrelated
AT5G47390 MYBrelated
AT5G02460 C2C2dof
AT4G38000 C2C2dof
AT4G01540 NAC
AT4G00250 GeBP
AT3G55370 C2C2dof
AT3G47500 C2C2dof
AT3G45610 C2C2dof
AT3G18990 ABI3VP1
AT2G38090 MYBrelated
AT2G17410 ARID
AT1G74840 MYBrelated
AT1G69570 C2C2dof
AT1G66420 GeBP
AT1G51700 C2C2dof
AT1G49480 REM
AT1G19000 MYBrelated
AT1G04880 ARID
AT5G23280 TCP
AT5G08330 TCP
AT5G08070 TCP
AT4G36740 HB
AT3G47620 TCP
AT3G45150 TCP
AT3G27010 TCP
AT3G25990 Trihelix
AT3G02150 TCP
AT1G72010 TCP
AT1G69690 TCP
AT1G67260 TCP
AT1G34190 NAC
AT1G34180 NAC
AT5G66940 C2C2dof
AT5G65130 AP2EREBP
AT5G59990 C2C2COlike
AT5G25190 AP2EREBP
AT4G38000 C2C2dof
AT3G22760 CPP
AT3G21175 C2C2gata
AT3G20310 AP2EREBP
AT3G18990 ABI3VP1
AT2G20110 CPP
AT1G75490 AP2EREBP
AT1G68550 AP2EREBP
AT1G53170 AP2EREBP
AT1G50640 AP2EREBP
AT1G49480 REM
AT1G28370 AP2EREBP
AT5G62380 NAC
AT5G60130 ABI3VP1
AT5G13180 NAC
AT5G03150 C2H2
AT4G10350 NAC
AT4G00730 HB
AT3G61150 Homeobox
AT3G18400 NAC
AT2G46770 NAC
AT2G24430 NAC
AT1G76110 ARID
AT5G64060 NAC
AT5G62320 MYB
AT5G61620 MYBrelated
AT5G58900 MYBrelated
AT5G56840 MYBrelated
AT5G56110 MYB
AT5G54230 MYB
AT5G47390 MYBrelated
AT5G46590 NAC
AT5G16600 MYB
AT5G14340 MYB
AT5G10280 MYB
AT5G08520 MYBrelated
AT5G04410 NAC
AT4G38620 MYB
AT4G29230 NAC
AT4G28110 MYB
AT4G26030 C2H2
AT4G17980 NAC
AT4G17785 MYB
AT4G05100 MYB
AT4G01540 NAC
AT3G61250 MYB
AT3G23250 MYB
AT3G18400 NAC
AT3G15170 NAC
AT3G12820 MYB
AT3G10480 NAC
AT3G03200 NAC
AT3G02940 MYB
AT2G42280 bHLH
AT2G17900 ND
AT1G76420 NAC
AT1G74840 MYBrelated
AT1G69180 C2C2YABBY
AT1G51140 bHLH
AT1G34670 MYB
AT1G32870 NAC
AT1G18960 MYBrelated
AT1G18570 MYB
AT1G06180 MYB
AT1G02250 NAC
AT1G02230 NAC
AT4G38170 ND
AT2G42280 bHLH
AT1G51140 bHLH
AT1G35460 bHLH
AT1G34370 C2H2
AT5G19790 AP2EREBP
AT4G34410 AP2EREBP
AT3G10030 Trihelix
AT1G08810 MYB

**Supplementary Data File 4**

Multiple sequence alignment of a *cis*-regulatory element common to all 14 promoters from plant-infecting viruses and bacteria. The red box ☐ highlights consensus binding motifs for the TGACG-motif binding (TGA) basic-leucine zipper (B-ZIP) transcription factors.

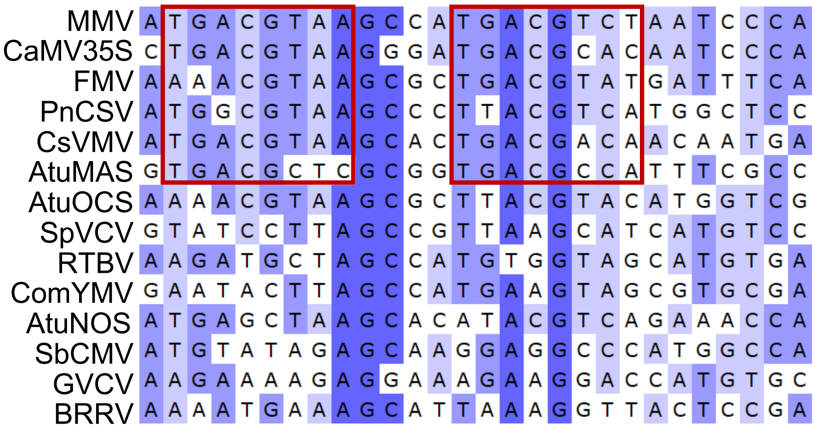

**Supplementary Data File 5**

Deletion of many candidate *cis*-regulatory elements (CRE) had no significant impact on expression. Error bars = 2 x standard error; P-values were calculated using unpaired two-tailed Student's t-test; *P<0.05, **P< 0.01, ***P< 0.001; n=3

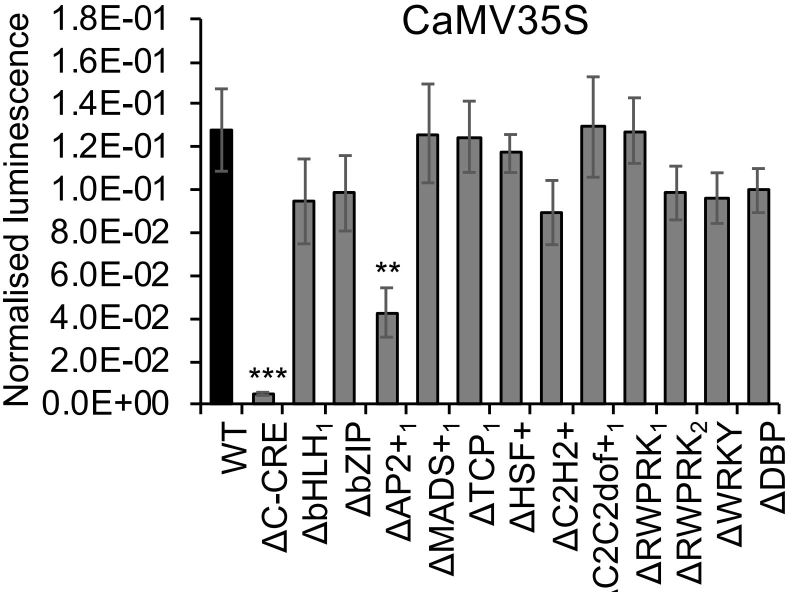


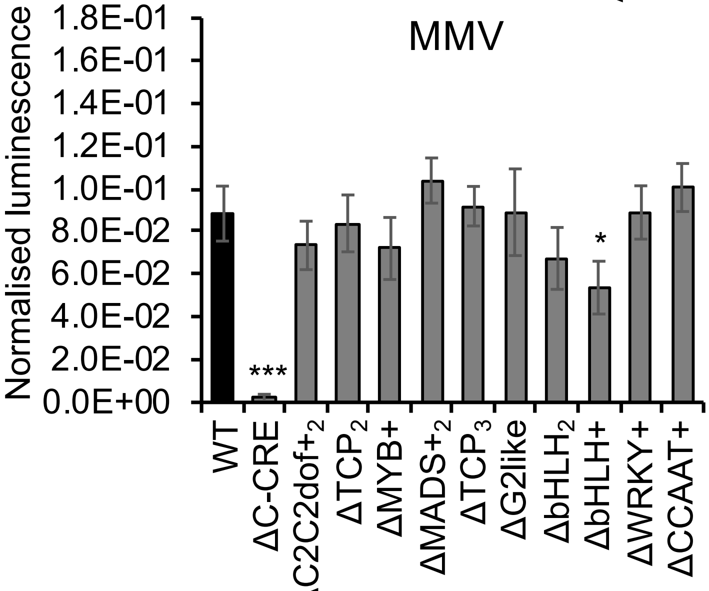


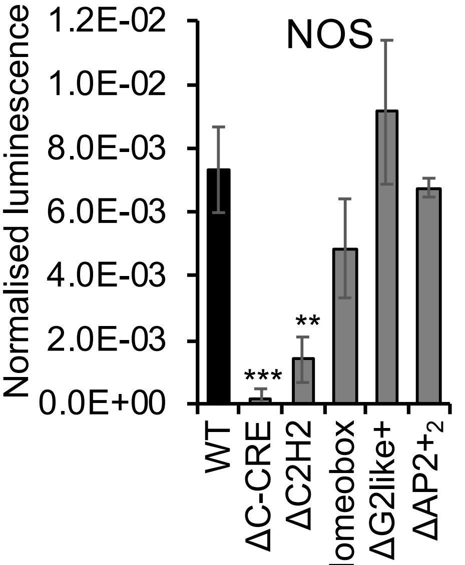

**Supplementary Data 6**

Combining certain *cis*-regulatory elements (CREs) into the variable regions of MinSyns did not result in significant expression

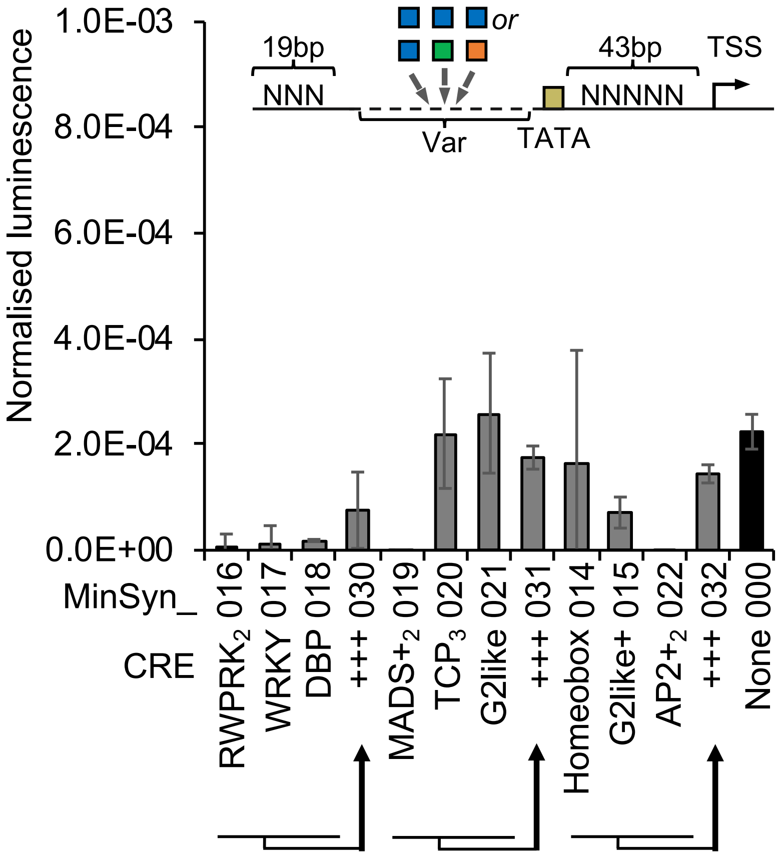

Supplementary Data 7. Sequences and predicted strengths of all computationally designed Minimal Synthetic Promoters (MinSyns)

>MinSyn_1492|Strength:0.000219375
GCGTGTCGTTTTAGTGAGGAACCACGTCTACAATGGTACTTGTGTACAGGGCTCACTGCTAGAGCAAG
TGGATGGAGGACCGATGCTGATCTTGCCCTATATAAGGTTTTGCTATTCATTGAAAGCAGTAGTGACT
GATTTGTATATA
>MinSyn_1126|Strength:0.000231594
GCGTGTCGTTTTAGTGAGGGTGGGAGCCACCAAACTGGTACTTGTGTACAGGGCTCACTGCTGAAGCA
TCTTCCAAGACCTCTACAAAACTGGTACTTGCTATATAAGGTTTTGCTATTCATTGAAAGCAGTAGTG
ACTGATTTGTATATA
>MinSyn_1298|Strength:0.000237417
GCGTGTCGTTTTAGTGAGGGTGGGAGCCACCAGATGCTGATCTTGCCTGCCTTTGAAGCATCTTCCGA
CCTCTACAAAACTGGTACTTGTGTTTTCAACAAACCACTATGCCCAATTAGGTTGTCTGAGCAAGTGG
ATACCACTATGCCCAATTAGGTTGTCTGCCTATATAAGGTTTTGCTATTCATTGAAAGCAGTAGTGAC
TGATTTGTATATA
>MinSyn_1485|Strength:0.000240934
GCGTGTCGTTTTAGTGAGGTCACTATCAGCTTAGGTTGTCTGCACCTCACAAACCATTATTGCGGGAG
GACCGATGCTGATCTTGCCTGCTATATAAGGTTTTGCTATTCATTGAAAGCAGTAGTGACTGATTTGT
ATATA
>MinSyn_1872|Strength:0.000243971
GCGTGTCGTTTTAGTGAGGCAGTGGTCCCTCCACTTAGGTTGTCTGCACCTCACATGTAGGCTGTGGG
AGCCACCAAATTAGGTTGTCTGCACCTCACACTATATAAGGTTTTGCTATTCATTGAAAGCAGTAGTG
ACTGATTTGTATATA
>MinSyn_1884|Strength:0.000249068
GCGTGTCGTTTTAGTGAGGCAAATATTTCTTGTAGGGCTCACTGCTAGGAGGACCGATGCTAACCATT
ATTGCGGACCACTATGCCCAATTAGGCTATATAAGGTTTTGCTATTCATTGAAAGCAGTAGTGACTGA
TTTGTATATA
>MinSyn_1562|Strength:0.000257005
GCGTGTCGTTTTAGTGAGGCAGCCACTTGTGTTCAGCTTAGCAAGGTGGGAGCCACCATGTAGGCTAT
CAGCTTAGCAAGACCTCTACCTATATAAGGTTTTGCTATTCATTGAAAGCAGTAGTGACTGATTTGTA
TATA
>MinSyn_1295|Strength:0.000263
GCGTGTCGTTTTAGTGAGGAGGTGGCTCCTACACCTCTACAAAACTGGTACTTTGGTGGAGCACGACA
TTGTGTACAGGGCTCACTGCTAGGAGGACTATATAAGGTTTTGCTATTCATTGAAAGCAGTAGTGACT
GATTTGTATATA
>MinSyn_1924|Strength:0.000265358
GCGTGTCGTTTTAGTGAGGTCTCTCTGCCGACAGTGGTCCCAAAGGTAGGTCACGACCACTATGCCCA
ATTAGCAAGTGGATGGTTGTCTGCACCTCACATGTAGGCTACTATATAAGGTTTTGCTATTCATTGAA
AGCAGTAGTGACTGATTTGTATATA
>MinSyn_1651|Strength:0.000267033
GCGTGTCGTTTTAGTGAGGTCCATCAACAAATAATCCAAGTAAGATGCCCAATTAGGTTGTCTGCACC
TCACATCAGCCACTTGTGTCGGTAGGTCACGACCACTATGCCCAATTAGCTATATAAGGTTTTGCTAT
TCATTGAAAGCAGTAGTGACTGATTTGTATATA
>MinSyn_1975|Strength:0.000271156
GCGTGTCGTTTTAGTGAGGAAAAATGTCAAAGATAGTAGGTCACGACCCAGCCACTTGTGTAAACTGG
TACTTGTGTACAGGGCTCACTGCCTATATAAGGTTTTGCTATTCATTGAAAGCAGTAGTGACTGATTT
GTATATA
>MinSyn_1178|Strength:0.000274599
GCGTGTCGTTTTAGTGAGGAGCAAGTGGATGGACCGATGCTGATCTTGCCTGCCTTGAACCACGTCTA
CAATAGCAAGACCCTATATAAGGTTTTGCTATTCATTGAAAGCAGTAGTGACTGATTTGTATATA
>MinSyn_1713|Strength:0.000275232
GCGTGTCGTTTTAGTGAGGCAGCCACTTGTGTTGGTACTTGTAACCATTATTGCGCCAATTAGGTTGT
CTGCACCTCACATGAACCACGTCTACAACCACTATGCCCAATTAGGTTGTCTGCTATATAAGGTTTTG

CTATTCATTGAAAGCAGTAGTGACTGATTTGTATATA
>MinSyn_1893|Strength:0.000276264
GCGTGTCGTTTTAGTGAGGAATTTCGGGAAACCTCCTCGTGCTAGGAGGACCGATGCTGTGGGAGCCA
CCATGTCTGCACCTCACATGTAGGCTATCAGCCTATATAAGGTTTTGCTATTCATTGAAAGCAGTAGT
GACTGATTTGTATATA
>MinSyn_1292|Strength:0.000282055
GCGTGTCGTTTTAGTGAGGATTGCGATAAAGGAAAGGCTGGTACTTGTGTACAAACCACGTCTACAAT
GCACCTCACATGTAGGCTATCAGCTTAGCCTATATAAGGTTTTGCTATTCATTGAAAGCAGTAGTGAC
TGATTTGTATATA
>MinSyn_1245|Strength:0.000283159
GCGTGTCGTTTTAGTGAGGTGGTGGAGCACGACAACAGGGCTCAAACCACGTCTACAAAGGAGGACCG
ATGCTGATCTTGCCTAAAAATGTCAAAGATATGTGTACAGGGCTCACTGCTAGGAGGAAGGTGGCTCC
TACGTTGTCTGCACCTCACATGTAGGCTATCCTATATAAGGTTTTGCTATTCATTGAAAGCAGTAGTG
ACTGATTTGTATATA
>MinSyn_1390|Strength:0.000283902
GCGTGTCGTTTTAGTGAGGTTTTCAACAACCTCTACAAAACTGGTACTTGTGTGGTGGAGCACGACAT
GCCTGCCTTGATCTATATAAGGTTTTGCTATTCATTGAAAGCAGTAGTGACTGATTTGTATATA
>MinSyn_1135|Strength:0.000284524
GCGTGTCGTTTTAGTGAGGTTATGACCCCCGCCGATGACGCGGGAGGCTCACTGCTAGGAGGACCGAT
GCTGAAGCATCTTCCCCTCACATGTAGGCTATCAGCTTAGCAAGACTATATAAGGTTTTGCTATTCAT
TGAAAGCAGTAGTGACTGATTTGTATATA
>MinSyn_1588|Strength:0.000286198
GCGTGTCGTTTTAGTGAGGTTTTCAACAACTCACTGCTAGGAGGACCGAATCGAAAGGACAGTACTGC
ACCTCACATGTAGGCTATCAGCAGTGGTCCCTCCACGACCACTATGCCCAATTAGGTTGTCTGCACAA
ATATTTCTTGTACTGCTAGGAGGACCGATGCCTATATAAGGTTTTGCTATTCATTGAAAGCAGTAGTG
ACTGATTTGTATATA
>MinSyn_1980|Strength:0.000287752
GCGTGTCGTTTTAGTGAGGATTGCGATAAAGGAAAGGTGCCCAATTAGGTTGTCTCAAATATTTCTTG
TGCTTAGCAAGACCTCTACAAAACTGGCTATATAAGGTTTTGCTATTCATTGAAAGCAGTAGTGACTG
ATTTGTATATA
>MinSyn_1949|Strength:0.000292956
GCGTGTCGTTTTAGTGAGGTTATGACCCCCGCCGATGACGCGGGAGGACCGATGCTGATCTTGCCTGC
CTTGAAACCACGTCTACAATATCAGCTTAGCAAGACCTCTACAAAACTATATAAGGTTTTGCTATTCA
TTGAAAGCAGTAGTGACTGATTTGTATATA
>MinSyn_129|Strength:0.000296041
GCGTGTCGTTTTAGTGAGGTTATGACCCCCGCCGATGACGCGGGATAGGCTATCAGCTTAGCAAGACC
TAAAAATGTCAAAGATAACCACTATGCCCAATTAGGTTGTCTTCACTATCAGCGGCTCACTGCTAGGA
GGACCGATCTATATAAGGTTTTGCTATTCATTGAAAGCAGTAGTGACTGATTTGTATATA
>MinSyn_1806|Strength:0.000296047
GCGTGTCGTTTTAGTGAGGTCACTATCAGCTATCAGCTTAGCAAGACCTAGCAAGTGGATTAGCAAGA
CCTCTACAAAAAATGTCAAAGATAAGGTCACGACCACTATGCCCAATTCTATATAAGGTTTTGCTATT
CATTGAAAGCAGTAGTGACTGATTTGTATATA
>MinSyn_1467|Strength:0.000298493
GCGTGTCGTTTTAGTGAGGGGAAAAAGAAGAGGTTGCCCAATTAGGTCAAATATTTCTTGTCCCAATT
AGGTTGTCTGCACCTCACATGTATGAAGCATCTTCCTCACATGTAGGCTATCAGCTTAGCAAGATTGC
GATAAAGGAAAGGACAAAACTGGTACTTGTGTACAGGGCCTATATAAGGTTTTGCTATTCATTGAAAG
CAGTAGTGACTGATTTGTATATA
>MinSyn_1692|Strength:0.000303832
GCGTGTCGTTTTAGTGAGGCAGTGGTCCCTCCACATCTTGCTCACTATCAGCCACTATGCCCAATTAG
GTTGTCTGGGAAAAAGAAGAGGTCATGTAGGCTATCAGCTTAATCGAAAGGACAGTACTGGTACTTGT
GTACAGGGCTCATTTTCAACAAGACCGATGCTGATCTTGCCTGCTATATAAGGTTTTGCTATTCATTG
AAAGCAGTAGTGACTGATTTGTATATA
>MinSyn_194|Strength:0.000313038
GCGTGTCGTTTTAGTGAGGGGAAAAAGAAGAGGTGTCAGCCACTTGTGTTACAGGGCTCACTGCTAGG

AGGACCGAGCAAGTGGATCCAATTAGGTTGTCTGCACCTCACATGTCAGAAGATCAAAGGGCTACTAG
GAGGACCGATGCTGATCTTGCCTATATAAGGTTTTGCTATTCATTGAAAGCAGTAGTGACTGATTTGT
ATATA
>MinSyn_1144|Strength:0.000313721
GCGTGTCGTTTTAGTGAGGTGAAGATAAGATAATAATGTTGAAGATAAGACACGACCACTATGCCCAA
TTAGGTTGTCTGAAGCATCTTCCGTAGGCTATCAGCTTAGCAAGACCTCTATATAAGGTTTTGCTATT
CATTGAAAGCAGTAGTGACTGATTTGTATATA
>MinSyn_1404|Strength:0.000319449
GCGTGTCGTTTTAGTGAGGTTTTCAACAAGTAGGCTATCAGCTTAGCAAGACCTCAGCCACTTGTGTG
TAGGTCACGACCACTATGCCCAATTACAAATATTTCTTGTGAACCATTATTGCGCGATGCTGAGCAAG
TGGATGCTGATCTTGCCTGCCTTGATGGGAAAAAGAAGAGGTTATCAGCTTAGCAAGACCTCTACAAA
ACTCACTATCAGCAGGTTGTCTGCACCTCACATGTAGGCCTATATAAGGTTTTGCTATTCATTGAAAG
CAGTAGTGACTGATTTGTATATA
>MinSyn_1938|Strength:0.000321911
GCGTGTCGTTTTAGTGAGGTTATGACCCCCGCCGATGACGCGGGAGGAGGACCGATGCTGATCTTGCC
TGCCTTGTGAAGCATCTTCCGGCTATCAGCTTACTATATAAGGTTTTGCTATTCATTGAAAGCAGTAG
TGACTGATTTGTATATA
>MinSyn_1727|Strength:0.000322209
GCGTGTCGTTTTAGTGAGGATTGCGATAAAGGAAAGGCCGATGCTGATCTAGGTGGCTCCTACACCGA
TGCTGATCTTGCCAGTGGTCCCTCCACTGTGTACAGGGCTCACTGCTAGGAGCTATATAAGGTTTTGC
TATTCATTGAAAGCAGTAGTGACTGATTTGTATATA
>MinSyn_1796|Strength:0.000324512
GCGTGTCGTTTTAGTGAGGTCAGAAGATCAAAGGGCTAAAAACTGGTACTTGTGTACTGGTGGAGCAC
GACATTGTCTGCACCTCACATGCTATATAAGGTTTTGCTATTCATTGAAAGCAGTAGTGACTGATTTG
TATATA
>MinSyn_116|Strength:0.00032615
GCGTGTCGTTTTAGTGAGGAATTTCGGGAAACCTCCTCGTTCAGAAGATCAAAGGGCTAGCTATCAGC
TTAGCAAGACCTCAACCATTATTGCGACAAAACTGGTACTTGTGTACAGGGCTCACCTATATAAGGTT
TTGCTATTCATTGAAAGCAGTAGTGACTGATTTGTATATA
>MinSyn_153|Strength:0.000326571
GCGTGTCGTTTTAGTGAGGGTGGGAGCCACCATTGTGTACAGGGCTCACTGCTATTATGACCCCCGCC
GATGACGCGGGACCGATGCTGATCTTGCCTGCCTTGATATCGAAAGGACAGTACTGCTAGGAGGACCG
ATGAGGTGGCTCCTACGCTATCAGCTTAGCAAGACCTCTACACTATATAAGGTTTTGCTATTCATTGA
AAGCAGTAGTGACTGATTTGTATATA
>MinSyn_1690|Strength:0.000327105
GCGTGTCGTTTTAGTGAGGTCCATCAACAAATAATCCAAGTAAGGGTACTTGTGTATTGCGATAAAGG
AAAGGCTAGGAGGACCGATGCTGATCAGGTGGCTCCTACCCCAATTAGGTTGTCTGCACCTCACATGT
ACTATATAAGGTTTTGCTATTCATTGAAAGCAGTAGTGACTGATTTGTATATA
>MinSyn_171|Strength:0.00033418
GCGTGTCGTTTTAGTGAGGAGCAAGTGGATAGCAAGACCTCTACAAAACTGGTACTTGTGGTGGGAGC
CACCAACAGGGCTCACTGCTAGGAGCAGCCACTTGTGTTAGCAAGACCTCTACAAAACAGGTGGCTCC
TACCTGGTACTTGTGTACAGCTATATAAGGTTTTGCTATTCATTGAAAGCAGTAGTGACTGATTTGTA
TATA
>MinSyn_1971|Strength:0.000334958
GCGTGTCGTTTTAGTGAGGGCACACCAGCATGTGTTGATCACCAGCTGCTATCAGCTTAGCAAGACAT
CGAAAGGACAGTACTTGTGTACAGGGCTCACTGCTAGGAGGCTATATAAGGTTTTGCTATTCATTGAA
AGCAGTAGTGACTGATTTGTATATA
>MinSyn_1868|Strength:0.000335447
GCGTGTCGTTTTAGTGAGGAAGATTGATGAAAAGTCAAAAACAAAAATCAATTATACCGATGCTGATC
TTGCCTGTGAAGCATCTTCCCTATCAGCTTAGCAAGACCTCTACAGGTGGCTCCTACGGCTCACTGCT
AGGAGGACCGATGCTAATTTCGGGAAACCTCCTCGCGGTAGGTCACGACCACTATGCCAAATATTTCT
TGTCTATGCCCAATTAGGTTGTCTGCACCTCTGGTGGAGCACGACATACAAAACTGGTACTTGTGTAC
AGGGCTCTATATAAGGTTTTGCTATTCATTGAAAGCAGTAGTGACTGATTTGTATATA
>MinSyn_11|Strength:0.000336236

GCGTGTCGTTTTAGTGAGGTTTTCAACAAGGGCTCACTGCTGGAAAAAGAAGAGGTCGATGCTGATCT
TGCCTGCCTTGATGAAATTTCGGGAAACCTCCTCGGCTATCAGCTTAGCAAGACCTATATAAGGTTTT
GCTATTCATTGAAAGCAGTAGTGACTGATTTGTATATA
>MinSyn_182|Strength:0.00033916
GCGTGTCGTTTTAGTGAGGTCCATCAACAAATAATCCAAGTAAGTCACGACCACTATTCAGAAGATCA
AAGGGCTAGGTTGTCAGCAAGTGGATCAGCTTAGCAAGACCTCTACAAAACTGGTCACTATCAGCTGA
TCTTGCCTGCCTTGATGATGGTGGAGCACGACACAAAACTGGTACTTGTGTACAGGGCCTATATAAGG
TTTTGCTATTCATTGAAAGCAGTAGTGACTGATTTGTATATA
>MinSyn_1250|Strength:0.000339958
GCGTGTCGTTTTAGTGAGGCAAATATTTCTTGTTGTCTGCACCTCACATGTAGGCTATCATCAGAAGA
TCAAAGGGCTAGGTTGTCTGCACCTTTTCAACAAAGGTCACGACCACTATGCCCAATTAGGTAGGTGG
CTCCTACGCACCTCACATGTAGGGGAAAAAGAAGAGGTCACTGCTAGGAGGACCGATGCTGATCTTTC
TCTCTGCCGACAGTGGTCCCAAACCCAATTAGGTTGTCTGCACCTCACATAACCACGTCTACAACCCA
ATTAGGTTGTCTGCACCTCACATCTATATAAGGTTTTGCTATTCATTGAAAGCAGTAGTGACTGATTT
GTATATA
>MinSyn_1828|Strength:0.000340023
GCGTGTCGTTTTAGTGAGGAGCAAGTGGATCACCTCACATGTAGGCTATCAGCTTACAAATATTTCTT
GTTGTACAGGGCTCACTGCTAGGAGGACCGTCTCTCTGCCGACAGTGGTCCCAAACACAGTGGGAGCC
ACCACACATGTAGGCTATCAGCTTAGCAAGACCTAACCACGTCTACAAAGCAAGACCTCTACAAAACT
GGTACTATATAAGGTTTTGCTATTCATTGAAAGCAGTAGTGACTGATTTGTATATA
>MinSyn_1767|Strength:0.000340443
GCGTGTCGTTTTAGTGAGGTTATGACCCCCGCCGATGACGCGGGAATGCTGATCTTGCCTGCCTTAAA
AATGTCAAAGATAGTACTTGTGTACAGGGCTCACTGCTCTATATAAGGTTTTGCTATTCATTGAAAGC
AGTAGTGACTGATTTGTATATA
>MinSyn_1693|Strength:0.000340747
GCGTGTCGTTTTAGTGAGGGCACACCAGCATGTGTTGATCACCAGCTCTTGTGTACAGGGCTCACTGC
TAGGAGGACAACCACGTCTACAATGCTGATCTTGCCCTATATAAGGTTTTGCTATTCATTGAAAGCAG
TAGTGACTGATTTGTATATA
>MinSyn_118|Strength:0.000341026
GCGTGTCGTTTTAGTGAGGTCCATCAACAAATAATCCAAGTAAGACAGGGCTCACTGCTAGGGGAAAA
AGAAGAGGTGGTTGTCTGCACCTCACATGTAGCTATATAAGGTTTTGCTATTCATTGAAAGCAGTAGT
GACTGATTTGTATATA
>MinSyn_1391|Strength:0.000342397
GCGTGTCGTTTTAGTGAGGTGGTGGAGCACGACAAGGCTACAGTGGTCCCTCCACCTGCCTTGATATC
GAAAGGACAGTAGCACCTCACATGTAGGCTATCAGCTTAGCCTATATAAGGTTTTGCTATTCATTGAA
AGCAGTAGTGACTGATTTGTATATA
>MinSyn_1969|Strength:0.000344584
GCGTGTCGTTTTAGTGAGGCAGTGGTCCCTCCACCAGGGCTCACTGCTAGGAGGAGGTGGCTCCTACT
AGGTCAATTGCGATAAAGGAAAGGCTGGTACTTGTGTACAGGGCTCACTGCTCTATATAAGGTTTTGC
TATTCATTGAAAGCAGTAGTGACTGATTTGTATATA
>MinSyn_1878|Strength:0.000346939
GCGTGTCGTTTTAGTGAGGCAAATATTTCTTGTCCTCTACAAATTTCGGGAAACCTCCTCGCTTAGCA
AGACCTCTACAAAACTGGTACTTCTATATAAGGTTTTGCTATTCATTGAAAGCAGTAGTGACTGATTT
GTATATA
>MinSyn_1109|Strength:0.000347557
GCGTGTCGTTTTAGTGAGGTGAAGATAAGATAATAATGTTGAAGATAAGAGCTTAGCAAGACCTCTAC
AAAACTGGTATCAGAAGATCAAAGGGCTAACTTGTGTACAGGGCTCACTGCTAGGAGGCTATATAAGG
TTTTGCTATTCATTGAAAGCAGTAGTGACTGATTTGTATATA
>MinSyn_1770|Strength:0.000351647
GCGTGTCGTTTTAGTGAGGATCCTTACCGCTATGGGTAAGATTCATGTAGGCTATCAGCTTAGCAAGG
TGGGAGCCACCAATTAGGTTGTCTGCACCTCACACAGTGGTCCCTCCACGTCTGCACCTCACATGTAG
GCTAATTTCGGGAAACCTCCTCGAGACCTCTACAAAACTGGTACTTCTATATAAGGTTTTGCTATTCA
TTGAAAGCAGTAGTGACTGATTTGTATATA
>MinSyn_1432|Strength:0.000351717

GCGTGTCGTTTTAGTGAGGGGAAAAAGAAGAGGTTAGGCTATCAGCTTAGCAAGACCTCTATTATGAC
CCCCGCCGATGACGCGGGATTGTGTACAGGGCTCACTGCTAGGAGGACCCTATATAAGGTTTTGCTAT
TCATTGAAAGCAGTAGTGACTGATTTGTATATA
>MinSyn_1753|Strength:0.000352007
GCGTGTCGTTTTAGTGAGGTGGTGGAGCACGACAGACCTCTACAAAACTGGTACTTGTGAGCAAGTGG
ATTCGCGGTAGGTCACGACCACTTTATGACCCCCGCCGATGACGCGGGACCTCACATGTAGGCTATCA
GTGGGAGCCACCAAGCTTAGCAAGACCTCTATCGAAAGGACAGTAGCCCAATTAGGTTGTCTGCACCT
CACCTATATAAGGTTTTGCTATTCATTGAAAGCAGTAGTGACTGATTTGTATATA
>MinSyn_1253|Strength:0.00035376
GCGTGTCGTTTTAGTGAGGTTTTCAACAACCTCTACAAAACTGGTACTTTCTCTCTGCCGACAGTGGT
CCCAAAACCTCTACAAAACTGGTACTTGTGTACAGGAACCACGTCTACAACGCGGTAGGTCACGACCA
CTATGCCCAGCAAGTGGATGTCACGACCACTATATAAGGTTTTGCTATTCATTGAAAGCAGTAGTGAC
TGATTTGTATATA
>MinSyn_1654|Strength:0.000353822
GCGTGTCGTTTTAGTGAGGATTGCGATAAAGGAAAGGTAGCAAGACCTCTACAAAACTGGTACTTGAA
CCATTATTGCGCCGATGCTGATCTTGCCTGCCTTGATGACAGCCACTTGTGTCTGCACCAACCACGTC
TACAAGGTCACGACCACTATGAAAAATGTCAAAGATATAGGTCACGACCACTATGCCCCTATATAAGG
TTTTGCTATTCATTGAAAGCAGTAGTGACTGATTTGTATATA
>MinSyn_1945|Strength:0.000354317
GCGTGTCGTTTTAGTGAGGAAAAATGTCAAAGATAACTTGTGAATTTCGGGAAACCTCCTCGAGCAAG
ACCTCTACAAAACTGGTACCAAATATTTCTTGTAATTAGGTTGTCTGCCTATATAAGGTTTTGCTATT
CATTGAAAGCAGTAGTGACTGATTTGTATATA
>MinSyn_1324|Strength:0.000358314
GCGTGTCGTTTTAGTGAGGTGGTGGAGCACGACATCGCGGTAGGTCACGACCACTATGCTCACTATCA
GCGCCCAATTAGGTTGTCTGCATCGAAAGGACAGTACTTGTGTACAGGGCTCACTAACCACGTCTACA
AGCTATCAGCTTAGCAAGACCTCTACAAAAAAAAATGTCAAAGATATCAGCTTAGCAAGACCTCTACA
AAACTGAAGCATCTTCCTAGGAGGACCAATTTCGGGAAACCTCCTCGACCGATGCTGATCTTGCCTGC
CTTGCTATATAAGGTTTTGCTATTCATTGAAAGCAGTAGTGACTGATTTGTATATA
>MinSyn_1361|Strength:0.000358475
GCGTGTCGTTTTAGTGAGGCAAATATTTCTTGTCTAACCATTATTGCGCTAGCAAGTGGATTACAGGG
CTCACTGCTAGGAGGACCGTCAGAAGATCAAAGGGCTAGGTAGGTCACGACCACTATGCCCAACTATA
TAAGGTTTTGCTATTCATTGAAAGCAGTAGTGACTGATTTGTATATA
>MinSyn_1815|Strength:0.000367835
GCGTGTCGTTTTAGTGAGGAATTTCGGGAAACCTCCTCGTGGTACTTGTGTACAGGGCTCACTGCTAT
CCTTACCGCTATGGGTAAGATTAGGGCTCACTGCTAGGAGGACCGCTATATAAGGTTTTGCTATTCAT
TGAAAGCAGTAGTGACTGATTTGTATATA
>MinSyn_1877|Strength:0.000369607
GCGTGTCGTTTTAGTGAGGCAGTGGTCCCTCCACATTCAGAAGATCAAAGGGCTACGCGGTAGGTCAC
GACATCCTTACCGCTATGGGTAAGATTTGGTACTTGTGTACAGGGCTCACTGCTAATCGAAAGGACAG
TAGTCACGACCACTTTTCAACAACCAATTAGGTTGTCTGCACCTCACATGTAGATTGCGATAAAGGAA
AGGTAGGAGGACCGATGCTGATCCTATATAAGGTTTTGCTATTCATTGAAAGCAGTAGTGACTGATTT
GTATATA
>MinSyn_1876|Strength:0.000371438
GCGTGTCGTTTTAGTGAGGATCCTTACCGCTATGGGTAAGATTGCAAGACCTCCAAATATTTCTTGTT
AGGTCACGACCACTATGGAAAAAGAAGAGGTCGGTAGGTCACGACCACTATCTATATAAGGTTTTGCT
ATTCATTGAAAGCAGTAGTGACTGATTTGTATATA
>MinSyn_1430|Strength:0.000373039
GCGTGTCGTTTTAGTGAGGATTGCGATAAAGGAAAGGAGGTAGGTGGCTCCTACTTAGCAAGACCTCT
ACAAAACTGGTATGAAGCATCTTCCGTAGGCTATCAGCTTAGCAAGACTGAAGATAAGATAATAATGT
TGAAGATAAGATACAAAACTGGTACTTGTGTGGGAGCCACCATGCAACCATTATTGCGATGCCCAATT
AGGTTGTCTGCACCTAAAAATGTCAAAGATATCACGACCACTATGCCCAATTCAGTGGTCCCTCCACA
CTGCTAGGAGGACCGATGCTTCACTATCAGCTCACTGCTAGGAGGACCGATGCTGATCTTCTATATAA
GGTTTTGCTATTCATTGAAAGCAGTAGTGACTGATTTGTATATA
>MinSyn_1873|Strength:0.000373543

GCGTGTCGTTTTAGTGAGGAAAAATGTCAAAGATAGTACAGGGCTCAATCGAAAGGACAGTACGATGC
TGATCTTGCCTGCCTTGATGATCTCTCTGCCGACAGTGGTCCCAAACTCACTGCTAGGAGGACCGATG
CTGAACCATTATTGCGTGCACCTCACATGTAGGCTATCATTATGACCCCCGCCGATGACGCGGGATGT
GTACAGGGCTCACTGCTAGTGGGAGCCACCATTAGCAAGACCTCTACAAAACTGCTATATAAGGTTTT
GCTATTCATTGAAAGCAGTAGTGACTGATTTGTATATA
>MinSyn_1687|Strength:0.000374085
GCGTGTCGTTTTAGTGAGGAATTTCGGGAAACCTCCTCGCTGATCTTGCCTGCCTTGATGAAGGTGGC
TCCTACCAGCTTAGCAAGACCTCTACATGAAGCATCTTCCACCGATGCTGTGGTGGAGCACGACACAG
GGCTCACTGCTAGGAGGACCAAAAATGTCAAAGATATGATCTTGCCTGCCTATATAAGGTTTTGCTAT
TCATTGAAAGCAGTAGTGACTGATTTGTATATA
>MinSyn_1774|Strength:0.000374206
GCGTGTCGTTTTAGTGAGGTTTTCAACAACACCTCACATGTAGGCTATCAGCTTAGCAAGATTGATGA
AAAGTCAAAAACAAAAATCAATTATTCACTGCTAGGAGGACCGATGCTGATAGGTGGCTCCTACTACA
GGCAGTGGTCCCTCCACACTTGTGTACAGGGCTCACTGCCAGCCACTTGTGTAGGCTATCAGCTTAGC
AAGGGAAAAAGAAGAGGTCAGCTTAGCAAGACCTCTACAAAACTCTATATAAGGTTTTGCTATTCATT
GAAAGCAGTAGTGACTGATTTGTATATA
>MinSyn_1910|Strength:0.00037519
GCGTGTCGTTTTAGTGAGGTCCATCAACAAATAATCCAAGTAAGCTTAGCAAGACCTCTACAATTTCG
GGAAACCTCCTCGTACTTGTGTACAGGGCTCACTGCTATCAGAAGATCAAAGGGCTATCACATGTAGG
CTATCAGCTTACTATATAAGGTTTTGCTATTCATTGAAAGCAGTAGTGACTGATTTGTATATA
>MinSyn_1288|Strength:0.000375621
GCGTGTCGTTTTAGTGAGGAACCACGTCTACAATTGCCTGCCTTGATGATTCTCTCTGCCGACAGTGG
TCCCAAATGTAGGCTATCAGCTTAGCAAGACCTCCTATATAAGGTTTTGCTATTCATTGAAAGCAGTA
GTGACTGATTTGTATATA
>MinSyn_1667|Strength:0.000375703
GCGTGTCGTTTTAGTGAGGCAAATATTTCTTGTTTGTGTACAGGGCTCACTGCTAGATTGCGATAAAG
GAAAGGCCACTATGCTCAGAAGATCAAAGGGCTACGACCACTATGCCCAATTAGGTTCTATATAAGGT
TTTGCTATTCATTGAAAGCAGTAGTGACTGATTTGTATATA
>MinSyn_1226|Strength:0.000375737
GCGTGTCGTTTTAGTGAGGAAAAATGTCAAAGATATAGGTTGTCTGCACCTCACAAACCATTATTGCG
AGGTCACGACCACAGCAAGTGGATTGTACAGGGCTCACTGCTAGGAGGACCTTATGACCCCCGCCGAT
GACGCGGGAATGCCCAATTAGGTTGTCTGCTATATAAGGTTTTGCTATTCATTGAAAGCAGTAGTGAC
TGATTTGTATATA
>MinSyn_1848|Strength:0.000382766
GCGTGTCGTTTTAGTGAGGATCGAAAGGACAGTAACATGTAGGCTATCAGCTTAGCAAGACGCTTTGT
CAAAAGCTAAAAAAGATGATGCCATGTAGGCTATCAGCTTAGCAAGGGAAAAAGAAGAGGTACCACTA
TGCCCAATTAGGTTGTCTGCTCACTATCAGCCTAGGAGGACCTATATAAGGTTTTGCTATTCATTGAA
AGCAGTAGTGACTGATTTGTATATA
>MinSyn_1974|Strength:0.000383031
GCGTGTCGTTTTAGTGAGGGTGGGAGCCACCACTTAGCAAGACCTCTACAAAACTGGAAAAAGAAGAG
GTTTGATTGAAGCATCTTCCCGATGCTGATCTTGCCTGCCTTGATGCTATATAAGGTTTTGCTATTCA
TTGAAAGCAGTAGTGACTGATTTGTATATA
>MinSyn_1940|Strength:0.000385118
GCGTGTCGTTTTAGTGAGGTCTCTCTGCCGACAGTGGTCCCAAATTGTGTACAGGGCTCACTGCTATC
CATCAACAAATAATCCAAGTAAGGGTAGGTCACGACCACTATGCCCTTATGACCCCCGCCGATGACGC
GGGACTCACTGCTAGAACCACGTCTACAAGTACAGGGCTCACTGCTAGGAGGACCGATCAGCCACTTG
TGTTTAGGTTGTCTGCACCTCATGGTGGAGCACGACACACGACCACTATGCCCAATTAGGCTATATAA
GGTTTTGCTATTCATTGAAAGCAGTAGTGACTGATTTGTATATA
>MinSyn_1235|Strength:0.000387116
GCGTGTCGTTTTAGTGAGGTCTCTCTGCCGACAGTGGTCCCAAACAGGGCTCACTGCTAGGAGGCAGT
GGTCCCTCCACGGTTGTCTGCACCTCACATGTAGGCTAATCCTTACCGCTATGGGTAAGATTGGTTTT
CAACAAAAGACCTCTACAAAACTGGTACTTGTGATTGCGATAAAGGAAAGGCAAAACTGGTACTTGTG
TACAGGCTATATAAGGTTTTGCTATTCATTGAAAGCAGTAGTGACTGATTTGTATATA
>MinSyn_1640|Strength:0.000388057

GCGTGTCGTTTTAGTGAGGTCTCTCTGCCGACAGTGGTCCCAAACCCAATTAGGTTGTCTGCACCTCA
TCGAAAGGACAGTAACTGCTAGGAGGACCGATGCTGATCTAGCAAGTGGATCTTTATGACCCCCGCCG
ATGACGCGGGAGTCTGCACCTCACATGTAGGAGGTGGCTCCTACGTCACGACCACTATGCCCAATTAC
AGTGGTCCCTCCACAGGACCGATGCTGATCTTGCCCTATATAAGGTTTTGCTATTCATTGAAAGCAGT
AGTGACTGATTTGTATATA
>MinSyn_1198|Strength:0.000388376
GCGTGTCGTTTTAGTGAGGTCACTATCAGCCACATGTATTTTCAACAAGTCAGGTGGCTCCTACACAT
GTAGGCTATCAGCTTAGCAACTATATAAGGTTTTGCTATTCATTGAAAGCAGTAGTGACTGATTTGTA
TATA
>MinSyn_1558|Strength:0.000388673
GCGTGTCGTTTTAGTGAGGGCTTTGTCAAAAGCTAAAAAAGATGATGCTATCAGCTTGGTGGAGCACG
ACAACTTGTGTACAGGGAAAAAGAAGAGGTGCGGTAGGTCACGACCACTATGCCCAATCTATATAAGG
TTTTGCTATTCATTGAAAGCAGTAGTGACTGATTTGTATATA
>MinSyn_1470|Strength:0.000389896
GCGTGTCGTTTTAGTGAGGAGGTGGCTCCTACTCACATGTAGGCTATCAGCTTAGCAATTGCGATAAA
GGAAAGGCGGTAGGTCACGACCACTTGAAGCATCTTCCCATGTAGGCTATCAGCTTCTATATAAGGTT
TTGCTATTCATTGAAAGCAGTAGTGACTGATTTGTATATA
>MinSyn_188|Strength:0.00039094
GCGTGTCGTTTTAGTGAGGAGGTGGCTCCTACTGTGATCCTTACCGCTATGGGTAAGATTGGTTGTCT
GCACCTCACATGTAGGCAAATATTTCTTGTGCTAGGAGGACCGATGCTGATCTTAAAAATGTCAAAGA
TATGCTAGGAGGACCGATGCTATTGCGATAAAGGAAAGGCCAATGGAAAAAGAAGAGGTGGACCGATG
CTGATCTTATCGAAAGGACAGTACTCTACAAAACTGGTACTTGTGTACAGGGCCTATATAAGGTTTTG
CTATTCATTGAAAGCAGTAGTGACTGATTTGTATATA
>MinSyn_178|Strength:0.000391657
GCGTGTCGTTTTAGTGAGGAACCATTATTGCGCAATTAGGTTGTCTGCACCTCAAAGATTGATGAAAA
GTCAAAAACAAAAATCAATTATAGGGCTCACTGCTAGGAGGACCGGAAAAAGAAGAGGTAAGACCTCT
ACAAAATGGTGGAGCACGACACGATGCTGATCTTGCCTGCCTTGACTATATAAGGTTTTGCTATTCAT
TGAAAGCAGTAGTGACTGATTTGTATATA
>MinSyn_1223|Strength:0.000392707
GCGTGTCGTTTTAGTGAGGAATTTCGGGAAACCTCCTCGGGTAGGTCACGACTCAGAAGATCAAAGGG
CTATACAGGGCTCACTAACCACGTCTACAACCAATTAAAAAATGTCAAAGATACGATGCTGATCTTGC
CTGCCTTTGAAGATAAGATAATAATGTTGAAGATAAGAGTACAGGGCTCACTGCTAGGAGGAGCAAGT
GGATCGACCACTATGCCCAATTAGGTTGAACCATTATTGCGAACTGGTACTTGTGTACAGGGCCTATA
TAAGGTTTTGCTATTCATTGAAAGCAGTAGTGACTGATTTGTATATA
>MinSyn_1962|Strength:0.000392928
GCGTGTCGTTTTAGTGAGGAATTTCGGGAAACCTCCTCGGATTCCATCAACAAATAATCCAAGTAAGG
GCTATCAGCTTAGCAAGACCTCTACAGGAAAAAGAAGAGGTTCACGACCACTATGCCCAATTAGGTTG
TCAGTGGTCCCTCCACAGGTCACGACCACTATTGCGATAAAGGAAAGGTGTGTACAGGGCATCGAAAG
GACAGTAGGAGGACCGATGCTGATCTTGCCAACCATTATTGCGCGGTAGGTCACCAAATATTTCTTGT
GAGGACCGATGCTGATCTTGCCTGCCCTATATAAGGTTTTGCTATTCATTGAAAGCAGTAGTGACTGA
TTTGTATATA
>MinSyn_1760|Strength:0.000393391
GCGTGTCGTTTTAGTGAGGTTATGACCCCCGCCGATGACGCGGGAATGTAGGCTATTCACTATCAGCT
ACTTGTGTACAGGGCTCACTGCTAGGAGGTGGGAGCCACCAACTGGCAGTGGTCCCTCCACCTCACAT
GTAGGCTATCAGTCCATCAACAAATAATCCAAGTAAGAAAACTGGTACTTGTGTACAGGCAGCCACTT
GTGTAAGACCTCTACAAAACTGGTACTATATAAGGTTTTGCTATTCATTGAAAGCAGTAGTGACTGAT
TTGTATATA
>MinSyn_1748|Strength:0.000394079
GCGTGTCGTTTTAGTGAGGGTGGGAGCCACCAACCGATGCTGATCTTGCCTGCCTTGATGATTTATGA
CCCCCGCCGATGACGCGGGAGCTGATCTTGCTATATAAGGTTTTGCTATTCATTGAAAGCAGTAGTGA
CTGATTTGTATATA
>MinSyn_1786|Strength:0.000395566
GCGTGTCGTTTTAGTGAGGTTATGACCCCCGCCGATGACGCGGGACCGATGCTGATCTTGCCTGCCTT
GATGATGAAGCATCTTCCCCTCTACAAAAGTGGGAGCCACCATGCTAGGAGGACCGATGCTATATAAG

GTTTTGCTATTCATTGAAAGCAGTAGTGACTGATTTGTATATA
>MinSyn_1607|Strength:0.000395696
GCGTGTCGTTTTAGTGAGGATTGCGATAAAGGAAAGGTTTTTCAACAAGCCCAATTAGGTTGTCTGCA
CCTTCACTATCAGCGTCTGCACCTCCTATATAAGGTTTTGCTATTCATTGAAAGCAGTAGTGACTGAT
TTGTATATA
>MinSyn_1582|Strength:0.000395822
GCGTGTCGTTTTAGTGAGGAGCAAGTGGATAGGTCACGCACACCAGCATGTGTTGATCACCAGCTGGT
AGGTCACGACCACTATGCCCAATTAGGTTTTCAACAAGGTACTTGTGTACAGGGCTCACTGCTTATGA
CCCCCGCCGATGACGCGGGAGATGCTGATCTTGCCTGCCTTGATAATTTCGGGAAACCTCCTCGTGTA
CAGGGCTCACTGCCTATATAAGGTTTTGCTATTCATTGAAAGCAGTAGTGACTGATTTGTATATA
>MinSyn_1729|Strength:0.000397165
GCGTGTCGTTTTAGTGAGGTGGTGGAGCACGACACTACAAAACTGGTACTTGTGTACAGGGAACCATT
ATTGCGCGACCACTATGCCCAATTAGGTTAGGTGGCTCCTACACCGATGCTGATCTAAAAATGTCAAA
GATACTAGGAGGACCGATGCTGATCTTGCCTGCCGCACACCAGCATGTGTTGATCACCAGCTGGTTGT
CTGCACCTCAGTGGGAGCCACCAAGACCTCTACAAAACTGGTACTTGTGTACATCACTATCAGCCTAC
AAAACTTTTTCAACAAGTAGGCTATATAAGGTTTTGCTATTCATTGAAAGCAGTAGTGACTGATTTGT
ATATA
>MinSyn_168|Strength:0.000397619
GCGTGTCGTTTTAGTGAGGGTGGGAGCCACCAGATGCTGATCTTGCCTGCATTGCGATAAAGGAAAGG
CACTGCTAGGAGGACCGATCCTTACCGCTATGGGTAAGATTTAGGTTGTCTGCACAACCATTATTGCG
CGACCATGGTGGAGCACGACACACTGCTAGGAGGACCGATGCTGATCTTGCCTATATAAGGTTTTGCT
ATTCATTGAAAGCAGTAGTGACTGATTTGTATATA
>MinSyn_1954|Strength:0.000397828
GCGTGTCGTTTTAGTGAGGAATTTCGGGAAACCTCCTCGACCACTATTTTTCAACAAATGCTGATCTT
GCCTGCCTAGGTGGCTCCTACGGCTCACTAGCAAGTGGATCTCTACAAAACTGGTACTCTATATAAGG
TTTTGCTATTCATTGAAAGCAGTAGTGACTGATTTGTATATA
>MinSyn_145|Strength:0.00039851
GCGTGTCGTTTTAGTGAGGAGCAAGTGGATAACTGGTACTTGTGTACAGGGCATTGCGATAAAGGAAA
GGAAGACCTCTACAAAACTGGTACTTCAGCCACTTGTGTGGAGGACCGATGCTGATCTTGCCTGCCTT
GTCCATCAACAAATAATCCAAGTAAGGACCTCTAACCACGTCTACAAGACCGATGCTGATCTTGCCTG
CCTTATGACCCCCGCCGATGACGCGGGACAATTAGGTTGTCTGCACCTCACAAACCATTATTGCGGGT
CACGACCACTATGCCCTATATAAGGTTTTGCTATTCATTGAAAGCAGTAGTGACTGATTTGTATATA
>MinSyn_1839|Strength:0.00039859
GCGTGTCGTTTTAGTGAGGTCCATCAACAAATAATCCAAGTAAGCACTGCTAGGAGGACCGATGCTGA
TTCAGAAGATCAAAGGGCTAACCTCACATGTAGGCTATCAGCTTAGCAAGAACCATTATTGCGTGTGG
GAGCCACCACACGTCACTATCAGCTACTTGTGTACAGGGCTCACTGCTAGCAGCCACTTGTGTGGGCT
CACTGCTAGGAGGACCGATGCTGATTCTCTCTGCCGACAGTGGTCCCAAACACCTCACACAGTGGTCC
CTCCACACTATGCCCAATTAGGTTGTCTGCACCTCTATATAAGGTTTTGCTATTCATTGAAAGCAGTA
GTGACTGATTTGTATATA
>MinSyn_1801|Strength:0.000399579
GCGTGTCGTTTTAGTGAGGGTGGGAGCCACCAAGGACCGATGCTGATCTTGCCTGGCACACCAGCATG
TGTTGATCACCAGCTCTATGCCCAATTAGGTTGTCTGCACCTAACCACGTCTACAATAGGCTATCAGC
TTAGCAGGTGGCTCCTACTGATGAAAAATGTCAAAGATAACTGCTAGGAGGACCGATGCTGATTATGA
CCCCCGCCGATGACGCGGGACTAGGAGGACCGATGCTGATCTTGCCTGTCACTATCAGCCATGTAGGC
TATCAGCTTAGCAAAACCATTATTGCGGTGTACAGGGCTCACTATATAAGGTTTTGCTATTCATTGAA
AGCAGTAGTGACTGATTTGTATATA
>MinSyn_1705|Strength:0.000400042
GCGTGTCGTTTTAGTGAGGCAGCCACTTGTGTGCTGATCTTGCCTGCCTTATTGCGATAAAGGAAAGG
TAGCAAGACCTCTACAAAACTGGTACTTGAACCATTATTGCGCATGTAGGCAACCACGTCTACAAGTA
CAGGGCTCACTGAGCAAGTGGATCAAGACCTCCTATATAAGGTTTTGCTATTCATTGAAAGCAGTAGT
GACTGATTTGTATATA
>MinSyn_185|Strength:0.000400853
GCGTGTCGTTTTAGTGAGGTTATGACCCCCGCCGATGACGCGGGATCTACAAAACTGGTACATCCTTA
CCGCTATGGGTAAGATTTAGGAGGACCGATGCTGATCTTGCCTGCCAGTGGTCCCTCCACGCCCAATT

AGGTTGTCTGCACCTCAATTGCGATAAAGGAAAGGACTGGTACTTGTGTACAGAGCAAGTGGATGCTA
GGAGGACCGACTATATAAGGTTTTGCTATTCATTGAAAGCAGTAGTGACTGATTTGTATATA
>MinSyn_1257|Strength:0.000401923
GCGTGTCGTTTTAGTGAGGTCTCTCTGCCGACAGTGGTCCCAAACACATGTAGGCTAAATTTCGGGAA
ACCTCCTCGGGCTCACTGCTAGGAGGACCGATCTATATAAGGTTTTGCTATTCATTGAAAGCAGTAGT
GACTGATTTGTATATA
>MinSyn_1225|Strength:0.000402301
GCGTGTCGTTTTAGTGAGGGGAAAAAGAAGAGGTGTAGGCTATCAGCTTTATGACCCCCGCCGATGAC
GCGGGAACTGGTACTTGTGTACAGGGCTCAATTGCGATAAAGGAAAGGCTGCACCTCACATGTAGGCT
TCTCTCTGCCGACAGTGGTCCCAAACGGTAGGTCACGACCACTATGCCCAATTCTATATAAGGTTTTG
CTATTCATTGAAAGCAGTAGTGACTGATTTGTATATA
>MinSyn_1921|Strength:0.000402616
GCGTGTCGTTTTAGTGAGGATCCTTACCGCTATGGGTAAGATTCCTCTACAAAAACCACGTCTACAAG
GTACTTGTGTACAGGGCAATTTCGGGAAACCTCCTCGGGTGCACACCAGCATGTGTTGATCACCAGCT
CCTCTACAAAACTGGTACTTGTGTACAGATCGAAAGGACAGTATATGCCCAATTAGGTTGTCTGCAAA
CCATTATTGCGTGCCTGCCTTGATGATATGAAGCATCTTCCCTGATCTTGCCTGCCGGAAAAAGAAGA
GGTTCTGCACCTCACATGTAGGCTATCACTATATAAGGTTTTGCTATTCATTGAAAGCAGTAGTGACT
GATTTGTATATA
>MinSyn_1902|Strength:0.000402621
GCGTGTCGTTTTAGTGAGGTCCATCAACAAATAATCCAAGTAAGTGCCCAAGCACACCAGCATGTGTT
GATCACCAGCTCTATCAGCTTAGCAAGACCTCTCTCTCTGCCGACAGTGGTCCCAAACAGGGCTCACT
GCTAGGAGGAATCGAAAGGACAGTAATCAGCTTAGCAAGACCTCTACAAAACTGCTATATAAGGTTTT
GCTATTCATTGAAAGCAGTAGTGACTGATTTGTATATA
>MinSyn_1122|Strength:0.000402748
GCGTGTCGTTTTAGTGAGGTGAAGCATCTTCCGCGGTAGGTCACGACCAGTGGTCCCTCCACTGTACA
GGGCTCACTGCAAAAATGTCAAAGATACACCTCGTGGGAGCCACCACTATGCCCAATTAGGTTGTCTG
CACCTCTATATAAGGTTTTGCTATTCATTGAAAGCAGTAGTGACTGATTTGTATATA
>MinSyn_1220|Strength:0.000402781
GCGTGTCGTTTTAGTGAGGAGGTGGCTCCTACGACCCAGTGGTCCCTCCACAGCAAGACCTCTACAAA
ACTGGTACTCAAATATTTCTTGTGACCTCTACAAAACTGGTTCACTATCAGCCTCACTGCTAGGAGGA
CCGATGCTCAGCCACTTGTGTCTGGTGGAGCACGACATCTTGCCTGCCTTGATGCTATATAAGGTTTT
GCTATTCATTGAAAGCAGTAGTGACTGATTTGTATATA
>MinSyn_1381|Strength:0.000403421
GCGTGTCGTTTTAGTGAGGGCACACCAGCATGTGTTGATCACCAGCTGTTGTCTGCACCTCACATGTA
GGAACCATTATTGCGATGCTGATCTTGCCTGCCTTGATATTGCGATAAAGGAAAGGCAGCTTAGCAAG
ACCTCTACAGCCACTTGTGTGTCTGCACCTCACATGTAGGCTAAAGATTGATGAAAAGTCAAAAACAA
AAATCAATTATGCCCAATTAGGTTGTCTGCGTGGGAGCCACCATCTGCACCTCACATGTAGGCTATCA
GCTATATAAGGTTTTGCTATTCATTGAAAGCAGTAGTGACTGATTTGTATATA
>MinSyn_1636|Strength:0.000404655
GCGTGTCGTTTTAGTGAGGATTGCGATAAAGGAAAGGCTCAGAAGATCAAAGGGCTAGACAACCATTA
TTGCGGACCACTATGCCCAATTAGGAATTTCGGGAAACCTCCTCGAGGTCACGACCACTATGCCCAAT
TAGGTTTCTCTCTGCCGACAGTGGTCCCAAATTAGCAAGACCTCTACAAAACTGGTACTTCTATATAA
GGTTTTGCTATTCATTGAAAGCAGTAGTGACTGATTTGTATATA
>MinSyn_1422|Strength:0.000404971
GCGTGTCGTTTTAGTGAGGAACCACGTCTACAACTGATCTTGCCTGCCTGCTTTGTCAAAAGCTAAAA
AAGATGATGCGGTACTTGTGTACAGGGCTCACTGCTAGGACTATATAAGGTTTTGCTATTCATTGAAA
GCAGTAGTGACTGATTTGTATATA
>MinSyn_1650|Strength:0.000405286
GCGTGTCGTTTTAGTGAGGTGAAGATAAGATAATAATGTTGAAGATAAGACTTAGCAAGACCTCTACA
AAACTGGATCCTTACCGCTATGGGTAAGATTTACAGGGCTCACTGCTAGGAGGACCGAATCGAAAGGA
CAGTAATCTTGCCTCTATATAAGGTTTTGCTATTCATTGAAAGCAGTAGTGACTGATTTGTATATA
>MinSyn_1130|Strength:0.000406384
GCGTGTCGTTTTAGTGAGGGCACACCAGCATGTGTTGATCACCAGCTAGGGCTCACTGCTAGGAGGAC
CGATGAGGTGGCTCCTACTAGGTTGTCTGCACCTCACATGTAGGCTAACCACGTCTACAAACCTTCAG

AAGATCAAAGGGCTACACATGTAGGCTATCAGCTTAGCAAGACTATATAAGGTTTTGCTATTCATTGA
AAGCAGTAGTGACTGATTTGTATATA
>MinSyn_1313|Strength:0.00040645
GCGTGTCGTTTTAGTGAGGCAGTGGTCCCTCCACGTCTGCACCTCTCCATCAACAAATAATCCAAGTA
AGCAAGACCTCTGGAAAAAGAAGAGGTACTGCTAGGAGGACCTCAGAAGATCAAAGGGCTATACAAAA
CTGGTACTTGTGTACAGGGCTCTATATAAGGTTTTGCTATTCATTGAAAGCAGTAGTGACTGATTTGT
ATATA
>MinSyn_137|Strength:0.000407341
GCGTGTCGTTTTAGTGAGGTCCATCAACAAATAATCCAAGTAAGAAGACCTCTACAAAAAATGTCAAA
GATAGCTCACTGCTAGGAGGGTGGGAGCCACCATGCTAGGAGGACCGATGCTGATAACCACGTCTACA
ACCACTATGCCCAATTTCTCTCTGCCGACAGTGGTCCCAAAGCAGTGGTCCCTCCACCTAGGAGGACC
GATGCTGAAGCATCTTCCTAGGAGGACCGATGCTGATCTTGCCTGATTGCGATAAAGGAAAGGTAGGA
GGACCGATGCTGATCTTGATCGAAAGGACAGTACTTAGCAAGACCTCTACAAAACTGGCTATATAAGG
TTTTGCTATTCATTGAAAGCAGTAGTGACTGATTTGTATATA
>MinSyn_1959|Strength:0.000407585
GCGTGTCGTTTTAGTGAGGGCTTTGTCAAAAGCTAAAAAAGATGATGCATCAGTTATGACCCCCGCCG
ATGACGCGGGACTTGTGTACAGGGCTCACTGCTAGGAGGGAAAAAGAAGAGGTCCACTATGCCCAAGG
TGGCTCCTACAGGACCGATGCTGATCTTGCCTGTGGTGGAGCACGACAACCTCTACAAATATTTCTTG
TTTGTCTGCACCTCACATGTAGGCAAAAATGTCAAAGATATTGTGTACAGGGCTCACTGCTAGGAATC
CTTACCGCTATGGGTAAGATTCAATTAGGTTGTCTGTCACTATCAGCTAGGAGGACCGATGCTGATCT
TGCCCTATATAAGGTTTTGCTATTCATTGAAAGCAGTAGTGACTGATTTGTATATA
>MinSyn_1342|Strength:0.000408132
GCGTGTCGTTTTAGTGAGGAGGTGGCTCCTACCTGCACCTCACATGTTCCATCAACAAATAATCCAAG
TAAGGCTCACTGCTAGGAGGACCTATATAAGGTTTTGCTATTCATTGAAAGCAGTAGTGACTGATTTG
TATATA
>MinSyn_1124|Strength:0.000408161
GCGTGTCGTTTTAGTGAGGTTATGACCCCCGCCGATGACGCGGGATAGCAAGACCTCTACAAAACTGG
TACTTGTGGTGGAGCACGACATGCCCAATTAGGTTGTCTGCACCTCAGCCACTTGTGTGTCACAGCAA
GTGGATCAAGACCTCTATATAAGGTTTTGCTATTCATTGAAAGCAGTAGTGACTGATTTGTATATA
>MinSyn_1379|Strength:0.000408859
GCGTGTCGTTTTAGTGAGGTGAAGATAAGATAATAATGTTGAAGATAAGAGCGGTAGGTCACGACCAC
TATCGAAAGGACAGTACATGTAGGCTATCAGCATCCTTACCGCTATGGGTAAGATTCGCGGTAGGTCA
CGACCACTATGCCCAATCTATATAAGGTTTTGCTATTCATTGAAAGCAGTAGTGACTGATTTGTATAT
A
>MinSyn_1762|Strength:0.000410888
GCGTGTCGTTTTAGTGAGGTGAAGCATCTTCCCTTGAAGATAAGATAATAATGTTGAAGATAAGAGCG
GTAGGTCACGACCACTATGCCCAATTGCACACCAGCATGTGTTGATCACCAGCTGTATTGCGATAAAG
GAAAGGGGCTATCAGCTTAGCAAGATTTTCAACAACTGCTAGGAGGACCGTGGTGGAGCACGACAAGA
CCTCTACAAAACTGGTACTAGGTGGCTCCTACTCACGACCACTATGCCCAATTAGGCTATATAAGGTT
TTGCTATTCATTGAAAGCAGTAGTGACTGATTTGTATATA
>MinSyn_1513|Strength:0.0004118
GCGTGTCGTTTTAGTGAGGAACCACGTCTACAATTAGCAAGACCATCGAAAGGACAGTAAGGTTGTCT
GCACCTCACATGTAGGAAAAAGAAGAGGTCTGATCTTGCCTCCATCAACAAATAATCCAAGTAAGACC
TCACATGTAGGCTATCAAATTTCGGGAAACCTCCTCGTGAACCATTATTGCGGCTCACTGCTAGTGAA
GCATCTTCCGCACCTCACATGTAGGCTATCAGCCAAATATTTCTTGTCTTGTGTACAGGGCTCACTGC
TAGGAGCTATATAAGGTTTTGCTATTCATTGAAAGCAGTAGTGACTGATTTGTATATA
>MinSyn_1358|Strength:0.000412011
GCGTGTCGTTTTAGTGAGGAATTTCGGGAAACCTCCTCGTTAGCAAGACCTCTACATTTTCAACAAAA
GACCTCTACAAAACCAGCCACTTGTGTACCGATGCTGGCTTTGTCAAAAGCTAAAAAAGATGATGCCT
CAGAAGATCAAAGGGCTATATCAGCTTAGCAAGACCTCTACAAAAGGTGGCTCCTACCGCGGTAGGTC
ACGACCACTATGCCCACAGTGGTCCCTCCACCAGGAACCATTATTGCGCACGACCACTATGCCCAATT
AGGTTTGAAGCATCTTCCGTTGTCTGCACCTCACATGTCTATATAAGGTTTTGCTATTCATTGAAAGC
AGTAGTGACTGATTTGTATATA
>MinSyn_1769|Strength:0.000412194

GCGTGTCGTTTTAGTGAGGAATTTCGGGAAACCTCCTCGGTGTACAGGGCTCACTGCTAGGAGGGCAC
ACCAGCATGTGTTGATCACCAGCTAGGTCACGACCACTTCAGAAGATCAAAGGGCTAAGCAAGACCTC
TACAAAACTGGTCTATATAAGGTTTTGCTATTCATTGAAAGCAGTAGTGACTGATTTGTATATA
>MinSyn_1644|Strength:0.000412312
GCGTGTCGTTTTAGTGAGGGGAAAAAGAAGAGGTGTCACGACCACCAGTGGTCCCTCCACTAGGAGGA
CCGATGCTGATCTGTGGGAGCCACCAACCTCTTCTCTCTGCCGACAGTGGTCCCAAAACCACTATGCC
CAATGCACACCAGCATGTGTTGATCACCAGCTCGATGCTGATCTTGCCTGCCTCAGCCACTTGTGTGC
TAGGAGGACCGATGCTGATCTTGCCTGCTATATAAGGTTTTGCTATTCATTGAAAGCAGTAGTGACTG
ATTTGTATATA
>MinSyn_1929|Strength:0.000412994
GCGTGTCGTTTTAGTGAGGAGCAAGTGGATAGAAGATTGATGAAAAGTCAAAAACAAAAATCAATTAT
AGACCTCTACAAAACTGGTACTTGTGTACTCTCTCTGCCGACAGTGGTCCCAAAGGACCGATGCTGAT
CTTGCCTGCCAGCCACTTGTGTCTTGTGTACAGGGATTGCGATAAAGGAAAGGCGGTAGGTCACGAAA
CCATTATTGCGACGACCACTATGCCCAATTAGGTTGCTATATAAGGTTTTGCTATTCATTGAAAGCAG
TAGTGACTGATTTGTATATA
>MinSyn_1477|Strength:0.000413378
GCGTGTCGTTTTAGTGAGGAGGTGGCTCCTACGACCACTATGCCCAATTAGGTTGTTGGTGGAGCACG
ACAACATGTAGGCTATCAGCTTGGAAAAAGAAGAGGTCACGACCACTATGCCCAATCGAAAGGACAGT
ACAAGACCCTATATAAGGTTTTGCTATTCATTGAAAGCAGTAGTGACTGATTTGTATATA
>MinSyn_16|Strength:0.000414286
GCGTGTCGTTTTAGTGAGGTGAAGCATCTTCCCTATCAGCTCAAATATTTCTTGTGACCACTATGCCC
AATTAGGTTCTATATAAGGTTTTGCTATTCATTGAAAGCAGTAGTGACTGATTTGTATATA
>MinSyn_1330|Strength:0.000414286
GCGTGTCGTTTTAGTGAGGGTGGGAGCCACCAGTCAAATATTTCTTGTATGTAGGCTATCAGCTTAGC
AAGACCTCTCTATATAAGGTTTTGCTATTCATTGAAAGCAGTAGTGACTGATTTGTATATA
>MinSyn_1568|Strength:0.000415048
GCGTGTCGTTTTAGTGAGGCAGTGGTCCCTCCACGTCACGACCACTATGAAAAATGTCAAAGATACTC
TACAAAACTGGTACTTGTAATTTCGGGAAACCTCCTCGACAAAACTGGTACTTGTGTACAGGCAAATA
TTTCTTGTTGTCTGCACCTCACATGTAATCGAAAGGACAGTAGTAGGTCCTATATAAGGTTTTGCTAT
TCATTGAAAGCAGTAGTGACTGATTTGTATATA
>MinSyn_1646|Strength:0.000415132
GCGTGTCGTTTTAGTGAGGCAAATATTTCTTGTGTTGTCTGCACCTCACATGTAAAAAATGTCAAAGA
TACGCGGTAGGTCGCACACCAGCATGTGTTGATCACCAGCTAAAACTGGTACTTGTGTACAGGGCTAT
CCTTACCGCTATGGGTAAGATTATTAGGTTGTCTGCACCTCACATGTCTATATAAGGTTTTGCTATTC
ATTGAAAGCAGTAGTGACTGATTTGTATATA
>MinSyn_1596|Strength:0.000415649
GCGTGTCGTTTTAGTGAGGGTGGGAGCCACCAGCTAGGAGGACCGATGCTGATCTTGCCATCCTTACC
GCTATGGGTAAGATTCGATGCTGATCTTAACCATTATTGCGATTAGGTTGAAGCATCTTCCGACCTCT
ACAAAACTGGTACTTCTATATAAGGTTTTGCTATTCATTGAAAGCAGTAGTGACTGATTTGTATATA
>MinSyn_1102|Strength:0.000416871
GCGTGTCGTTTTAGTGAGGTGAAGATAAGATAATAATGTTGAAGATAAGACTCACATGTAGGCAAATA
TTTCTTGTAGGCTATCAGCTTAGCAAGACTTATGACCCCCGCCGATGACGCGGGAACTTGTGTACAAT
CGAAAGGACAGTATTGTGTACAGGGCTCACTGCTAGGAGGACTATATAAGGTTTTGCTATTCATTGAA
AGCAGTAGTGACTGATTTGTATATA
>MinSyn_1649|Strength:0.000417073
GCGTGTCGTTTTAGTGAGGTTTTCAACAAAGGGCTCCAGTGGTCCCTCCACTCTACAAAACTGGTACC
TATATAAGGTTTTGCTATTCATTGAAAGCAGTAGTGACTGATTTGTATATA
>MinSyn_1776|Strength:0.000417106
GCGTGTCGTTTTAGTGAGGTCACTATCAGCGGTACTTGTGTACAGGGCTCAGCAAGTGGATTGCAGCC
ACTTGTGTTGGTACTTGTGATCGAAAGGACAGTATGATCTTGCCTGCCTTGATGATTTTCAACAATTG
TGTACTATATAAGGTTTTGCTATTCATTGAAAGCAGTAGTGACTGATTTGTATATA
>MinSyn_1297|Strength:0.000418307
GCGTGTCGTTTTAGTGAGGAACCACGTCTACAACATGTAGGCTAATCGAAAGGACAGTATGCACCTCA
CATGTAGGCGTGGGAGCCACCACACATGTAGGCGCTTTGTCAAAAGCTAAAAAAGATGATGCGCAAGA

CCTCTACAAAACTGGTACTTGTGTCTATATAAGGTTTTGCTATTCATTGAAAGCAGTAGTGACTGATT
TGTATATA
>MinSyn_1221|Strength:0.000418533
GCGTGTCGTTTTAGTGAGGATTGCGATAAAGGAAAGGGGTTGTCTGCACCTCACATGTAGGCCAGCCA
CTTGTGTACTATTTTTCAACAAGCTATCAGCTTAGCAAGACAAATATTTCTTGTTACTTGTGTACAGG
GCTCATCTCTCTGCCGACAGTGGTCCCAAAGTCTGCACCTCACATGTAGGCTATCAGTCAGAAGATCA
AAGGGCTACCTCTACAACTATATAAGGTTTTGCTATTCATTGAAAGCAGTAGTGACTGATTTGTATAT
A
>MinSyn_1706|Strength:0.000419038
GCGTGTCGTTTTAGTGAGGCAGCCACTTGTGTCTCAGAAGATCAAAGGGCTAGTAGGCTATCAGCTTA
GCAAGACCTCTGCTTTGTCAAAAGCTAAAAAAGATGATGCACATGTAGGAATTTCGGGAAACCTCCTC
GAGCAAGACGTGGGAGCCACCACAAGACAAATATTTCTTGTTGTGTACAGGGCTCACTGCTAGGAGGA
AGGTGGCTCCTACGCTTAGCAAGACCTCTACAAAATCGAAAGGACAGTACAAGACCTCTACAAAACTG
GTACCTATATAAGGTTTTGCTATTCATTGAAAGCAGTAGTGACTGATTTGTATATA
>MinSyn_1560|Strength:0.000419042
GCGTGTCGTTTTAGTGAGGTGAAGCATCTTCCCTGGTACTTGTGTACAGGGCTCTCTCTCTGCCGACA
GTGGTCCCAAAAGGTCACGCTATATAAGGTTTTGCTATTCATTGAAAGCAGTAGTGACTGATTTGTAT
ATA
>MinSyn_1676|Strength:0.000419042
GCGTGTCGTTTTAGTGAGGTGAAGCATCTTCCGTTTTCAACAACACTGCTAGGAGGACCGATGCTGAT
CAAATATTTCTTGTCTCACCTATATAAGGTTTTGCTATTCATTGAAAGCAGTAGTGACTGATTTGTAT
ATA
>MinSyn_143|Strength:0.000419131
GCGTGTCGTTTTAGTGAGGATCCTTACCGCTATGGGTAAGATTCTCTACAAAACTGGTACTTGTGTCC
ATCAACAAATAATCCAAGTAAGGGCTCACTGCTAGGAGGACCTATATAAGGTTTTGCTATTCATTGAA
AGCAGTAGTGACTGATTTGTATATA
>MinSyn_150|Strength:0.000419499
GCGTGTCGTTTTAGTGAGGTCACTATCAGCGGACCGATGCTGATCTTGCCTGCCTTCCATCAACAAAT
AATCCAAGTAAGGTCCTATATAAGGTTTTGCTATTCATTGAAAGCAGTAGTGACTGATTTGTATATA
>MinSyn_195|Strength:0.000420504
GCGTGTCGTTTTAGTGAGGGGAAAAAGAAGAGGTATGCCCAATTAGGTTGTCTGCACCTCACATATTG
CGATAAAGGAAAGGACCTCACATGTAGGCTCAGCCACTTGTGTACCACCTATATAAGGTTTTGCTATT
CATTGAAAGCAGTAGTGACTGATTTGTATATA
>MinSyn_1393|Strength:0.000420653
GCGTGTCGTTTTAGTGAGGTGAAGATAAGATAATAATGTTGAAGATAAGACGACCAGCAAGTGGATTC
AAATATTTCTTGTGCTGATCTTGGCTTTGTCAAAAGCTAAAAAAGATGATGCTATGCAACCACGTCTA
CAACACCTCACATGTAGGCTATCAGCTTAGAACCATTATTGCGCTAGGAGGACCGATGCTGATCTTGC
CTAGGTGGCTCCTACATGCTGATCCAGTGGTCCCTCCACTAGGAGGACCGATGCTGATCTTGCCTGCT
ATATAAGGTTTTGCTATTCATTGAAAGCAGTAGTGACTGATTTGTATATA
>MinSyn_197|Strength:0.000420941
GCGTGTCGTTTTAGTGAGGCAGTGGTCCCTCCACGACCTCTACAAATCCTTACCGCTATGGGTAAGAT
TAGGTTGTCTGCACCTCACCTATATAAGGTTTTGCTATTCATTGAAAGCAGTAGTGACTGATTTGTAT
ATA
>MinSyn_1714|Strength:0.000422424
GCGTGTCGTTTTAGTGAGGAAGATTGATGAAAAGTCAAAAACAAAAATCAATTATCCGATGCTGATCT
TGAAGCATCTTCCATGCCCAATTAGGTTGTCTGCACCTCAAGGTGGCTCCTACATGTAGGCTATCAGC
TTAGCATCCTTACCGCTATGGGTAAGATTTACTTGTGTACAGGGCCTATATAAGGTTTTGCTATTCAT
TGAAAGCAGTAGTGACTGATTTGTATATA
>MinSyn_1112|Strength:0.000422666
GCGTGTCGTTTTAGTGAGGATTGCGATAAAGGAAAGGCTGCACCTCACATGTAGGCTATCATCAGAAG
ATCAAAGGGCTAGCAAACCATTATTGCGTCTGCACCTCACATGTAGGCTATCAGCTTTTCAACAAACA
GGGCTCACTGCTAGCTTTGTCAAAAGCTAAAAAAGATGATGCACCTCACATGTAGGCAAATATTTCTT
GTCAAAACTGGTACTTCACTATCAGCAACTGGTACTTGTGTACAGGGCTCCTATATAAGGTTTTGCTA
TTCATTGAAAGCAGTAGTGACTGATTTGTATATA

>MinSyn_1967|Strength:0.000423182
GCGTGTCGTTTTAGTGAGGATTGCGATAAAGGAAAGGCAGGGCTCACTGCACACCAGCATGTGTTGAT
CACCAGCTTACAAAACTGGTACTTGTGTACAGGGCTCACTATATAAGGTTTTGCTATTCATTGAAAGC
AGTAGTGACTGATTTGTATATA
>MinSyn_1123|Strength:0.000423327
GCGTGTCGTTTTAGTGAGGGCACACCAGCATGTGTTGATCACCAGCTAGGCTATCAGCTTAGCAACCA
TTATTGCGCACCTCACATGTAGGCTATATCCTTACCGCTATGGGTAAGATTGCTCACTGCTAGGAGGA
CCGATGTGGTGGAGCACGACACCTGCCTTGTTATGACCCCCGCCGATGACGCGGGATACAGGGCTCGG
AAAAAGAAGAGGTGTCTGCACCTCACATGTAGTCAGAAGATCAAAGGGCTAGGAGGACCGATGCTGAT
CTAGCAAGTGGATTAGGCTATCAGCTTAGCAAGACCTCTACCTATATAAGGTTTTGCTATTCATTGAA
AGCAGTAGTGACTGATTTGTATATA
>MinSyn_1121|Strength:0.000425261
GCGTGTCGTTTTAGTGAGGAGCAAGTGGATGCCTGCCTTTTTTCAACAACAGCTTAGGCTTTGTCAAA
AGCTAAAAAAGATGATGCTTAGGTTGTCTGCACCTCACATGTAGGCTAAGGTGGCTCCTACCAGCTTA
GCAAGCACACCAGCATGTGTTGATCACCAGCTTGTACAGGGCTCACTGCTAGGAGGACCGACTATATA
AGGTTTTGCTATTCATTGAAAGCAGTAGTGACTGATTTGTATATA
>MinSyn_1389|Strength:0.0004256
GCGTGTCGTTTTAGTGAGGATCGAAAGGACAGTAGTTGTCTGCAGCAAGTGGATGGTCACGACCACTA
TCTATATAAGGTTTTGCTATTCATTGAAAGCAGTAGTGACTGATTTGTATATA
>MinSyn_1499|Strength:0.000425928
GCGTGTCGTTTTAGTGAGGGTGGGAGCCACCATAGGCTATGAAGATAAGATAATAATGTTGAAGATAA
GATCACGACCACTATGCCCAATTATTATGACCCCCGCCGATGACGCGGGAACCTCACATGTAGGCTAT
CAGCTTAGCAAATTGCGATAAAGGAAAGGCAATTAGGTTGTCTGCACCTCACATGTCCATCAACAAAT
AATCCAAGTAAGCGCGGTAGGTCACGACCACCTATATAAGGTTTTGCTATTCATTGAAAGCAGTAGTG
ACTGATTTGTATATA
>MinSyn_1977|Strength:0.000427144
GCGTGTCGTTTTAGTGAGGGCTTTGTCAAAAGCTAAAAAAGATGATGCTAGGCTATCAGCTTAGTCCA
TCAACAAATAATCCAAGTAAGACCTCACATGTAGGCTATCAGCTTAGCACTATATAAGGTTTTGCTAT
TCATTGAAAGCAGTAGTGACTGATTTGTATATA
>MinSyn_1259|Strength:0.000427174
GCGTGTCGTTTTAGTGAGGAAGATTGATGAAAAGTCAAAAACAAAAATCAATTATGCTAGGAGGACCG
ATGGCTTTGTCAAAAGCTAAAAAAGATGATGCAAAACTGGTACTTGTGTACAGTGGTGGAGCACGACA
GATGCTGATCTTGCCTGCCTTGATGAGCACACCAGCATGTGTTGATCACCAGCTTCACGACCACTATG
CCCCAGCCACTTGTGTCAAGACCTCTACAAAACTGGTACTTCACTATCAGCGTTGTCTGCACCTCACA
TGTAGGCTAATCGAAAGGACAGTAGTAGGTCACTATATAAGGTTTTGCTATTCATTGAAAGCAGTAGT
GACTGATTTGTATATA
>MinSyn_1832|Strength:0.000427205
GCGTGTCGTTTTAGTGAGGTCTCTCTGCCGACAGTGGTCCCAAAAGCTTAGCTTTTCAACAATAAACC
ATTATTGCGACATGTAGGCTCAAATATTTCTTGTGCCTGCCTTGATGGCACACCAGCATGTGTTGATC
ACCAGCTCTCACTGCTAGGAGTCACTATCAGCGACCTCTACAAAACTGGTACTTGTGTACATCCATCA
ACAAATAATCCAAGTAAGAGGAGGACCGATGCTGATCTTGCCTGCCTTAGCAAGTGGATTCACGACCA
CTATGCCTTATGACCCCCGCCGATGACGCGGGATCACATGTAGGCTATCAGCTTAGCAACTATATAAG
GTTTTGCTATTCATTGAAAGCAGTAGTGACTGATTTGTATATA
>MinSyn_1994|Strength:0.000428997
GCGTGTCGTTTTAGTGAGGTGGTGGAGCACGACAATCAGCTTAGCAAGACCTCTACAAATCGAAAGGA
CAGTATGATCTTGCCTGCCTTGATGATATCAGAAGATCAAAGGGCTAAGGTCACGACCACTATGCCCG
CTTTGTCAAAAGCTAAAAAAGATGATGCTGCTAGGAGGACCGACTATATAAGGTTTTGCTATTCATTG
AAAGCAGTAGTGACTGATTTGTATATA
>MinSyn_1193|Strength:0.000429735
GCGTGTCGTTTTAGTGAGGTGGTGGAGCACGACAGGCTATCAGCTTAGCAAGACCTCTACAAGCTTTG
TCAAAAGCTAAAAAAGATGATGCTTGTGTACAGGCTATATAAGGTTTTGCTATTCATTGAAAGCAGTA
GTGACTGATTTGTATATA
>MinSyn_1199|Strength:0.000429735
GCGTGTCGTTTTAGTGAGGGGAAAAAGAAGAGGTTCACATGTAGGCTATTGGTGGAGCACGACACTTG

CCTGCCTTTGAAGCATCTTCCACGACCACTATGCCTATATAAGGTTTTGCTATTCATTGAAAGCAGTA
GTGACTGATTTGTATATA
>MinSyn_1201|Strength:0.000430205
GCGTGTCGTTTTAGTGAGGTTATGACCCCCGCCGATGACGCGGGATAGATTGCGATAAAGGAAAGGTG
TAGGCTATCAGCTTAGTCCATCAACAAATAATCCAAGTAAGGGACCGAAACCATTATTGCGCCGATGC
TGATCTTGCCTGCGGAAAAAGAAGAGGTCACCTCACATGTAGGCTATCAGCCTATATAAGGTTTTGCT
ATTCATTGAAAGCAGTAGTGACTGATTTGTATATA
>MinSyn_1322|Strength:0.000430613
GCGTGTCGTTTTAGTGAGGTTTTCAACAATACAGGGCTCACTGCTACAGCCACTTGTGTTCTACAAAA
CTGGTACTTGTGTACTCAGAAGATCAAAGGGCTATAGGTCACGACCACTATGCCCAATTGCTTTGTCA
AAAGCTAAAAAAGATGATGCTACAAAACTGGTACTTGTGTACAGGGCTCAAAAAAATGTCAAAGATAAC
TTGTGTACAGGGCTCACTAATTTCGGGAAACCTCCTCGCAGCTTAGCAAGACCTCAAGATTGATGAAA
AGTCAAAAACAAAAATCAATTATGCTTAGCAAGACCTCTACAAAACAACCACGTCTACAAAGGACCGA
TGCTGATCTTGCCTCTATATAAGGTTTTGCTATTCATTGAAAGCAGTAGTGACTGATTTGTATATA
>MinSyn_1626|Strength:0.000431092
GCGTGTCGTTTTAGTGAGGGTGGGAGCCACCACACGACCACTATGCCCAGCTTTGTCAAAAGCTAAAA
AAGATGATGCTCACATGTAGGCTATCAGCTCTATATAAGGTTTTGCTATTCATTGAAAGCAGTAGTGA
CTGATTTGTATATA
>MinSyn_1612|Strength:0.000431136
GCGTGTCGTTTTAGTGAGGAAAAATGTCAAAGATAGGTACTTGTGTACAAGATTGATGAAAAGTCAAA
AACAAAAATCAATTATGTACTTGTGTACAGGGCTCACTGCTAGGATCGAAAGGACAGTATATCAGCTT
ACTATATAAGGTTTTGCTATTCATTGAAAGCAGTAGTGACTGATTTGTATATA
>MinSyn_1852|Strength:0.000432025
GCGTGTCGTTTTAGTGAGGAACCATTATTGCGAGACCTCTACAAAACTGGTACTTGTCAAATATTTCT
TGTACTGCTAGGAGGACCTGGTGGAGCACGACAACCACTGGAAAAAGAAGAGGTTGCACCTCACATGT
ATTATGACCCCCGCCGATGACGCGGGAGCTATCAGCTTAGCAAGACCTCTCTATATAAGGTTTTGCTA
TTCATTGAAAGCAGTAGTGACTGATTTGTATATA
>MinSyn_1685|Strength:0.000432266
GCGTGTCGTTTTAGTGAGGTTTTCAACAACACCTCACATGTAGAACCATTATTGCGCGACCACTATGC
CAGGTGGCTCCTACCTACTATATAAGGTTTTGCTATTCATTGAAAGCAGTAGTGACTGATTTGTATAT
A
>MinSyn_1737|Strength:0.000432481
GCGTGTCGTTTTAGTGAGGTTTTCAACAAGACCACTATGCCATCCTTACCGCTATGGGTAAGATTAAG
GTGGCTCCTACTCACGACCACTATGCCCAATTAGGTTGTCGCACACCAGCATGTGTTGATCACCAGCT
CACGACCACTATGCCCAATTAGGCTATATAAGGTTTTGCTATTCATTGAAAGCAGTAGTGACTGATTT
GTATATA
>MinSyn_1490|Strength:0.000432892
GCGTGTCGTTTTAGTGAGGAACCATTATTGCGCAAGACCTCTACAAAACAAAAATGTCAAAGATAACC
TCTACAAAACTGGTACTTGTGTAGGTGGCTCCTACCCAATTAGGTTGTCTGCACCTCATCCTTACCGC
TATGGGTAAGATTTTGCCTGCCTTGATGTGAAGATAAGATAATAATGTTGAAGATAAGAACCGATGCT
GATCTTGCCTGCCTTGACTATATAAGGTTTTGCTATTCATTGAAAGCAGTAGTGACTGATTTGTATAT
A
>MinSyn_1488|Strength:0.000433088
GCGTGTCGTTTTAGTGAGGAGGTGGCTCCTACCGGTAGGTCACGACCACTATGCCCAATTAGAAAAAT
GTCAAAGATAAGCTATATAAGGTTTTGCTATTCATTGAAAGCAGTAGTGACTGATTTGTATATA
>MinSyn_1927|Strength:0.000433566
GCGTGTCGTTTTAGTGAGGTGGTGGAGCACGACAGTGAACCATTATTGCGAGGGCTCACGGAAAAAGA
AGAGGTGTTGTCTGCACCTCACACAAATATTTCTTGTACCGAAGGTGGCTCCTACCTGCACCTCACAT
GTAGGCTATCAGCTTACAGTGGTCCCTCCACCATTTTCAACAAAATTAGGTTGTCTGCACTATATAAG
GTTTTGCTATTCATTGAAAGCAGTAGTGACTGATTTGTATATA
>MinSyn_1157|Strength:0.000434341
GCGTGTCGTTTTAGTGAGGTGAAGCATCTTCCGCTAGGAGGACCGATGCTGGTGGGAGCCACCACATG
TAGGCTATCAGCAATTTCGGGAAACCTCCTCGTGCTAGGAGGACCGATGCTGATCTTGTGAAGATAAG
ATAATAATGTTGAAGATAAGACCACTATGCCCAATTAGGTTGTGGTGGAGCACGACAAAACTGGTACT

TGTATCGAAAGGACAGTATTGTGTACAGGGCTCACTGCTAGGAAGCAAGTGGATTGTAGGCTTTTTCA
ACAAGTTGGGAAAAAGAAGAGGTTGATCTTGCCTGCCTTGATGATCTATATAAGGTTTTGCTATTCAT
TGAAAGCAGTAGTGACTGATTTGTATATA
>MinSyn_1134|Strength:0.000434425
GCGTGTCGTTTTAGTGAGGAGGTGGCTCCTACACTGGTACTTGTGTACAGTCCATCAACAAATAATCC
AAGTAAGCTGCTAGGAGGACCGATGAAGATAAGATAATAATGTTGAAGATAAGAGGTAGGTCACGACC
ACTATGAGCAAGTGGATGTAGGCTATCAGCCAAATATTTCTTGTTAGGTTGTCTGCACCTCACATGCT
ATATAAGGTTTTGCTATTCATTGAAAGCAGTAGTGACTGATTTGTATATA
>MinSyn_1587|Strength:0.000435102
GCGTGTCGTTTTAGTGAGGAATTTCGGGAAACCTCCTCGAAACTGGTACTTGTGTACAGGGCTTCAGA
AGATCAAAGGGCTATGCATGGTGGAGCACGACAGTAGATTGCGATAAAGGAAAGGCCTCACATGTAGG
CTATCAGCTTAGCAAGATTTTCAACAACACTATGTTATGACCCCCGCCGATGACGCGGGAGTGTACAG
GGCTCACTGCTAGGAGGACCGCTATATAAGGTTTTGCTATTCATTGAAAGCAGTAGTGACTGATTTGT
ATATA
>MinSyn_1750|Strength:0.000435115
GCGTGTCGTTTTAGTGAGGATCGAAAGGACAGTAGCTAGGAGGACCGATGCTGATCAGGTGGCTCCTA
CAACTGGCTATATAAGGTTTTGCTATTCATTGAAAGCAGTAGTGACTGATTTGTATATA
>MinSyn_1241|Strength:0.000435566
GCGTGTCGTTTTAGTGAGGTGAAGATAAGATAATAATGTTGAAGATAAGATAGCTTTGTCAAAAGCTA
AAAAAGATGATGCTAAACCACGTCTACAACTTAGCAAGACCTCTACAAAACTGGTGGAGCACGACACA
AGACCTCTACAAATCACTATCAGCAAGACCTCTACAAAACTGGTACTTTTTTCAACAAGACCTCTACA
AAACTGGTACTTGTGTACAGGAAAAAGAAGAGGTGGCTATATAAGGTTTTGCTATTCATTGAAAGCAG
TAGTGACTGATTTGTATATA
>MinSyn_1916|Strength:0.000435691
GCGTGTCGTTTTAGTGAGGATCCTTACCGCTATGGGTAAGATTATGCCCAATTAGGAACCACGTCTAC
AACACTGCTAGGAGGACCGATGCTGAAGCAAGTGGATGGTAGGTCACGACCACTATGCCCAATTAGGG
CTTTGTCAAAAGCTAAAAAAGATGATGCCTACAAAACTGGTACTTATCGAAAGGACAGTAACCACTAT
GCGGAAAAAGAAGAGGTCAAAACTGGTACTTGTGTACAAATTTCGGGAAACCTCCTCGTATGCCCAAT
TAGGTTGTCTGCTCACTATCAGCGCTAGGAGGAAACCATTATTGCGCTTGCCTGCCTTGCTATATAAG
GTTTTGCTATTCATTGAAAGCAGTAGTGACTGATTTGTATATA
>MinSyn_1789|Strength:0.000436324
GCGTGTCGTTTTAGTGAGGTTATGACCCCCGCCGATGACGCGGGATAGGCTATCAGCTTAGCAATTGC
GATAAAGGAAAGGCTTAGCAAGACCTCTACAAAAAGCAAGTGGATGTACAGGGCTCCTATATAAGGTT
TTGCTATTCATTGAAAGCAGTAGTGACTGATTTGTATATA
>MinSyn_1414|Strength:0.000436428
GCGTGTCGTTTTAGTGAGGGCTTTGTCAAAAGCTAAAAAAGATGATGCCTAGGAGGACTGAAGATAAG
ATAATAATGTTGAAGATAAGACTATGCCCAATTAGGTTGTCTGCACCCAGTGGTCCCTCCACTTGTCT
GCACCTCACATGTAGGCTATCAAGCAAGTGGATGCAAGACCTCTTCACTATCAGCCGGCACACCAGCA
TGTGTTGATCACCAGCTTACTTGTGTACAGGGCTCACTGCTAAATTTCGGGAAACCTCCTCGCCTCAC
ATGTAGGCTATCAGCTATCCTTACCGCTATGGGTAAGATTTACAAAACTGGTACTTGTGTACAGGGCA
GCCACTTGTGTGCTGATCTTGCCTGCCTTCTATATAAGGTTTTGCTATTCATTGAAAGCAGTAGTGAC
TGATTTGTATATA
>MinSyn_1512|Strength:0.000436561
GCGTGTCGTTTTAGTGAGGTGAAGATAAGATAATAATGTTGAAGATAAGACTGCACCTCACATGTAGG
CTATTTTCAACAAGTAGGCTATCAGCTTTGTCAAAAGCTAAAAAAGATGATGCGTTGTCTGCACCTCA
CATGTAGGCTATCAGTTATGACCCCCGCCGATGACGCGGGATCGCGGTAGGTCACGACCACTACTATA
TAAGGTTTTGCTATTCATTGAAAGCAGTAGTGACTGATTTGTATATA
>MinSyn_1797|Strength:0.000437288
GCGTGTCGTTTTAGTGAGGCAGTGGTCCCTCCACACGACCACTATGCCCAATTAGGTTGTCATTGCGA
TAAAGGAAAGGGTAATCGAAAGGACAGTATGTACAGGGCTCACTGCTAGTCAGAAGATCAAAGGGCTA
CGACAGGTGGCTCCTACTTAGCAAGACCTCTACAAAACTGGTACTCTATATAAGGTTTTGCTATTCAT
TGAAAGCAGTAGTGACTGATTTGTATATA
>MinSyn_1442|Strength:0.000438006
GCGTGTCGTTTTAGTGAGGGCACACCAGCATGTGTTGATCACCAGCTGCAAGACCTCTAGGAAAAAGA

AGAGGTATCTTGCCTGCCAATTTCGGGAAACCTCCTCGGCCCAATTAGGTTGTCTGCACCTCACATGT
TTTTCAACAACTATGCCCAATTAGGTTGTCAAATATTTCTTGTTGATCTTGCCTGCCTTGATGATGAA
GCATCTTCCCACTATGCCCAATTAGGTTGTCTGCACTGAAGATAAGATAATAATGTTGAAGATAAGAA
GGGCCTATATAAGGTTTTGCTATTCATTGAAAGCAGTAGTGACTGATTTGTATATA
>MinSyn_110|Strength:0.000438462
GCGTGTCGTTTTAGTGAGGAGCAAGTGGATCTCTACAAAACTGGTACAAATATTTCTTGTCCTATATA
AGGTTTTGCTATTCATTGAAAGCAGTAGTGACTGATTTGTATATA
>MinSyn_1463|Strength:0.000438462
GCGTGTCGTTTTAGTGAGGAACCATTATTGCGAGACCTCTACAAAACTGTGGTGGAGCACGACATGGT
ACTTGTCTATATAAGGTTTTGCTATTCATTGAAAGCAGTAGTGACTGATTTGTATATA
>MinSyn_1565|Strength:0.00043893
GCGTGTCGTTTTAGTGAGGAGGTGGCTCCTACCTTAGCAAGACCTCTACAAAACTGTGGGAGCCACCA
GTCTGCACCATCGAAAGGACAGTAAGGACCGATGCTGATCTTGCCTGGGAAAAAGAAGAGGTGCTATA
TAAGGTTTTGCTATTCATTGAAAGCAGTAGTGACTGATTTGTATATA
>MinSyn_1240|Strength:0.000439125
GCGTGTCGTTTTAGTGAGGCAGCCACTTGTGTCTACACAAATATTTCTTGTTACTTGTGTACGCACAC
CAGCATGTGTTGATCACCAGCTTGTAGGCTATCAGCTTAGCAAGACCTCCTATATAAGGTTTTGCTAT
TCATTGAAAGCAGTAGTGACTGATTTGTATATA
>MinSyn_1520|Strength:0.00043945
GCGTGTCGTTTTAGTGAGGATCCTTACCGCTATGGGTAAGATTTTAGCAAGACCTCTACAAAACTGTG
AAGCATCTTCCTGCTGATCTTGCCGCTTTGTCAAAAGCTAAAAAAGATGATGCGCTCACTGCTAGGAT
TGCGATAAAGGAAAGGTATCAGCTTAGCAAGACCTCTACTATATAAGGTTTTGCTATTCATTGAAAGC
AGTAGTGACTGATTTGTATATA
>MinSyn_1464|Strength:0.000440105
GCGTGTCGTTTTAGTGAGGGGAAAAAGAAGAGGTAATTAGGTTGTCTGCACATCGAAAGGACAGTACG
GTAGGTCACGACGCACACCAGCATGTGTTGATCACCAGCTGATCTTGCCTGCTGGTGGAGCACGACAA
CCTCACATGTAGGCTATCAGCTTAAGCAAGTGGATGCTCACTGCTCTATATAAGGTTTTGCTATTCAT
TGAAAGCAGTAGTGACTGATTTGTATATA
>MinSyn_1787|Strength:0.000440456
GCGTGTCGTTTTAGTGAGGAGGTGGCTCCTACCGGTAGGTCACGACCACTATGCCCAATAAGATTGAT
GAAAAGTCAAAAACAAAAATCAATTATCTGCACCTCACATGTAGGCTATCAGCCTATATAAGGTTTTG
CTATTCATTGAAAGCAGTAGTGACTGATTTGTATATA
>MinSyn_1578|Strength:0.000440553
GCGTGTCGTTTTAGTGAGGTCAGAAGATCAAAGGGCTAAAGACCTCTACAAAACTGGTTTTTCAACAA
ATAACCATTATTGCGGCTTAGCAAGACCTCTACAAAACTGGTACTATTGCGATAAAGGAAAGGCTCAA
GCAAGTGGATTCACGACCACTATATAAGGTTTTGCTATTCATTGAAAGCAGTAGTGACTGATTTGTAT
ATA
>MinSyn_1820|Strength:0.000440629
GCGTGTCGTTTTAGTGAGGTTATGACCCCCGCCGATGACGCGGGAATCAGCTTAGCAAGACCTCTCAA
ATATTTCTTGTGCGGTAGGTCACGGGAAAAAGAAGAGGTTGTACATGAAGCATCTTCCTTAGGTTGTC
TGCACCTCACATGTAGAACCACGTCTACAAATCTTGCCTGTCTCTCTGCCGACAGTGGTCCCAAAGTC
TGCACCTCACATGTAGGCTATCCTTACCGCTATGGGTAAGATTTACACAGCCACTTGTGTCGATGCTG
ATCTTGCCTGCCTTGATGATAAAAATGTCAAAGATACTCTACAAAACTGGTACTTGTCTATATAAGGT
TTTGCTATTCATTGAAAGCAGTAGTGACTGATTTGTATATA
>MinSyn_1745|Strength:0.00044159
GCGTGTCGTTTTAGTGAGGTGAAGATAAGATAATAATGTTGAAGATAAGACCTGCCTTGATGATCAGA
AGATCAAAGGGCTAAGGAGGACCGATGCTGATCTTGCCCAAATATTTCTTGTGGACCGATGCTGATCT
TGCCTGCCTTGTTTTCAACAATAGGATTGCGATAAAGGAAAGGACCTCTACAACAGCCACTTGTGTCT
GATATCCTTACCGCTATGGGTAAGATTTACAAAACTGAATTTCGGGAAACCTCCTCGGTACAGGGCTC
ACTGCTAGGAGGACCGAAGCAAGTGGATTTAGGTTGTCTGCACCTCACATGCTATATAAGGTTTTGCT
ATTCATTGAAAGCAGTAGTGACTGATTTGTATATA
>MinSyn_1708|Strength:0.000442092
GCGTGTCGTTTTAGTGAGGGGAAAAAGAAGAGGTTGCACCCAAATATTTCTTGTCTCTAATTTCGGGA
AACCTCCTCGCTGCACCTCACATGTAGGCTATCATTGCGATAAAGGAAAGGAAACTGGTACTTGTGTA

CAGTCTCTCTGCCGACAGTGGTCCCAAACTGCATGGTGGAGCACGACAATAACCATTATTGCGGCTCA
CTGCTAGGAGGACCGATGCTCACTATCAGCCTGGTACTTGTGTACAGGGCTCCTATATAAGGTTTTGC
TATTCATTGAAAGCAGTAGTGACTGATTTGTATATA
>MinSyn_1540|Strength:0.000442612
GCGTGTCGTTTTAGTGAGGTCACTATCAGCCTGCTAGGAGGACCGATGCCAGTGGTCCCTCCACTCAC
GACCACTGGTGGAGCACGACAGCACCTCTGAAGCATCTTCCTCACATGTAGGCTATCAGCTTAAAAAA
TGTCAAAGATACTGCAAATATTTCTTGTAATCCTTACCGCTATGGGTAAGATTCGACCACTATGCCCA
AAATTTCGGGAAACCTCCTCGACAAAACTGGTACTTGTGTACAGGGCTAACCACGTCTACAATTAGGT
TGTCTGCACCTCACATGTCTATATAAGGTTTTGCTATTCATTGAAAGCAGTAGTGACTGATTTGTATA
TA
>MinSyn_13|Strength:0.00044263
GCGTGTCGTTTTAGTGAGGAAAAATGTCAAAGATATATCAGCTTAGCAAGGGAAAAAGAAGAGGTGGC
TCACTTGGTGGAGCACGACATACTTGTGTACAGGGCTCACTATATAAGGTTTTGCTATTCATTGAAAG
CAGTAGTGACTGATTTGTATATA
>MinSyn_1755|Strength:0.000442646
GCGTGTCGTTTTAGTGAGGTCTCTCTGCCGACAGTGGTCCCAAACTAGGAGGACCGATGCTGATCTTG
CCTGTCAGAAGATCAAAGGGCTACTTGATGATAAGCAAGTGGATGAGGACCGATGCTGATCAAAAATG
TCAAAGATATCTTGCCTGCTATATAAGGTTTTGCTATTCATTGAAAGCAGTAGTGACTGATTTGTATA
TA
>MinSyn_1955|Strength:0.000442857
GCGTGTCGTTTTAGTGAGGAAAAATGTCAAAGATACTTGATGATGTGGGAGCCACCATGCACCTCACA
TGTAGGCTACTATATAAGGTTTTGCTATTCATTGAAAGCAGTAGTGACTGATTTGTATATA
>MinSyn_1858|Strength:0.000443189
GCGTGTCGTTTTAGTGAGGAACCATTATTGCGTAGGCTTATGACCCCCGCCGATGACGCGGGATGTGT
ACAGGGCTCACTGCTAGGAGGACCAGGTGGCTCCTACCAAGACCTCTACTGGTGGAGCACGACAGCTG
ATCTTCTATATAAGGTTTTGCTATTCATTGAAAGCAGTAGTGACTGATTTGTATATA
>MinSyn_1997|Strength:0.000443271
GCGTGTCGTTTTAGTGAGGTCAGAAGATCAAAGGGCTATCTACAAAACTGGTACTTTGAAGATAAGAT
AATAATGTTGAAGATAAGACTATGCCCAATTAGGTTGTCTGAAGATTGATGAAAAGTCAAAAACAAAA
ATCAATTATCTGCCCAGCCACTTGTGTGGTAGGTCACGACCAAACCACGTCTACAATTAGCAAGACCT
CTACAAAACTGGTGTGGGAGCCACCAGTTGTCTGCACCTCAAATTTCGGGAAACCTCCTCGAAACTGG
TACTTGTGTAATCCTTACCGCTATGGGTAAGATTCCTCACATGTAGGCTATCCAAATATTTCTTGTCT
ATGCCCAATTAGGTTGTCTGCACCTCACCTATATAAGGTTTTGCTATTCATTGAAAGCAGTAGTGACT
GATTTGTATATA
>MinSyn_187|Strength:0.000444791
GCGTGTCGTTTTAGTGAGGGCACACCAGCATGTGTTGATCACCAGCTGTACTTGTGTACAGGGCTCAC
TGCTAGGAGCAAATATTTCTTGTCTATCAGCTTAGCAAGTCAGAAGATCAAAGGGCTACAAGACCTCT
ACCTATATAAGGTTTTGCTATTCATTGAAAGCAGTAGTGACTGATTTGTATATA
>MinSyn_1189|Strength:0.000444856
GCGTGTCGTTTTAGTGAGGAAAAATGTCAAAGATACTGGTACTTGTTTTTCAACAATGTAGGCTATCA
GCCAAATATTTCTTGTGGCTCACTGCTAGGAGGACCGATATTGCGATAAAGGAAAGGTGCTGGGAAAA
AGAAGAGGTGGTAGGTCACGACCACTATGCCCATCCTTACCGCTATGGGTAAGATTCTACAAAACTGG
TACTTGTGTACAGGGCTCAGGTGGCTCCTACTAACCACGTCTACAAGACCACTATGCCCAATTAGGTT
GTCGCACACCAGCATGTGTTGATCACCAGCTATCTTGCCTGCCTTGATGATCTATATAAGGTTTTGCT
ATTCATTGAAAGCAGTAGTGACTGATTTGTATATA
>MinSyn_1732|Strength:0.000445036
GCGTGTCGTTTTAGTGAGGCAAATATTTCTTGTTGCCCAATTAGGTTGTCTGCAAGATTGATGAAAAG
TCAAAAACAAAAATCAATTATACTGCTAGGAGGACCGAAACCACGTCTACAAGACCGATGCTGATCTT
CAGAAGATCAAAGGGCTATAGTTTTCAACAATCGCGGTAGGTCACGACCAGGAAAAAGAAGAGGTAGG
AGGACCGATGCTGATCCTATATAAGGTTTTGCTATTCATTGAAAGCAGTAGTGACTGATTTGTATATA
>MinSyn_1269|Strength:0.000445045
GCGTGTCGTTTTAGTGAGGCAGCCACTTGTGTTCACGACCACTTCACTATCAGCACTATATAAGGTTT
TGCTATTCATTGAAAGCAGTAGTGACTGATTTGTATATA
>MinSyn_1917|Strength:0.000445054

GCGTGTCGTTTTAGTGAGGGCACACCAGCATGTGTTGATCACCAGCTCGCGGTAGGTCACGACCACTA
TGCCCAATGCTTTGTCAAAAGCTAAAAAAGATGATGCAGACCTCTACAAAACTGGTACTTGTGTATGA
AGATAAGATAATAATGTTGAAGATAAGATCTACAAAACTGGTACTTGTTGGTGGAGCACGACATGCTC
AGAAGATCAAAGGGCTACGCGGTAGGTCACGACCACGTGGGAGCCACCAGGTACTTGTGTACAAGCAA
GTGGATTGTCTGCACCTCACATCGAAAGGACAGTACTATGCCCAATTAGGTTGTCTGCCTATATAAGG
TTTTGCTATTCATTGAAAGCAGTAGTGACTGATTTGTATATA
>MinSyn_1111|Strength:0.000445172
GCGTGTCGTTTTAGTGAGGGTGGGAGCCACCAACCACTATGCCCAATTAGGTTGTCTGCATCTCTCTG
CCGACAGTGGTCCCAAAGACCTCTACAAAACTGGTACTTAACCACGTCTACAATAACCATTATTGCGT
TGTCTGCACCTCACATGTAGGCTATCAGCGCACACCAGCATGTGTTGATCACCAGCTCACTGCTAGGA
CTATATAAGGTTTTGCTATTCATTGAAAGCAGTAGTGACTGATTTGTATATA
>MinSyn_1625|Strength:0.000445222
GCGTGTCGTTTTAGTGAGGTCCATCAACAAATAATCCAAGTAAGTACAAAACTGGTACTTGGGAAAAA
GAAGAGGTGCAGGTGGCTCCTACTTGTGTACAGGGCTCACTGCTAGGAGGCAGCCACTTGTGTCGACC
ACTATGCCCAATTAGGTTGTCACTATCAGCCTACAAAACTGGTGGGAGCCACCAACATGTAGGCTATC
AGCAAGATTGATGAAAAGTCAAAAACAAAAATCAATTATCGATGCTGATCTTGCTCTCTCTGCCGACA
GTGGTCCCAAATCACTGCTAGGAGGACAACCATTATTGCGCTCACATGTAGGCTATCAGCTTAGCAAG
CTATATAAGGTTTTGCTATTCATTGAAAGCAGTAGTGACTGATTTGTATATA
>MinSyn_1581|Strength:0.000445296
GCGTGTCGTTTTAGTGAGGCAAATATTTCTTGTGCTGATCTTGCCTCAGAAGATCAAAGGGCTATTGT
GTACAGGGCTCACTGCTAGGAGGATCACTATCAGCGGACCGGCTTTGTCAAAAGCTAAAAAAGATGAT
GCTCGCGGTAGGTCACGTGGTGGAGCACGACATCTGCACCTCACATGTAGGCTATCAGCCACTTGTGT
AAGACCTCTACAAAACTGGGAAAAAGAAGAGGTCTGCACCGTGGGAGCCACCAGGACCGATTGAAGCA
TCTTCCGTGTACAGGGCTCACTGCTAGGACTATATAAGGTTTTGCTATTCATTGAAAGCAGTAGTGAC
TGATTTGTATATA
>MinSyn_1517|Strength:0.000447223
GCGTGTCGTTTTAGTGAGGATTGCGATAAAGGAAAGGGCCTTGATGATAAACCACGTCTACAATGGTA
CTTGTGTACAGGGCTCACTGCTAGTGGTGGAGCACGACATTGCCTGCCTTGATGATATCCTTACCGCT
ATGGGTAAGATTCTCACTTTTCAACAAGGTGGGAGCCACCATGTACAGGGCTCACTGCTAGGAGGACT
ATATAAGGTTTTGCTATTCATTGAAAGCAGTAGTGACTGATTTGTATATA
>MinSyn_1188|Strength:0.000447732
GCGTGTCGTTTTAGTGAGGCAGTGGTCCCTCCACGATGCTGAGGTGGCTCCTACTATCAGCTTAGCAA
GAAAAAATGTCAAAGATAGGCTCACTGCTAGGAGGACCTGAAGCATCTTCCTGTGCTATATAAGGTTT
TGCTATTCATTGAAAGCAGTAGTGACTGATTTGTATATA
>MinSyn_1880|Strength:0.000447994
GCGTGTCGTTTTAGTGAGGCAGTGGTCCCTCCACAGGAGGAACCACGTCTACAACGATGCTGATCTTG
CCTGCCTTGATGATATTTTCAACAAAGGTTGTCTGCACCTCACATGTAGGCTATCATCGAAAGGACAG
TACCAATTAGGTTGAAGATTGATGAAAAGTCAAAAACAAAAATCAATTATTGCTGATCTTGCCTGCCT
TAACCATTATTGCGTACAGGGCTCACTGTGAAGCATCTTCCAGGCTATCAGCCTATATAAGGTTTTGC
TATTCATTGAAAGCAGTAGTGACTGATTTGTATATA
>MinSyn_1315|Strength:0.000448819
GCGTGTCGTTTTAGTGAGGTGAAGCATCTTCCCTGCTAGGAGGACCGATGCTGGTGGAGCACGACACC
GATCTATATAAGGTTTTGCTATTCATTGAAAGCAGTAGTGACTGATTTGTATATA
>MinSyn_1928|Strength:0.00044951
GCGTGTCGTTTTAGTGAGGTTATGACCCCCGCCGATGACGCGGGATACTTGTGTACAGGGCTCACAGT
GGTCCCTCCACCACTATGCCCAATTAGGTTGTCTGCAGGTGGCTCCTACGGTAGGTCACGACCACTAT
ATCCTTACCGCTATGGGTAAGATTCAAGACCAGCCACTTGTGTTGTAGGCTATCAGCTTAAATTTCGG
GAAACCTCCTCGACTGGTACTTGTGTACAGGGCTCACTGAAGATAAGATAATAATGTTGAAGATAAGA
GTACAGGGCTCACTGCTAGGAGGATTGCGATAAAGGAAAGGTGCCCAATTAGGTTGTCTGCTATATAA
GGTTTTGCTATTCATTGAAAGCAGTAGTGACTGATTTGTATATA
>MinSyn_1976|Strength:0.000449618
GCGTGTCGTTTTAGTGAGGTCAGAAGATCAAAGGGCTATGATGATATTTTCAACAACAAAACTGGTAC
TTGTGTACTATATAAGGTTTTGCTATTCATTGAAAGCAGTAGTGACTGATTTGTATATA
>MinSyn_1197|Strength:0.000449799

GCGTGTCGTTTTAGTGAGGATGACGTAAGCCATGACGTCTACTCTACAAAACTGGTACTTGTGTAGCA
AGTGGATACTATGCCCAATTAGGTTGTCTGCACCTTGAAGATAAGATAATAATGTTGAAGATAAGATA
TCTCTCTGCCGACAGTGGTCCCAAACCTCTACAAAACTTCAGAAGATCAAAGGGCTAGTTGTCTGCAT
TATGACCCCCGCCGATGACGCGGGAGTCTGGTGGGAGCCACCACAAAACTGGTACTTGTGTACAGGGC
TCACTTGAAGCATCTTCCTTAGCAAGACCTCTAAACCACGTCTACAAACAAAACTGGTACTTGTGTAC
AGGGCTCACCTATATAAGGTTTTGCTATTCATTGAAAGCAGTAGTGACTGATTTGTATATA
>MinSyn_186|Strength:0.000449905
GCGTGTCGTTTTAGTGAGGAACCACGTCTACAATATCAGCTTATGGTGGAGCACGACAGGACCGATGC
TGATCTTGCACACCAGCATGTGTTGATCACCAGCTCAGCTTAGCAAGACCTCTACAAAAATTTCGGGA
AACCTCCTCGACAGTGGGAGCCACCAGTTGTCTGCACCTCACATGTAGGCTATCAGCTATATAAGGTT
TTGCTATTCATTGAAAGCAGTAGTGACTGATTTGTATATA
>MinSyn_1309|Strength:0.000450571
GCGTGTCGTTTTAGTGAGGAAGATTGATGAAAAGTCAAAAACAAAAATCAATTATTAGGTTGTCTGCA
CCTCACATCCTTACCGCTATGGGTAAGATTTGTGTACAGGGCTCACTGGGAAAAAGAAGAGGTATGCC
CAATTAGGTTGTCTGCTTTGTCAAAAGCTAAAAAAGATGATGCTAGGTTGTCTGCACCTCACATGTAG
GCTCAGTGGTCCCTCCACGCAAGACCTCTACAAAACTGGTACTTTTTTCAACAAGTACAGGGCTCACC
AAATATTTCTTGTACTATGCCCAATGCACACCAGCATGTGTTGATCACCAGCTACTATGCCCAATTAG
GTTGTCCTATATAAGGTTTTGCTATTCATTGAAAGCAGTAGTGACTGATTTGTATATA
>MinSyn_1279|Strength:0.000450835
GCGTGTCGTTTTAGTGAGGAATTTCGGGAAACCTCCTCGGGGCTCACTGCTAGGAGGACCGATGTGGG
AGCCACCATCTACAAAACTGGTACTTGTGTAAGCAAGTGGATCAGGTGAAGATAAGATAATAATGTTG
AAGATAAGACGATGCTGATCTTGCCTGCCTTGATGATAATTGCGATAAAGGAAAGGTCTGCTATATAA
GGTTTTGCTATTCATTGAAAGCAGTAGTGACTGATTTGTATATA
>MinSyn_1212|Strength:0.000451427
GCGTGTCGTTTTAGTGAGGAACCACGTCTACAATAGGTCACGACCACTATGCCAAGATTGATGAAAAG
TCAAAAACAAAAATCAATTATGTACAGGGCTCACTGGGAAAAAGAAGAGGTCACATGTAGGCTAGCAC
ACCAGCATGTGTTGATCACCAGCTGCTGATCTTGCCTGCGTGGGAGCCACCAGCTATCAGCTTAGCAA
GACCTCTACAAAACTCACTATCAGCGTACTTGTGTACAACCATTATTGCGGCTATCAGCTCTATATAA
GGTTTTGCTATTCATTGAAAGCAGTAGTGACTGATTTGTATATA
>MinSyn_130|Strength:0.000452226
GCGTGTCGTTTTAGTGAGGTCAGAAGATCAAAGGGCTAGGTTGTCTGCACCTCACATGTAGGCTATGC
TTTGTCAAAAGCTAAAAAAGATGATGCAACTGGTAATTTCGGGAAACCTCCTCGACTATGCCCAATTA
GGTTGTTGAAGATAAGATAATAATGTTGAAGATAAGAAGCTTAGCAAGACCTCTACAAAAATTGCGAT
AAAGGAAAGGCGCGGTAGGTCACGACCACTATCTATATAAGGTTTTGCTATTCATTGAAAGCAGTAGT
GACTGATTTGTATATA
>MinSyn_1718|Strength:0.000452239
GCGTGTCGTTTTAGTGAGGTGAAGCATCTTCCCCGATGCTGATCTTGCCTGCCTTGATGATATGAAGA
TAAGATAATAATGTTGAAGATAAGAGCTCACCTATATAAGGTTTTGCTATTCATTGAAAGCAGTAGTG
ACTGATTTGTATATA
>MinSyn_1507|Strength:0.000452783
GCGTGTCGTTTTAGTGAGGTGAAGATAAGATAATAATGTTGAAGATAAGATTCCATCAACAAATAATC
CAAGTAAGCTACAAAACTGGTACTCAAATATTTCTTGTAGCTTAGCAAGACCTCTACAAACTATATAA
GGTTTTGCTATTCATTGAAAGCAGTAGTGACTGATTTGTATATA
>MinSyn_1741|Strength:0.000452977
GCGTGTCGTTTTAGTGAGGAGGTGGCTCCTACCACGACCACTATGCCAAAAATGTCAAAGATAGCTAG
GAGATTGCGATAAAGGAAAGGCACTGCTAGGAGGACCGATGCTTTTCAACAAAAAACTGGTACTTGTG
TACAGGATCCTTACCGCTATGGGTAAGATTTGGTACTTGGGAAAAAGAAGAGGTATGTAGGCTATCAG
CTAACCACGTCTACAACAGGGCTCTCTCTCTGCCGACAGTGGTCCCAAACACCTCACATGTAGGCTAT
CAGCTTAGCCTATATAAGGTTTTGCTATTCATTGAAAGCAGTAGTGACTGATTTGTATATA
>MinSyn_1237|Strength:0.000453104
GCGTGTCGTTTTAGTGAGGTCCATCAACAAATAATCCAAGTAAGGACCACTATGCCCAATTAGGTTGT
TGAAGATAAGATAATAATGTTGAAGATAAGATACTTGTGTACAGGGCTCACTGCTTTATGACCCCCGC
CGATGACGCGGGAACCTCTACAAAACTGGTACCTATATAAGGTTTTGCTATTCATTGAAAGCAGTAGT
GACTGATTTGTATATA

>MinSyn_121|Strength:0.000453313
GCGTGTCGTTTTAGTGAGGTTATGACCCCCGCCGATGACGCGGGACCTGCCTTGAGGTGGCTCCTACC
CTGCCTTGATGATGTGGGAGCCACCAGCTAGGAGGACCGATGCTATATAAGGTTTTGCTATTCATTGA
AAGCAGTAGTGACTGATTTGTATATA
>MinSyn_1740|Strength:0.000453752
GCGTGTCGTTTTAGTGAGGCAGCCACTTGTGTTCTGCACCTCACATGTAGGTCTCTCTGCCGACAGTG
GTCCCAAATTTATGACCCCCGCCGATGACGCGGGATTGTGTACAGGGCTCACTGCTAGGAGATCCTTA
CCGCTATGGGTAAGATTACCTCACATGTAGGCTATCAGCTCTATATAAGGTTTTGCTATTCATTGAAA
GCAGTAGTGACTGATTTGTATATA
>MinSyn_1855|Strength:0.000453838
GCGTGTCGTTTTAGTGAGGATCGAAAGGACAGTAACTATGCCCAATTAGGTTGTCTTGAAGATAAGAT
AATAATGTTGAAGATAAGACAATTAGGTTGTCTGCACCTTGGTGGAGCACGACATGCTAGGAGGACCG
ATGCTGATTGAAGCATCTTCCAAGACCTCTACAAAACTGGTACTTGTGTCACTATCAGCTGAAAAATG
TCAAAGATAATCAGCTTAGCAAGACCTCTACAAAACCAGCCACTTGTGTTGATATCCATCAACAAATA
ATCCAAGTAAGCGATGCTGATCTTGCCTTTATGACCCCCGCCGATGACGCGGGACTAGGAGGACCGAT
GCTGATCTTGCCTATATAAGGTTTTGCTATTCATTGAAAGCAGTAGTGACTGATTTGTATATA
>MinSyn_1574|Strength:0.000454059
GCGTGTCGTTTTAGTGAGGGCACACCAGCATGTGTTGATCACCAGCTACCACTATCAGAAGATCAAAG
GGCTAGTAGGTCACGACCACTATGCCCAATTAGGAAAAAGAAGAGGTCACATGTAGGCTATCAGCTTA
GCATTGCGATAAAGGAAAGGACCACTATGCCCAGCCACTTGTGTCACTATGCCCAATTAGGTTCTCTC
TGCCGACAGTGGTCCCAAAACTGCTAGGAGGACCGATGCTGATATCGAAAGGACAGTAACTGGTACTT
GTCACTATCAGCGCCTTGAAAAATGTCAAAGATACAAGACCTCTACAAAACTGGTCTATATAAGGTTT
TGCTATTCATTGAAAGCAGTAGTGACTGATTTGTATATA
>MinSyn_1327|Strength:0.000454542
GCGTGTCGTTTTAGTGAGGGCACACCAGCATGTGTTGATCACCAGCTTACTTGTGTACAGGGCTCAAC
CATTATTGCGACCTCACATAAGATTGATGAAAAGTCAAAAACAAAAATCAATTATGCAAGACCTCTAC
AAATTATGACCCCCGCCGATGACGCGGGATACAAGGAAAAAGAAGAGGTCGATGCTGATCTTGCCTGC
CTTGATAGCAAGTGGATTACAGGGCTTCACTATCAGCTAGGAGGACCGATGCTGATCTTGCCTGCCTC
TATATAAGGTTTTGCTATTCATTGAAAGCAGTAGTGACTGATTTGTATATA
>MinSyn_1618|Strength:0.000454801
GCGTGTCGTTTTAGTGAGGGCACACCAGCATGTGTTGATCACCAGCTAAAACTGGTACTTGTGTACAT
GGTGGAGCACGACATCACATTCTCTCTGCCGACAGTGGTCCCAAAAGCTTAGCAAGACCTCTACAAGC
AAGTGGATCCGATGCGGAAAAAGAAGAGGTGCAAGACCTCTTTTTCAACAATCGCGGTAGGTCACGAC
CACTCTATATAAGGTTTTGCTATTCATTGAAAGCAGTAGTGACTGATTTGTATATA
>MinSyn_1720|Strength:0.000455346
GCGTGTCGTTTTAGTGAGGTCACTATCAGCTATCAGCTTAGCAAGACATCGAAAGGACAGTAGCACCT
CACATGTAGGCTCAGTGGTCCCTCCACCACGACCACTATGCCCAATTAGGTTGAGGTGGCTCCTACTC
GCGGTATCCTTACCGCTATGGGTAAGATTCACCTCACATGTCTATATAAGGTTTTGCTATTCATTGAA
AGCAGTAGTGACTGATTTGTATATA
>MinSyn_1886|Strength:0.000455346
GCGTGTCGTTTTAGTGAGGTGAAGATAAGATAATAATGTTGAAGATAAGACTGGTGGAGCACGACAGG
AGGAAAAATGTCAAAGATACAATTAGGTGTGGGAGCCACCAAAGACCTCTACAAAACTGGTACTTGTA
GCAAGTGGATCACGACCACTATGCCCAATTAGGTTGTCTTCAGAAGATCAAAGGGCTACCTCACATGT
AGGCTATCATGAAGCATCTTCCGGCTATCAGCTTAGCAAGACCTCTACAAGCACACCAGCATGTGTTG
ATCACCAGCTTCTTGCCTGCCTTGATTCTCTCTGCCGACAGTGGTCCCAAATATGCCCAATTAGGCTA
TATAAGGTTTTGCTATTCATTGAAAGCAGTAGTGACTGATTTGTATATA
>MinSyn_1317|Strength:0.000456148
GCGTGTCGTTTTAGTGAGGTCACTATCAGCGTAGGCTATCAGCTTGGAAAAAGAAGAGGTACAAAACT
TCTCTCTGCCGACAGTGGTCCCAAACAAGACCTCTACAAAACTGGTGGTGGAGCACGACAGATCTTGC
CTGCCTTGATGATAAGATTGATGAAAAGTCAAAAACAAAAATCAATTATTACTTGTGTACATTATGAC
CCCCGCCGATGACGCGGGAGGCTATCAGCTTAGCAAGACCTCTACAAAAAATTTCGGGAAACCTCCTC
GATTAGGTTGTCTGCACCTCACATGCTATATAAGGTTTTGCTATTCATTGAAAGCAGTAGTGACTGAT
TTGTATATA
>MinSyn_1161|Strength:0.000456466

GCGTGTCGTTTTAGTGAGGGCTTTGTCAAAAGCTAAAAAAGATGATGCTGCACCTCACATGTAGGCTA
TCAGCTTAGCGTGGGAGCCACCAACATGTAGGCTATCAGCAACCATTATTGCGCTCACTGCTAGGAGG
CAAATATTTCTTGTGATGCTGATCTTGCCTGCCTTGATTGCGATAAAGGAAAGGTACAGGGAAAAAGA
AGAGGTGTCACGACCACTATGCCCAATTAGGTGAAGCATCTTCCACATGAAGATAAGATAATAATGTT
GAAGATAAGAAGGAGGACCGATGCTGATCTTATCGAAAGGACAGTATCTACAAAACTGGTACTTGCTA
TATAAGGTTTTGCTATTCATTGAAAGCAGTAGTGACTGATTTGTATATA
>MinSyn_1710|Strength:0.000457329
GCGTGTCGTTTTAGTGAGGAAAAATGTCAAAGATACTCATGGTGGAGCACGACACAGCTTAGCAAGAA
GGTGGCTCCTACCCTCTACAAAACTGGTAAGCAAGTGGATACCTATATAAGGTTTTGCTATTCATTGA
AAGCAGTAGTGACTGATTTGTATATA
>MinSyn_176|Strength:0.000457337
GCGTGTCGTTTTAGTGAGGTGGTGGAGCACGACAATTAGGTTGTCTGCACCTTGAAGATAAGATAATA
ATGTTGAAGATAAGACCAATTAGGTTGTCTGCACTATATAAGGTTTTGCTATTCATTGAAAGCAGTAG
TGACTGATTTGTATATA
>MinSyn_1487|Strength:0.000459371
GCGTGTCGTTTTAGTGAGGAGGTGGCTCCTACACCACTATGCCCGGAAAAAGAAGAGGTTAGCAAGAC
CTCTATGAAGATAAGATAATAATGTTGAAGATAAGATAGGAGGACCGATGCTGATCTTGTGGTGGAGC
ACGACAACCTCTACAAAACTGGTACTTGTGTGGGAGCCACCAATGCCCAATTAGGTTGAAAAATGTCA
AAGATAACTGGTACTTGTGTCAAATATTTCTTGTATGTCTATATAAGGTTTTGCTATTCATTGAAAGC
AGTAGTGACTGATTTGTATATA
>MinSyn_1664|Strength:0.0004594
GCGTGTCGTTTTAGTGAGGGTGGGAGCCACCAGTCACGACCACTATGCCCAATTAGGTTGTCAACCAC
GTCTACAAATTAGGTTGTCTGCACCTCACATGCAGCCACTTGTGTGCTATCAGCTTAGCAAGACCTCT
TCCATCAACAAATAATCCAAGTAAGAGGACCGATGCTGATCTTGCCTGCCTATCCTTACCGCTATGGG
TAAGATTCTCACTGCTAGGAGGACCGATGCATTGCGATAAAGGAAAGGCTAGGAGGACCGATGCTGAT
CTTGCCTGCCGCACACCAGCATGTGTTGATCACCAGCTTAGGAGGACCGAGGTGGCTCCTACTATCGA
AAGGACAGTATCTGCACCTCACATGTAGCTATATAAGGTTTTGCTATTCATTGAAAGCAGTAGTGACT
GATTTGTATATA
>MinSyn_1113|Strength:0.000459401
GCGTGTCGTTTTAGTGAGGGGAAAAAGAAGAGGTCTGCACCTCACATGTAGGCTATCAGCTTAAAGAT
TGATGAAAAGTCAAAAACAAAAATCAATTATTGTAGGCTATCAGCTTAGCAAGACCATTGCGATAAAG
GAAAGGATGCCCAATTAGGTTGTCTGCACCTCACAAAAATGTCAAAGATATCTACAAAACTGGTACTT
GTTCCATCAACAAATAATCCAAGTAAGAACTGGTAGCTTTGTCAAAAGCTAAAAAAGATGATGCACCT
CACATGTAGGCTATCAGCTTAGCACTATATAAGGTTTTGCTATTCATTGAAAGCAGTAGTGACTGATT
TGTATATA
>MinSyn_123|Strength:0.000460108
GCGTGTCGTTTTAGTGAGGAACCATTATTGCGCAGTCAGAAGATCAAAGGGCTATCTACAGTGGGAGC
CACCATGTACAGGGCTCACTGCTAGGAGGACCAAAAATGTCAAAGATAGATCTTGCCAGGTGGCTCCT
ACACAGGGCTCCTATATAAGGTTTTGCTATTCATTGAAAGCAGTAGTGACTGATTTGTATATA
>MinSyn_1849|Strength:0.000460631
GCGTGTCGTTTTAGTGAGGAAGATTGATGAAAAGTCAAAAACAAAAATCAATTATCTGATCTTGCCTT
CACTATCAGCCTGAAGCATCTTCCAGGGCTCACTGCTTCTCTCTGCCGACAGTGGTCCCAAAAAGAAG
CAAGTGGATAAAACTGGTACTTGTGTACAGGGCTGGAAAAAGAAGAGGTACAGGGCTCACTGCTAGGA
CTATATAAGGTTTTGCTATTCATTGAAAGCAGTAGTGACTGATTTGTATATA
>MinSyn_1593|Strength:0.000460644
GCGTGTCGTTTTAGTGAGGAACCACGTCTACAAATTAGGTTGTCTGCTGAAGATAAGATAATAATGTT
GAAGATAAGAGGAGGACCGATGCTGATCTTGCCTGGTGGAGCACGACACAGGGCTCACTGAATTTCGG
GAAACCTCCTCGCTAGGAGCAGCCACTTGTGTTCTTGCCTGCCTTGATGTTATGACCCCCGCCGATGA
CGCGGGAGACCACTATGCCCAACAGTGGTCCCTCCACTAGGAGGACCGATGCTGATCTTGCTATATAA
GGTTTTGCTATTCATTGAAAGCAGTAGTGACTGATTTGTATATA
>MinSyn_1857|Strength:0.000461199
GCGTGTCGTTTTAGTGAGGGCACACCAGCATGTGTTGATCACCAGCTTGCTGATCTTGCCTGCCTTAA
CCACGTCTACAATATGCCCAATTAGGTTGTCTGCTGAAGCATCTTCCAGACCTCTCCATCAACAAATA
ATCCAAGTAAGCGATGCTGATCTTGCCTGCATTGCGATAAAGGAAAGGAGACCTCTACAAGATTGATG

AAAAGTCAAAAACAAAAATCAATTATAATTAGGTTGTAGCAAGTGGATTGATCTTGCCTGCCTTGATG
ATCTCTCTGCCGACAGTGGTCCCAAAACAGGGCTCACTGCTAGGAGGACCGACAGCCACTTGTGTAAC
TGGTACTTGTGTACAGGGCCTATATAAGGTTTTGCTATTCATTGAAAGCAGTAGTGACTGATTTGTAT
ATA
>MinSyn_1401|Strength:0.000461706
GCGTGTCGTTTTAGTGAGGTTTTCAACAATGTCTGCACCTCACATGTAGGGCTTTGTCAAAAGCTAAA
AAAGATGATGCAGGACCGATGCTGATCTTGCCTGCCTCAGTGGTCCCTCCACAGCTTAGCAAGACCTC
TACAAAACTGGTATTGCGATAAAGGAAAGGCTGGTACTTGTGTACAGGTCCATCAACAAATAATCCAA
GTAAGCAGGGCTCACTGCATCCTTACCGCTATGGGTAAGATTGCTATCAGCTTAGTGGGAGCCACCAG
TACAGGGCTCACTGCTAGGAGCTATATAAGGTTTTGCTATTCATTGAAAGCAGTAGTGACTGATTTGT
ATATA
>MinSyn_122|Strength:0.000461739
GCGTGTCGTTTTAGTGAGGGTGGGAGCCACCATCTACAAAACTGGTACTTGTGTACATTGCGATAAAG
GAAAGGAATTAGGTTGTCTGCACCTCACATGTAGGGGAAAAAGAAGAGGTTGCCAGCAAGTGGATTCG
CGGTAGGTTCACTATCAGCGACCGATGCTGATCTTGCCTGCCTTGATGCAGCCACTTGTGTAGGCTAT
CAATTTCGGGAAACCTCCTCGCCTGCCTTGATGATGAAGCATCTTCCACCTATATAAGGTTTTGCTAT
TCATTGAAAGCAGTAGTGACTGATTTGTATATA
>MinSyn_1527|Strength:0.000462033
GCGTGTCGTTTTAGTGAGGAAGATTGATGAAAAGTCAAAAACAAAAATCAATTATGTCTGCACCTCAC
ATGTAGGCTATCATCAGAAGATCAAAGGGCTACAGCAAGTGGATGCCTGCCTTGATGATAATCCTTAC
CGCTATGGGTAAGATTCTGCACCTCACATGTAGGCTATCACTATATAAGGTTTTGCTATTCATTGAAA
GCAGTAGTGACTGATTTGTATATA
>MinSyn_1589|Strength:0.000462613
GCGTGTCGTTTTAGTGAGGGGAAAAAGAAGAGGTCACTATGCCCAATTAGGTTGTCTGCACCCAGTGG
TCCCTCCACCTTAGCAAGACCTCTACAAAACTGGTACTTCTCTCTGCCGACAGTGGTCCCAAACCTCA
CATGTAGGCTATTGCGATAAAGGAAAGGTGGTACTTGCTATATAAGGTTTTGCTATTCATTGAAAGCA
GTAGTGACTGATTTGTATATA
>MinSyn_1543|Strength:0.000462821
GCGTGTCGTTTTAGTGAGGTCCATCAACAAATAATCCAAGTAAGCTGCACCTCACATGTAGGCTATCT
CTCTCTGCCGACAGTGGTCCCAAAGCACCTCACTATATAAGGTTTTGCTATTCATTGAAAGCAGTAGT
GACTGATTTGTATATA
>MinSyn_1374|Strength:0.00046294
GCGTGTCGTTTTAGTGAGGGTGGGAGCCACCAGCAAGACCTCTACAAAACTGGTGGTGGAGCACGACA
TACAAAACTGGTACTTGGCTTTGTCAAAAGCTAAAAAAGATGATGCAGTCTCTCTGCCGACAGTGGTC
CCAAACTGCTAGGAGGACCAACCATTATTGCGAGCTTAGCAAGACCTCTACAAAACTGGCAGTGGTCC
CTCCACGGTTGTCTGCTATATAAGGTTTTGCTATTCATTGAAAGCAGTAGTGACTGATTTGTATATA
>MinSyn_1472|Strength:0.000463386
GCGTGTCGTTTTAGTGAGGATCGAAAGGACAGTATACAGGGCTTGAAGCATCTTCCCTTGTGTACAGG
GCTCACTGTGGGAGCCACCACTCACTGCTAGGAGGACCGATGCTCAGCCACTTGTGTCCTCACTCACT
ATCAGCTGTAGGGCACACCAGCATGTGTTGATCACCAGCTTGCTAGGAGGAAATTTCGGGAAACCTCC
TCGAGACCTCTACAAAACTGGTACTTGCTATATAAGGTTTTGCTATTCATTGAAAGCAGTAGTGACTG
ATTTGTATATA
>MinSyn_1355|Strength:0.000463415
GCGTGTCGTTTTAGTGAGGTGAAGCATCTTCCGCACCTCACATCAGTGGTCCCTCCACAGGTTGTCTC
TATATAAGGTTTTGCTATTCATTGAAAGCAGTAGTGACTGATTTGTATATA
>MinSyn_184|Strength:0.000463477
GCGTGTCGTTTTAGTGAGGGCTTTGTCAAAAGCTAAAAAAGATGATGCAGCAAGACCTCTACAAAACT
GGTACTTCACTATCAGCACTTGTGTACAGGGCCAAATATTTCTTGTAGCAAGACCTCTACAAAACTGG
TACTTGCACACCAGCATGTGTTGATCACCAGCTAGGAGGACCGATGCTGATCTTTTCAACAATCTGCA
CCTCACATAATTTCGGGAAACCTCCTCGGCGGCTATATAAGGTTTTGCTATTCATTGAAAGCAGTAGT
GACTGATTTGTATATA
>MinSyn_1505|Strength:0.000463507
GCGTGTCGTTTTAGTGAGGAAAAATGTCAAAGATATCTGCACCTCACATGTAGGTGGGAGCCACCATA
GGAGGACCGATGCTGATCTTGCCTCAGTGGTCCCTCCACTTAGCAAATCGAAAGGACAGTACACCTCA

CATGTAGGAACCACGTCTACAAACCACTATGCCCAGCAAGTGGATCTTAGGCTTTGTCAAAAGCTAAA
AAAGATGATGCTCGCGGTAGGTCACGACCACTATGGTGGAGCACGACAATTAGGTTGTCTGCACCTTC
CATCAACAAATAATCCAAGTAAGAGGGCTCACTGCTAGGAGGACCCTATATAAGGTTTTGCTATTCAT
TGAAAGCAGTAGTGACTGATTTGTATATA
>MinSyn_1881|Strength:0.000463701
GCGTGTCGTTTTAGTGAGGCAGCCACTTGTGTGCTCACTGCTAGGAGGTGGCTCCTACCAAAACTGGT
ACTTGTGTACAGGGCTCTTATGACCCCCGCCGATGACGCGGGACGATGCTGATCTTGCCTGCCTTGAT
GAATTGCGATAAAGGAAAGGCAGCTTAGCAAGACCTCTACAAAACTGGTAGCTTTGTCAAAAGCTAAA
AAAGATGATGCACCACTATGCCCAATATCCTTACCGCTATGGGTAAGATTTGTGTACAGGGCAGTGGT
CCCTCCACTGTACAGGGCTCACTGCTAGTCCATCAACAAATAATCCAAGTAAGGCCCAATTAGGTTGT
CTGCACTATATAAGGTTTTGCTATTCATTGAAAGCAGTAGTGACTGATTTGTATATA
>MinSyn_1345|Strength:0.000464107
GCGTGTCGTTTTAGTGAGGTGAAGATAAGATAATAATGTTGAAGATAAGACGATGTCTCTCTGCCGAC
AGTGGTCCCAAATGCACCTCACATGTAGGCTATTCACTATCAGCCAATTAGGTTGTCTGCACCTCACA
TGTAGTCAGAAGATCAAAGGGCTACTATCAGCTTAGCAAGACCTCTACAATTATGACCCCCGCCGATG
ACGCGGGATATCTATATAAGGTTTTGCTATTCATTGAAAGCAGTAGTGACTGATTTGTATATA
>MinSyn_18|Strength:0.000464916
GCGTGTCGTTTTAGTGAGGATCCTTACCGCTATGGGTAAGATTACATAAGATTGATGAAAAGTCAAAA
ACAAAAATCAATTATCAGCTTAGCAACCACGTCTACAATACTTGTGTACAGGGCTCACTGCTAGCAGT
GGTCCCTCCACACTTGTGTACAGGCTATATAAGGTTTTGCTATTCATTGAAAGCAGTAGTGACTGATT
TGTATATA
>MinSyn_1985|Strength:0.000464946
GCGTGTCGTTTTAGTGAGGTCCATCAACAAATAATCCAAGTAAGGCCCAATTAGGTTGTCTGCACCTC
ACATGAATTTCGGGAAACCTCCTCGCGACCACTATGCCCAATTAGGTTGTCTGCAAACCACGTCTACA
ACCTCACATGTAGGCTATCAGCTTAGCAGGTGGCTCCTACCTGCCTTGATGATACAAATATTTCTTGT
CGACCACTATGCCCAAATCGAAAGGACAGTATAGGTCACGAAAGATTGATGAAAAGTCAAAAACAAAA
ATCAATTATCTTGTGTACAGGGCTCCTATATAAGGTTTTGCTATTCATTGAAAGCAGTAGTGACTGAT
TTGTATATA
>MinSyn_1743|Strength:0.000465044
GCGTGTCGTTTTAGTGAGGTGAAGCATCTTCCGTAGGTCACGACCACTATGCCCAATTAGTGAAGATA
AGATAATAATGTTGAAGATAAGAAACTGGTACTTGTGTACAGGGCTCACTGTCCATCAACAAATAATC
CAAGTAAGCTTGTGTACAGGGCTTCTCTCTGCCGACAGTGGTCCCAAAGATGCTGATCTTGCCCTATA
TAAGGTTTTGCTATTCATTGAAAGCAGTAGTGACTGATTTGTATATA
>MinSyn_144|Strength:0.000465274
GCGTGTCGTTTTAGTGAGGTGGTGGAGCACGACATCTACAAAACTGGTACTTGTGAACCATTATTGCG
TGCCCAATTAGGTTGTCTGCACCTCAGTGGTCCCTCCACCTGATCTTGCCTGCCTTTATGACCCCCGC
CGATGACGCGGGACGCGGTAGGTCACGACCACTATGCCGCACACCAGCATGTGTTGATCACCAGCTTG
CCTTGAAAAATGTCAAAGATAGCTTTTTCAACAATATGCCCAATTAGGTTGTCTGCACCTCTATATAA
GGTTTTGCTATTCATTGAAAGCAGTAGTGACTGATTTGTATATA
>MinSyn_1620|Strength:0.000465346
GCGTGTCGTTTTAGTGAGGAAAAATGTCAAAGATATACTTGTGTACAGGGGGAAAAAGAAGAGGTAAA
TTTCGGGAAACCTCCTCGTGCACCTCACATGTAGGCTATTATGACCCCCGCCGATGACGCGGGAAGGT
TGTCTGCACCTCACATGTAGGCTATCTCCATCAACAAATAATCCAAGTAAGACTATGCCCAATTAGGT
TGTCTGCACCTCAAGATTGATGAAAAGTCAAAAACAAAAATCAATTATGTTGTCTGCACCTCACATGT
AGGCTATATAAGGTTTTGCTATTCATTGAAAGCAGTAGTGACTGATTTGTATATA
>MinSyn_1984|Strength:0.000465529
GCGTGTCGTTTTAGTGAGGATTGCGATAAAGGAAAGGTCTTGCCTGCCTTGGCTTTGTCAAAAGCTAA
AAAAGATGATGCCATGTAGGCTATCAGCTTAGCAAGACTGAAGCATCTTCCAGCTTAGCAAGACCTCT
ACAAAACTAACCACGTCTACAACTGCTCTCTCTGCCGACAGTGGTCCCAAAGACCACTATGCCCAATT
AGGTCTATATAAGGTTTTGCTATTCATTGAAAGCAGTAGTGACTGATTTGTATATA
>MinSyn_1842|Strength:0.000465637
GCGTGTCGTTTTAGTGAGGTCCATCAACAAATAATCCAAGTAAGCTTGTGTACAGGGCTCACTGCTAG
GAGGAATTGCGATAAAGGAAAGGTACTTGTGTACAGGGTCACTATCAGCATGAATCCTTACCGCTATG
GGTAAGATTCACTATGCCCAATTAGGTTAGGTGGCTCCTACGGGCTCACTGCTAGTGGTGGAGCACGA

CACTGCCTTGATGAGCAAGTGGATGTACAGGGCTCCTATATAAGGTTTTGCTATTCATTGAAAGCAGT
AGTGACTGATTTGTATATA
>MinSyn_1306|Strength:0.000465666
GCGTGTCGTTTTAGTGAGGATCGAAAGGACAGTACACTGCTAGGAGGACCGATGCTGATGCACACCAG
CATGTGTTGATCACCAGCTTACTATATAAGGTTTTGCTATTCATTGAAAGCAGTAGTGACTGATTTGT
ATATA
>MinSyn_1559|Strength:0.000465724
GCGTGTCGTTTTAGTGAGGGGAAAAAGAAGAGGTAGGTCACGACCACTTCCATCAACAAATAATCCAA
GTAAGGGGCTCATCGAAAGGACAGTATGTAGGCTATCAGCTTAGCAAGACAACCACGTCTACAATTGT
CTGCACCTCACATGTAGGCTATCAAAAATGTCAAAGATAGGTACTTAATTTCGGGAAACCTCCTCGAC
AGCCACTTGTGTCTTAGCAAGACCTCTACAAAACTGGCTATATAAGGTTTTGCTATTCATTGAAAGCA
GTAGTGACTGATTTGTATATA
>MinSyn_141|Strength:0.000466777
GCGTGTCGTTTTAGTGAGGGCTTTGTCAAAAGCTAAAAAAGATGATGCCTAGGAGGACCGATGCTGAT
CTGAAGCATCTTCCAGGACCGATGCTGATCTTGCCTGCATCCTTACCGCTATGGGTAAGATTCAGGGC
TCACTGCTAGGAGGACCGATAAAAATGTCAAAGATATCACTGCTAGGAGGACCGATGCTGATCAGGTG
GCTCCTACGTAGGAATTTCGGGAAACCTCCTCGACCTATATAAGGTTTTGCTATTCATTGAAAGCAGT
AGTGACTGATTTGTATATA
>MinSyn_1703|Strength:0.000466905
GCGTGTCGTTTTAGTGAGGTCAGAAGATCAAAGGGCTAGATCCTTACCGCTATGGGTAAGATTTAGGC
TATCAGCTTAGCAAGAAATTTCGGGAAACCTCCTCGAACTGGTACTTGTGTATGAAGATAAGATAATA
ATGTTGAAGATAAGAAGGGCTCACTGCTAGGAGGACCGATGCTGCTATATAAGGTTTTGCTATTCATT
GAAAGCAGTAGTGACTGATTTGTATATA
>MinSyn_1939|Strength:0.000466949
GCGTGTCGTTTTAGTGAGGAAAAATGTCAAAGATACCCAATTAGGTTGTTATGACCCCCGCCGATGAC
GCGGGAACGAAATTTCGGGAAACCTCCTCGTCACATGTAGGCTATCAGCTTCCATCAACAAATAATCC
AAGTAAGAAACTGGTACTTGTGTAATCCTTACCGCTATGGGTAAGATTACTGGTACTTGTGTACAGGG
CTCACTGCTATATAAGGTTTTGCTATTCATTGAAAGCAGTAGTGACTGATTTGTATATA
>MinSyn_1561|Strength:0.000467289
GCGTGTCGTTTTAGTGAGGTGAAGCATCTTCCTATGCCCAATTAGGTTGTCTGCACCTCACTCCATCA
ACAAATAATCCAAGTAAGGACCTCTACAAAACTTCACTATCAGCTGTAGGCTATCAGCTATCGAAAGG
ACAGTAGATCTTGTCTCTCTGCCGACAGTGGTCCCAAACGACCACTATGCCCAATTAGGTTGTTGAAG
ATAAGATAATAATGTTGAAGATAAGAGCTAGGAGGACCGATGCTGATCCTATATAAGGTTTTGCTATT
CATTGAAAGCAGTAGTGACTGATTTGTATATA
>MinSyn_179|Strength:0.000467529
GCGTGTCGTTTTAGTGAGGATCCTTACCGCTATGGGTAAGATTCACCTCACATGTAGGCTATCAGCTT
AGCAAAGATTGATGAAAAGTCAAAAACAAAAATCAATTATTCTACAAAACTGGTACTTGTGTCTATAT
AAGGTTTTGCTATTCATTGAAAGCAGTAGTGACTGATTTGTATATA
>MinSyn_1337|Strength:0.000467587
GCGTGTCGTTTTAGTGAGGTCTCTCTGCCGACAGTGGTCCCAAAGGTTGTCTGCACCTCACATGTAGG
CTATAAGATTGATGAAAAGTCAAAAACAAAAATCAATTATCTAAAAATGTCAAAGATATCTTGCCTGC
ACACCAGCATGTGTTGATCACCAGCTCTAGGAGGACCGATGCTGATCTTGCCTAGCAAGTGGATTGCC
CAATTAGGTTGTCTGCTATATAAGGTTTTGCTATTCATTGAAAGCAGTAGTGACTGATTTGTATATA
>MinSyn_1167|Strength:0.000468044
GCGTGTCGTTTTAGTGAGGGTGGGAGCCACCACTTTTATGACCCCCGCCGATGACGCGGGAACTTGTG
TACAGGGCTCACTGCTAGAATTTCGGGAAACCTCCTCGGCTATCAGCTTAGCAAGACCCAAATATTTC
TTGTTCAGCTTAGCAAGACCTCTACAAAACTGGTATCCTTACCGCTATGGGTAAGATTCTGGTACTGG
AAAAAGAAGAGGTGCTCACTGCTAGGCTTTGTCAAAAGCTAAAAAAGATGATGCGCTTAGCAAGACCT
CTATCACTATCAGCGTACTAGGTGGCTCCTACACCACTATGCCCAATTAGGTTGTCTGCTATATAAGG
TTTTGCTATTCATTGAAAGCAGTAGTGACTGATTTGTATATA
>MinSyn_1669|Strength:0.000469287
GCGTGTCGTTTTAGTGAGGAGCAAGTGGATCTATCAGCTTAGCAAGACGCACACCAGCATGTGTTGAT
CACCAGCTAGGTTGTCTGCACTTATGACCCCCGCCGATGACGCGGGATAGGAGAAAAATGTCAAAGAT
AATGGTGGAGCACGACAACATGTAGGCTAAGGTGGCTCCTACCACATGTAGGCTATCAGCTTAGCAAA

ATTTCGGGAAACCTCCTCGAATTAGGTTGTCTGCACCTCACACTATATAAGGTTTTGCTATTCATTGA
AAGCAGTAGTGACTGATTTGTATATA
>MinSyn_1709|Strength:0.00046944
GCGTGTCGTTTTAGTGAGGGGAAAAAGAAGAGGTTCTTGCCTGCCTTGATGTCTCTCTGCCGACAGTG
GTCCCAAACTATCAGCTTAGCAAGACCTCTACAAAACCATTATTGCGTCTGCACCTCACATGTAGGCT
ATCAGATCCTTACCGCTATGGGTAAGATTTGTAGGCTATCATTGCGATAAAGGAAAGGGGAGGACCGA
TGCTATATAAGGTTTTGCTATTCATTGAAAGCAGTAGTGACTGATTTGTATATA
>MinSyn_1332|Strength:0.000471137
GCGTGTCGTTTTAGTGAGGTTATGACCCCCGCCGATGACGCGGGAATTATGAAGATAAGATAATAATG
TTGAAGATAAGAACTGCTAGGAGGACCGATGCTGACAGTGGTCCCTCCACTGCACCTCACATGTAGGC
TATCAGCTTGTGGGAGCCACCAACAAAACTGGTACTTCAGAAGATCAAAGGGCTAAATTAGGTTGTCT
GCACCTCACATGATCGAAAGGACAGTAGACCACTATGCCCAATTAATCCTTACCGCTATGGGTAAGAT
TAGGACCGACTATATAAGGTTTTGCTATTCATTGAAAGCAGTAGTGACTGATTTGTATATA
>MinSyn_152|Strength:0.0004712
GCGTGTCGTTTTAGTGAGGTCACTATCAGCTACAAAACATTGCGATAAAGGAAAGGTCACTGCTAGGA
GCTATATAAGGTTTTGCTATTCATTGAAAGCAGTAGTGACTGATTTGTATATA
>MinSyn_1294|Strength:0.0004712
GCGTGTCGTTTTAGTGAGGCAAATATTTCTTGTACTGCTAGGACAGTGGTCCCTCCACCTATGCCCAA
TCTATATAAGGTTTTGCTATTCATTGAAAGCAGTAGTGACTGATTTGTATATA
>MinSyn_1468|Strength:0.000471788
GCGTGTCGTTTTAGTGAGGTCAGAAGATCAAAGGGCTAATGCTGATCTTGCCTGCCTTGATGATTCTC
TCTGCCGACAGTGGTCCCAAAGTCACGACCACTATGCCCAATTAGGTTGCAGTGGTCCCTCCACCTAT
CAGCTTAGCAAGACCTCTACAAAATTTTCAACAAAAGACCTCTACAATTGCGATAAAGGAAAGGCACT
GCTAGGAGGAAAAATGTCAAAGATAGGTTGTCTGCACCTCACATGTAAAGATTGATGAAAAGTCAAAA
ACAAAAATCAATTATCTCACTGCTAGGAGTGAAGATAAGATAATAATGTTGAAGATAAGATGCTAGGA
GGACCGATGCTGATCTTGCCCTATATAAGGTTTTGCTATTCATTGAAAGCAGTAGTGACTGATTTGTA
TATA
>MinSyn_1352|Strength:0.000472288
GCGTGTCGTTTTAGTGAGGAAGATTGATGAAAAGTCAAAAACAAAAATCAATTATGAAACCATTATTG
CGCCTCTACAAAAATTTCGGGAAACCTCCTCGCGATGCTGATCTTGCCTGCCTTGATTGAAGATAAGA
TAATAATGTTGAAGATAAGAGTGTACAGGGCTCACTGCTAGTTTTCAACAACTATCGAAAGGACAGTA
TGGTACTTGTGTACAGGGCTCACTGCTACTATATAAGGTTTTGCTATTCATTGAAAGCAGTAGTGACT
GATTTGTATATA
>MinSyn_128|Strength:0.000473289
GCGTGTCGTTTTAGTGAGGTGGTGGAGCACGACACCGATGCTGATCTTGCCAAGATTGATGAAAAGTC
AAAAACAAAAATCAATTATGTACTTGTGTACAGGGCTCACTGCTATCTCTCTGCCGACAGTGGTCCCA
AACTGATCTTGCAGCCACTTGTGTCATGTAGGCTATCAGCTTAGAATTTCGGGAAACCTCCTCGAGAC
CTCTACAACAGTGGTCCCTCCACCTACAAAACTGGTACTTGTGTACCTATATAAGGTTTTGCTATTCA
TTGAAAGCAGTAGTGACTGATTTGTATATA
>MinSyn_1353|Strength:0.000473452
GCGTGTCGTTTTAGTGAGGAAAAATGTCAAAGATATACTTGTGTACAGGGCTCAATCCTTACCGCTAT
GGGTAAGATTTAGCAAGACCTCGTGGGAGCCACCAGGCTCACTGCTATGGTGGAGCACGACACCTCTA
CAAACTATATAAGGTTTTGCTATTCATTGAAAGCAGTAGTGACTGATTTGTATATA
>MinSyn_1691|Strength:0.000474925
GCGTGTCGTTTTAGTGAGGAGCAAGTGGATGTCTGCACCTCACATGTAGATCGAAAGGACAGTATTAG
GTTGTCTGCACCAATTTCGGGAAACCTCCTCGGGCTATCAGCTTAGCAAGACTGAAGCATCTTCCTCA
GGAAAAAGAAGAGGTGACCTCTACAAAACTATTGCGATAAAGGAAAGGTGCTGATCTTGCCTATATAA
GGTTTTGCTATTCATTGAAAGCAGTAGTGACTGATTTGTATATA
>MinSyn_166|Strength:0.000475
GCGTGTCGTTTTAGTGAGGCAAATATTTCTTGTACGACCAGGAAAAAGAAGAGGTGTAGGTCACGACC
CTATATAAGGTTTTGCTATTCATTGAAAGCAGTAGTGACTGATTTGTATATA
>MinSyn_1919|Strength:0.000475547
GCGTGTCGTTTTAGTGAGGGTGGAGCCACCATCTGCACCAGTGGTCCCTCCACAAGACCTCTACAAA
ACTGGTACTTGTGAAAAATGTCAAAGATACTCACTGCTAGGAGGACGGAAAAAGAAGAGGTTGCACCT

GAAGATAAGATAATAATGTTGAAGATAAGAACCGATGCTGATCTTGCCCTATATAAGGTTTTGCTATT
CATTGAAAGCAGTAGTGACTGATTTGTATATA
>MinSyn_1476|Strength:0.000475555
GCGTGTCGTTTTAGTGAGGTCACTATCAGCCACGACCACTATGCCCAAGCACACCAGCATGTGTTGAT
CACCAGCTCTGCACCTCACATGTAGGCTTGAAGATAAGATAATAATGTTGAAGATAAGACTGATCTTG
CCTGCCTTGCTATATAAGGTTTTGCTATTCATTGAAAGCAGTAGTGACTGATTTGTATATA
>MinSyn_1104|Strength:0.000475659
GCGTGTCGTTTTAGTGAGGTCTCTCTGCCGACAGTGGTCCCAAAGCGGTAGGTCACAACCATTATTGC
GGACCACTATGCTTTGTCAAAAGCTAAAAAAGATGATGCAGGAGGACCGATGCTGATCTTGCCCTATA
TAAGGTTTTGCTATTCATTGAAAGCAGTAGTGACTGATTTGTATATA
>MinSyn_1744|Strength:0.000475817
GCGTGTCGTTTTAGTGAGGGTGGGAGCCACCACACTGCTTGAAGCATCTTCCAAACTGGTACTTGAAG
ATAAGATAATAATGTTGAAGATAAGAAGGGCTCACTGCTAGGAGGACCTATATAAGGTTTTGCTATTC
ATTGAAAGCAGTAGTGACTGATTTGTATATA
>MinSyn_1211|Strength:0.000476084
GCGTGTCGTTTTAGTGAGGTTATGACCCCCGCCGATGACGCGGGAACTGCTAGGAGGACCGATGCTGA
TCTTGCAACCACGTCTACAAGACCACTAAAGATTGATGAAAAGTCAAAAACAAAAATCAATTATCTGA
TCTGAAGCATCTTCCTAGGTCACGACCACTAAGGTGGCTCCTACTGTACATCACTATCAGCTGCTGAT
CTTGCCTGCCTTGATATTGCGATAAAGGAAAGGTAGGAGGACCGATGCCTATATAAGGTTTTGCTATT
CATTGAAAGCAGTAGTGACTGATTTGTATATA
>MinSyn_1304|Strength:0.000476522
GCGTGTCGTTTTAGTGAGGAGGTGGCTCCTACCACTATGCCCAATTAGGTTGTCTGCTTTGTCAAAAG
CTAAAAAAGATGATGCTATGCCCAATTAGGTGAAGCATCTTCCCCACTATGCCCAATTAGGTTGTCCT
ATATAAGGTTTTGCTATTCATTGAAAGCAGTAGTGACTGATTTGTATATA
>MinSyn_1276|Strength:0.000476687
GCGTGTCGTTTTAGTGAGGTGGTGGAGCACGACATCACATGTAGGCTTGAAGATAAGATAATAATGTT
GAAGATAAGAGGCTCACTGCTAGGAGGACCGATGCTGATAACCACGTCTACAACTGCTAGGAGGACCG
ATGAAAAATGTCAAAGATACCACTATGAGCAAGTGGATATCTTAATTTCGGGAAACCTCCTCGACGCA
CACCAGCATGTGTTGATCACCAGCTTCGCGGTAGGTCACGACCACTATGCCCCAAATATTTCTTGTGC
AAGACCTCTACAAAACTCTATATAAGGTTTTGCTATTCATTGAAAGCAGTAGTGACTGATTTGTATAT
A
>MinSyn_111|Strength:0.000477718
GCGTGTCGTTTTAGTGAGGTCACTATCAGCCTCTACAAAACTGGTACTTGTGAAGATTGATGAAAAGT
CAAAAACAAAAATCAATTATTGATCTTGCAAAAATGTCAAAGATACCTTGATGATCAGCCACTTGTGT
CTATGCCCAATTAAACCATTATTGCGTCACGACCACTATTTTCAACAAGTACAGGGCTCACCTATATA
AGGTTTTGCTATTCATTGAAAGCAGTAGTGACTGATTTGTATATA
>MinSyn_1771|Strength:0.000478862
GCGTGTCGTTTTAGTGAGGATCGAAAGGACAGTAATCTCAAATATTTCTTGTCGACCACTATGCCCAC
TATATAAGGTTTTGCTATTCATTGAAAGCAGTAGTGACTGATTTGTATATA
>MinSyn_1419|Strength:0.000479029
GCGTGTCGTTTTAGTGAGGGCACACCAGCATGTGTTGATCACCAGCTGTTGTCTGCACCTCACATGGA
AAAAGAAGAGGTATGTAGGCTATCAGCTTAAAGATTGATGAAAAGTCAAAAACAAAAATCAATTATTA
CAAAACTGGTACTTGCAGTGGTCCCTCCACTGGTACTTGTGTATGAAGATAAGATAATAATGTTGAAG
ATAAGACTTGTGTACAGGGCTCACTGCTAGGAGGATCCATCAACAAATAATCCAAGTAAGGTTGTCTG
CACCTCACATGTAGGCTATCAAGGTGGCTCCTACCCCAATTAGGTTGTAAAAATGTCAAAGATACTGA
TCTTGCTATATAAGGTTTTGCTATTCATTGAAAGCAGTAGTGACTGATTTGTATATA
>MinSyn_1778|Strength:0.00047947
GCGTGTCGTTTTAGTGAGGTGGTGGAGCACGACAAGGTTGTCTGCACCTCAACCATTATTGCGGGCTC
ACTGCTAGGAGGACCGATGCTGAAGATAAGATAATAATGTTGAAGATAAGAGACCACTATGCCCAATT
AGGAATTTCGGGAAACCTCCTCGTGTGTACAGGGCTCACTGCTAGGAGGACCGGCTTTGTCAAAAGCT
AAAAAAGATGATGCTAGCAAGACCTCTACAATCCTTACCGCTATGGGTAAGATTGGACCGATGCTCTA
TATAAGGTTTTGCTATTCATTGAAAGCAGTAGTGACTGATTTGTATATA
>MinSyn_1663|Strength:0.000479697
GCGTGTCGTTTTAGTGAGGCAAATATTTCTTGTACTAAGATTGATGAAAAGTCAAAAACAAAAATCAA

TTATCTAGGAGGACCGATGCTGATGTGGGAGCCACCAAAACCATTATTGCGCTTTTCAACAACTATCA
GCTTAGCAAGAATTGCGATAAAGGAAAGGCATGTAGGCTATCAGCTTAGCACTATATAAGGTTTTGCT
ATTCATTGAAAGCAGTAGTGACTGATTTGTATATA
>MinSyn_1617|Strength:0.000480272
GCGTGTCGTTTTAGTGAGGAACCACGTCTACAACAGCAAGATTGATGAAAAGTCAAAAACAAAAATCA
ATTATCCTCACATGTAGGCTATCAGTGAAGATAAGATAATAATGTTGAAGATAAGAGCTATCAGCTTA
GCAAGACCTCTATCCATCAACAAATAATCCAAGTAAGGCGTCTCTCTGCCGACAGTGGTCCCAAATGG
TACTTGTGTACAGGGCTCACTGCTAGCAAGTGGATCTAGGAGGACCGAGTGGGAGCCACCATTGTCTG
CACCTCAATTGCGATAAAGGAAAGGCTATCAGCTTAGCAAGACCTCTACAAAACTTTTCAACAAAGGC
CTATATAAGGTTTTGCTATTCATTGAAAGCAGTAGTGACTGATTTGTATATA
>MinSyn_1734|Strength:0.00048031
GCGTGTCGTTTTAGTGAGGTTATGACCCCCGCCGATGACGCGGGATGAACCATTATTGCGTCACATGT
AGGCTAATCGAAAGGACAGTATAGGAGGACCGATGCTGATCACTATCAGCTCACTTCAGAAGATCAAA
GGGCTACTACAAAACTGGTACTTGCTATATAAGGTTTTGCTATTCATTGAAAGCAGTAGTGACTGATT
TGTATATA
>MinSyn_1400|Strength:0.000480627
GCGTGTCGTTTTAGTGAGGTTATGACCCCCGCCGATGACGCGGGATACATGAAGCATCTTCCTCACTG
CTAGGTCCATCAACAAATAATCCAAGTAAGCCTCTACAAAACTGGTACTTGCTATATAAGGTTTTGCT
ATTCATTGAAAGCAGTAGTGACTGATTTGTATATA
>MinSyn_1344|Strength:0.000480865
GCGTGTCGTTTTAGTGAGGTGAAGCATCTTCCTGTCTGCACCTCACATGTAGTCAGAAGATCAAAGGG
CTAACTCTCTCTGCCGACAGTGGTCCCAAATGCACCTCACATGTAGGCTTCACTATCAGCTACAGGGC
TCACTGCTAGGAGGACCTTATGACCCCCGCCGATGACGCGGGATTAGGTTGTCTGCACCTCACATGTA
GGCTCAGTGGTCCCTCCACAGGCTATCAAGGTGGCTCCTACAGGACGCACACCAGCATGTGTTGATCA
CCAGCTAACTGGTACTTGTGTACAGGCTATATAAGGTTTTGCTATTCATTGAAAGCAGTAGTGACTGA
TTTGTATATA
>MinSyn_114|Strength:0.000480868
GCGTGTCGTTTTAGTGAGGATCCTTACCGCTATGGGTAAGATTCGGTGGGAGCCACCATGATCTTGCC
TGCCTTGATCAAATATTTCTTGTCTGCACCTGCACACCAGCATGTGTTGATCACCAGCTTGTGTACAG
GGCTCTTTTCAACAAGTACAGGGCTCACTGCTATCACTATCAGCTCTGCACTATATAAGGTTTTGCTA
TTCATTGAAAGCAGTAGTGACTGATTTGTATATA
>MinSyn_172|Strength:0.000481204
GCGTGTCGTTTTAGTGAGGATCCTTACCGCTATGGGTAAGATTGTAGGTCACGACCACTATGCCCAAT
TCTCTCTGCCGACAGTGGTCCCAAATCTATATAAGGTTTTGCTATTCATTGAAAGCAGTAGTGACTGA
TTTGTATATA
>MinSyn_1987|Strength:0.000481914
GCGTGTCGTTTTAGTGAGGAGCAAGTGGATGCGGTAGGTCACTATCAGCAGGCTATCAGCAACCATTA
TTGCGCTCGCTTTGTCAAAAGCTAAAAAAGATGATGCACCTCACATGTAGGGAAAAAGAAGAGGTTCT
ACAAAACTGGTACTTAATTTCGGGAAACCTCCTCGCTCACATGTAGGCTCTATATAAGGTTTTGCTAT
TCATTGAAAGCAGTAGTGACTGATTTGTATATA
>MinSyn_1624|Strength:0.000482787
GCGTGTCGTTTTAGTGAGGAAAAATGTCAAAGATAGATGAACCATTATTGCGACCGATGCTGATCTCT
ATATAAGGTTTTGCTATTCATTGAAAGCAGTAGTGACTGATTTGTATATA
>MinSyn_1854|Strength:0.000483051
GCGTGTCGTTTTAGTGAGGGGAAAAAGAAGAGGTAGGTGGGAGCCACCACATGTAGGCTATCCTATAT
AAGGTTTTGCTATTCATTGAAAGCAGTAGTGACTGATTTGTATATA
>MinSyn_1321|Strength:0.000484043
GCGTGTCGTTTTAGTGAGGTCTCTCTGCCGACAGTGGTCCCAAATCATCCTTACCGCTATGGGTAAGA
TTGGAGGAAGCAAGTGGATTCAAGATTGATGAAAAGTCAAAAACAAAAATCAATTATCACGACCACTA
TGCCCAATCAGAAGATCAAAGGGCTATAGGTTGTCTGCACCTCACATGTAGGCTTATGACCCCCGCCG
ATGACGCGGGAGGACCGATGCTGATCTTGCTTTTCAACAATCTACAAAAGGTGGCTCCTACCCACTAT
GCCCAATTAGGTTGTCTGCACCCAGCCACTTGTGTGATGATCTATATAAGGTTTTGCTATTCATTGAA
AGCAGTAGTGACTGATTTGTATATA
>MinSyn_1129|Strength:0.000484503

GCGTGTCGTTTTAGTGAGGATCCTTACCGCTATGGGTAAGATTAAGACCTCTACAAAATTATGACCCC
CGCCGATGACGCGGGACTGCTAGGAGGAAGCAAGTGGATTACAAAACTGGTACTTGTGTACAGGGCTC
AGCTTTGTCAAAAGCTAAAAAAGATGATGCACTGCTAGGAGGACCGATGCTGATCTGGTGGAGCACGA
CAAGGAGGACCGATGCTGATGAAGCATCTTCCATCAGAAGATCAAAGGGCTAATTAGGTTGTCTGCTA
TATAAGGTTTTGCTATTCATTGAAAGCAGTAGTGACTGATTTGTATATA
>MinSyn_1869|Strength:0.000485092
GCGTGTCGTTTTAGTGAGGTGGTGGAGCACGACACACCAGCCACTTGTGTTGGTACTTGTGTACAGGG
CTCACTATCCTTACCGCTATGGGTAAGATTCTACAAAACTGGTACTTGTAACCACGTCTACAATGCAC
CTTATGACCCCCGCCGATGACGCGGGAGGGCTCACTGCTCACTATCAGCGGTCACGACCACTATGCCC
AATCTATATAAGGTTTTGCTATTCATTGAAAGCAGTAGTGACTGATTTGTATATA
>MinSyn_1662|Strength:0.000485092
GCGTGTCGTTTTAGTGAGGAGGTGGCTCCTACACAAAACTGGTATTATGACCCCCGCCGATGACGCGG
GACAGCTTAGCAAGACCTCTACAATCTCTCTGCCGACAGTGGTCCCAAAGGAGGACTATATAAGGTTT
TGCTATTCATTGAAAGCAGTAGTGACTGATTTGTATATA
>MinSyn_1788|Strength:0.000485114
GCGTGTCGTTTTAGTGAGGGCACACCAGCATGTGTTGATCACCAGCTGGTTGTCTGTCTCTCTGCCGA
CAGTGGTCCCAAAAAAACTGGTACTTGTGTACACTATATAAGGTTTTGCTATTCATTGAAAGCAGTAG
TGACTGATTTGTATATA
>MinSyn_1840|Strength:0.00048517
GCGTGTCGTTTTAGTGAGGAAAAATGTCAAAGATAAGCAAGACCTCTACAAATCCTTACCGCTATGGG
TAAGATTATCTTGCCTTATGACCCCCGCCGATGACGCGGGAGCAAGACCTCTACAAAACTCTATATAA
GGTTTTGCTATTCATTGAAAGCAGTAGTGACTGATTTGTATATA
>MinSyn_1473|Strength:0.000485175
GCGTGTCGTTTTAGTGAGGGTGGGAGCCACCACTGCACCTCACATGTAGGCTATCAGCTGCTTTGTCA
AAAGCTAAAAAAGATGATGCTGAAGATTGATGAAAAGTCAAAAACAAAAATCAATTATAGGACCGATT
GGTGGAGCACGACAACTGGTACTTGTGTACAGCAAATATTTCTTGTGCCTGCCTTGATGATTCAGAAG
ATCAAAGGGCTAGTACAGGGCTCACTGCTAGGAGCTATATAAGGTTTTGCTATTCATTGAAAGCAGTA
GTGACTGATTTGTATATA
>MinSyn_1899|Strength:0.000485401
GCGTGTCGTTTTAGTGAGGTGGTGGAGCACGACATGCACCTATTGCGATAAAGGAAAGGGGTTGTCTG
CACCTCACATGTACTATATAAGGTTTTGCTATTCATTGAAAGCAGTAGTGACTGATTTGTATATA
>MinSyn_1999|Strength:0.000485401
GCGTGTCGTTTTAGTGAGGAACCACGTCTACAACTACAAAACTGGTACTTGTGTATCAGAAGATCAAA
GGGCTACGCGGTACTATATAAGGTTTTGCTATTCATTGAAAGCAGTAGTGACTGATTTGTATATA
>MinSyn_1497|Strength:0.000486019
GCGTGTCGTTTTAGTGAGGTGAAGATAAGATAATAATGTTGAAGATAAGAGTAGGCTATCAGCTTAGC
AAGACCTCAATTTCGGGAAACCTCCTCGAGACCTCTACAAAACTGGAAAAATGTCAAAGATATTGTCT
GCACCTCACATGTAGGCTTATGACCCCCGCCGATGACGCGGGAACGACCAGCTTTGTCAAAAGCTAAA
AAAGATGATGCCCTCACATGTAGGCTATCAGCTTACTATATAAGGTTTTGCTATTCATTGAAAGCAGT
AGTGACTGATTTGTATATA
>MinSyn_1631|Strength:0.000486145
GCGTGTCGTTTTAGTGAGGATGACGTAAGCCATGACGTCTACAAGACCTCTACAAAACTGGTACTTGT
GTTATGACCCCCGCCGATGACGCGGGAGATCTTGCCTGCCTTGATAACCACGTCTACAAGTGTACAGG
GCTCACTGCTAGGAGGATGAAGCATCTTCCAGGAGGAGCACACCAGCATGTGTTGATCACCAGCTGGT
AGGTCACGACCACTATGAAGATTGATGAAAAGTCAAAAACAAAAATCAATTATCCAATTAGGTTGTCT
TTTCAACAAGCAAGACCTCTACAAAACTGGTATTGCGATAAAGGAAAGGACTGCTAGGAGCAAGTGGA
TCTAGGAGGACCGATGCTATATAAGGTTTTGCTATTCATTGAAAGCAGTAGTGACTGATTTGTATATA
>MinSyn_1808|Strength:0.000486183
GCGTGTCGTTTTAGTGAGGAACCACGTCTACAACAAAACCATTATTGCGGCACCTCACATGTAGGCTA
TTGGTGGAGCACGACACACTGCTAGGAGGACAGCAAGTGGATCGCGGTAGGTCACGACCACTATGCCC
AAGCTTTGTCAAAAGCTAAAAAAGATGATGCATGCCCAATTAGGTTGTCTGCTGAAGATAAGATAATA
ATGTTGAAGATAAGATGCTGTGAAGCATCTTCCGCCCAATTAGGTTGTCTGCACCTCTATATAAGGTT
TTGCTATTCATTGAAAGCAGTAGTGACTGATTTGTATATA
>MinSyn_1535|Strength:0.000486371

GCGTGTCGTTTTAGTGAGGGCTTTGTCAAAAGCTAAAAAAGATGATGCGGCTATCAGCTTAGCAAGAC
CTCTCTCTCTGCCGACAGTGGTCCCAAATTGTAGGTGGCTCCTACTAGGCTATCAGCTTAGCAAGACC
TCTACACTATATAAGGTTTTGCTATTCATTGAAAGCAGTAGTGACTGATTTGTATATA
>MinSyn_1600|Strength:0.000486475
GCGTGTCGTTTTAGTGAGGGTGGGAGCCACCAGTCACGACCACTATGCCCAATTAGGCAGTGGTCCCT
CCACGCATCAGAAGATCAAAGGGCTACGCGGTAGGCTATATAAGGTTTTGCTATTCATTGAAAGCAGT
AGTGACTGATTTGTATATA
>MinSyn_1642|Strength:0.000486495
GCGTGTCGTTTTAGTGAGGGGAAAAAGAAGAGGTGGACCGATGCTGATCTTGCCTGCCTTGATTCCAT
CAACAAATAATCCAAGTAAGGGCTAGGTGGCTCCTACGGAGGACCGATGCTGATCCAGTGGTCCCTCC
ACCCGATGCTGATCTTGCCTGCCAAGATTGATGAAAAGTCAAAAACAAAAATCAATTATCGATGCTGT
GGTGGAGCACGACAGGTAGGTCACGACCACTATGCAACCATTATTGCGACCTCACATGTAGGCTATAT
AAGGTTTTGCTATTCATTGAAAGCAGTAGTGACTGATTTGTATATA
>MinSyn_1406|Strength:0.000487535
GCGTGTCGTTTTAGTGAGGCAGTGGTCCCTCCACTCACGACCACTATTCCATCAACAAATAATCCAAG
TAAGTCACTGCTATCTCTCTGCCGACAGTGGTCCCAAAGACCGATGCTGATCTTGCCTATATAAGGTT
TTGCTATTCATTGAAAGCAGTAGTGACTGATTTGTATATA
>MinSyn_1333|Strength:0.000487611
GCGTGTCGTTTTAGTGAGGCAAATATTTCTTGTTTGTGGGAGCCACCAAAAACTGGTCTATATAAGGT
TTTGCTATTCATTGAAAGCAGTAGTGACTGATTTGTATATA
>MinSyn_1273|Strength:0.000487876
GCGTGTCGTTTTAGTGAGGGTGGGAGCCACCAAGACCTCTACAAAACTGGTGCACACCAGCATGTGTT
GATCACCAGCTTTAGGTTATTGCGATAAAGGAAAGGGGTCACGACCACTATGTGAAGATAAGATAATA
ATGTTGAAGATAAGACCTCACATGTAGGCTATCAGCTTAGCTATATAAGGTTTTGCTATTCATTGAAA
GCAGTAGTGACTGATTTGTATATA
>MinSyn_117|Strength:0.000488036
GCGTGTCGTTTTAGTGAGGTGAAGCATCTTCCAGACCTCTACAAAACTGGTACTTGTGTAATTGCGAT
AAAGGAAAGGTAGGAGGACCGATGCTCAGTGGTCCCTCCACACCGATGCTGATCTCAGAAGATCAAAG
GGCTAGGAGGACCGATGCTGTGAAGATAAGATAATAATGTTGAAGATAAGACTATTTTCAACAAAGGT
CACGACCACTATGCCCAATTAGCTATATAAGGTTTTGCTATTCATTGAAAGCAGTAGTGACTGATTTG
TATATA
>MinSyn_1143|Strength:0.000488286
GCGTGTCGTTTTAGTGAGGCAAATATTTCTTGTCACGACCACAGCCACTTGTGTTGCCCAATTAGGCA
GTGGTCCCTCCACTGCTCTATATAAGGTTTTGCTATTCATTGAAAGCAGTAGTGACTGATTTGTATAT
A
>MinSyn_1343|Strength:0.000488971
GCGTGTCGTTTTAGTGAGGAATTTCGGGAAACCTCCTCGACGACCACTGTGGGAGCCACCAAGGACCG
ATGCTGATCTTGCTATATAAGGTTTTGCTATTCATTGAAAGCAGTAGTGACTGATTTGTATATA
>MinSyn_1282|Strength:0.000490274
GCGTGTCGTTTTAGTGAGGAAGATTGATGAAAAGTCAAAAACAAAAATCAATTATCTCTACAAAACTG
GTACTTGAGGTGGCTCCTACTAGGCTATCAGCTTAGCAACAAATATTTCTTGTGGCTATATAAGGTTT
TGCTATTCATTGAAAGCAGTAGTGACTGATTTGTATATA
>MinSyn_1399|Strength:0.000490325
GCGTGTCGTTTTAGTGAGGGCTTTGTCAAAAGCTAAAAAAGATGATGCGACCGATGCTGATGAAGATA
AGATAATAATGTTGAAGATAAGAACAGGGCTCAGTGGGAGCCACCAACTATGCCCAATAATTTCGGGA
AACCTCCTCGGACCTCTACAAAACTGGTACTTGTGTTTTCAACAAACTTGTGTACAGGGCTCACTGCT
GCACACCAGCATGTGTTGATCACCAGCTTAGGAGGACCGATGCTGATCTCAGCCACTTGTGTAAAACT
GTCAGAAGATCAAAGGGCTAGTCACGACCACTACTATATAAGGTTTTGCTATTCATTGAAAGCAGTAG
TGACTGATTTGTATATA
>MinSyn_1712|Strength:0.000491126
GCGTGTCGTTTTAGTGAGGTTATGACCCCCGCCGATGACGCGGGAGGAGGACCGATGCTGATCTTGCA
ACCATTATTGCGTCTACAAAACATTGCGATAAAGGAAAGGGTCTGCACCTCACATGTAGGCTATCAGC
TATCGAAAGGACAGTAGAACCACGTCTACAAACTATATAAGGTTTTGCTATTCATTGAAAGCAGTAGT
GACTGATTTGTATATA

>MinSyn_1989|Strength:0.000491193
GCGTGTCGTTTTAGTGAGGTCTCTCTGCCGACAGTGGTCCCAAAACTTGTGTACAGGGCTCACTGCTT
GGTGGAGCACGACAATGCTGATCTTGCCTGCCTTGATGGCACACCAGCATGTGTTGATCACCAGCTTT
AGCAAGAACCATTATTGCGACAGGGCTCACTGCCTATATAAGGTTTTGCTATTCATTGAAAGCAGTAG
TGACTGATTTGTATATA
>MinSyn_1623|Strength:0.000491487
GCGTGTCGTTTTAGTGAGGATCGAAAGGACAGTAGTAGGTCACGACCACTTTTCAACAATGGTACGTG
GGAGCCACCAGGCTATCAGCTTAGCAAGACCTCTCAGTGGTCCCTCCACTTGTGTACAGGGCTCACTG
CTAGGAGTCACTATCAGCCATGTCTCTCTGCCGACAGTGGTCCCAAAGTACTTGTGTACAGGGAAGAT
TGATGAAAAGTCAAAAACAAAAATCAATTATGGCTATCAGCTTAGCAAGACTATATAAGGTTTTGCTA
TTCATTGAAAGCAGTAGTGACTGATTTGTATATA
>MinSyn_1427|Strength:0.000492338
GCGTGTCGTTTTAGTGAGGGTGGGAGCCACCAATCAATTTCGGGAAACCTCCTCGGTAGGTCACGACA
TCCTTACCGCTATGGGTAAGATTGCTATCAGGTGGCTCCTACGCAAGACCTCTACAAAATGGTGGAGC
ACGACATGCTCAAATATTTCTTGTGGCTCACTGCTAGGAGGACCGTTTTCAACAAGCCCAATTAGGTT
GTCTGCACCTAAGATTGATGAAAAGTCAAAAACAAAAATCAATTATGTTGTCTGCACCTCACACTATA
TAAGGTTTTGCTATTCATTGAAAGCAGTAGTGACTGATTTGTATATA
>MinSyn_1482|Strength:0.000492826
GCGTGTCGTTTTAGTGAGGTGAAGATAAGATAATAATGTTGAAGATAAGAATCAGCTTAGCAAGACCT
CTACATCCATCAACAAATAATCCAAGTAAGTTGTCTGCACCTCACATGTAGGCTACAGCCACTTGTGT
ACCACTATGCCCAATTAGGTTTGGTGGAGCACGACAGACCTCTACAAAACTGGTCTCTCTGCCGACAG
TGGTCCCAAAGTACAGGGCTCACTGCTAGGAGGACCGATAACCACGTCTACAAGTCGGAAAAAGAAGA
GGTCTCTACAAAATCCTTACCGCTATGGGTAAGATTGCCTGCCTTGATGATCTATATAAGGTTTTGCT
ATTCATTGAAAGCAGTAGTGACTGATTTGTATATA
>MinSyn_1397|Strength:0.000493562
GCGTGTCGTTTTAGTGAGGAAAAATGTCAAAGATATAGGTCACGACCACTATGCCCAATTAGGTTCAG
AAGATCAAAGGGCTACTCACTGCTAGGATCTCTCTGCCGACAGTGGTCCCAAACTCACATGTAGGCTA
TGCTTTGTCAAAAGCTAAAAAAGATGATGCGCTAGGAGGACCGATGCTGAAACCACGTCTACAACCGA
TGCTGAGCACACCAGCATGTGTTGATCACCAGCTACTATGCCCAATTAGGTTGTGAAGATAAGATAAT
AATGTTGAAGATAAGAGCCCAATTAGGTTGTCTGCACCTCACATGCTATATAAGGTTTTGCTATTCAT
TGAAAGCAGTAGTGACTGATTTGTATATA
>MinSyn_1966|Strength:0.000494034
GCGTGTCGTTTTAGTGAGGCAGCCACTTGTGTGCGGTAGGTCAATCGAAAGGACAGTATCACGACCAC
TATGCCCAATTAGAGGTGGCTCCTACACCTCACATGTAGGCTATCAGCTTAGCAATGAAGATAAGATA
ATAATGTTGAAGATAAGAGCTAGGAGGACCGATGCTGATCTTGCCTGCAACCACGTCTACAAGCTATC
AGCTTAGCGTGGGAGCCACCAGGAGGACCGATGCTGATCTTGCCTGCCTCTCTCTGCCGACAGTGGTC
CCAAACTATGCCCAATTAGGAAGATTGATGAAAAGTCAAAAACAAAAATCAATTATGCTACTATATAA
GGTTTTGCTATTCATTGAAAGCAGTAGTGACTGATTTGTATATA
>MinSyn_1906|Strength:0.000494188
GCGTGTCGTTTTAGTGAGGCAGCCACTTGTGTGGTTGTCTGCATTTTCAACAACACTATGCCCAATTA
GGTTGTCTGCGCACACCAGCATGTGTTGATCACCAGCTCTACAAAACTGGTACTTGTGTACTGGTGGA
GCACGACAGGCTATATAAGGTTTTGCTATTCATTGAAAGCAGTAGTGACTGATTTGTATATA
>MinSyn_1387|Strength:0.000494208
GCGTGTCGTTTTAGTGAGGGCACACCAGCATGTGTTGATCACCAGCTGTCTGCACCTCACATGAAGAT
TGATGAAAAGTCAAAAACAAAAATCAATTATAGGAGGACCGATGCTGATCTTGCCTGCCTTCTATATA
AGGTTTTGCTATTCATTGAAAGCAGTAGTGACTGATTTGTATATA
>MinSyn_1554|Strength:0.000494573
GCGTGTCGTTTTAGTGAGGAAGATTGATGAAAAGTCAAAAACAAAAATCAATTATAACTGGTATGAAG
ATAAGATAATAATGTTGAAGATAAGAAGGTTGTCTGGAAAAAGAAGAGGTTGTCTGCACTCCATCAAC
AAATAATCCAAGTAAGTGGTACTTGTTGAAGCATCTTCCGCCCAATTAGGTTGTCTGCACCTCACATT
CTCTCTGCCGACAGTGGTCCCAAAAACTGGTACTTGTGTACAGGAACCACGTCTACAACACCTCACAT
GTAGGCTATCAGCTGCTTTGTCAAAAGCTAAAAAAGATGATGCGCTGATCTTGCCTGCCTCTATATAA
GGTTTTGCTATTCATTGAAAGCAGTAGTGACTGATTTGTATATA
>MinSyn_1837|Strength:0.000494766

GCGTGTCGTTTTAGTGAGGGCTTTGTCAAAAGCTAAAAAAGATGATGCGATGCTGATCTTGCCTGCCT
TGATCACTATCAGCACTGGTACTTGTGTACAGGGCTCACTGCATTGCGATAAAGGAAAGGTTAGCAAG
ACCTCTACAAAACTGGTTGGTGGAGCACGACATAGGCTATCAGCTTAGAAGATTGATGAAAAGTCAAA
AACAAAAATCAATTATATCTTGCCTGGCACACCAGCATGTGTTGATCACCAGCTGACTGAAGATAAGA
TAATAATGTTGAAGATAAGACGGTAGGTCACGACCACTATGCCCTGAAGCATCTTCCCGACCACTATG
CCCAATTAGGTTGTCTATATAAGGTTTTGCTATTCATTGAAAGCAGTAGTGACTGATTTGTATATA
>MinSyn_1998|Strength:0.000495183
GCGTGTCGTTTTAGTGAGGGTGGGAGCCACCATGATCTTGCGCTTTGTCAAAAGCTAAAAAAGATGAT
GCACAGGGCTCACTGCTAGTCAGAAGATCAAAGGGCTACCTATCGAAAGGACAGTAATTAGGTTGTCT
GCACCTCACATGTACTATATAAGGTTTTGCTATTCATTGAAAGCAGTAGTGACTGATTTGTATATA
>MinSyn_1942|Strength:0.000495811
GCGTGTCGTTTTAGTGAGGAAGATTGATGAAAAGTCAAAAACAAAAATCAATTATTGCTAGGAGGACG
TGGGAGCCACCAAGCAAAAAAATGTCAAAGATACAAGACCTCTACAAAACTGGTACTATCCTTACCGC
TATGGGTAAGATTCCACTATGCCCCTATATAAGGTTTTGCTATTCATTGAAAGCAGTAGTGACTGATT
TGTATATA
>MinSyn_1168|Strength:0.000496149
GCGTGTCGTTTTAGTGAGGATCCTTACCGCTATGGGTAAGATTTGTGAAGCATCTTCCTGCACACCAG
CATGTGTTGATCACCAGCTGACCACTATGCCCAATTAGGTTGATCGAAAGGACAGTATGTCCATCAAC
AAATAATCCAAGTAAGAGCAAGACCTCTACAAAACAAAAATGTCAAAGATATGTCTGCACCTCACATG
TAGGGTGGGAGCCACCACTCACATGTGGAAAAAGAAGAGGTACAGGGCTCACTGCTAGGAGGTGGCTC
CTACCCTTGATGATCTATATAAGGTTTTGCTATTCATTGAAAGCAGTAGTGACTGATTTGTATATA
>MinSyn_1101|Strength:0.00049649
GCGTGTCGTTTTAGTGAGGGTGGGAGCCACCATAGGAGGAACCATTATTGCGCTGCACCTCACTTATG
ACCCCCGCCGATGACGCGGGAGCTGATCTTGCTTTGTCAAAAGCTAAAAAAGATGATGCAAACTGGTA
CTTGTGTACAGGGCTCCTATATAAGGTTTTGCTATTCATTGAAAGCAGTAGTGACTGATTTGTATATA
>MinSyn_1756|Strength:0.000496767
GCGTGTCGTTTTAGTGAGGAAAAATGTCAAAGATAACCATGGTGGAGCACGACACGGTAGGTCACGAC
CACTATGCCCAATTACAGTGGTCCCTCCACCTATGCCCAATTAGGTTGTCTGCACCTCAGGTGGCTCC
TACGCCTTGATGATAACCACGTCTACAAAGCTTAGGCACACCAGCATGTGTTGATCACCAGCTGCTTA
GCACTATATAAGGTTTTGCTATTCATTGAAAGCAGTAGTGACTGATTTGTATATA
>MinSyn_1695|Strength:0.000496809
GCGTGTCGTTTTAGTGAGGTTATGACCCCCGCCGATGACGCGGGAAGGACCGATGCTTGAAGCATCTT
CCTCTACAAAACTGGTACTTGTGCTTTGTCAAAAGCTAAAAAAGATGATGCTATGCCCAATTAGGTTG
TCTGCACCTTGAAGATAAGATAATAATGTTGAAGATAAGAAGGCTATCAGCTTAGCAAGACCTCTACA
AAGCACACCAGCATGTGTTGATCACCAGCTCAAGACCTATATAAGGTTTTGCTATTCATTGAAAGCAG
TAGTGACTGATTTGTATATA
>MinSyn_1213|Strength:0.000496953
GCGTGTCGTTTTAGTGAGGGGAAAAAGAAGAGGTGCACCTCACATGTAGGCTATCCAGCCACTTGTGT
CCGATTTTCAACAACGCGGTAGGTCACGACCACTAATTGCGATAAAGGAAAGGCTGATGGTGGAGCAC
GACACTATCTATATAAGGTTTTGCTATTCATTGAAAGCAGTAGTGACTGATTTGTATATA
>MinSyn_1831|Strength:0.000496976
GCGTGTCGTTTTAGTGAGGAATTTCGGGAAACCTCCTCGGGCTATAAGATTGATGAAAAGTCAAAAAC
AAAAATCAATTATAAACTGGTACTTGTGTACAGGGCTCAACCACGTCTACAAGCAAGACCTCTACAAA
ACTGGTACTCAAATATTTCTTGTCTCTACAAAATTATGACCCCCGCCGATGACGCGGGACTGATCTTG
CCTGCCTTGATGATTCCATCAACAAATAATCCAAGTAAGACCGATGCTATATAAGGTTTTGCTATTCA
TTGAAAGCAGTAGTGACTGATTTGTATATA
>MinSyn_1982|Strength:0.000497211
GCGTGTCGTTTTAGTGAGGAACCACGTCTACAAATGTAGGCTATCAGCGCACACCAGCATGTGTTGAT
CACCAGCTAATTAGAAAAATGTCAAAGATACAGGGCTCACTGCTAGGAGTGGTGGAGCACGACATGTA
CAGGGCTCACTGCTACAGCCACTTGTGTGGGAATTTCGGGAAACCTCCTCGGCACCTCACATGTAGGC
TATATAAGGTTTTGCTATTCATTGAAAGCAGTAGTGACTGATTTGTATATA
>MinSyn_1996|Strength:0.00049876
GCGTGTCGTTTTAGTGAGGGTGGGAGCCACCAGGTCACGACCACTATGCCCAATTAGGTTGTAAAAAT
GTCAAAGATACTTGCCTGCCTTGATGATCGAAAGGACAGTAAGCAAGACCTCTATTTTCAACAACGTC

AGAAGATCAAAGGGCTATGTGTACAGGGCTCACTGCTATCACTATCAGCGCTTAGCAAGACCTCTACA
AAACTCAGCCACTTGTGTCAGCTTAGCAAGACCTCTACAAATCCATCAACAAATAATCCAAGTAAGGC
ACCTCACATCCTTACCGCTATGGGTAAGATTGCTCCTATATAAGGTTTTGCTATTCATTGAAAGCAGT
AGTGACTGATTTGTATATA
>MinSyn_1584|Strength:0.000499458
GCGTGTCGTTTTAGTGAGGCAGCCACTTGTGTAATCACTATCAGCCGCGGTAGGTCACGACCACTATT
TTCAACAACCTCTACAAAACTGGTAATCGAAAGGACAGTAGTGTACAGGGCTCAATCCTTACCGCTAT
GGGTAAGATTACTCTATATAAGGTTTTGCTATTCATTGAAAGCAGTAGTGACTGATTTGTATATA
>MinSyn_169|Strength:0.000499512
GCGTGTCGTTTTAGTGAGGATTGCGATAAAGGAAAGGTGCCTTGATTGGTGGAGCACGACAATGCCCA
ATTCCATCAACAAATAATCCAAGTAAGGTAGGCTATCAGCTTTGTCAAAAGCTAAAAAAGATGATGCG
CGGTAGGTCACGACCACTATGCCCCTATATAAGGTTTTGCTATTCATTGAAAGCAGTAGTGACTGATT
TGTATATA
>MinSyn_1164|Strength:0.000499977
GCGTGTCGTTTTAGTGAGGAAAAATGTCAAAGATACTTAGCAAGACCTCTGCTTTGTCAAAAGCTAAA
AAAGATGATGCGAGGACCCTATATAAGGTTTTGCTATTCATTGAAAGCAGTAGTGACTGATTTGTATA
TA
>MinSyn_1586|Strength:0.000500124
GCGTGTCGTTTTAGTGAGGAACCATTATTGCGACTGCTAGGAGGACCGATGCTGAGCACACCAGCATG
TGTTGATCACCAGCTCGCGGTAGGTCATCAGAAGATCAAAGGGCTAACCGATGCTGATCTTGCCTGCC
TTGATGTCCATCAACAAATAATCCAAGTAAGTCTACAAAACTGGTACTTGATCCTTACCGCTATGGGT
AAGATTACAGGGCTCACTGCTAGGAGGATCGAAAGGACAGTACACTATATAAGGTTTTGCTATTCATT
GAAAGCAGTAGTGACTGATTTGTATATA
>MinSyn_1420|Strength:0.000500832
GCGTGTCGTTTTAGTGAGGTCTCTCTGCCGACAGTGGTCCCAAACCAATTAGGTTGTCTGCACCTCTT
TTCAACAATGCTGGAAAAAGAAGAGGTACTATGCCCAATTAGGTTGTCTGCCAGTGGTCCCTCCACCT
TGTGTACAGGGAAAAATGTCAAAGATACCTCACAATTGCGATAAAGGAAAGGGTGTACCTATATAAGG
TTTTGCTATTCATTGAAAGCAGTAGTGACTGATTTGTATATA
>MinSyn_1218|Strength:0.00050086
GCGTGTCGTTTTAGTGAGGCAGTGGTCCCTCCACGGTACTTGTGTACAGGCTTTGTCAAAAGCTAAAA
AAGATGATGCGGGCTCACTGCTAGGAGGACCGTTATGACCCCCGCCGATGACGCGGGAACCTCTCTAT
ATAAGGTTTTGCTATTCATTGAAAGCAGTAGTGACTGATTTGTATATA
>MinSyn_1764|Strength:0.000501227
GCGTGTCGTTTTAGTGAGGAATTTCGGGAAACCTCCTCGGATGCGCTTTGTCAAAAGCTAAAAAAGAT
GATGCCGTGGGAGCCACCAGTGTACAGGGCTCACTGTTATGACCCCCGCCGATGACGCGGGAAGGACC
GATGCTGATCTTGCCTGCCTATATAAGGTTTTGCTATTCATTGAAAGCAGTAGTGACTGATTTGTATA
TA
>MinSyn_1871|Strength:0.000501502
GCGTGTCGTTTTAGTGAGGGCTTTGTCAAAAGCTAAAAAAGATGATGCTCACGACATCCTTACCGCTA
TGGGTAAGATTAGCTTAGCAAGACCTCTACAAATTTTCAACAATCAGCTTAGCAAGAATTTCGGGAAA
CCTCCTCGCTCACTATCAGCCCACTATGCCCAATTAGGTTCAGTGGTCCCTCCACTGCTAGGAGGTTA
TGACCCCCGCCGATGACGCGGGAAAACTGGTACTTGTGTACAGGGCTCAGTGGGAGCCACCATGATAA
AAATGTCAAAGATAGGACCGATGCTGATCCTATATAAGGTTTTGCTATTCATTGAAAGCAGTAGTGAC
TGATTTGTATATA
>MinSyn_1731|Strength:0.000502602
GCGTGTCGTTTTAGTGAGGCAGCCACTTGTGTGATGATCAGAAGATCAAAGGGCTATAGCAAGACTCA
CTATCAGCGTCACGACCACTATGTGGGAGCCACCATGCTCAGTGGTCCCTCCACTGTAGGATTGCGAT
AAAGGAAAGGGGCTATCAGCTTAGCAAGACCTCTACAAAATCCATCAACAAATAATCCAAGTAAGACA
GGGCTAAAAATGTCAAAGATAATCAGCTAACCACGTCTACAAGGAGGACCGATGCTGATCTTCTATAT
AAGGTTTTGCTATTCATTGAAAGCAGTAGTGACTGATTTGTATATA
>MinSyn_1986|Strength:0.000503427
GCGTGTCGTTTTAGTGAGGGCACACCAGCATGTGTTGATCACCAGCTGTACTTGTGTACAGGGCTCAA
GCAAGTGGATAGGTCACTCACTATCAGCGGACCGATGCTGATCTTGTCCATCAACAAATAATCCAAGT
AAGGTAGGTCACGACCACTATGCCCATTATGACCCCCGCCGATGACGCGGGAACGTGGGAGCCACCAC

CCAATTAGGTTGTCTGCACTATATAAGGTTTTGCTATTCATTGAAAGCAGTAGTGACTGATTTGTATA
TA
>MinSyn_1216|Strength:0.000504066
GCGTGTCGTTTTAGTGAGGTCCATCAACAAATAATCCAAGTAAGTGTACCAGTGGTCCCTCCACCCCA
ATTAGGTTGTCTGCACCTCACATGTAATCGAAAGGACAGTAAGGGCTCACTGCTAGGAGGACCGAGCA
AGTGGATTATGCCCAATTAGGTTGTCTAAGATTGATGAAAAGTCAAAAACAAAAATCAATTATTGCCC
AATTATGAAGCATCTTCCGCTCACTGCCTATATAAGGTTTTGCTATTCATTGAAAGCAGTAGTGACTG
ATTTGTATATA
>MinSyn_1550|Strength:0.000504213
GCGTGTCGTTTTAGTGAGGTGAAGCATCTTCCCAGCTTAGTGGTGGAGCACGACATACAGGGCTCACT
GCTAGGAGGACCGATGCTCAGAAGATCAAAGGGCTAGCAAGACCTCTACAAAACTGGTAGCTTTGTCA
AAAGCTAAAAAAGATGATGCGGCTCACTGGGAAAAAGAAGAGGTCACATGTAGGCTATCAGCAATTTC
GGGAAACCTCCTCGGGGCTCTATATAAGGTTTTGCTATTCATTGAAAGCAGTAGTGACTGATTTGTAT
ATA
>MinSyn_1850|Strength:0.000504548
GCGTGTCGTTTTAGTGAGGTGGTGGAGCACGACACTACAAAACTGGTACTTGAAGCATCTTCCAGGGC
TCACTGCTAGGAGGAGCACACCAGCATGTGTTGATCACCAGCTACTGCTAGGAGGTGGCTCCTACCCG
ATGCTGATCCTATATAAGGTTTTGCTATTCATTGAAAGCAGTAGTGACTGATTTGTATATA
>MinSyn_1883|Strength:0.000505066
GCGTGTCGTTTTAGTGAGGAAAAATGTCAAAGATACCACTATGCCCAATTGAAGATAAGATAATAATG
TTGAAGATAAGAGCTGATCTATTGCGATAAAGGAAAGGTACTTGTGTACAGGGCTCACTGAACCATTA
TTGCGCTTGATGATCTATATAAGGTTTTGCTATTCATTGAAAGCAGTAGTGACTGATTTGTATATA
>MinSyn_1639|Strength:0.000505957
GCGTGTCGTTTTAGTGAGGAGGTGGCTCCTACGTAGGTCACGACCACTAAGCAAGTGGATAGGCTATC
AGCTTAGCATGAAGCATCTTCCACTATTGCGATAAAGGAAAGGGTACAGGGCATCGAAAGGACAGTAT
CTTGCCCTATATAAGGTTTTGCTATTCATTGAAAGCAGTAGTGACTGATTTGTATATA
>MinSyn_1320|Strength:0.000506667
GCGTGTCGTTTTAGTGAGGTCCATCAACAAATAATCCAAGTAAGTCAGCTTAGCAAGACCTCTACAAA
ACTGGTAACCATTATTGCGTCTACAACTATATAAGGTTTTGCTATTCATTGAAAGCAGTAGTGACTGA
TTTGTATATA
>MinSyn_1156|Strength:0.000506897
GCGTGTCGTTTTAGTGAGGTCCATCAACAAATAATCCAAGTAAGGGACCGGCTTTGTCAAAAGCTAAA
AAAGATGATGCTAGGCTATCAGCTTAGCAAGACCTATATAAGGTTTTGCTATTCATTGAAAGCAGTAG
TGACTGATTTGTATATA
>MinSyn_1752|Strength:0.000507108
GCGTGTCGTTTTAGTGAGGCAAATATTTCTTGTGACCGATGCTGATCTTGCCTGCCTTCAGAAGATCA
AAGGGCTAAGGAAAAAGAAGAGGTGCTAGTGAAGATAAGATAATAATGTTGAAGATAAGATCAGTGGT
CCCTCCACAATTAGGTTGTCTGTCACTATCAGCCTGCTAGGAGGACCGATAGCAAGTGGATCCACTAT
GCCCAATTAGGTTGTCTGCACATCCTTACCGCTATGGGTAAGATTGTAGGCTAATTTCGGGAAACCTC
CTCGTTGTCTGCACCTCACATGTAGGCCTATATAAGGTTTTGCTATTCATTGAAAGCAGTAGTGACTG
ATTTGTATATA
>MinSyn_1214|Strength:0.000507693
GCGTGTCGTTTTAGTGAGGTCTCTCTGCCGACAGTGGTCCCAAACTTTTTCAACAACTGCACCTCACA
TGTAGGCTATCAGAACCACGTCTACAATAGGTCACGACCACTATGCCCAATTAGTTATGACCCCCGCC
GATGACGCGGGATGCTAGGAGGACCGATGCTGATTGCGATAAAGGAAAGGCTCACTGCTAGGAGGACC
GCACACCAGCATGTGTTGATCACCAGCTTAGGCTATCAGCTTTGAAGATAAGATAATAATGTTGAAGA
TAAGATCTACAAAACTGGATCCTTACCGCTATGGGTAAGATTCTGCACCTCACATGTAGGCTACTATA
TAAGGTTTTGCTATTCATTGAAAGCAGTAGTGACTGATTTGTATATA
>MinSyn_1671|Strength:0.000507923
GCGTGTCGTTTTAGTGAGGATCGAAAGGACAGTACCTTGATGTGGGAGCCACCAGTACTTGTGTACAG
GGCTCATTGCGATAAAGGAAAGGCATGGTGGAGCACGACAAGGAGGACCGATGCTGATCTTGCCTGCA
CACCAGCATGTGTTGATCACCAGCTACCAGTGGTCCCTCCACGGCTCACTGCTAGGAGGACCGATGCA
AATATTTCTTGTTGCTAGGAGGACCGATGAAGATTGATGAAAAGTCAAAAACAAAAATCAATTATTGC
TGATCTTGCCTCTCTCTGCCGACAGTGGTCCCAAAATGCTGATCTTGCCTGCCTTGATGACTATATAA

GGTTTTGCTATTCATTGAAAGCAGTAGTGACTGATTTGTATATA
>MinSyn_12|Strength:0.000508055
GCGTGTCGTTTTAGTGAGGTTTTCAACAAGCCCAATTAGGTTGTCTGCACCAACCATTATTGCGGTAC
AGGGCTCACTGCTAGGAGGACCGAGGTGGCTCCTACTGTAGGATCCTTACCGCTATGGGTAAGATTCT
CTACAAAAATGTCAAAGATATAGGTCACCTATATAAGGTTTTGCTATTCATTGAAAGCAGTAGTGACT
GATTTGTATATA
>MinSyn_1549|Strength:0.00051013
GCGTGTCGTTTTAGTGAGGCAGCCACTTGTGTGCATCCTTACCGCTATGGGTAAGATTCACCTCACAT
GTAGGCTATCAGATCGAAAGGACAGTAACTGGTACTTGTGTACAGGGCTCACAATTTCGGGAAACCTC
CTCGAAACCATTATTGCGCTTACTATATAAGGTTTTGCTATTCATTGAAAGCAGTAGTGACTGATTTG
TATATA
>MinSyn_1909|Strength:0.000511078
GCGTGTCGTTTTAGTGAGGATCCTTACCGCTATGGGTAAGATTCCACTATGCCCAATTAGGTTGTCTG
TCAGAAGATCAAAGGGCTAGCTCACTGCTAGGAGGACCGGCTTTGTCAAAAGCTAAAAAAGATGATGC
GACATCGAAAGGACAGTATACAGGGCTCAGCCACTTGTGTATGTAGGCTATCAGCTCTATATAAGGTT
TTGCTATTCATTGAAAGCAGTAGTGACTGATTTGTATATA
>MinSyn_1372|Strength:0.000511303
GCGTGTCGTTTTAGTGAGGAAGATTGATGAAAAGTCAAAAACAAAAATCAATTATTGCCCAATTAGGT
TGTCTGCACCTCACATTCACTATCAGCCACTATGCCCAATTAATTGCGATAAAGGAAAGGTGCACCTC
ACATGTAAAAAATGTCAAAGATATGCTGATAACCACGTCTACAACCTATATAAGGTTTTGCTATTCAT
TGAAAGCAGTAGTGACTGATTTGTATATA
>MinSyn_1958|Strength:0.000511538
GCGTGTCGTTTTAGTGAGGCAAATATTTCTTGTATATCAGAAGATCAAAGGGCTAGGTACTTGTGTAC
AGGGCTCTATATAAGGTTTTGCTATTCATTGAAAGCAGTAGTGACTGATTTGTATATA
>MinSyn_1510|Strength:0.000511558
GCGTGTCGTTTTAGTGAGGTCAGAAGATCAAAGGGCTAACAGGGCTCACTGCTAGGAAAAAGAAGAGG
TGTTGTCTGCACCTCACATGTAGGCTATCCAAATATTTCTTGTCAATTCCATCAACAAATAATCCAAG
TAAGATGCTGCTATATAAGGTTTTGCTATTCATTGAAAGCAGTAGTGACTGATTTGTATATA
>MinSyn_1438|Strength:0.000511664
GCGTGTCGTTTTAGTGAGGTGAAGATAAGATAATAATGTTGAAGATAAGACTATGCCCAATTAGGTTG
TCAAAAATGTCAAAGATAGCTCACTATCAGCTACTTGTGTACAGGGCTCACTGCTAGGAAGATTGATG
AAAAGTCAAAAACAAAAATCAATTATTCTGCACCTCACATGTAGGCTAAGGTGGCTCCTACTAGGTTG
TCTGCACCTATATAAGGTTTTGCTATTCATTGAAAGCAGTAGTGACTGATTTGTATATA
>MinSyn_1429|Strength:0.000513677
GCGTGTCGTTTTAGTGAGGAATTTCGGGAAACCTCCTCGTAGGAGGACCGATGCTGATCTTGCATCCT
TACCGCTATGGGTAAGATTTCTTAGGTGGCTCCTACAGGCTATCAGCTTAGCAAGACCATCGAAAGGA
CAGTATCACGCTTTGTCAAAAGCTAAAAAAGATGATGCTACAAAACTGGTACTTGTGTACTTATGACC
CCCGCCGATGACGCGGGAGCCTGCCTTGATGCTATATAAGGTTTTGCTATTCATTGAAAGCAGTAGTG
ACTGATTTGTATATA
>MinSyn_1566|Strength:0.000513997
GCGTGTCGTTTTAGTGAGGATCCTTACCGCTATGGGTAAGATTTGTAGGCTATCAGCTTAGCAAGACC
TCTATCAGAAGATCAAAGGGCTATGCTAGGAGGACCGATGCTGATCTTGCCTAATTTCGGGAAACCTC
CTCGTGAGCAAGTGGATAGGTCACGACCACTATGGTGGAGCACGACAGATCTCTCTGCCGACAGTGGT
CCCAAAGGTTGTCTGCACCTCACCTATATAAGGTTTTGCTATTCATTGAAAGCAGTAGTGACTGATTT
GTATATA
>MinSyn_120|Strength:0.00051431
GCGTGTCGTTTTAGTGAGGGCTTTGTCAAAAGCTAAAAAAGATGATGCCACAAGATTGATGAAAAGTC
AAAAACAAAAATCAATTATTTGTGTACAGGGCTGTGGGAGCCACCAGGTAGGTCACGAAGGTGGCTCC
TACGGACCGATGCTGATCTTGCCTTCAGAAGATCAAAGGGCTATAGGAGGACCGATGCTGATCTTGCC
TGGCACACCAGCATGTGTTGATCACCAGCTGCAACTATATAAGGTTTTGCTATTCATTGAAAGCAGTA
GTGACTGATTTGTATATA
>MinSyn_1452|Strength:0.000515092
GCGTGTCGTTTTAGTGAGGGTGGGAGCCACCACACTGCTAGGAGGACCGTCAGAAGATCAAAGGGCTA
TCACATGTAGGAACCACGTCTACAAGGTAGGTCACGACCACTATCCTTACCGCTATGGGTAAGATTGT

AGGTCCTATATAAGGTTTTGCTATTCATTGAAAGCAGTAGTGACTGATTTGTATATA
>MinSyn_1583|Strength:0.000515698
GCGTGTCGTTTTAGTGAGGTGAAGCATCTTCCGATTGCGATAAAGGAAAGGAATTAGGTTGTCTGCAC
CTTCCATCAACAAATAATCCAAGTAAGTACAGGGCTCACTGCTAGGATCCTTACCGCTATGGGTAAGA
TTCAAAACGCACACCAGCATGTGTTGATCACCAGCTGTTGTCTGCACCTCACTTATGACCCCCGCCGA
TGACGCGGGACCGATGCTGATCTTGCCTGCCTTGCTATATAAGGTTTTGCTATTCATTGAAAGCAGTA
GTGACTGATTTGTATATA
>MinSyn_1501|Strength:0.000516784
GCGTGTCGTTTTAGTGAGGTGAAGATAAGATAATAATGTTGAAGATAAGAGTAGGTCACGACCAATTG
CGATAAAGGAAAGGTGTCTGTTTTCAACAATATGAAGCATCTTCCCACGACCACTATGCCCAATTAGG
TTGTCTTCAGAAGATCAAAGGGCTAGCTTAGCAAGACCTCTACAATCGAAAGGACAGTAACGACCACC
AGCCACTTGTGTACCTCTACAAAAGCAAGTGGATCTCTACAGGTGGCTCCTACTGATCTTCTATATAA
GGTTTTGCTATTCATTGAAAGCAGTAGTGACTGATTTGTATATA
>MinSyn_1107|Strength:0.000517843
GCGTGTCGTTTTAGTGAGGTCCATCAACAAATAATCCAAGTAAGCGATGCTGATCTTGCCTGCCTTGA
TGCACACCAGCATGTGTTGATCACCAGCTGCTGATCTTGCCTGCCTTGTTTTCAACAACTTGCCTTAT
GACCCCCGCCGATGACGCGGGACGCGGTAGGTCACCTATATAAGGTTTTGCTATTCATTGAAAGCAGT
AGTGACTGATTTGTATATA
>MinSyn_1147|Strength:0.000518182
GCGTGTCGTTTTAGTGAGGATTGCGATAAAGGAAAGGTGCTGATCTTGCCTAAAAATGTCAAAGATAA
AAACTGGTCTATATAAGGTTTTGCTATTCATTGAAAGCAGTAGTGACTGATTTGTATATA
>MinSyn_1846|Strength:0.000518202
GCGTGTCGTTTTAGTGAGGATCGAAAGGACAGTAGTTGTCCATCAACAAATAATCCAAGTAAGAACTG
GTACTTTTTTCAACAAAGGCGCTTTGTCAAAAGCTAAAAAAGATGATGCCAAAACTGGAAGATTGATG
AAAAGTCAAAAACAAAAATCAATTATTCACTGCTAGGAGGACCGATGCTTTATGACCCCCGCCGATGA
CGCGGGAAGACCTCTCACTATCAGCGTAGGCTATCATCCTTACCGCTATGGGTAAGATTTCACGACCA
CTATGCCCAATTAGGTAACCACGTCTACAATGTGTACAGGGCTCACTGCTATATAAGGTTTTGCTATT
CATTGAAAGCAGTAGTGACTGATTTGTATATA
>MinSyn_1514|Strength:0.000518555
GCGTGTCGTTTTAGTGAGGATTGCGATAAAGGAAAGGCCCAATTAGGTTGTCTGCACCTCACGGAAAA
AGAAGAGGTATGCTGATCTTGCCGCTTTGTCAAAAGCTAAAAAAGATGATGCTACAGGGCAGTGGTCC
CTCCACGATGCTGATCTTGCCTGCCTTCACTATCAGCTACTTGTGTACAGGGCTCACTGCTATTATGA
CCCCCGCCGATGACGCGGGAGTGTACATGAAGATAAGATAATAATGTTGAAGATAAGACAATTAGGTT
GTCTCTATATAAGGTTTTGCTATTCATTGAAAGCAGTAGTGACTGATTTGTATATA
>MinSyn_1981|Strength:0.000518869
GCGTGTCGTTTTAGTGAGGGCACACCAGCATGTGTTGATCACCAGCTCCCAATTAGGTTGTGTGGGAG
CCACCAGTAGGCTATCAGCTTAGCATTTTCAACAAAGGTCACGAGCTTTGTCAAAAGCTAAAAAAGAT
GATGCCTTGTGTACAGGGCTCACTGCTAGTGGTGGAGCACGACACCACTATGCCCAATTAGGTTGTCT
CAGCCACTTGTGTTATGCCCAATTAGGTTGTCTGCACCTCACATCCTTACCGCTATGGGTAAGATTAG
CTTAGCAAGACCTCTACAAAACTGGTAAGATTGATGAAAAGTCAAAAACAAAAATCAATTATCACTGC
TAGGAGGACCTTATGACCCCCGCCGATGACGCGGGACTCCTATATAAGGTTTTGCTATTCATTGAAAG
CAGTAGTGACTGATTTGTATATA
>MinSyn_1284|Strength:0.000519069
GCGTGTCGTTTTAGTGAGGTTTTCAACAAAGGCTATAACCATTATTGCGGTCACGACCGGAAAAAGAA
GAGGTAAACTGGTACTTGTGTACAGGGCAAGATTGATGAAAAGTCAAAAACAAAAATCAATTATTGCT
AGGAGGACCGATGCTGATCTTGCCTGAAGATAAGATAATAATGTTGAAGATAAGATAGGTCTATATAA
GGTTTTGCTATTCATTGAAAGCAGTAGTGACTGATTTGTATATA
>MinSyn_1993|Strength:0.000519938
GCGTGTCGTTTTAGTGAGGTCAGAAGATCAAAGGGCTACAGCTTAGCAAGGCACACCAGCATGTGTTG
ATCACCAGCTGTAGGCTATCAGCTTAGCAAGACTGAAGCATCTTCCGGTAGGTCACGACTTATGACCC
CCGCCGATGACGCGGGAGTAGAGGTGGCTCCTACGCACCTCACATGTAGGCCTATATAAGGTTTTGCT
ATTCATTGAAAGCAGTAGTGACTGATTTGTATATA
>MinSyn_1606|Strength:0.000520184
GCGTGTCGTTTTAGTGAGGAAGATTGATGAAAAGTCAAAAACAAAAATCAATTATGTACAGGGCTGCT

TTGTCAAAAGCTAAAAAAGATGATGCGGTACTTGTGTACAGGAGGTGGCTCCTACCACTGCTAGGAGG
ACCGCACACCAGCATGTGTTGATCACCAGCTATCAGCTTAGCAAGACCCTATATAAGGTTTTGCTATT
CATTGAAAGCAGTAGTGACTGATTTGTATATA
>MinSyn_126|Strength:0.000521004
GCGTGTCGTTTTAGTGAGGGCTTTGTCAAAAGCTAAAAAAGATGATGCGGTACTTGTGTACAGGGCTC
ACTGTGGGAGCCACCAAGCTTATCCTTACCGCTATGGGTAAGATTTCACGACCACTATGCCCTATATA
AGGTTTTGCTATTCATTGAAAGCAGTAGTGACTGATTTGTATATA
>MinSyn_124|Strength:0.000521194
GCGTGTCGTTTTAGTGAGGGCTTTGTCAAAAGCTAAAAAAGATGATGCTGTAGGCTATCAGCTTAGCA
AGACTGAAGCATCTTCCGTCACGAATTTCGGGAAACCTCCTCGGGTTGTCTGCACCTCACATGTAGTC
AGAAGATCAAAGGGCTACTGGCTATATAAGGTTTTGCTATTCATTGAAAGCAGTAGTGACTGATTTGT
ATATA
>MinSyn_1781|Strength:0.000521266
GCGTGTCGTTTTAGTGAGGTCCATCAACAAATAATCCAAGTAAGAGCAATCGAAAGGACAGTATGCCC
AATTAGGTTGTCTGCACCTCACAGCTTTGTCAAAAGCTAAAAAAGATGATGCGCTATATAAGGTTTTG
CTATTCATTGAAAGCAGTAGTGACTGATTTGTATATA
>MinSyn_1152|Strength:0.0005222
GCGTGTCGTTTTAGTGAGGATCGAAAGGACAGTATTGTGTACAGGGCTCACTGCTAGGAGGGTGGGAG
CCACCACTGCACCTCACATGTAGGCTATCAGCTTAGATTGCGATAAAGGAAAGGCTAGCTTTGTCAAA
AGCTAAAAAAGATGATGCTGTCTGCTATATAAGGTTTTGCTATTCATTGAAAGCAGTAGTGACTGATT
TGTATATA
>MinSyn_175|Strength:0.000522821
GCGTGTCGTTTTAGTGAGGGTGGGAGCCACCAAGGTCAAAAATGTCAAAGATAAAGACCTCTCACTAT
CAGCCAATCCATCAACAAATAATCCAAGTAAGTTAGGTTGTCTTGAAGATAAGATAATAATGTTGAAG
ATAAGATCTACAAAACTGGTACTTGTGGCTTTGTCAAAAGCTAAAAAAGATGATGCCACATGTAGGCT
ATCAGCTTAGCCTATATAAGGTTTTGCTATTCATTGAAAGCAGTAGTGACTGATTTGTATATA
>MinSyn_1597|Strength:0.000524138
GCGTGTCGTTTTAGTGAGGTTTTCAACAAGGCTAATTTCGGGAAACCTCCTCGCACGACCCTATATAA
GGTTTTGCTATTCATTGAAAGCAGTAGTGACTGATTTGTATATA
>MinSyn_1125|Strength:0.000524583
GCGTGTCGTTTTAGTGAGGCAAATATTTCTTGTGCTATGTGGGAGCCACCACCCAATTAGGTTGTTAT
GACCCCCGCCGATGACGCGGGATTAGCAAGAAATTTCGGGAAACCTCCTCGTTAGCAGCCACTTGTGT
CACATGTAGGCTATCATCCATCAACAAATAATCCAAGTAAGGCCCAATTAGGTTAACCATTATTGCGG
ATCTTGCCTGCCTTGATGACTATATAAGGTTTTGCTATTCATTGAAAGCAGTAGTGACTGATTTGTAT
ATA
>MinSyn_1542|Strength:0.000525664
GCGTGTCGTTTTAGTGAGGAATTTCGGGAAACCTCCTCGTGCACCTCACATGTAGGCTATCAGCTTAT
GGTGGAGCACGACACACTATGCCCAATTAGGTGTGGGAGCCACCAATCAGCAAAAATGTCAAAGATAG
ACCTCTACAAAACTGGTACTTAACCATTATTGCGTGCACCTCGCACACCAGCATGTGTTGATCACCAG
CTCCTATATAAGGTTTTGCTATTCATTGAAAGCAGTAGTGACTGATTTGTATATA
>MinSyn_1205|Strength:0.000526351
GCGTGTCGTTTTAGTGAGGTTATGACCCCCGCCGATGACGCGGGAACTGGTACTTGTGTACAGGGCTC
ACTTGAAGCATCTTCCAGACCTCTCTATATAAGGTTTTGCTATTCATTGAAAGCAGTAGTGACTGATT
TGTATATA
>MinSyn_1629|Strength:0.000526517
GCGTGTCGTTTTAGTGAGGCAGCCACTTGTGTAGCAAGACCTCTACAAAACTGGTACTTGTAATTTCG
GGAAACCTCCTCGGCTTAGCAAGACCTCTACAAAACTGGTACAACCATTATTGCGCACATGTAGGCTA
TCAGCATTGCGATAAAGGAAAGGAGGTCACGAAGATTGATGAAAAGTCAAAAACAAAAATCAATTATC
ACTGCTAGGAGGACCGATGAAGATAAGATAATAATGTTGAAGATAAGAATGCTGATCATCCTTACCGC
TATGGGTAAGATTACCGATGCTGATGAAGCATCTTCCTCTACAAAACTGGTACTTGTGTTTTTCAACA
AAATTACTATATAAGGTTTTGCTATTCATTGAAAGCAGTAGTGACTGATTTGTATATA
>MinSyn_1475|Strength:0.000527332
GCGTGTCGTTTTAGTGAGGATCGAAAGGACAGTAGATCTTGCCTGCCTTGATGAAAGATTGATGAAAA
GTCAAAAACAAAAATCAATTATATGCTGAGTGGGAGCCACCATTGTGTACAGGGCTCACTGCTAGGAG

GACTGAAGATAAGATAATAATGTTGAAGATAAGATGCCTTGATGATTTTTCAACAAACTGCTAGGAGG
ACCGATAATTTCGGGAAACCTCCTCGACTATCAGCCACTTGTGTACCTCACATTGCGATAAAGGAAAG
GCCTCTACAAAACTAGCAAGTGGATAATTAGGTTGTCTATATAAGGTTTTGCTATTCATTGAAAGCAG
TAGTGACTGATTTGTATATA
>MinSyn_1915|Strength:0.000527422
GCGTGTCGTTTTAGTGAGGAACCACGTCTACAAGCTGATCTTGCCTGCCTTGATGATACAGCCACTTG
TGTCTAGCAAGTGGATTCACATGTAGGCTATCAGCTTAGCATCCTTACCGCTATGGGTAAGATTTGCA
CCTCACATGTGCTTTGTCAAAAGCTAAAAAAGATGATGCTGTAGGCTATCAGCTTAGCCAGTGGTCCC
TCCACTGCTGTGAAGATAAGATAATAATGTTGAAGATAAGACGCGGTAGGTCACGACCTATATAAGGT
TTTGCTATTCATTGAAAGCAGTAGTGACTGATTTGTATATA
>MinSyn_1926|Strength:0.000528194
GCGTGTCGTTTTAGTGAGGGCACACCAGCATGTGTTGATCACCAGCTCTGCACCTCACATGCAGTGGT
CCCTCCACCTGCCTAATTTCGGGAAACCTCCTCGAGGTCACGACCACTATGCCTATATAAGGTTTTGC
TATTCATTGAAAGCAGTAGTGACTGATTTGTATATA
>MinSyn_1138|Strength:0.000528436
GCGTGTCGTTTTAGTGAGGTCAGAAGATCAAAGGGCTACTCACTGCTAGGAGGACCGATGCTGTGGTG
GAGCACGACAGGAAAAATGTCAAAGATACCTCTACATCTCTCTGCCGACAGTGGTCCCAAAAATTAAG
GTGGCTCCTACCAAGACCTCTACAAAGATTGATGAAAAGTCAAAAACAAAAATCAATTATATGTAGGC
TATCAGCTTAGCAAGACCTCTCTATATAAGGTTTTGCTATTCATTGAAAGCAGTAGTGACTGATTTGT
ATATA
>MinSyn_127|Strength:0.000530142
GCGTGTCGTTTTAGTGAGGTTATGACCCCCGCCGATGACGCGGGAACTGGTACTTGTGTACATCAGAA
GATCAAAGGGCTAGTAGTCACTATCAGCTTGATGACTATATAAGGTTTTGCTATTCATTGAAAGCAGT
AGTGACTGATTTGTATATA
>MinSyn_1979|Strength:0.000531128
GCGTGTCGTTTTAGTGAGGGGAAAAAGAAGAGGTGATGCTGATTGGTGGAGCACGACAAACTGGTAAA
TTTCGGGAAACCTCCTCGTGCTGATCTTGCAGTGGTCCCTCCACGATCGAAAGGACAGTAAGGACCGA
TGCTGATCTTATGACCCCCGCCGATGACGCGGGATCTGCACCTCACATGTAGGCTGAAGCATCTTCCA
CTGCTAACCATTATTGCGGGCTATCACTATATAAGGTTTTGCTATTCATTGAAAGCAGTAGTGACTGA
TTTGTATATA
>MinSyn_1215|Strength:0.000531175
GCGTGTCGTTTTAGTGAGGTGGTGGAGCACGACAACCTCACATGTAGGCTAAACCACGTCTACAAACT
GGTACTTGTGTACAGGGCTAAAAATGTCAAAGATAGCTTAGCAGCTTTGTCAAAAGCTAAAAAAGATG
ATGCACTGCTAGGAGGTCCATCAACAAATAATCCAAGTAAGCTACAAAACTGGTATCACTATCAGCCA
AAACTGCTATATAAGGTTTTGCTATTCATTGAAAGCAGTAGTGACTGATTTGTATATA
>MinSyn_191|Strength:0.000531882
GCGTGTCGTTTTAGTGAGGATTGCGATAAAGGAAAGGGCCTGCCTTGTTATGACCCCCGCCGATGACG
CGGGAGGTACTTGTGTACAGGATCGAAAGGACAGTAGATGCTGTCACTATCAGCTGTGAAGATAAGAT
AATAATGTTGAAGATAAGATATCAGCTTAGCAAGACCTCTACCTATATAAGGTTTTGCTATTCATTGA
AAGCAGTAGTGACTGATTTGTATATA
>MinSyn_1131|Strength:0.000532348
GCGTGTCGTTTTAGTGAGGGGAAAAAGAAGAGGTACCACTATGCCCAATTAGGTTGTTATGACCCCCG
CCGATGACGCGGGATGCCTTGATGATTGCGATAAAGGAAAGGAACTGGTACTTGTGTCTATATAAGGT
TTTGCTATTCATTGAAAGCAGTAGTGACTGATTTGTATATA
>MinSyn_1296|Strength:0.000532991
GCGTGTCGTTTTAGTGAGGAACCACGTCTACAACTATGCCCAATTAGGTTGTCTGTTATGACCCCCGC
CGATGACGCGGGAGCGGTAGGTCAGGTGGCTCCTACTACAGGGCTCACTGCTGCTTTGTCAAAAGCTA
AAAAAGATGATGCCACGACCACTAAGATTGATGAAAAGTCAAAAACAAAAATCAATTATACTTGTGTA
CAGGGCTCACTGCTACTATATAAGGTTTTGCTATTCATTGAAAGCAGTAGTGACTGATTTGTATATA
>MinSyn_1318|Strength:0.000533672
GCGTGTCGTTTTAGTGAGGAGCAAGTGGATAGGACCGATGCTGATCTTGCCTGCCTGGAAAAAGAAGA
GGTTAGGCTATCAGCTTAGCAAGAGGTGGCTCCTACCTCACATGTAGGCTATCAGCTTAGCAAGCAGT
GGTCCCTCCACGTGAAGCATCTTCCTGGTATCAGAAGATCAAAGGGCTACTCACTGCTAGGAGGACCG
ATGCTGATCTTAAGATTGATGAAAAGTCAAAAACAAAAATCAATTATCCTCTACAAAACTAAAAATGT

CAAAGATAGCCCAATTAGGTTGTTCTCTCTGCCGACAGTGGTCCCAAATATCACTATATAAGGTTTTG
CTATTCATTGAAAGCAGTAGTGACTGATTTGTATATA
>MinSyn_1283|Strength:0.000534427
GCGTGTCGTTTTAGTGAGGTCACTATCAGCTACTTGTGTACAGGGCTCACTGCTAGAACCACGTCTAC
AAAGGTTTGAAGATAAGATAATAATGTTGAAGATAAGACACGGAAAAAGAAGAGGTCAAAACTGGTAC
TTGTGTACTTTTCAACAAACCGATGCTGAATTTCGGGAAACCTCCTCGGCCTGCCTTGAAGCATCTTC
CTAGGCTATCTATATAAGGTTTTGCTATTCATTGAAAGCAGTAGTGACTGATTTGTATATA
>MinSyn_1591|Strength:0.000536024
GCGTGTCGTTTTAGTGAGGAAAAATGTCAAAGATATCGCGGTAATCCTTACCGCTATGGGTAAGATTT
TTGAAGCATCTTCCATGCTGATCTTGCCTGCCTTGACTATATAAGGTTTTGCTATTCATTGAAAGCAG
TAGTGACTGATTTGTATATA
>MinSyn_1376|Strength:0.00053629
GCGTGTCGTTTTAGTGAGGCAAATATTTCTTGTGTAGGCTCAGAAGATCAAAGGGCTACCTCTACAAA
CTATATAAGGTTTTGCTATTCATTGAAAGCAGTAGTGACTGATTTGTATATA
>MinSyn_148|Strength:0.000536421
GCGTGTCGTTTTAGTGAGGAACCACGTCTACAATATGCCCAATTAGGTTGTCTGCACCATTGCGATAA
AGGAAAGGTGATCTTGCCTGCACACCAGCATGTGTTGATCACCAGCTGCTAGGTTATGACCCCCGCCG
ATGACGCGGGACTCACTGCTAGATCCTTACCGCTATGGGTAAGATTGTCTGCACCTCACATGTAGGCT
ATCAGCTTTCCATCAACAAATAATCCAAGTAAGCCAATTAGGTGGTGGAGCACGACACGACCACTATG
CCCAATTCAGTGGTCCCTCCACTAGGCTATATAAGGTTTTGCTATTCATTGAAAGCAGTAGTGACTGA
TTTGTATATA
>MinSyn_1493|Strength:0.000536749
GCGTGTCGTTTTAGTGAGGGCACACCAGCATGTGTTGATCACCAGCTGGGCTCACTTTATGACCCCCG
CCGATGACGCGGGACTCTACAAAACTGGTACTTGTGTACAGGGTGAAGCATCTTCCCGGAAAAAGAAG
AGGTTGCTGATCACTATCAGCGGCTCACTGCTAGCTATATAAGGTTTTGCTATTCATTGAAAGCAGTA
GTGACTGATTTGTATATA
>MinSyn_1530|Strength:0.000537459
GCGTGTCGTTTTAGTGAGGAGCAAGTGGATTCACATGTAGGCTATCAGCTAATTTCGGGAAACCTCCT
CGACGACCACTATGCCCAAGATTGATGAAAAGTCAAAAACAAAAATCAATTATACCGATGGAAAAAGA
AGAGGTGGTACTCAGCCACTTGTGTGGTTGTCTGCACCTCACCTATATAAGGTTTTGCTATTCATTGA
AAGCAGTAGTGACTGATTTGTATATA
>MinSyn_1622|Strength:0.000537689
GCGTGTCGTTTTAGTGAGGATTGCGATAAAGGAAAGGTGGTACTTGTTCTCTCTGCCGACAGTGGTCC
CAAAGCCCAATTGAAGCATCTTCCTCACTGCTAGGAGGACCGATGCAAGATTGATGAAAAGTCAAAAA
CAAAAATCAATTATAATTAGGTTGTCCTATATAAGGTTTTGCTATTCATTGAAAGCAGTAGTGACTGA
TTTGTATATA
>MinSyn_1864|Strength:0.000538583
GCGTGTCGTTTTAGTGAGGTCAGAAGATCAAAGGGCTAGTACTTGTGTACAGGCAGTGGTCCCTCCAC
GCTCTATATAAGGTTTTGCTATTCATTGAAAGCAGTAGTGACTGATTTGTATATA
>MinSyn_1270|Strength:0.000538591
GCGTGTCGTTTTAGTGAGGGCACACCAGCATGTGTTGATCACCAGCTTACAGAACCATTATTGCGGGC
TCACTGCTAGGGGAAAAAGAAGAGGTTATCAGCTATATAAGGTTTTGCTATTCATTGAAAGCAGTAGT
GACTGATTTGTATATA
>MinSyn_1696|Strength:0.00053894
GCGTGTCGTTTTAGTGAGGATCCTTACCGCTATGGGTAAGATTACCGATGCTGATCTTGCCTGCCTTG
ATGATATTGCGATAAAGGAAAGGCGACCACTATGCCCTTATGACCCCCGCCGATGACGCGGGATAAGA
TTGATGAAAAGTCAAAAACAAAAATCAATTATAGACCTCTACAAAACTGGTACTTGTCTATATAAGGT
TTTGCTATTCATTGAAAGCAGTAGTGACTGATTTGTATATA
>MinSyn_1181|Strength:0.000539645
GCGTGTCGTTTTAGTGAGGAGGTGGCTCCTACATGCACACCAGCATGTGTTGATCACCAGCTTAGGAG
GACCGATGCTGATCTTGCTTATGACCCCCGCCGATGACGCGGGACACATGTAGAACCACGTCTACAAA
ACTGGTACTTGGAAAAAGAAGAGGTAGGAGGACCGATGCCAAATATTTCTTGTGGCTATATAAGGTTT
TGCTATTCATTGAAAGCAGTAGTGACTGATTTGTATATA
>MinSyn_1657|Strength:0.000540517

GCGTGTCGTTTTAGTGAGGTCACTATCAGCGGTACTAATTTCGGGAAACCTCCTCGACCACTATATAA
GGTTTTGCTATTCATTGAAAGCAGTAGTGACTGATTTGTATATA
>MinSyn_1354|Strength:0.00054134
GCGTGTCGTTTTAGTGAGGCAGTGGTCCCTCCACTGATCTTCAGAAGATCAAAGGGCTACAATTAGGT
TGTCTGCTTATGACCCCCGCCGATGACGCGGGATCACATGTAGGCTATTGAAGCATCTTCCCCTTGAT
GATTCTCTCTGCCGACAGTGGTCCCAAAGGTAGCAAGTGGATCAGGGCTCACTGCTAGGAAGATTGAT
GAAAAGTCAAAAACAAAAATCAATTATAACTGGTACTTGTGTACAGGGCCTATATAAGGTTTTGCTAT
TCATTGAAAGCAGTAGTGACTGATTTGTATATA
>MinSyn_1534|Strength:0.000541439
GCGTGTCGTTTTAGTGAGGCAAATATTTCTTGTCCGATGCTGATCTTGAAGATTGATGAAAAGTCAAA
AACAAAAATCAATTATGATCTTGCCTGCCTTTTTCAACAAACAGGGCTCACTGCTAGCTTTGTCAAAA
GCTAAAAAAGATGATGCAGATTGCGATAAAGGAAAGGGCTTAGCAAGACCTCTACAAATCTCTCTGCC
GACAGTGGTCCCAAAAGCAAGACCTCTACAAAACTGGTACTTGTGCACACCAGCATGTGTTGATCACC
AGCTAGGTCACAGGTGGCTCCTACAGGGCTCACTGCTAGGAAACCATTATTGCGTCCTATATAAGGTT
TTGCTATTCATTGAAAGCAGTAGTGACTGATTTGTATATA
>MinSyn_1930|Strength:0.000541642
GCGTGTCGTTTTAGTGAGGATTGCGATAAAGGAAAGGTATCATCAGAAGATCAAAGGGCTAGATCTTG
CCTGCCTTGATAGCAAGTGGATTCTGCACCTCACATGTAGGCTATCAGCTTTGTCAAAAGCTAAAAAA
GATGATGCCTTAGCAAGACCTCTACAAAACTGAATTTCGGGAAACCTCCTCGGACCGATGCTGATTTA
TGACCCCCGCCGATGACGCGGGACCTATATAAGGTTTTGCTATTCATTGAAAGCAGTAGTGACTGATT
TGTATATA
>MinSyn_1217|Strength:0.000541698
GCGTGTCGTTTTAGTGAGGTCTCTCTGCCGACAGTGGTCCCAAACGGTAGGTCACCAGTGGTCCCTCC
ACCTTAGCAAGACCTCTATTATGACCCCCGCCGATGACGCGGGACTATTGCGATAAAGGAAAGGCTTG
TGTACAGGGCTCACTGCTAGGACTATATAAGGTTTTGCTATTCATTGAAAGCAGTAGTGACTGATTTG
TATATA
>MinSyn_1496|Strength:0.000542839
GCGTGTCGTTTTAGTGAGGATCGAAAGGACAGTACAATTAGGTTGTCTGCACCAAGATTGATGAAAAG
TCAAAAACAAAAATCAATTATCTGCCTCTATATAAGGTTTTGCTATTCATTGAAAGCAGTAGTGACTG
ATTTGTATATA
>MinSyn_1174|Strength:0.000543276
GCGTGTCGTTTTAGTGAGGAACCATTATTGCGAAATCCTTACCGCTATGGGTAAGATTTATCAGCTTA
GCAAGACCTCTACAAATGAAGATAAGATAATAATGTTGAAGATAAGAAAGCAAGTGGATGGACCGATA
AGATTGATGAAAAGTCAAAAACAAAAATCAATTATGCTCACTGCTAGGAGGACCGATGCTGATCTAGG
TGGCTCCTACAGGCTATCAGCTTAGCAAGACCTCCATCAACAAATAATCCAAGTAAGCACTTCAGAAG
ATCAAAGGGCTACCCAATTAGGTTGTCTCTATATAAGGTTTTGCTATTCATTGAAAGCAGTAGTGACT
GATTTGTATATA
>MinSyn_1179|Strength:0.000543854
GCGTGTCGTTTTAGTGAGGGCTTTGTCAAAAGCTAAAAAAGATGATGCGTAGGTCACGACCACTATGC
CCAAGATTGATGAAAAGTCAAAAACAAAAATCAATTATTAGGCTATCAGCTTAGCAAGACCTCTCCAT
CAACAAATAATCCAAGTAAGCCTATATAAGGTTTTGCTATTCATTGAAAGCAGTAGTGACTGATTTGT
ATATA
>MinSyn_1567|Strength:0.000544288
GCGTGTCGTTTTAGTGAGGTCCATCAACAAATAATCCAAGTAAGGTTGTCTGCTGAAGATAAGATAAT
AATGTTGAAGATAAGATAGCAAGACCTCTACAAAACTGGTACTTGTATTGCGATAAAGGAAAGGTAAA
AATGTCAAAGATAATTAGGTTGTCTGCACCTCACATCGAAAGGACAGTATTGGTGGAGCACGACAGAG
GACCCTATATAAGGTTTTGCTATTCATTGAAAGCAGTAGTGACTGATTTGTATATA
>MinSyn_134|Strength:0.000544462
GCGTGTCGTTTTAGTGAGGTCAGAAGATCAAAGGGCTATTGTTCACTATCAGCACTGCTAGGAGGAAT
TGCGATAAAGGAAAGGTTGGGAAAAAGAAGAGGTAGGCTATCAGCTTAGAGCAAGTGGATTCTGCACC
AATTTCGGGAAACCTCCTCGTAGGCTCTATATAAGGTTTTGCTATTCATTGAAAGCAGTAGTGACTGA
TTTGTATATA
>MinSyn_164|Strength:0.000544667
GCGTGTCGTTTTAGTGAGGTTATGACCCCCGCCGATGACGCGGGAGGTAGGTCACGGAAAAAGAAGAG

GTCTTAGCAAGACCTCTACAAAACTGCTATATAAGGTTTTGCTATTCATTGAAAGCAGTAGTGACTGA
TTTGTATATA
>MinSyn_1370|Strength:0.000545512
GCGTGTCGTTTTAGTGAGGAAAAATGTCAAAGATAGCTCAGAAGATCAAAGGGCTACTTGCCTGCCTT
GATCACTATCAGCCTCTACAAAACTGGCAAATATTTCTTGTTGATCTTGCCTGCGGAAAAAGAAGAGG
TCGCTTTGTCAAAAGCTAAAAAAGATGATGCTCAGCTTAGCAAGACCTCAGCCACTTGTGTTTAGGTT
GTCTGCTATATAAGGTTTTGCTATTCATTGAAAGCAGTAGTGACTGATTTGTATATA
>MinSyn_1937|Strength:0.00054576
GCGTGTCGTTTTAGTGAGGTCTCTCTGCCGACAGTGGTCCCAAATACAAAACTGGTACTTGTGTTGAA
GATAAGATAATAATGTTGAAGATAAGATGCCTATATAAGGTTTTGCTATTCATTGAAAGCAGTAGTGA
CTGATTTGTATATA
>MinSyn_1491|Strength:0.000545999
GCGTGTCGTTTTAGTGAGGCAGCCACTTGTGTATGATCGAAAGGACAGTAACAAGATTGATGAAAAGT
CAAAAACAAAAATCAATTATCAGCTTAGCAAGACCTCTACAAAACTTTATGACCCCCGCCGATGACGC
GGGACATTTTCAACAACCTCTACAAAACTGGTACTTGTGTACTGAAGATAAGATAATAATGTTGAAGA
TAAGAGACCGATGCCTATATAAGGTTTTGCTATTCATTGAAAGCAGTAGTGACTGATTTGTATATA
>MinSyn_1892|Strength:0.000546105
GCGTGTCGTTTTAGTGAGGGGAAAAAGAAGAGGTAGGCTATCAGCTTAGCAAGACCTCTACAAAAAGA
TTGATGAAAAGTCAAAAACAAAAATCAATTATATCAGCTTAGCAAGCAGTGGTCCCTCCACTCACATG
TAGGCTATCAGGCTTTGTCAAAAGCTAAAAAAGATGATGCCGCGTGAAGCATCTTCCGACATTGCGAT
AAAGGAAAGGGGTTGTCTGCACCTCACATGCTATATAAGGTTTTGCTATTCATTGAAAGCAGTAGTGA
CTGATTTGTATATA
>MinSyn_1665|Strength:0.000547049
GCGTGTCGTTTTAGTGAGGCAAATATTTCTTGTCTCTACAAAACTGGTACTTGAAGATTGATGAAAAG
TCAAAAACAAAAATCAATTATCTCTACAAAACTGTGAAGCATCTTCCTAGGAGGACCGATGCTGATCC
TATATAAGGTTTTGCTATTCATTGAAAGCAGTAGTGACTGATTTGTATATA
>MinSyn_1266|Strength:0.000547899
GCGTGTCGTTTTAGTGAGGAACCACGTCTACAACAAGACCTCTGCTTTGTCAAAAGCTAAAAAAGATG
ATGCACAAAACTTTTTCAACAATGCTAGGAGGACCGCTATATAAGGTTTTGCTATTCATTGAAAGCAG
TAGTGACTGATTTGTATATA
>MinSyn_1504|Strength:0.000547956
GCGTGTCGTTTTAGTGAGGTTATGACCCCCGCCGATGACGCGGGAACCGATGCTATCCTTACCGCTAT
GGGTAAGATTGTGTACAGGGCTCACATTGCGATAAAGGAAAGGGATGCTGATCAAATATTTCTTGTAT
CAGCTTCTATATAAGGTTTTGCTATTCATTGAAAGCAGTAGTGACTGATTTGTATATA
>MinSyn_1694|Strength:0.000549676
GCGTGTCGTTTTAGTGAGGTCCATCAACAAATAATCCAAGTAAGCTATCAGAAGATCAAAGGGCTAAC
CACTATGCCCAATTAGGTTGTCTGCAGCAAGTGGATACAGCTTTGTCAAAAGCTAAAAAAGATGATGC
TCACATGTAGGCTCTATATAAGGTTTTGCTATTCATTGAAAGCAGTAGTGACTGATTTGTATATA
>MinSyn_1719|Strength:0.000550095
GCGTGTCGTTTTAGTGAGGCAGCCACTTGTGTCCTCACATAACCACGTCTACAATGCCTGCCTTGATG
TCCATCAACAAATAATCCAAGTAAGGCTTAGCAAGACCTCTACAAAACTGGTGCACACCAGCATGTGT
TGATCACCAGCTGTGTACAGGGTGAAGATAAGATAATAATGTTGAAGATAAGACTGAAGCATCTTCCT
CACTGCTAGGAGGACCGATAATTTCGGGAAACCTCCTCGTAGGATTTTCAACAACTTAGCAAGACCTC
CTATATAAGGTTTTGCTATTCATTGAAAGCAGTAGTGACTGATTTGTATATA
>MinSyn_1983|Strength:0.000551135
GCGTGTCGTTTTAGTGAGGTCTCTCTGCCGACAGTGGTCCCAAATGGTACTTGTGAAGATAAGATAAT
AATGTTGAAGATAAGAGGACCGATGCTGATCTTGCCTGCCTTATGACCCCCGCCGATGACGCGGGAGG
AATTTCGGGAAACCTCCTCGGAGGACCGATGCTGATCTTGCCTGCCTTCTATATAAGGTTTTGCTATT
CATTGAAAGCAGTAGTGACTGATTTGTATATA
>MinSyn_1555|Strength:0.000551145
GCGTGTCGTTTTAGTGAGGAGCAAGTGGATACATGTAGGCTATCAGCTTTCTCTCTGCCGACAGTGGT
CCCAAACCTATATAAGGTTTTGCTATTCATTGAAAGCAGTAGTGACTGATTTGTATATA
>MinSyn_1192|Strength:0.000551265
GCGTGTCGTTTTAGTGAGGCAGCCACTTGTGTATCTTGCCTGCCTTGATTGAAGATAAGATAATAATG

TTGAAGATAAGATGGTCAAATATTTCTTGTGGTACTTGTGTACAGGGCTCACCTATATAAGGTTTTGC
TATTCATTGAAAGCAGTAGTGACTGATTTGTATATA
>MinSyn_1367|Strength:0.000552027
GCGTGTCGTTTTAGTGAGGGCACACCAGCATGTGTTGATCACCAGCTGCACCTCACATGTCAGCCACT
TGTGTAGGTTGTCTGCACCTCACACTATATAAGGTTTTGCTATTCATTGAAAGCAGTAGTGACTGATT
TGTATATA
>MinSyn_1775|Strength:0.000552041
GCGTGTCGTTTTAGTGAGGTTTTCAACAATCAGCTTAGCAAGACCTCTACAAAATCAGAAGATCAAAG
GGCTACTGCCTTGATGATCCATCAACAAATAATCCAAGTAAGTAGCTATATAAGGTTTTGCTATTCAT
TGAAAGCAGTAGTGACTGATTTGTATATA
>MinSyn_1378|Strength:0.000552435
GCGTGTCGTTTTAGTGAGGTTATGACCCCCGCCGATGACGCGGGACCCAATTAGGTTGTCTGCTTTGT
CAAAAGCTAAAAAAGATGATGCCTCACTATCAGCAGCAAGACCTCTACAAAACTGGCTATATAAGGTT
TTGCTATTCATTGAAAGCAGTAGTGACTGATTTGTATATA
>MinSyn_1175|Strength:0.000553914
GCGTGTCGTTTTAGTGAGGTGAAGATAAGATAATAATGTTGAAGATAAGAAGCAGTGGTCCCTCCACG
TACAGGGCTCTCCATCAACAAATAATCCAAGTAAGCAATTAGGTTGTCTGCACCTGAAGCATCTTCCG
TACTTGTGTACAGGGCTCACTGCTAGAGCAAGTGGATGTACTTCAGAAGATCAAAGGGCTACTTGGTG
GAGCACGACAAGCAAGACCTCTACAAAACTGGTACTTGCACACCAGCATGTGTTGATCACCAGCTCTG
CTAGGAGGACCGATGCTGATTATGACCCCCGCCGATGACGCGGGAGTCTATATAAGGTTTTGCTATTC
ATTGAAAGCAGTAGTGACTGATTTGTATATA
>MinSyn_1825|Strength:0.000555323
GCGTGTCGTTTTAGTGAGGATCCTTACCGCTATGGGTAAGATTTTAGCAAGACCTCTACAAAAAACCA
TTATTGCGCTAGGAGGACCGAAATTTCGGGAAACCTCCTCGTAGGGGAAAAAGAAGAGGTATGCTGAT
CTTTTTTCAACAATACAAAACCTATATAAGGTTTTGCTATTCATTGAAAGCAGTAGTGACTGATTTGT
ATATA
>MinSyn_1403|Strength:0.000555882
GCGTGTCGTTTTAGTGAGGGTGGGAGCCACCAAGAGCAAGTGGATTATTGCGATAAAGGAAAGGAGGC
TATCAGCTTTGTCAAAAGCTAAAAAAGATGATGCTATCAGCTTAGCAAGACTGAAGCATCTTCCTTGT
CTGCACCTCACACTATATAAGGTTTTGCTATTCATTGAAAGCAGTAGTGACTGATTTGTATATA
>MinSyn_1364|Strength:0.000556439
GCGTGTCGTTTTAGTGAGGTGAAGATAAGATAATAATGTTGAAGATAAGACCGATGCTGATCTTGCCT
GCCAGCCACTTGTGTAAGGTGGCTCCTACACCTCTACAAAACTGGTACATCCTTACCGCTATGGGTAA
GATTTCACTGCTAAAAAATGTCAAAGATATCACATGCTATATAAGGTTTTGCTATTCATTGAAAGCAG
TAGTGACTGATTTGTATATA
>MinSyn_1115|Strength:0.000556951
GCGTGTCGTTTTAGTGAGGTTTTCAACAATACAAAACTGGTACTTGTGTACAGGGCTCAGCACACCAG
CATGTGTTGATCACCAGCTTAGGTTGTCTGCACCTCACAATCCTTACCGCTATGGGTAAGATTTGGTA
CCTATATAAGGTTTTGCTATTCATTGAAAGCAGTAGTGACTGATTTGTATATA
>MinSyn_1289|Strength:0.000557363
GCGTGTCGTTTTAGTGAGGTTATGACCCCCGCCGATGACGCGGGAACCTCACATGTAGGCTATCAGCT
TTCCATCAACAAATAATCCAAGTAAGGGTTGTCTGCACCTCACTGAAGCATCTTCCGTTGTGGGAGCC
ACCATGTAGGCTATCAGCTTAGCAAGACAAAAATGTCAAAGATATTGTGTACAGGGCTCACTGCTAAA
GATTGATGAAAAGTCAAAAACAAAAATCAATTATAGGACCGAAGGTGGCTCCTACGGTAGGTCACATC
CTTACCGCTATGGGTAAGATTGTACAGGAACCACGTCTACAAACAAAACTGGCTATATAAGGTTTTGC
TATTCATTGAAAGCAGTAGTGACTGATTTGTATATA
>MinSyn_1280|Strength:0.000559587
GCGTGTCGTTTTAGTGAGGTCTCTCTGCCGACAGTGGTCCCAAACTCAGAAGATCAAAGGGCTAGTCT
GCACCTCACATGTAGGCTATGGAAAAAGAAGAGGTTCACTGCTAGGAGGACCGATGCTGAAGATTGAT
GAAAAGTCAAAAACAAAAATCAATTATGTAGGTCACGACCACCAGCCACTTGTGTGCTATATAAGGTT
TTGCTATTCATTGAAAGCAGTAGTGACTGATTTGTATATA
>MinSyn_1538|Strength:0.000559589
GCGTGTCGTTTTAGTGAGGGCACACCAGCATGTGTTGATCACCAGCTCAATTAGGTTGTCTGCACCTC
ACAGCCACTTGTGTTGCACCTCCTATATAAGGTTTTGCTATTCATTGAAAGCAGTAGTGACTGATTTG

TATATA
>MinSyn_1603|Strength:0.000561479
GCGTGTCGTTTTAGTGAGGTGGTGGAGCACGACACACCTCACATGTAGGAAAAAGAAGAGGTGGTAGG
TTCCATCAACAAATAATCCAAGTAAGCTCTACAAAACCTATATAAGGTTTTGCTATTCATTGAAAGCA
GTAGTGACTGATTTGTATATA
>MinSyn_1232|Strength:0.000562025
GCGTGTCGTTTTAGTGAGGTCCATCAACAAATAATCCAAGTAAGATCAGCTTAGCAATGGTGGAGCAC
GACATCACTGCTAGGAGGACCGATGCTGATTGAAGATAAGATAATAATGTTGAAGATAAGAAGGTCAC
GACCACTATGCCCTTATGACCCCCGCCGATGACGCGGGATCTTGCCTGCCTTGCTTTGTCAAAAGCTA
AAAAAGATGATGCCTGATCTTGCCTGCCTTGATCTCTCTGCCGACAGTGGTCCCAAATCAGCTTAGCA
TTGCGATAAAGGAAAGGACTATGCCCACTATATAAGGTTTTGCTATTCATTGAAAGCAGTAGTGACTG
ATTTGTATATA
>MinSyn_163|Strength:0.000563448
GCGTGTCGTTTTAGTGAGGGCACACCAGCATGTGTTGATCACCAGCTTAGGTTGTCTGCACCTCACAT
GTAGGCCAGCCACTTGTGTGCCTATATAAGGTTTTGCTATTCATTGAAAGCAGTAGTGACTGATTTGT
ATATA
>MinSyn_1443|Strength:0.000565057
GCGTGTCGTTTTAGTGAGGATTGCGATAAAGGAAAGGAAAACTGGTACTTGTGTACAGGGTCAGAAGA
TCAAAGGGCTAGATCTTGCTTATGACCCCCGCCGATGACGCGGGAACCACTATGCCCAAATCGAAAGG
ACAGTAGTCTATATAAGGTTTTGCTATTCATTGAAAGCAGTAGTGACTGATTTGTATATA
>MinSyn_1340|Strength:0.000565481
GCGTGTCGTTTTAGTGAGGTGAAGCATCTTCCATGCCCAATTAGGTTGTCTTCACTATCAGCACTTGT
GTACAGGGCTCACTAAAAATGTCAAAGATACACATGTAGGCTATCAGCTTAGTCAGAAGATCAAAGGG
CTACAGCTTAGCAAGTGAAGATAAGATAATAATGTTGAAGATAAGATCGCAAATATTTCTTGTACAAA
ACTGGTACGGAAAAAGAAGAGGTGCAGCAAGTGGATTCTTGCCCTATATAAGGTTTTGCTATTCATTG
AAAGCAGTAGTGACTGATTTGTATATA
>MinSyn_147|Strength:0.000566225
GCGTGTCGTTTTAGTGAGGGCACACCAGCATGTGTTGATCACCAGCTCCTCACATGTAGGCTATCAGC
TTAGCAAGGGAAAAAGAAGAGGTCCTGCTATATAAGGTTTTGCTATTCATTGAAAGCAGTAGTGACTG
ATTTGTATATA
>MinSyn_1528|Strength:0.000567164
GCGTGTCGTTTTAGTGAGGAACCATTATTGCGCTTGCCTGCCTTGATTCCATCAACAAATAATCCAAG
TAAGCTTGTGCTATATAAGGTTTTGCTATTCATTGAAAGCAGTAGTGACTGATTTGTATATA
>MinSyn_1117|Strength:0.000570385
GCGTGTCGTTTTAGTGAGGGTGGGAGCCACCAAATTAGGTTGTCTGCACCTCACAATTGCGATAAAGG
AAAGGCAAGACCTCTACAAAACTGGTACTTGAAGCATCTTCCGTAGGCTATCAGCTTAGCAAGAGCAC
ACCAGCATGTGTTGATCACCAGCTTAACCATTATTGCGAATTCAGTGGTCCCTCCACACTATGCTATA
TAAGGTTTTGCTATTCATTGAAAGCAGTAGTGACTGATTTGTATATA
>MinSyn_1450|Strength:0.000570555
GCGTGTCGTTTTAGTGAGGATTGCGATAAAGGAAAGGCCTCTACAAAACTGGTACTTTCCATCAACAA
ATAATCCAAGTAAGCTCACTGCTAGGCAGCCACTTGTGTGCCTGCCTTGATGCAGTGGTCCCTCCACG
GACCGAGGTGGCTCCTACCATGTGAAGATAAGATAATAATGTTGAAGATAAGATGCGTGGGAGCCACC
AACCTCTACAAAACTGGTACTTGCTATATAAGGTTTTGCTATTCATTGAAAGCAGTAGTGACTGATTT
GTATATA
>MinSyn_1645|Strength:0.000570947
GCGTGTCGTTTTAGTGAGGAGCAAGTGGATGATGCTGATCTTGCCTGCCTTGATGATTGAAGCATCTT
CCCTGCACCTCACATGTAGGCTATCTTATGACCCCCGCCGATGACGCGGGAGGTAGGTCACGACCACT
ATGCCCAAATTTCGGGAAACCTCCTCGCCTGCCTCTCTCTGCCGACAGTGGTCCCAAAAAACTGGTAC
TTGTGTACAGGGTCCATCAACAAATAATCCAAGTAAGCAAAACTGGTACTTGTGTACAGCACACCAGC
ATGTGTTGATCACCAGCTGGGCTCACTGCTAGTGAAGATAAGATAATAATGTTGAAGATAAGATGTCC
TATATAAGGTTTTGCTATTCATTGAAAGCAGTAGTGACTGATTTGTATATA
>MinSyn_1736|Strength:0.000572008
GCGTGTCGTTTTAGTGAGGCAAATATTTCTTGTGATGATAAAAAATGTCAAAGATAGATCTTAAGATT
GATGAAAAGTCAAAAACAAAAATCAATTATTCAGCTTAGCAAGACTATATAAGGTTTTGCTATTCATT

GAAAGCAGTAGTGACTGATTTGTATATA
>MinSyn_1759|Strength:0.000573826
GCGTGTCGTTTTAGTGAGGGCACACCAGCATGTGTTGATCACCAGCTACTTGTGTACGGAAAAAGAAG
AGGTCTTGTGTACAGGGCTCACTGCCTATATAAGGTTTTGCTATTCATTGAAAGCAGTAGTGACTGAT
TTGTATATA
>MinSyn_1206|Strength:0.000574078
GCGTGTCGTTTTAGTGAGGCAAATATTTCTTGTACAGGGCTCACTGCGCTTTGTCAAAAGCTAAAAAA
GATGATGCATGTAGGCTATCAGCTTAGCAATTGCGATAAAGGAAAGGCTCTACAAAACTGTTATGACC
CCCGCCGATGACGCGGGATGGTATCAGAAGATCAAAGGGCTATTAGCAAGACCCTATATAAGGTTTTG
CTATTCATTGAAAGCAGTAGTGACTGATTTGTATATA
>MinSyn_1424|Strength:0.000574204
GCGTGTCGTTTTAGTGAGGATCGAAAGGACAGTACAAATTTTCAACAATGATCTTGCCTGCCTTGAAA
CCATTATTGCGACGCACACCAGCATGTGTTGATCACCAGCTGTGTACAGGGCTCACTGCTTGAAGATA
AGATAATAATGTTGAAGATAAGATCTACAAAACTGAAGCATCTTCCTATCAGCTTAGCAAGTCAGAAG
ATCAAAGGGCTATGCTATATAAGGTTTTGCTATTCATTGAAAGCAGTAGTGACTGATTTGTATATA
>MinSyn_1653|Strength:0.000576289
GCGTGTCGTTTTAGTGAGGTTATGACCCCCGCCGATGACGCGGGAATGAAGATAAGATAATAATGTTG
AAGATAAGAGCCTTGATGATATGAAGCATCTTCCACGACCACTATGCCCAATTAGCAAGTGGATCCTG
CCTTGATCTCTCTGCCGACAGTGGTCCCAAATAGGCTATATAAGGTTTTGCTATTCATTGAAAGCAGT
AGTGACTGATTTGTATATA
>MinSyn_1632|Strength:0.000577108
GCGTGTCGTTTTAGTGAGGAGGTGGCTCCTACATCAGCTTAGCAAGACCTCTACAAAACTGGTTATGA
CCCCCGCCGATGACGCGGGAACATTGCGATAAAGGAAAGGAACTATATAAGGTTTTGCTATTCATTGA
AAGCAGTAGTGACTGATTTGTATATA
>MinSyn_1733|Strength:0.000577204
GCGTGTCGTTTTAGTGAGGTGAAGATAAGATAATAATGTTGAAGATAAGACTATCAGCTTAGCAAGTG
GTGGAGCACGACACCTGCCTTAGGTGGCTCCTACCTGCTAGGAGGATCCTTACCGCTATGGGTAAGAT
TCTTAGCTATATAAGGTTTTGCTATTCATTGAAAGCAGTAGTGACTGATTTGTATATA
>MinSyn_180|Strength:0.000577577
GCGTGTCGTTTTAGTGAGGTGAAGATAAGATAATAATGTTGAAGATAAGAATAAAAATGTCAAAGATA
GTAGGTCACGACCACTATGCCCAATATCGAAAGGACAGTAGCTATCAGCTTAAACCATTATTGCGCAA
AACTGGTACTTGTGTACAGGGCTTCTCTCTGCCGACAGTGGTCCCAAATAGGTTGTCTGCAAGGTGGC
TCCTACCTGTGAAGCATCTTCCGCGGTAGGGCACACCAGCATGTGTTGATCACCAGCTCACTATATAA
GGTTTTGCTATTCATTGAAAGCAGTAGTGACTGATTTGTATATA
>MinSyn_1489|Strength:0.000582963
GCGTGTCGTTTTAGTGAGGAAAAATGTCAAAGATAACGACCACTATCAGAAGATCAAAGGGCTATTGA
TGAAGATTGATGAAAAGTCAAAAACAAAAATCAATTATCGATGCTGATCTTGCCTGCCTTTTTTCAAC
AAGATGCTGCAAATATTTCTTGTGTAGGTCACGTTATGACCCCCGCCGATGACGCGGGACTCTACACT
ATATAAGGTTTTGCTATTCATTGAAAGCAGTAGTGACTGATTTGTATATA
>MinSyn_1323|Strength:0.000583186
GCGTGTCGTTTTAGTGAGGTCACTATCAGCGCAAGACCTCTACAAAACTGGTACCAGCCACTTGTGTA
TGTAGGTCTCTCTGCCGACAGTGGTCCCAAACTACAAAACATCGAAAGGACAGTAATGATTTATGACC
CCCGCCGATGACGCGGGACCTCTACAGCAAGTGGATAGCTTAGAAGATTGATGAAAAGTCAAAAACAA
AAATCAATTATTTAGGTTGTCTGCACCTCACATGTAGTCAGAAGATCAAAGGGCTAGCAAATTTCGGG
AAACCTCCTCGCGGTAGGTCACGACCACTATATAAGGTTTTGCTATTCATTGAAAGCAGTAGTGACTG
ATTTGTATATA
>MinSyn_1243|Strength:0.000584045
GCGTGTCGTTTTAGTGAGGATCCTTACCGCTATGGGTAAGATTAACTGGTACTTGTGTACAGGGCTCA
CTGCAAGATTGATGAAAAGTCAAAAACAAAAATCAATTATAATCGAAAGGACAGTAACCTCACATGTA
GGCTCCATCAACAAATAATCCAAGTAAGGTCACGACCACTATGCCCTTATGACCCCCGCCGATGACGC
GGGATGCTAGGAGGAGCTTTGTCAAAAGCTAAAAAAGATGATGCATCAGCTTAGCAAGACCTCTACAA
TTGCGATAAAGGAAAGGTATCACTATATAAGGTTTTGCTATTCATTGAAAGCAGTAGTGACTGATTTG
TATATA
>MinSyn_1116|Strength:0.000585616

GCGTGTCGTTTTAGTGAGGGCACACCAGCATGTGTTGATCACCAGCTACCTATCGAAAGGACAGTAGA
TGCTGATCTTGCCTGCCTTGATCTATATAAGGTTTTGCTATTCATTGAAAGCAGTAGTGACTGATTTG
TATATA
>MinSyn_133|Strength:0.000585879
GCGTGTCGTTTTAGTGAGGCAAATATTTCTTGTCACCTCACTCTCTCTGCCGACAGTGGTCCCAAACG
ACCACTATGCCCAATTTGAAGATAAGATAATAATGTTGAAGATAAGACACTATGCCCTCCATCAACAA
ATAATCCAAGTAAGACCACTATGCCCAATTAGGCTTTGTCAAAAGCTAAAAAAGATGATGCCCTCACA
TGTAGGCTATCAGCTTTCAGAAGATCAAAGGGCTACAAGATTGATGAAAAGTCAAAAACAAAAATCAA
TTATACTTGTGTACAGGGCTCACTGCTGCACACCAGCATGTGTTGATCACCAGCTTGTCAGCCACTTG
TGTCGCGGTAGGTCACGACCACCTATATAAGGTTTTGCTATTCATTGAAAGCAGTAGTGACTGATTTG
TATATA
>MinSyn_1575|Strength:0.000586832
GCGTGTCGTTTTAGTGAGGAGGTGGCTCCTACTGCAAAAATGTCAAAGATAACTTGTGTACAGGGCTC
ACTGCTAGGAGGAGCACACCAGCATGTGTTGATCACCAGCTAGCAATCTCTCTGCCGACAGTGGTCCC
AAACTCACTGCTAGATTGCGATAAAGGAAAGGGACCGATCTATATAAGGTTTTGCTATTCATTGAAAG
CAGTAGTGACTGATTTGTATATA
>MinSyn_1659|Strength:0.00058877
GCGTGTCGTTTTAGTGAGGCAAATATTTCTTGTACCTCTACAAAACTGGTACTTGTGTATGAAGCATC
TTCCCTACAAAAGGTGGCTCCTACCTTAGCAAGACCTCTACAAAACTCAGTGGTCCCTCCACGGTAGG
TCACGACCACTATTATGACCCCCGCCGATGACGCGGGAGTAGGTCACGACCACTATGCCAAAAATGTC
AAAGATAGGTAGGTCACAAGATTGATGAAAAGTCAAAAACAAAAATCAATTATGGTAGGTCATCCTTA
CCGCTATGGGTAAGATTACTCTATATAAGGTTTTGCTATTCATTGAAAGCAGTAGTGACTGATTTGTA
TATA
>MinSyn_1508|Strength:0.000589747
GCGTGTCGTTTTAGTGAGGGCACACCAGCATGTGTTGATCACCAGCTACAGGGCTCGTGGGAGCCACC
ATCACATGTAAAGATTGATGAAAAGTCAAAAACAAAAATCAATTATCTATCAGCTTAGCAAGACCTAT
CGAAAGGACAGTAAACTGGTACTTGTGCTATATAAGGTTTTGCTATTCATTGAAAGCAGTAGTGACTG
ATTTGTATATA
>MinSyn_167|Strength:0.000590225
GCGTGTCGTTTTAGTGAGGTTTTCAACAATCAGCTTAGCGCTTTGTCAAAAGCTAAAAAAGATGATGC
GTGGTGGAGCACGACATGATCTTGCCTGCCTTGATCTATATAAGGTTTTGCTATTCATTGAAAGCAGT
AGTGACTGATTTGTATATA
>MinSyn_156|Strength:0.000592104
GCGTGTCGTTTTAGTGAGGAAAAATGTCAAAGATAGTGTACAATTTCGGGAAACCTCCTCGTCTACAA
AACTGGTACTTGTAAGATTGATGAAAAGTCAAAAACAAAAATCAATTATCTTGTGTACAGGGCTCACT
CAGTGGTCCCTCCACCTTACTATATAAGGTTTTGCTATTCATTGAAAGCAGTAGTGACTGATTTGTAT
ATA
>MinSyn_1941|Strength:0.000593477
GCGTGTCGTTTTAGTGAGGAACCATTATTGCGGCAAGACCTCTACATCCATCAACAAATAATCCAAGT
AAGCTATGGTGGGAGCCACCACTACGCACACCAGCATGTGTTGATCACCAGCTACTGCTAGGTGAAGA
TAAGATAATAATGTTGAAGATAAGACCGATGCTGATCTTGCCTATATAAGGTTTTGCTATTCATTGAA
AGCAGTAGTGACTGATTTGTATATA
>MinSyn_1933|Strength:0.000595927
GCGTGTCGTTTTAGTGAGGAAAAATGTCAAAGATAACCACTATGCCCAATAACCACGTCTACAAAGGA
GGACCGATGCTGATCTTGCGTGGGAGCCACCACAATTATCGAAAGGACAGTAAGGGCTCACTGCTAGG
AATTTCGGGAAACCTCCTCGCTATCAGCTTAGGCTTTGTCAAAAGCTAAAAAAGATGATGCCAAGACC
TCTACTGAAGATAAGATAATAATGTTGAAGATAAGACAGGGCTCAGGAAAAAGAAGAGGTCTACAAAA
CTGGTCAGAAGATCAAAGGGCTAACTATATAAGGTTTTGCTATTCATTGAAAGCAGTAGTGACTGATT
TGTATATA
>MinSyn_1812|Strength:0.000596173
GCGTGTCGTTTTAGTGAGGCAGCCACTTGTGTCTACAAAACTGGTACTTGTGTGGAAAAAGAAGAGGT
GGCTATCAGAGGTGGCTCCTACCTTCAGAAGATCAAAGGGCTAGGACCGATGCTGAATCGAAAGGACA
GTATCTATATAAGGTTTTGCTATTCATTGAAAGCAGTAGTGACTGATTTGTATATA
>MinSyn_1227|Strength:0.00059635

GCGTGTCGTTTTAGTGAGGAAAAATGTCAAAGATACACCTCACATGTAGGCTCCATCAACAAATAATC
CAAGTAAGTATGCCTATATAAGGTTTTGCTATTCATTGAAAGCAGTAGTGACTGATTTGTATATA
>MinSyn_183|Strength:0.000596353
GCGTGTCGTTTTAGTGAGGAAGATTGATGAAAAGTCAAAAACAAAAATCAATTATGTCACGACCACTA
TGCCCAATTAGGTTGTCAACCATTATTGCGACTGCTAGTGGGAGCCACCACACATGTAGGCTTCCATC
AACAAATAATCCAAGTAAGGGTACTTGTTGAAGATAAGATAATAATGTTGAAGATAAGATGCCCAATT
AGGTTGTCTGCACGCACACCAGCATGTGTTGATCACCAGCTCTAGTGAAGCATCTTCCCGGTACTATA
TAAGGTTTTGCTATTCATTGAAAGCAGTAGTGACTGATTTGTATATA
>MinSyn_1349|Strength:0.00059967
GCGTGTCGTTTTAGTGAGGGGAAAAAGAAGAGGTTACTTGTGTACAGGGCTCACTGAAGATTGATGAA
AAGTCAAAAACAAAAATCAATTATGACCTCTACAAACAGCCACTTGTGTGACCCTATATAAGGTTTTG
CTATTCATTGAAAGCAGTAGTGACTGATTTGTATATA
>MinSyn_1766|Strength:0.000603221
GCGTGTCGTTTTAGTGAGGTTTTCAACAACTGATCGCACACCAGCATGTGTTGATCACCAGCTGGTCA
CGACCACTATGCCCCAGCCACTTGTGTTCAGCTTAGCAAGGAAAAAGAAGAGGTTAGGTTGTCTGCAC
CTTATGACCCCCGCCGATGACGCGGGATGGTCAAATATTTCTTGTACGACCACTATAACCATTATTGC
GCCTATATAAGGTTTTGCTATTCATTGAAAGCAGTAGTGACTGATTTGTATATA
>MinSyn_1634|Strength:0.000605072
GCGTGTCGTTTTAGTGAGGGTGGGAGCCACCAAGGGCTCACTGCTAGGAGGACCGCACACCAGCATGT
GTTGATCACCAGCTCTTGAAAAATGTCAAAGATAGCAGCCACTTGTGTAGGTCCAAATATTTCTTGTA
TCTTCACTATCAGCTACTTGTGTACAGCTATATAAGGTTTTGCTATTCATTGAAAGCAGTAGTGACTG
ATTTGTATATA
>MinSyn_161|Strength:0.000609519
GCGTGTCGTTTTAGTGAGGTCTCTCTGCCGACAGTGGTCCCAAATCACTGCTAGGAGGACCGAATCCT
TACCGCTATGGGTAAGATTTGCACCTCACATGTAGGCTATCATCCATCAACAAATAATCCAAGTAAGC
AGCTTAGCAAGACCTCTAGGAAAAAGAAGAGGTCATGTAGGCTATCAGCTTAGCGCTTTGTCAAAAGC
TAAAAAAGATGATGCACCGATATTGCGATAAAGGAAAGGGCGGTAGGTCACGATGAAGATAAGATAAT
AATGTTGAAGATAAGAGATCGAAAGGACAGTATGTCTATATAAGGTTTTGCTATTCATTGAAAGCAGT
AGTGACTGATTTGTATATA
>MinSyn_1494|Strength:0.000610683
GCGTGTCGTTTTAGTGAGGATTGCGATAAAGGAAAGGTGATCTTGCCTGCCTTGATGATCCTTACCGC
TATGGGTAAGATTAGGAGGACCGATGCTGATCAGAAGATCAAAGGGCTACTTGAAAAAATGTCAAAGA
TACTAGGACAGTGGTCCCTCCACACTGCTATATAAGGTTTTGCTATTCATTGAAAGCAGTAGTGACTG
ATTTGTATATA
>MinSyn_1794|Strength:0.000613386
GCGTGTCGTTTTAGTGAGGCAGTGGTCCCTCCACTGCTAGGAGTTTTCAACAAGAACCATTATTGCGC
CGACTATATAAGGTTTTGCTATTCATTGAAAGCAGTAGTGACTGATTTGTATATA
>MinSyn_1310|Strength:0.0006135
GCGTGTCGTTTTAGTGAGGAGCAAGTGGATACTATCCTTACCGCTATGGGTAAGATTGCTTAGCAAGA
CCTCTATCAGAAGATCAAAGGGCTACGCCTATATAAGGTTTTGCTATTCATTGAAAGCAGTAGTGACT
GATTTGTATATA
>MinSyn_1889|Strength:0.00061389
GCGTGTCGTTTTAGTGAGGGGAAAAAGAAGAGGTCTACAAAACTGGTACTGTGGGAGCCACCATGTAG
GCTATCAGCTTAGCAAGACTCACTATCAGCTGTACAGGGCTCACTGCTAGGAGGACCGTCCATCAACA
AATAATCCAAGTAAGCCCAATTAGGTTGCTTTGTCAAAAGCTAAAAAAGATGATGCTTGAAGATAAGA
TAATAATGTTGAAGATAAGAACTCTATATAAGGTTTTGCTATTCATTGAAAGCAGTAGTGACTGATTT
GTATATA
>MinSyn_1207|Strength:0.000615862
GCGTGTCGTTTTAGTGAGGAATTTCGGGAAACCTCCTCGACCGATGCTTCCATCAACAAATAATCCAA
GTAAGTCACGACCACTATGCCCTATATAAGGTTTTGCTATTCATTGAAAGCAGTAGTGACTGATTTGT
ATATA
>MinSyn_1347|Strength:0.000616084
GCGTGTCGTTTTAGTGAGGCAGCCACTTGTGTGGCTATCAGCTTAGCAAGACCTCTACAAAATTGCGA
TAAAGGAAAGGGTAGCAAGTGGATTGTAGGCTATCAGCTTAGCAAGACAAGATTGATGAAAAGTCAAA

AACAAAAATCAATTATTAGGCTATCAGCTTAGCAAGACCTCTACAAACCATTATTGCGATTGAAGCAT
CTTCCGTTATGACCCCCGCCGATGACGCGGGAGCGGTAGGTGAAGATAAGATAATAATGTTGAAGATA
AGACCACTATGCCTATATAAGGTTTTGCTATTCATTGAAAGCAGTAGTGACTGATTTGTATATA
>MinSyn_1246|Strength:0.000616216
GCGTGTCGTTTTAGTGAGGATCGAAAGGACAGTATTCAGAAGATCAAAGGGCTAACTATATAAGGTTT
TGCTATTCATTGAAAGCAGTAGTGACTGATTTGTATATA
>MinSyn_1802|Strength:0.000618939
GCGTGTCGTTTTAGTGAGGGTGGGAGCCACCACATGTAGGCTGCACACCAGCATGTGTTGATCACCAG
CTCACGACCTATATAAGGTTTTGCTATTCATTGAAAGCAGTAGTGACTGATTTGTATATA
>MinSyn_1992|Strength:0.000622954
GCGTGTCGTTTTAGTGAGGATCCTTACCGCTATGGGTAAGATTTCACGACCACAAGATTGATGAAAAG
TCAAAAACAAAAATCAATTATCTCTACAAAACTGGTACTTGTTGGTGGAGCACGACATGCTAGGCTAT
ATAAGGTTTTGCTATTCATTGAAAGCAGTAGTGACTGATTTGTATATA
>MinSyn_1311|Strength:0.000627252
GCGTGTCGTTTTAGTGAGGAGGTGGCTCCTACCACATGTAGGCTAGCTTTGTCAAAAGCTAAAAAAGA
TGATGCGGTATGGTGGAGCACGACAGAACCATTATTGCGACCACTATGTGAAGATAAGATAATAATGT
TGAAGATAAGAAAGACCTCTTCAGAAGATCAAAGGGCTAAAACTGGTACTTGTGTACCTATATAAGGT
TTTGCTATTCATTGAAAGCAGTAGTGACTGATTTGTATATA
>MinSyn_1410|Strength:0.000633333
GCGTGTCGTTTTAGTGAGGCAAATATTTCTTGTGGCACACCAGCATGTGTTGATCACCAGCTTGTGTA
CAGGGCTCCTATATAAGGTTTTGCTATTCATTGAAAGCAGTAGTGACTGATTTGTATATA
>MinSyn_1564|Strength:0.000642446
GCGTGTCGTTTTAGTGAGGTGGTGGAGCACGACATCACTGCTAGGAAGGTGGCTCCTACTGGTAAAAA
ATGTCAAAGATAACACTATATAAGGTTTTGCTATTCATTGAAAGCAGTAGTGACTGATTTGTATATA
>MinSyn_1334|Strength:0.000642857
GCGTGTCGTTTTAGTGAGGATCCTTACCGCTATGGGTAAGATTCACTATTCAGAAGATCAAAGGGCTA
ACAGGGCTCCTATATAAGGTTTTGCTATTCATTGAAAGCAGTAGTGACTGATTTGTATATA
>MinSyn_1822|Strength:0.000643475
GCGTGTCGTTTTAGTGAGGGTGGGAGCCACCACCCAAATATTTCTTGTCTTGTGTACAGGGCTCACGG
AAAAAGAAGAGGTTGCAAGATTGATGAAAAGTCAAAAACAAAAATCAATTATACCTCACATCTATATA
AGGTTTTGCTATTCATTGAAAGCAGTAGTGACTGATTTGTATATA
>MinSyn_1895|Strength:0.000643802
GCGTGTCGTTTTAGTGAGGATCCTTACCGCTATGGGTAAGATTTATCACAGTGGTCCCTCCACGGCTA
TATAAGGTTTTGCTATTCATTGAAAGCAGTAGTGACTGATTTGTATATA
>MinSyn_1821|Strength:0.000644068
GCGTGTCGTTTTAGTGAGGTCCATCAACAAATAATCCAAGTAAGGCATGAAGCATCTTCCCACTATAT
AAGGTTTTGCTATTCATTGAAAGCAGTAGTGACTGATTTGTATATA
>MinSyn_1875|Strength:0.000645322
GCGTGTCGTTTTAGTGAGGATTGCGATAAAGGAAAGGTAGGAGGACCAAATATTTCTTGTTGTATCTC
TCTGCCGACAGTGGTCCCAAAGGTACTTGTGTACAGGGCTCATGAAGATAAGATAATAATGTTGAAGA
TAAGAGGAAAAATGTCAAAGATAATCCTATATAAGGTTTTGCTATTCATTGAAAGCAGTAGTGACTGA
TTTGTATATA
>MinSyn_1844|Strength:0.000655392
GCGTGTCGTTTTAGTGAGGAGGTGGCTCCTACCGACAAATATTTCTTGTCGACCGCTTTGTCAAAAGC
TAAAAAAGATGATGCATGTAGGCTATATAAGGTTTTGCTATTCATTGAAAGCAGTAGTGACTGATTTG
TATATA
>MinSyn_1803|Strength:0.000656391
GCGTGTCGTTTTAGTGAGGAGCAAGTGGATGGACCGATGCTGATCTTGCCTGCCTTGATGTCACTATC
AGCCCTCACATGTAAGATTGATGAAAAGTCAAAAACAAAAATCAATTATAAACTGTGAAGATAAGAT
AATAATGTTGAAGATAAGATCTTGCCTGCCCTATATAAGGTTTTGCTATTCATTGAAAGCAGTAGTGA
CTGATTTGTATATA
>MinSyn_1860|Strength:0.000662791
GCGTGTCGTTTTAGTGAGGGCACACCAGCATGTGTTGATCACCAGCTTAGGTCACGAGGAAAAAGAAG
AGGTTCTATATAAGGTTTTGCTATTCATTGAAAGCAGTAGTGACTGATTTGTATATA

>MinSyn_1436|Strength:0.000683436
GCGTGTCGTTTTAGTGAGGAAGATTGATGAAAAGTCAAAAACAAAAATCAATTATAAATCGAAAGGAC
AGTACTTGATGATGCTTTGTCAAAAGCTAAAAAAGATGATGCATCAGAAGATCAAAGGGCTAGGAGGA
CCGATGCTGACTATATAAGGTTTTGCTATTCATTGAAAGCAGTAGTGACTGATTTGTATATA
>MinSyn_1290|Strength:0.000684
GCGTGTCGTTTTAGTGAGGAAGATTGATGAAAAGTCAAAAACAAAAATCAATTATCTGCTAGGAGGAC
CGATAAAAATGTCAAAGATAGATGCTCTATATAAGGTTTTGCTATTCATTGAAAGCAGTAGTGACTGA
TTTGTATATA
>MinSyn_1248|Strength:0.000685262
GCGTGTCGTTTTAGTGAGGCAGTGGTCCCTCCACGGTACTGTGGGAGCCACCACCAAATATTTCTTGT
ATGTAGGCTAAATTTCGGGAAACCTCCTCGACTATATAAGGTTTTGCTATTCATTGAAAGCAGTAGTG
ACTGATTTGTATATA
>MinSyn_1757|Strength:0.000697656
GCGTGTCGTTTTAGTGAGGTTATGACCCCCGCCGATGACGCGGGAAAACTTCAGAAGATCAAAGGGCT
ACAACTATATAAGGTTTTGCTATTCATTGAAAGCAGTAGTGACTGATTTGTATATA
>MinSyn_1139|Strength:0.000722122
GCGTGTCGTTTTAGTGAGGGTGGGAGCCACCAGCTCACTGCTAGGAGGACTGAAGATAAGATAATAAT
GTTGAAGATAAGAACAAAGGAAAAAGAAGAGGTAGGAGGTGGCTCCTACCCTATATAAGGTTTTGCTA
TTCATTGAAAGCAGTAGTGACTGATTTGTATATA
>MinSyn_1498|Strength:0.000737638
GCGTGTCGTTTTAGTGAGGATGACGTAAGCCATGACGTCTATACTTGTGTATCAGAAGATCAAAGGGC
TACCGATGCTGATCTTGCCTGCCTTGCAGCCACTTGTGTGTACTTGTGTACAGGGCTCACTGCTAGGA
GAGGTGGCTCCTACTTAGGTAATTTCGGGAAACCTCCTCGGAGCTTTGTCAAAAGCTAAAAAAGATGA
TGCATGTAGGCTATCAGCTTAGCAAGACCTAACCACGTCTACAACTGCACCTCACATGTAGGCTATCA
GCGTGGGAGCCACCATGCTAGGAGGACCGATGCTGATCTTGATCGAAAGGACAGTATACAACTATATA
AGGTTTTGCTATTCATTGAAAGCAGTAGTGACTGATTTGTATATA
>MinSyn_1274|Strength:0.000809032
GCGTGTCGTTTTAGTGAGGAAGATTGATGAAAAGTCAAAAACAAAAATCAATTATCTAGGAGGAGCAC
ACCAGCATGTGTTGATCACCAGCTAAACTGGCTATATAAGGTTTTGCTATTCATTGAAAGCAGTAGTG
ACTGATTTGTATATA
>MinSyn_1281|Strength:0.000867417
GCGTGTCGTTTTAGTGAGGATGACGTAAGCCATGACGTCTACACCTCACACAGCCACTTGTGTGCTCA
CTGCTAGGAGAACCACGTCTACAAGCTCACTGCTAGGAGGACCGATGCTGATCTAACCATTATTGCGG
ACCACTATGCCCAATTAATTTCGGGAAACCTCCTCGGCGGTAGGTCACGACCACGTGGGAGCCACCAT
ACAGGGCTCACTGCTATCACTATCAGCTTGCCTGCCTTGATGATATCCTTACCGCTATGGGTAAGATT
ATGTAGGCTATCAGCTTTATGACCCCCGCCGATGACGCGGGAAATTAGGTTGTCTGCACTATATAAGG
TTTTGCTATTCATTGAAAGCAGTAGTGACTGATTTGTATATA
>MinSyn_158|Strength:0.001731679
GCGTGTCGTTTTAGTGAGGCTGACGTAAGGGATGACGCACAGACCTCTACAAAACTGGTACTTGTGTC
AGAAGATCAAAGGGCTAACTGGTACTTGTGTACAGGGCTATCGAAAGGACAGTATGATGTGGGAGCCA
CCACCTGCCTTGCAAATATTTCTTGTCTTGCCTGCCAAAAATGTCAAAGATATTAGCAAGACCTCTAC
AAAACTGGTACTGAAGCATCTTCCACTGCTAGGAGGACCGATGCTGAACCATTATTGCGTAGCAAGAC
CTCTACAAAACTGGTACTTGCAGCCACTTGTGTCTACTATATAAGGTTTTGCTATTCATTGAAAGCAG
TAGTGACTGATTTGTATATA
>MinSyn_1628|Strength:0.001809417
GCGTGTCGTTTTAGTGAGGCTGACGTAAGGGATGACGCACAAAACTGGTACTTGTGTACAGGGCTCAC
GTGGGAGCCACCACATCGAAAGGACAGTAGACCACTATGCCTGGTGGAGCACGACAGTACAGGGCTCA
CTGCTAGGAGGACCGATCAGCCACTTGTGTTAGGTCAGGAAAAAGAAGAGGTCCTGCCTGCACACCAG
CATGTGTTGATCACCAGCTCACTATGCCCAATCCATCAACAAATAATCCAAGTAAGTCTACAAAATTG
CGATAAAGGAAAGGGCTCACTGCTAGGAGGACCGATGCTATATAAGGTTTTGCTATTCATTGAAAGCA
GTAGTGACTGATTTGTATATA
>MinSyn_1548|Strength:0.001854156
GCGTGTCGTTTTAGTGAGGCTGACGTAAGGGATGACGCACAAGCAAGACCTCTACTGGTGGAGCACGA
CACCTCTACAAAACTGGTACTTGTGTACTGAAGCATCTTCCCAGGGCTCACTGCTAGGAGGACCGATG

CTAATTTCGGGAAACCTCCTCGCCTTAAGATTGATGAAAAGTCAAAAACAAAAATCAATTATACTGCT
AGGAGGACCGGGAAAAAGAAGAGGTCCTCTACAAAGTGGGAGCCACCACGACCACTATGCCCAATTAG
GTTGTCTGCAGCCACTTGTGTACATGTAGGCTATCTATATAAGGTTTTGCTATTCATTGAAAGCAGTA
GTGACTGATTTGTATATA
>MinSyn_1303|Strength:0.002065551
GCGTGTCGTTTTAGTGAGGTCCATCAACAAATAATCCAAGTAAGGCTCACTGCTAGGAGGACCGGTGG
GAGCCACCAGACCACTATGCCCAATTAGGCTGACGTAAGGGATGACGCACACTGCACCTCACATGTAG
AAGATTGATGAAAAGTCAAAAACAAAAATCAATTATAGCAAGACCTCTACAAAACTGGTACATCCTTA
CCGCTATGGGTAAGATTGCACCTCACATGTAGGCTATCAGCTTAGTGAAGCATCTTCCCAAGACCTCT
ACAAAACTGGTACTGGAAAAAGAAGAGGTTGTAGGCTATCAGCTTAGTTATGACCCCCGCCGATGACG
CGGGATTGTCTGCACCTCACATGTAGGCTATCTATATAAGGTTTTGCTATTCATTGAAAGCAGTAGTG
ACTGATTTGTATATA
>MinSyn_125|Strength:0.00219106
GCGTGTCGTTTTAGTGAGGAACCACGTCTACAAGCTTAGCAAGACCTCTACAAAACATGACGTAAGCC
ATGACGTCTAAGCTTAGCATCCATCAACAAATAATCCAAGTAAGGCTAGGAGGACCGATGCTGATCTT
AACCATTATTGCGCGACCACTATGCCCAATTAGGTTGTCTGCATTGCGATAAAGGAAAGGCTCACATG
TAGGCTATCAGCTTTCAGAAGATCAAAGGGCTACCGAATTTCGGGAAACCTCCTCGCCTCTACAAAAC
TGGTACTGCTTTGTCAAAAGCTAAAAAAGATGATGCTACAAAACTGGTACTTGTGTACAGCTATATAA
GGTTTTGCTATTCATTGAAAGCAGTAGTGACTGATTTGTATATA
>MinSyn_1461|Strength:0.002253715
GCGTGTCGTTTTAGTGAGGTTATGACCCCCGCCGATGACGCGGGACAATTAGGTTGTCTGCACTGACG
TAAGGGATGACGCACATGTACAGGGCTCACTGCTAGGAGGATGGTGGAGCACGACAGGAGGACCGATG
CTGATCTTGCCTGCCTAAGATTGATGAAAAGTCAAAAACAAAAATCAATTATCTGGTACTTGTGTACA
GGGCTCACTGCTCAGCCACTTGTGTATCAGCTTAGCAAGACCTCTACAAAACAGCAAGTGGATACTGC
TAGGAGGACCGATGCTGATCTTGTGAAGCATCTTCCGTTGTCTGCACCTCACATGTAGGCTCTATATA
AGGTTTTGCTATTCATTGAAAGCAGTAGTGACTGATTTGTATATA
>MinSyn_1944|Strength:0.002306431
GCGTGTCGTTTTAGTGAGGATGACGTAAGCCATGACGTCTACATCGAAAGGACAGTACGGGGAAAAAG
AAGAGGTTCTGCACCTTCCATCAACAAATAATCCAAGTAAGACCTCTACAAATGGTGGAGCACGACAC
CAATTAGGTTGTCTGTGAAGCATCTTCCGCTAGGAGGACCGATGCTGGCACACCAGCATGTGTTGATC
ACCAGCTGGCAAATATTTCTTGTTTAGCAAGACCTCTACAAAACTGGTAAACCATTATTGCGCCTCTA
CAAAACTGGTACTTGTGTACAGGGCTATATAAGGTTTTGCTATTCATTGAAAGCAGTAGTGACTGATT
TGTATATA
>MinSyn_1905|Strength:0.002335471
GCGTGTCGTTTTAGTGAGGCTGACGTAAGGGATGACGCACACGCGGTAGGTCAGAAGATCAAAGGGCT
ACAAGATCGAAAGGACAGTACTCAGTGGTCCCTCCACGACCTCTACAAAACTGGTAGCACACCAGCAT
GTGTTGATCACCAGCTCTATGCCCAATTAGGTTGTCTGCACCTCACATTGCGATAAAGGAAAGGGCTT
AGCAAGACAGCCACTTGTGTTAGGCTATCAGCTTAGCAAGACCTCTACGCTTTGTCAAAAGCTAAAAA
AGATGATGCGTCTGCACCTCACATCTATATAAGGTTTTGCTATTCATTGAAAGCAGTAGTGACTGATT
TGTATATA
>MinSyn_1807|Strength:0.002478067
GCGTGTCGTTTTAGTGAGGCTGACGTAAGGGATGACGCACAGCAAGACCTCTACAAAACTGGTTTTTC
AACAATGGTACTTGTGTACAGGGCTCACGCTTTGTCAAAAGCTAAAAAAGATGATGCATCAGAAGATC
AAAGGGCTAGCTCACTGCTAGGAGGACCGATGTCCATCAACAAATAATCCAAGTAAGTTGTCTGCACC
TCACATGTAGGCTTCTCTCTGCCGACAGTGGTCCCAAACCTCTACAAAACTGGTACTTATTGCGATAA
AGGAAAGGTGTCTGCACCTCACTATATAAGGTTTTGCTATTCATTGAAAGCAGTAGTGACTGATTTGT
ATATA
>MinSyn_1613|Strength:0.002614575
GCGTGTCGTTTTAGTGAGGGCACACCAGCATGTGTTGATCACCAGCTCAGCTTAGCAAGACCTCAGGT
GGCTCCTACAATTAGGTATGACGTAAGCCATGACGTCTAGGGCTCACTGCTAGGAGAAGATTGATGAA
AAGTCAAAAACAAAAATCAATTATATCTTGCCTGCCTTGATGAAAAATGTCAAAGATAATTAGGTTGT
CTGCACCTCACATGTAGGCTGTGGGAGCCACCATGATCTTGCCTGCCTTGATGGGAAAAAGAAGAGGT
CCACTATGCCCAATTAGATCCTTACCGCTATGGGTAAGATTAGGCTAACCACGTCTACAAGGGCTCAC
TGCTAGGACTATATAAGGTTTTGCTATTCATTGAAAGCAGTAGTGACTGATTTGTATATA

>MinSyn_1160|Strength:0.002819945
GCGTGTCGTTTTAGTGAGGGCTTTGTCAAAAGCTAAAAAAGATGATGCGCTCACTGCTAGGAGGACCG
ATGCTGATCATTGCGATAAAGGAAAGGAAACTGGTACTTGTGTACAGATGACGTAAGCCATGACGTCT
ACTCCATCAACAAATAATCCAAGTAAGATTAGGTTGTCTGCACCTCACATGTGAAGATAAGATAATAA
TGTTGAAGATAAGACTTAGCAAGACCTCTACAAAACTGATCGAAAGGACAGTAGCGGTACAAATATTT
CTTGTCTAGGAGGACCGATGCTGATCTTGCCAAAAATGTCAAAGATAAAGACCTCTACAAAACTGGTA
TCACTATCAGCCTATCAGCTTAGCAAGACTATATAAGGTTTTGCTATTCATTGAAAGCAGTAGTGACT
GATTTGTATATA
>MinSyn_1908|Strength:0.003027735
GCGTGTCGTTTTAGTGAGGATGACGTAAGCCATGACGTCTATGCACCTCACATGTAGGCTATCTGAAG
CATCTTCCACTGCTAGGAGGACCGATGCTGATCTTGCCAATTTCGGGAAACCTCCTCGTGTGTACAGG
GCTCATCCTTACCGCTATGGGTAAGATTCCTCTACAAAACTGGTAAACCATTATTGCGGCGGTGCTTT
GTCAAAAGCTAAAAAAGATGATGCTGCACCTCACATGTAGGCTATCAGCTTAGTTTTCAACAACAGCT
TAGCAAGACCTATATAAGGTTTTGCTATTCATTGAAAGCAGTAGTGACTGATTTGTATATA
>MinSyn_1371|Strength:0.003065516
GCGTGTCGTTTTAGTGAGGTCAGAAGATCAAAGGGCTATTGTGTACAGGGCTCACTGCTCTGACGTAA
GGGATGACGCACATGGTTCCATCAACAAATAATCCAAGTAAGACAAAACTGGTACTTGTGTATCGAAA
GGACAGTAAAAACTGGTACTTGTGTACAGGGCTTTTTCAACAAGTCACGACCACTAAATTTCGGGAAA
CCTCCTCGAATTAGGTTGTCTGCACCTCACATGTAGCTTTGTCAAAAGCTAAAAAAGATGATGCACCG
ATGCTGATCTCTCTCTGCCGACAGTGGTCCCAAATACAGGGCTCCTATATAAGGTTTTGCTATTCATT
GAAAGCAGTAGTGACTGATTTGTATATA
>MinSyn_1118|Strength:0.003267163
GCGTGTCGTTTTAGTGAGGATGACGTAAGCCATGACGTCTAATCAGCTTAGCAAGTCTCTCTGCCGAC
AGTGGTCCCAAAGTAGGTCACGACCACTATGCCCAATAATTTCGGGAAACCTCCTCGTCTGCACCTCA
CATGTAGGCTAAAAATGTCAAAGATATCGCGGTAGGTCACGACCACTATGCCCAGCTTTGTCAAAAGC
TAAAAAAGATGATGCGGTAGGTCACGACCACTATGCCCAATTAGTTATGACCCCCGCCGATGACGCGG
GACACCTATATAAGGTTTTGCTATTCATTGAAAGCAGTAGTGACTGATTTGTATATA
>MinSyn_1108|Strength:0.003273603
GCGTGTCGTTTTAGTGAGGCTGACGTAAGGGATGACGCACACAATTAGGTTGTCTGCACCTCACATGT
ACAGTGGTCCCTCCACCACCTCACATGTAGGCTATGGAAAAAGAAGAGGTACTGGTACTTGTGTACAG
GGCTCACTGCTCAAATATTTCTTGTCGGTAGGTCACGACCACTATGCCCATGAAGCATCTTCCCACCT
CACATGTAGGCTATCAGCTTAGCATCACTATCAGCATCTTGCCTTATGACCCCCGCCGATGACGCGGG
AAGCCTATATAAGGTTTTGCTATTCATTGAAAGCAGTAGTGACTGATTTGTATATA
>MinSyn_1136|Strength:0.003366746
GCGTGTCGTTTTAGTGAGGATCGAAAGGACAGTATGCCTGCCCAGCCACTTGTGTAATTAGGTTGTCT
GCACCTCAATTGCGATAAAGGAAAGGACCGATGCTGATCTTGCCTCTGACGTAAGGGATGACGCACAC
TCACATGTAGGCTATCAGCTTAGAGGTGGCTCCTACTCGCGGTAGGTCAATCCTTACCGCTATGGGTA
AGATTAAACTGGTACTTGTGTACAGGGCACACCAGCATGTGTTGATCACCAGCTGTACTTGTGTACAG
GAAGATTGATGAAAAGTCAAAAACAAAAATCAATTATTGCACCTCACATGTAGGCTATCAAAAATGTC
AAAGATAAGGAGGACCTATATAAGGTTTTGCTATTCATTGAAAGCAGTAGTGACTGATTTGTATATA
>MinSyn_1648|Strength:0.00390934
GCGTGTCGTTTTAGTGAGGAAAAATGTCAAAGATATAGGAGGACCGATGCTGATCTTGCGTGGGAGCC
ACCAGACCTCTACACTGACGTAAGGGATGACGCACACTAGGAGGACCGATGCTGATCTTATCGAAAGG
ACAGTAGTCAGAAGATCAAAGGGCTAACCTCTACATTGCGATAAAGGAAAGGAGGTTGAATTTCGGGA
AACCTCCTCGGTACTTGTGTACAGGGCTCACTGCTAGGAGGCACACCAGCATGTGTTGATCACCAGCT
TGCCTGCATCCTTACCGCTATGGGTAAGATTTAGGCTATCAGCTTACTATATAAGGTTTTGCTATTCA
TTGAAAGCAGTAGTGACTGATTTGTATATA
>MinSyn_1965|Strength:0.003918067
GCGTGTCGTTTTAGTGAGGCTGACGTAAGGGATGACGCACATCTACAAAACTGGTACTTGTGTATGGT
GGAGCACGACACTACAGGAAAAAGAAGAGGTGCCCAATTAGGTTGTCATCCTTACCGCTATGGGTAAG
ATTTCTCAGCCACTTGTGTGCTCAGTGGTCCCTCCACTCTGCACCTCAATTTCGGGAAACCTCCTCGT
GTACAGGTGGGAGCCACCAAGGGCTCACTGAAGATAAGATAATAATGTTGAAGATAAGATCTACCTAT
ATAAGGTTTTGCTATTCATTGAAAGCAGTAGTGACTGATTTGTATATA
>MinSyn_1440|Strength:0.003920533

GCGTGTCGTTTTAGTGAGGTCTCTCTGCCGACAGTGGTCCCAAAGGCTATCAGCTTTGTCAAAAGCTA
AAAAAGATGATGCGGTAGGTCATTTTCAACAACTTGTGTACAGGGCTCACTGCTAGTCACTATCAGCT
GCTAGGAGGACCGATGCTGATCTTATGACGTAAGCCATGACGTCTATATCAGCTTAGCAAGACCTCTA
CAAAACATCGAAAGGACAGTATTAGGTTGTCTGCTGAAGATAAGATAATAATGTTGAAGATAAGAGTA
CTTGTGTACAGTCCATCAACAAATAATCCAAGTAAGTGTACAGGGCTCACTGCTAGGAGGACCGTGGT
GGAGCACGACACCGATGCTGATCTTGCCTGCCTTGATGCTATATAAGGTTTTGCTATTCATTGAAAGC
AGTAGTGACTGATTTGTATATA
>MinSyn_1350|Strength:0.003962101
GCGTGTCGTTTTAGTGAGGAATTTCGGGAAACCTCCTCGCGACCACTATGCCCAATTAGGTTGTAACC
ACGTCTACAAACATGTAGGCTATCAGCTTAGCCTGACGTAAGGGATGACGCACACTCACATGTAGGCT
AGCTTTGTCAAAAGCTAAAAAAGATGATGCAGCAAAACCATTATTGCGCTATGCCCAATTAGGTTGTC
TGCACTCTCTCTGCCGACAGTGGTCCCAAACGCGGTAGGTCACGACCACTATGCCTCAGAAGATCAAA
GGGCTATTGTCTGCACCTCAGTGGTCCCTCCACGGTAGGTCACGACCACTATGCCCACTATATAAGGT
TTTGCTATTCATTGAAAGCAGTAGTGACTGATTTGTATATA
>MinSyn_1453|Strength:0.004008575
GCGTGTCGTTTTAGTGAGGATGACGTAAGCCATGACGTCTAGGAGGACCGATGCACACCAGCATGTGT
TGATCACCAGCTACCTCTACAAAACTGGTACTTGTGTCACTATCAGCCACGACCACTATGCCCCAGCC
ACTTGTGTCTAGGTGGCTCCTACTGTAGGCTATCAGCTTAGCAAGACCTCTACTTTTCAACAAATGCC
CAATTAGGTTGTCTGCATGAAGATAAGATAATAATGTTGAAGATAAGATCACGACCACTCTATATAAG
GTTTTGCTATTCATTGAAAGCAGTAGTGACTGATTTGTATATA
>MinSyn_1402|Strength:0.00408242
GCGTGTCGTTTTAGTGAGGCTGACGTAAGGGATGACGCACAGCTATCAGCTTAGCAAGATCTCTCTGC
CGACAGTGGTCCCAAATGGCACACCAGCATGTGTTGATCACCAGCTTCTTGCCCAGCCACTTGTGTTG
CCCAATTAGGTTGTCTGCACCTTATGACCCCCGCCGATGACGCGGGAAAGACCTCTACAAAACTAACC
ATTATTGCGGCTTAGCAATCACTATCAGCGTCACGACCACTATGCCCAATTAGGTTGTCTATATAAGG
TTTTGCTATTCATTGAAAGCAGTAGTGACTGATTTGTATATA
>MinSyn_1922|Strength:0.00411463
GCGTGTCGTTTTAGTGAGGCTGACGTAAGGGATGACGCACAGGACCGATGCTGATCTTGCCTGCCTGC
ACACCAGCATGTGTTGATCACCAGCTTTAGTCACTATCAGCGAGGACCGATGCTGATCTTGCCTTATG
ACCCCCGCCGATGACGCGGGACCTCATCCTTACCGCTATGGGTAAGATTCACTGCATTGCGATAAAGG
AAAGGTATCAGCTTAGCAAGAACCATTATTGCGCACATGTAGGCTATCAGCTTAGCAACTATATAAGG
TTTTGCTATTCATTGAAAGCAGTAGTGACTGATTTGTATATA
>MinSyn_1208|Strength:0.004189436
GCGTGTCGTTTTAGTGAGGCAGCCACTTGTGTGGTTGTCTGCACCTCACATGTAGGCTATCAATGACG
TAAGCCATGACGTCTACAGGGCTCACTGCTAGGAGGACCGATGGGAAAAAGAAGAGGTCTTGTGTACA
GTGGGAGCCACCATGCACCTCACATGTAGGCTATGAAGATAAGATAATAATGTTGAAGATAAGACACT
GCTAGGAGGACCGATGCTGAAGCATCTTCCCTGATCTTGCCTGCCTTGAACCATTATTGCGCGGTAGG
TCACGTCACTATCAGCCCTCTACTATATAAGGTTTTGCTATTCATTGAAAGCAGTAGTGACTGATTTG
TATATA
>MinSyn_1914|Strength:0.004246368
GCGTGTCGTTTTAGTGAGGATGACGTAAGCCATGACGTCTATGTGTACAGGGCAGCAAGTGGATGCTC
ACTGCTAGGAGGACCGATGCAAAAATGTCAAAGATACATGTAGGCTATCAATCGAAAGGACAGTAGCT
TAGCAAGACCTTGAAGCATCTTCCCATGTAGGGGAAAAAGAAGAGGTCTCACATGTAGGCTTCCATCA
ACAAATAATCCAAGTAAGCCTGCCTTGATGTTATGACCCCCGCCGATGACGCGGGAGCTATATAAGGT
TTTGCTATTCATTGAAAGCAGTAGTGACTGATTTGTATATA
>MinSyn_1356|Strength:0.004290488
GCGTGTCGTTTTAGTGAGGTGGTGGAGCACGACACCCACAAATATTTCTTGTTTAGCAAGACCTCTAC
AAAACTCTGACGTAAGGGATGACGCACACTACAAAACTGGTACTTGTGTACAGGGTGAAGCATCTTCC
CAGCTTAGTGGGAGCCACCATACTTGTGTACAGGGCTCACTGCTAGGTGGCTCCTACCAAAACTGGTA
CTCAGTGGTCCCTCCACTCTACAAAACTGCTTTGTCAAAAGCTAAAAAAGATGATGCGCCTGCCTTTT
TCAACAATCTGCACCTCACATGTAGGCTATCTATATAAGGTTTTGCTATTCATTGAAAGCAGTAGTGA
CTGATTTGTATATA
>MinSyn_1195|Strength:0.00429256
GCGTGTCGTTTTAGTGAGGCTGACGTAAGGGATGACGCACAGAGGACGCTTTGTCAAAAGCTAAAAAA

GATGATGCACCTCACATTGGTGGAGCACGACATACAAAACTGGCAGCCACTTGTGTCTGCACCTCACA
TGTAGGCTATCAGGGAAAAAGAAGAGGTTGCCTTTATGACCCCCGCCGATGACGCGGGAAGACCTCTA
CAAAACTGGTATCGAAAGGACAGTAAAAACTGGTACTTGTGTACAGGGCTCACTCTATATAAGGTTTT
GCTATTCATTGAAAGCAGTAGTGACTGATTTGTATATA
>MinSyn_1674|Strength:0.004371034
GCGTGTCGTTTTAGTGAGGCAAATATTTCTTGTGTACTTGTGTACAGGGCCTGACGTAAGGGATGACG
CACAGGTAGGTCACGACCACTATGCCCAATAACCATTATTGCGGGTTCCATCAACAAATAATCCAAGT
AAGAGGTTGTCTGCACCTCACATGTAGGCTATGGAAAAAGAAGAGGTAATTAGGTTGTCTGCACCTCA
CATGTAAGCAAGTGGATGACCTTTTCAACAACACTGCTAGGAGGACGCTTTGTCAAAAGCTAAAAAAG
ATGATGCCGCCTATATAAGGTTTTGCTATTCATTGAAAGCAGTAGTGACTGATTTGTATATA
>MinSyn_1621|Strength:0.00452854
GCGTGTCGTTTTAGTGAGGTGAAGCATCTTCCTAGGTCACGACCACTATGCCCAATTAGCAAGTGGAT
GCAAGACTGGTGGAGCACGACAGGACCGATGCTGATCTTGCCTGCCTTGAATGACGTAAGCCATGACG
TCTAGGGCTCACTGCTAGGAGGACGCACACCAGCATGTGTTGATCACCAGCTCTATGCCCAATTAGGT
TGTCTGCACCGTGGGAGCCACCAGGGCTCACTGCTAGGAGGACCGATGCATCGAAAGGACAGTACGGT
AGGTCACGACCAAAAAATGTCAAAGATAGGTTCACTATCAGCTAGGTTGTCTGCACCTCCTATATAAG
GTTTTGCTATTCATTGAAAGCAGTAGTGACTGATTTGTATATA
>MinSyn_162|Strength:0.004573168
GCGTGTCGTTTTAGTGAGGTTATGACCCCCGCCGATGACGCGGGAACATGTAGGCTATCAGCTTAATT
GCGATAAAGGAAAGGACCGATGCTGATCTTGCCTGACGTAAGGGATGACGCACAGCTGATCTTGCCTG
CCTTGATGATAAGGTGGCTCCTACTATCAGGCTTTGTCAAAAGCTAAAAAAGATGATGCAAGACCTCT
ACAAAACTGGCAGCCACTTGTGTGCTTAGCAGGAAAAAGAAGAGGTCTATCAGCTTAAACCATTATTG
CGGTTGTCTGCACCTCACATGTAGGCTATCAGAGCAAGTGGATTACTATATAAGGTTTTGCTATTCAT
TGAAAGCAGTAGTGACTGATTTGTATATA
>MinSyn_1683|Strength:0.00458846
GCGTGTCGTTTTAGTGAGGTCAGAAGATCAAAGGGCTAGACCACTATGCCCAATTAGGTTGTAGCAAG
TGGATTAGGTCACGCACACCAGCATGTGTTGATCACCAGCTTTAGGTTGTCTGCACCTCACATGTAGG
CATGACGTAAGCCATGACGTCTAAAGACCTCTACAAAACTGCAGTGGTCCCTCCACGCCTGCCAAGAT
TGATGAAAAGTCAAAAACAAAAATCAATTATCCTCACATGATTGCGATAAAGGAAAGGATGCCCAATT
AGGTTGTTATGACCCCCGCCGATGACGCGGGAAGGAGGACCGAACCATTATTGCGCTACAAAACTGGT
ACTTGTGCTATATAAGGTTTTGCTATTCATTGAAAGCAGTAGTGACTGATTTGTATATA
>MinSyn_1793|Strength:0.004607228
GCGTGTCGTTTTAGTGAGGGTGGGAGCCACCACACCTCACATGTAGGCTATCAGCTTAGCAGCACACC
AGCATGTGTTGATCACCAGCTCTATCAGCTTAGCAAGACCTCTACAAAACTATCGAAAGGACAGTATG
CTGATCTTGCCTGCCTTGATGATCTGACGTAAGGGATGACGCACAATGTAGGCTTTGTCAAAAGCTAA
AAAAGATGATGCCTGCACCAATTTCGGGAAACCTCCTCGCAAACCATTATTGCGAGCAAGACCTCTAC
AAAACTGATTGCGATAAAGGAAAGGACCTCTACAAAACTGGTACTTGTGTAAAAATGTCAAAGATACT
GATCTTGCCTGCCTTGATGATACTATATAAGGTTTTGCTATTCATTGAAAGCAGTAGTGACTGATTTG
TATATA
>MinSyn_1166|Strength:0.00464948
GCGTGTCGTTTTAGTGAGGATCCTTACCGCTATGGGTAAGATTTCACTGCTAGGAGGACCGATGCTGA
TCTTAATTTCGGGAAACCTCCTCGACTGGTACTTGTATGACGTAAGCCATGACGTCTATCACATGTAG
GCTATCAGCTTAGCAAGACTTTTCAACAATAGCAAGACCTCTACAAAACTGATTGCGATAAAGGAAAG
GCCTCACATGTAGGCTATCAGCTTAGCAAACCACGTCTACAATATGCCCAATTAGGTCAGCCACTTGT
GTGGCTCACTGCTAGGAGGACCGAGTGGGAGCCACCATCTTGCCTCTATATAAGGTTTTGCTATTCAT
TGAAAGCAGTAGTGACTGATTTGTATATA
>MinSyn_1730|Strength:0.004653749
GCGTGTCGTTTTAGTGAGGCAGCCACTTGTGTGGCTCACATCGAAAGGACAGTAACATGACGTAAGCC
ATGACGTCTATAAACCACGTCTACAATACAAAACTGGTACTTGTGTACAGGTGAAGATAAGATAATAA
TGTTGAAGATAAGAAGTGAAGCATCTTCCCTATGCCCAATTAGCAAATATTTCTTGTTTGTCTGCACC
TCACATGTAGGCTATCAGCAGCAAGTGGATCAAAAAAAATGTCAAAGATATTAGCAAGACCTCTACAA
AACTGGTACCTATATAAGGTTTTGCTATTCATTGAAAGCAGTAGTGACTGATTTGTATATA
>MinSyn_1672|Strength:0.004717724
GCGTGTCGTTTTAGTGAGGCAAATATTTCTTGTACAGGGCTCACTGCTAGGAGCTGACGTAAGGGATG

ACGCACACCCAATTATCGAAAGGACAGTATCGCTTTGTCAAAAGCTAAAAAAGATGATGCACAGGGCT
CACTGCTAGGATCACTATCAGCTCAGCTTAGCAAGACCTCCAGCCACTTGTGTCTCACTGCTAGGAGG
ACCGGCACACCAGCATGTGTTGATCACCAGCTACCGATGCTGATCTTGCCTGCAGCAAGTGGATGGCT
ATCAGCCTATATAAGGTTTTGCTATTCATTGAAAGCAGTAGTGACTGATTTGTATATA
>MinSyn_1723|Strength:0.004749634
GCGTGTCGTTTTAGTGAGGCAAATATTTCTTGTAGGCTATAAGATTGATGAAAAGTCAAAAACAAAAA
TCAATTATCGCGGTAGGTCACGATGACGTAAGCCATGACGTCTAAGGAGGACCGATGCACACCAGCAT
GTGTTGATCACCAGCTCACTATGCCCAATTAGGTTGTTGGTGGAGCACGACATGGTACTTTTTTCAAC
AACCACTATGCCCAATTAGGTTGTCTGCACCAAAAATGTCAAAGATAGACCTCTACAACAGTGGTCCC
TCCACTGGTACTTGTGTACAGGGCTCACTGCTACTATATAAGGTTTTGCTATTCATTGAAAGCAGTAG
TGACTGATTTGTATATA
>MinSyn_1222|Strength:0.004783589
GCGTGTCGTTTTAGTGAGGTTATGACCCCCGCCGATGACGCGGGATGTCTGCACCTCACATGTATGAC
GTAAGCCATGACGTCTAAACTGGTGGGAGCCACCAACCACTATGCCCAAATCGAAAGGACAGTACACG
ACCACTATGCCCAATTAGGTTGTCTGATCCTTACCGCTATGGGTAAGATTGCAAGACCTCTACAAAAC
TTCCATCAACAAATAATCCAAGTAAGCGGTAGGTCACGGCTTTGTCAAAAGCTAAAAAAGATGATGCC
GATGCTGATCTTGCTATATAAGGTTTTGCTATTCATTGAAAGCAGTAGTGACTGATTTGTATATA
>MinSyn_1456|Strength:0.004789159
GCGTGTCGTTTTAGTGAGGTGGTGGAGCACGACACTCACATGTAGGCTATCAGCTTAGCCTGACGTAA
GGGATGACGCACAAAAACTGGTACTTGTGTACAGGGCTCATCTCTCTGCCGACAGTGGTCCCAAAGCA
CCTCACACAGCCACTTGTGTGGTACTTGTGTACAGGGTCCATCAACAAATAATCCAAGTAAGCTCACT
GCTAGGAGGACCGATGCTGATAATTTCGGGAAACCTCCTCGGATGCTGATAGGTGGCTCCTACACTGC
TAGGAGGACCTATATAAGGTTTTGCTATTCATTGAAAGCAGTAGTGACTGATTTGTATATA
>MinSyn_1165|Strength:0.004858978
GCGTGTCGTTTTAGTGAGGCTGACGTAAGGGATGACGCACAAATTAGGTTGTCTGCCAAATATTTCTT
GTATGCTGATCTTGCCTGCCTTGAATCGAAAGGACAGTAAAGACCTCAGCAAGTGGATGGAATCCTTA
CCGCTATGGGTAAGATTTCACGACCACTATGCCCAATTAGGTTGTCTTGAAGCATCTTCCTGTACAGG
GCTCACTGCTAGGAGGACCTGGTGGAGCACGACAACATGTAGGCCTATATAAGGTTTTGCTATTCATT
GAAAGCAGTAGTGACTGATTTGTATATA
>MinSyn_132|Strength:0.005164033
GCGTGTCGTTTTAGTGAGGGCTTTGTCAAAAGCTAAAAAAGATGATGCCGACCACTATGCAGCCACTT
GTGTCTAGGAGGACCGATGCTGATCTTTCACTATCAGCTGCCCAATTAGGTTGTCTCTGACGTAAGGG
ATGACGCACAAGCAAGACCTCTACAAAACTGGTACTTGTGAACCACGTCTACAATCAGCTTAGCAAGA
CCTCTACAAAACTGGAGCAAGTGGATGATCTTATTGCGATAAAGGAAAGGCTACAAAACTGGTGAAGA
TAAGATAATAATGTTGAAGATAAGAGCCCAATTAGGTTGTCTGCACCTCACCTATATAAGGTTTTGCT
ATTCATTGAAAGCAGTAGTGACTGATTTGTATATA
>MinSyn_1258|Strength:0.005275893
GCGTGTCGTTTTAGTGAGGATTGCGATAAAGGAAAGGCTTCTCTCTGCCGACAGTGGTCCCAAATTGC
CCTGACGTAAGGGATGACGCACATACAGGGCTCACAGGTGGCTCCTACCGTGAAGCATCTTCCGACCT
CTACAAAACTATCCTTACCGCTATGGGTAAGATTGTACTTGTGTACAGGGCTCACTGCTGTGGGAGCC
ACCACTTGTGTACAGGCAGTGGTCCCTCCACGAGGACCGATGCTGATTTTTCAACAACTAGGAGGACC
GATGCTGACTATATAAGGTTTTGCTATTCATTGAAAGCAGTAGTGACTGATTTGTATATA
>MinSyn_1891|Strength:0.005324217
GCGTGTCGTTTTAGTGAGGTTTTCAACAATGATTGCGATAAAGGAAAGGCTTGTGTACAGGGCTCACT
GCTAATGACGTAAGCCATGACGTCTAGCTAGGAGGACCGATGCTGAGGTGGCTCCTACGCACCTCACA
TGTAGGCTATCCAAATATTTCTTGTTGCCCAATTTCACTATCAGCACTGGTACTTGTGTACAGGGCTC
AGAAGATCAAAGGGCTACATGTAGGCTATCAGCTTAGCAAGACCTTGGTGGAGCACGACAGTCTGCAC
CTCACATGCTATATAAGGTTTTGCTATTCATTGAAAGCAGTAGTGACTGATTTGTATATA
>MinSyn_1838|Strength:0.005331676
GCGTGTCGTTTTAGTGAGGAACCATTATTGCGCAGCTTAGCAAGACCTCTACAAAACTGGTATCCATC
AACAAATAATCCAAGTAAGAGGAGGAGGTGGCTCCTACTCGCGGTCTGACGTAAGGGATGACGCACAC
CGATGCTGATCTTGCCAACCACGTCTACAATGTACAGGGCTCACTGCTAGGAGGTGAAGCATCTTCCT
TAGCAAGACCTCTACAAAAATTTCGGGAAACCTCCTCGTGCCCAATTAGGTTGTCTTCAGAAGATCAA
AGGGCTACCTCTACAAAGCAAGTGGATCTACAAAACTGGTCTATATAAGGTTTTGCTATTCATTGAAA

GCAGTAGTGACTGATTTGTATATA
>MinSyn_1244|Strength:0.005358282
GCGTGTCGTTTTAGTGAGGCAGCCACTTGTGTGCTGATCTTGCCTGCCTTAGCAAGTGGATGGCTATC
AGCTTAGCAAGACCTCACTATCAGCTTAGGTTGTCTGCACCTCACAACCATTATTGCGCTGCACCTCA
CATGTAGGCTATCAGCTTAGCTGACGTAAGGGATGACGCACACACATGTAGGCTGAAGATAAGATAAT
AATGTTGAAGATAAGACTCACTGCTAGGAGGACCGATGCTGATGGTGGAGCACGACATACAAAACTGG
TACTTGTGTACAGGGCGTGGGAGCCACCATAGGTTGTCTGCACCTCACAATTGCGATAAAGGAAAGGT
GCCCTATATAAGGTTTTGCTATTCATTGAAAGCAGTAGTGACTGATTTGTATATA
>MinSyn_1570|Strength:0.005399062
GCGTGTCGTTTTAGTGAGGATCCTTACCGCTATGGGTAAGATTTATTGCGATAAAGGAAAGGGTAGGC
TATCAGCTTCTGACGTAAGGGATGACGCACAGTACAGGGCTCACTGCTAGGAGGAAACCATTATTGCG
GCTATCAGCTTAGCAAGACCTCTAAAGATTGATGAAAAGTCAAAAACAAAAATCAATTATTATGCCCA
ATTAGGTGCTTTGTCAAAAGCTAAAAAAGATGATGCCCCAATTAGGTTGTCTGCCAGTGGTCCCTCCA
CACGACCACTATGCTATATAAGGTTTTGCTATTCATTGAAAGCAGTAGTGACTGATTTGTATATA
>MinSyn_1319|Strength:0.005466357
GCGTGTCGTTTTAGTGAGGTCCATCAACAAATAATCCAAGTAAGATGCTGATCTTGCCTGCCTTGAGC
ACACCAGCATGTGTTGATCACCAGCTCCCAATTAGGTTGTCTGCACCTCACATGATGACGTAAGCCAT
GACGTCTAGAATTTCGGGAAACCTCCTCGCTCACATGTAGGCTATCAGCTTAGCAAGCAGTGGTCCCT
CCACCTGTCAGAAGATCAAAGGGCTAACTGCTAGGAGGAACCACGTCTACAAATGCTGATCTTGCCTG
CAAAAATGTCAAAGATAAGCAAGACCTCTACAAAACTGGTACTTCTATATAAGGTTTTGCTATTCATT
GAAAGCAGTAGTGACTGATTTGTATATA
>MinSyn_1901|Strength:0.005581576
GCGTGTCGTTTTAGTGAGGATGACGTAAGCCATGACGTCTACACCTCACATGTAGGCTATCAAAAATG
TCAAAGATACCTCACATGTAGGCTATCAATCGAAAGGACAGTATACAGTCTCTCTGCCGACAGTGGTC
CCAAAGGAGGACCGCACACCAGCATGTGTTGATCACCAGCTTAGCAAGACCTCTGCTTTGTCAAAAGC
TAAAAAAGATGATGCGATCTTGCCTGCCTTGATGACTATATAAGGTTTTGCTATTCATTGAAAGCAGT
AGTGACTGATTTGTATATA
>MinSyn_1861|Strength:0.005645225
GCGTGTCGTTTTAGTGAGGGGAAAAAGAAGAGGTTTGTCTGCACCTCACACTGACGTAAGGGATGACG
CACACCCAAGGTGGCTCCTACTCTACAAAACTGGTACTTGTGTACAGGGAGCAAGTGGATAAGACCTC
TACAAAACTGGAATTTCGGGAAACCTCCTCGTATGCCCAATTAGGTTGTCTGCACCTCAATCCTTACC
GCTATGGGTAAGATTCGATGCTGATCTTTTCAACAAGTACAGGGCTCACTGCTAGCTATATAAGGTTT
TGCTATTCATTGAAAGCAGTAGTGACTGATTTGTATATA
>MinSyn_1605|Strength:0.005934536
GCGTGTCGTTTTAGTGAGGTCCATCAACAAATAATCCAAGTAAGGCTAGGAGGACCGATGCTGATCTT
GCCTGCAATTTCGGGAAACCTCCTCGGTCACGACCACTATGCCCAATTAGTCACTATCAGCACATGTA
GCTGACGTAAGGGATGACGCACACTTGCCTGCCTTGATCGAAAGGACAGTAGATGCTTTGTCAAAAGC
TAAAAAAGATGATGCTCACTGCTAGGAGGACCGATGCTGATCTTGTTATGACCCCCGCCGATGACGCG
GGAACTTGAAGCATCTTCCCTATCAGCTTAGCAAGACCTCTACAAAACTATATAAGGTTTTGCTATTC
ATTGAAAGCAGTAGTGACTGATTTGTATATA
>MinSyn_1805|Strength:0.005937835
GCGTGTCGTTTTAGTGAGGTCTCTCTGCCGACAGTGGTCCCAAATACAGGGCTCACTGCTAGGAGGAC
CGATGTTTTCAACAATAGGAGATGACGTAAGCCATGACGTCTAGCGGTAGGAACCATTATTGCGATCA
GCAGTGGTCCCTCCACTCTACAAAACTGGTACTTGTGATCCTTACCGCTATGGGTAAGATTTAGGTCA
TCGAAAGGACAGTAGAATTTCGGGAAACCTCCTCGCACATGTAGGCTATCAGCTTAGAAAAATGTCAA
AGATACACATGTACTATATAAGGTTTTGCTATTCATTGAAAGCAGTAGTGACTGATTTGTATATA
>MinSyn_1465|Strength:0.005987796
GCGTGTCGTTTTAGTGAGGAACCATTATTGCGAGGACCGATGTCCATCAACAAATAATCCAAGTAAGT
AGCAAGACCTCTACAAAACTGGTGGTGGAGCACGACAACTGGTACTTGTGTACAGGGGAAAAAGAAGA
GGTCACTGCTAGGAGGACATGACGTAAGCCATGACGTCTATTTGAAGATAAGATAATAATGTTGAAGA
TAAGAGACATTGCGATAAAGGAAAGGGCTAGGAGGACCGATAATTTCGGGAAACCTCCTCGCAGCTTA
GCAAGACCTCTACAAAACAACCACGTCTACAAGCTTAGCAAGACCTCTACAAAACTGGCTATATAAGG
TTTTGCTATTCATTGAAAGCAGTAGTGACTGATTTGTATATA
>MinSyn_1785|Strength:0.006136774

GCGTGTCGTTTTAGTGAGGATGACGTAAGCCATGACGTCTATGAAACCACGTCTACAAGGGCTCACTG
CTAGGAGGACTCCATCAACAAATAATCCAAGTAAGTGTAGGCTATCAGCAGCAAGTGGATCGCACACC
AGCATGTGTTGATCACCAGCTCTCACTATCAGCGATCTTGCCTGGTGGGAGCCACCATCTGCACCTCA
CATGTAGGAACCATTATTGCGTGCCTGCTATATAAGGTTTTGCTATTCATTGAAAGCAGTAGTGACTG
ATTTGTATATA
>MinSyn_1773|Strength:0.006230396
GCGTGTCGTTTTAGTGAGGTTTTCAACAACGCGGTAGGTCACGACCACTATGCCCAATATCGAAAGGA
CAGTAAGGAGGACCGATGCTGATCTTGCCTGCTCCATCAACAAATAATCCAAGTAAGGGAGGACCGAT
GCTGATCTTGCCTGCCGCTTTGTCAAAAGCTAAAAAAGATGATGCCCTCACATGTAAGCAAGTGGATG
CTTAGCAAGACCTCTACAAAACTGGTACCAGTGGTCCCTCCACACCGATGCTGATCTTGCCTGCCTTG
ACTGACGTAAGGGATGACGCACACACGACCACTATGCCCAATTAGGTTGTCATTGCGATAAAGGAAAG
GTCACGACCACTATGCCCAATTAGGTTGAAGCATCTTCCTGTCTGCACCTCACATGTAGGCTATCAGC
CTATATAAGGTTTTGCTATTCATTGAAAGCAGTAGTGACTGATTTGTATATA
>MinSyn_1302|Strength:0.00639162
GCGTGTCGTTTTAGTGAGGCAAATATTTCTTGTCATTGCGATAAAGGAAAGGCTGCTAGGAGGACGCT
TTGTCAAAAGCTAAAAAAGATGATGCGCTAGGAGGACCGATCAGCCACTTGTGTCCTGACGTAAGGGA
TGACGCACAGTTGTCTGCACCTCACATGTAGGCTTGGTGGAGCACGACAAAAACTGGTACTTGTGTAC
AGGGCTCACTATCCTTACCGCTATGGGTAAGATTACATGTAGGCTATTCACTATCAGCACAGGGCTCA
CTGAGCAAGTGGATACGACCACTATGCCCTATATAAGGTTTTGCTATTCATTGAAAGCAGTAGTGACT
GATTTGTATATA
>MinSyn_1481|Strength:0.006415017
GCGTGTCGTTTTAGTGAGGAACCACGTCTACAAATTCCATCAACAAATAATCCAAGTAAGAATCAAAT
ATTTCTTGTTTGTATGACGTAAGCCATGACGTCTAGCTCACTGCTAGGAGGAAGATTGATGAAAAGTC
AAAAACAAAAATCAATTATTGTACAGGGCTCACTGCTAGGAAAAAGAAGAGGTCATGTAGGCTATCAG
CTTAAATTTCGGGAAACCTCCTCGGCTTAAAAATGTCAAAGATATGTGTACAGGGCTCACTGCTAGGC
TATATAAGGTTTTGCTATTCATTGAAAGCAGTAGTGACTGATTTGTATATA
>MinSyn_1782|Strength:0.006521126
GCGTGTCGTTTTAGTGAGGCAGCCACTTGTGTGTAGGCTATCAGCTTAGCAAGACCTCTATGAAGCAT
CTTCCAAAACTATGACGTAAGCCATGACGTCTATACAAAACTGGTACTTGTGTACAGGGCTCAATTTC
GGGAAACCTCCTCGCTGAAGATAAGATAATAATGTTGAAGATAAGAATGTAGGCTATCATCACTATCA
GCTTAGGTTGTCTGCACCTCACATAGCAAGTGGATCCTCTACAAAACTGGTACTTGTGTACACTATAT
AAGGTTTTGCTATTCATTGAAAGCAGTAGTGACTGATTTGTATATA
>MinSyn_1688|Strength:0.006538338
GCGTGTCGTTTTAGTGAGGGCACACCAGCATGTGTTGATCACCAGCTTCTTATGACCCCCGCCGATGA
CGCGGGAAGGAGGACCGATGCGGAAAAAGAAGAGGTGAGGACCGATGCTGATCTTGCCTCCATCAACA
AATAATCCAAGTAAGTGTGTACAGGGCTCACTGCTGACGTAAGGGATGACGCACAAAGACCTCTACAA
AAAACCACGTCTACAAAATTAGCAGTGGTCCCTCCACGCTATCAGCTTAGCAAGACCTATCCTTACCG
CTATGGGTAAGATTCTCTAAAAAATGTCAAAGATAGCACCTCACATGTAGGCTATCAGCCTATATAAG
GTTTTGCTATTCATTGAAAGCAGTAGTGACTGATTTGTATATA
>MinSyn_1836|Strength:0.006558965
GCGTGTCGTTTTAGTGAGGCTGACGTAAGGGATGACGCACAAACAGCAAGTGGATACTGGTACTTGTG
TACAGGGCTCACTTATGACCCCCGCCGATGACGCGGGAGCTAGGAGGACCGGCACACCAGCATGTGTT
GATCACCAGCTTCACGACCACTGCTTTGTCAAAAGCTAAAAAAGATGATGCTCGCGGTAGGTCACGAC
CTTTTCAACAAGACCTCTACACTATATAAGGTTTTGCTATTCATTGAAAGCAGTAGTGACTGATTTGT
ATATA
>MinSyn_1866|Strength:0.006626829
GCGTGTCGTTTTAGTGAGGTGAAGATAAGATAATAATGTTGAAGATAAGAGACCGATGCTGATCTTGC
CTTCTCTCTGCCGACAGTGGTCCCAAAGGCTATCAGCTTAGCAAGACCATCCTTACCGCTATGGGTAA
GATTGGCTCACTGCTAGGAGGACCGATGCTGAGCTTTGTCAAAAGCTAAAAAAGATGATGCCGACCAC
TATGCCCAATATGACGTAAGCCATGACGTCTATGCCTGAGGTGGCTCCTACTCAGCTTAGCAAGACCT
CTTGAAGCATCTTCCCTATGCCCAATTAGGTTGTCTGCAAATATTTCTTGTACAGGGCTCACTGCTAG
GAGAGCAAGTGGATCTCTACCTATATAAGGTTTTGCTATTCATTGAAAGCAGTAGTGACTGATTTGTA
TATA
>MinSyn_1511|Strength:0.006788753

GCGTGTCGTTTTAGTGAGGATTGCGATAAAGGAAAGGCACAATTTCGGGAAACCTCCTCGCTATGCCC
AATTAGGTTGTCTGCACCTCATGAAGCATCTTCCCAAAACTGGTACTTGTGTACAGGGCTCACTTGAA
GATAAGATAATAATGTTGAAGATAAGAAGGGCTCACTGCTAGGAGGGTGGGAGCCACCATAGGTTGTC
TGCACCTCATGACGTAAGCCATGACGTCTAAGCAAGACCTCTACAAAAACCACGTCTACAAGACCTCT
ACAAAACTGGTACTTGTGTTCAGAAGATCAAAGGGCTAGTCACGACCACTATGCCCAAAAAATGTCAA
AGATAGCGGTAGGTCCTATATAAGGTTTTGCTATTCATTGAAAGCAGTAGTGACTGATTTGTATATA
>MinSyn_1865|Strength:0.006836281
GCGTGTCGTTTTAGTGAGGAAGATTGATGAAAAGTCAAAAACAAAAATCAATTATTAGGTTGAAGATA
AGATAATAATGTTGAAGATAAGAAGGACCGATCAGTGGTCCCTCCACCCTCACATGTAGGCTATCAGC
TTAGCAAGCACACCAGCATGTGTTGATCACCAGCTACATGTAGGCTGAAGCATCTTCCCTGCCTTGAT
GATATGACGTAAGCCATGACGTCTATTGTCTGCACCTGCTTTGTCAAAAGCTAAAAAAGATGATGCTG
ATCTTTCTCTCTGCCGACAGTGGTCCCAAAACCTCTACAAAACTGGTACTTGTCAGAAGATCAAAGGG
CTAAATTAGGTTGCTATATAAGGTTTTGCTATTCATTGAAAGCAGTAGTGACTGATTTGTATATA
>MinSyn_1995|Strength:0.006870516
GCGTGTCGTTTTAGTGAGGCTGACGTAAGGGATGACGCACACACCTCACATGTAGGCTATCATGAAGC
ATCTTCCGTGTACAGGGCTCACTGCTAGGAATCCTTACCGCTATGGGTAAGATTCACAAGATTGATGA
AAAGTCAAAAACAAAAATCAATTATTGGTACTTGTGTACAGGGTCAGAAGATCAAAGGGCTAACCTCA
TTTTCAACAATAGGAGGCTATATAAGGTTTTGCTATTCATTGAAAGCAGTAGTGACTGATTTGTATAT
A
>MinSyn_1758|Strength:0.007020013
GCGTGTCGTTTTAGTGAGGTTATGACCCCCGCCGATGACGCGGGAGTAGGCTATCAGCTTAGCAAGAC
AAAAATGTCAAAGATACTGATCTTGCCTGCCGCTTTGTCAAAAGCTAAAAAAGATGATGCCCTGCCTT
GATGAAATTTCGGGAAACCTCCTCGACGACCACTATGCCCAATTAGGTTATGACGTAAGCCATGACGT
CTAGCCTTGAATTGCGATAAAGGAAAGGCTGCACCTCACATGTAGGCTATCAGCTGGTGGAGCACGAC
AATCTTGCCTGCCTTGTGAAGATAAGATAATAATGTTGAAGATAAGACTATCAGCTTAGCACTATATA
AGGTTTTGCTATTCATTGAAAGCAGTAGTGACTGATTTGTATATA
>MinSyn_1972|Strength:0.007022027
GCGTGTCGTTTTAGTGAGGTGAAGATAAGATAATAATGTTGAAGATAAGACAAAGATTGATGAAAAGT
CAAAAACAAAAATCAATTATTTAGGTTGTCTGCACCTCACCTGACGTAAGGGATGACGCACAACATGT
AGGCTATCAATCCTTACCGCTATGGGTAAGATTTCATCGAAAGGACAGTACCTCTACAAAACTGGTAC
TTGTGTACACAGCCACTTGTGTCACATGTAGGCTATAACCATTATTGCGCCTCTACAAAACTGGTACT
TGTGTACCTATATAAGGTTTTGCTATTCATTGAAAGCAGTAGTGACTGATTTGTATATA
>MinSyn_113|Strength:0.007030368
GCGTGTCGTTTTAGTGAGGGGAAAAAGAAGAGGTTGTGTACAGGGCTCACTGCTAGATGAGCTAAGCA
CATACGTCAGGTACTTGTGTACAGGGCTCACTTTATGACCCCCGCCGATGACGCGGGACAGCTTAGCA
AATCGAAAGGACAGTAAAAACTGGTACTTGTGTACAGGGCTCACTTCTCTCTGCCGACAGTGGTCCCA
AACACTGCTAGGAGGACCGATGCTGATCTTGGCACACCAGCATGTGTTGATCACCAGCTGGACCGATG
CTGACAGTGGTCCCTCCACGCCCAATTAGGTTGTCTGCACCTCACATGTGTGGGAGCCACCACTTGTG
TACAGGGCTCCATCAACAAATAATCCAAGTAAGTATCAGCTTAGCAAGACCTCTACCTATATAAGGTT
TTGCTATTCATTGAAAGCAGTAGTGACTGATTTGTATATA
>MinSyn_1813|Strength:0.007141751
GCGTGTCGTTTTAGTGAGGATTGCGATAAAGGAAAGGTGCTGCACACCAGCATGTGTTGATCACCAGC
TTCACTGCTAGGAGGACCGATGCTGATCTTATGACGTAAGCCATGACGTCTAATTAGGTTGTCTGCAC
CTCACATGTAGGCTGAAGCATCTTCCTACTTGTGTACAGGGCTCACTGCTCCATCAACAAATAATCCA
AGTAAGAATTAGGTTGTCTGCACCTCACATGTACAGCCACTTGTGTATCTTGCCTGCCTTGATGACTA
TATAAGGTTTTGCTATTCATTGAAAGCAGTAGTGACTGATTTGTATATA
>MinSyn_1190|Strength:0.007358442
GCGTGTCGTTTTAGTGAGGGCTTTGTCAAAAGCTAAAAAAGATGATGCGGCTATCAGCTTAGCAAGAC
CTCCAGTGGTCCCTCCACTCACGACCAACCATTATTGCGTCACGACCACTATGCCCAATTAGGTTGTC
TCTGACGTAAGGGATGACGCACACTGCATCCTTACCGCTATGGGTAAGATTTTGTGTACAGGGCTCAC
TGTCACTATCAGCCTCACTGCTAGGAGGACCGATGCTGATCCATCAACAAATAATCCAAGTAAGTGTG
TACAGGGCTCACTGCTAGGAGCTATATAAGGTTTTGCTATTCATTGAAAGCAGTAGTGACTGATTTGT
ATATA
>MinSyn_1761|Strength:0.007428958

GCGTGTCGTTTTAGTGAGGTTATGACCCCCGCCGATGACGCGGGAACCTCACTGACGTAAGGGATGAC
GCACACTCACATGTAGGCTATCAGCTTAGCGGAAAAAGAAGAGGTATCAGCTTAGCAAGACCTCTACA
AACCATTATTGCGCTGGTGGAGCACGACAGACCTCTACAAAACTGGTACTTGTCACTATCAGCTTAGG
TTGTCTGCAAACCACGTCTACAATACAAAACTATATAAGGTTTTGCTATTCATTGAAAGCAGTAGTGA
CTGATTTGTATATA
>MinSyn_1521|Strength:0.007431896
GCGTGTCGTTTTAGTGAGGAAGATTGATGAAAAGTCAAAAACAAAAATCAATTATGAGGACCGATGCT
GATTCCATCAACAAATAATCCAAGTAAGGACCGATATGACGTAAGCCATGACGTCTATAGGCTATCAG
CTTAGTCACTATCAGCCTATCAGCTTAGCAAGACCTCTACAACAGCCACTTGTGTGTAGGCTATCAGC
TTAGCAAGACCTCTACAGGTGGCTCCTACACTTGTGTACAGGGCTGGAAAAAGAAGAGGTACTGGCTA
TATAAGGTTTTGCTATTCATTGAAAGCAGTAGTGACTGATTTGTATATA
>MinSyn_1800|Strength:0.007451443
GCGTGTCGTTTTAGTGAGGTGAAGATAAGATAATAATGTTGAAGATAAGAGGTAGGTCACGATGGTGG
AGCACGACAAAACTGGTACTTGTGTAAGGTGGCTCCTACTGTAGGCTATCAGCTTAGCAAGACCTCTA
AAAAATGTCAAAGATAATCAGCTTAGCAAGACCTCTACAAAATCAGAAGATCAAAGGGCTACAGCTTA
GCAAGACATGACGTAAGCCATGACGTCTACCCAATTAGGTTGTCTGCACCATCGAAAGGACAGTACTA
CAAACAGCCACTTGTGTCCAATTAGGTTTCACTATCAGCACTATGCCCAATTAGGTTGTCTGCACCTC
ACTATATAAGGTTTTGCTATTCATTGAAAGCAGTAGTGACTGATTTGTATATA
>MinSyn_1867|Strength:0.007558677
GCGTGTCGTTTTAGTGAGGTCCATCAACAAATAATCCAAGTAAGTAACCATTATTGCGCGATGCTGAT
CTTGTGAAGATAAGATAATAATGTTGAAGATAAGAATTAGGTTGTCTGCACCTCACAAAGATTGATGA
AAAGTCAAAAACAAAAATCAATTATGCTATCAGCTTAGCTGACGTAAGGGATGACGCACAGACCTCTA
CAAAACTGGTACTTGTGTACCAGCCACTTGTGTGACCAGCAAGTGGATATTAGAGGTGGCTCCTACTT
GTGTACAGGGCTCAAACCACGTCTACAAATGCCCAATTAGGTTGCTATATAAGGTTTTGCTATTCATT
GAAAGCAGTAGTGACTGATTTGTATATA
>MinSyn_1923|Strength:0.00756904
GCGTGTCGTTTTAGTGAGGTGAAGATAAGATAATAATGTTGAAGATAAGATAGGAGGACCGATGCTGC
AGCCACTTGTGTCCTCTACAAAACATCCTTACCGCTATGGGTAAGATTTCACATGTAGGCTATCAGCT
TCTGACGTAAGGGATGACGCACAACGTTATGACCCCCGCCGATGACGCGGGAGCAAGACCTCTGGTGG
AGCACGACACAATTAGGTTGTCTGCAATTGCGATAAAGGAAAGGGTAAGATTGATGAAAAGTCAAAAA
CAAAAATCAATTATTTGATGACTATATAAGGTTTTGCTATTCATTGAAAGCAGTAGTGACTGATTTGT
ATATA
>MinSyn_1988|Strength:0.00756913
GCGTGTCGTTTTAGTGAGGGCTTTGTCAAAAGCTAAAAAAGATGATGCGTTGTCTGCACCTCAAGATT
GATGAAAAGTCAAAAACAAAAATCAATTATAGGACCGATGCTGAAGGTGGCTCCTACCTGCTAGAAAA
ATGTCAAAGATATGTCTGCACCTCACATGCAAATATTTCTTGTGCAAGACCTCTACATGACGTAAGCC
ATGACGTCTAAATTAGGTTGTCTCTCTCTGCCGACAGTGGTCCCAAAATCAGCTTAGCATTATGACCC
CCGCCGATGACGCGGGACTCTACAAAACTGGTACTTGTGTATGGTGGAGCACGACATCTATATAAGGT
TTTGCTATTCATTGAAAGCAGTAGTGACTGATTTGTATATA
>MinSyn_1675|Strength:0.007628141
GCGTGTCGTTTTAGTGAGGATCGAAAGGACAGTATCTACAAAACTGGTACTTGTGTTGAAGCATCTTC
CCGGTAGGTCACGACCACTATGCCCAAAGCAAGTGGATTGTGTACAGGGCTCACTGATGACGTAAGCC
ATGACGTCTACCAATTAGGTTGTCTGCACCTAGGTGGCTCCTACTCTTGCCTGCCTTGATGATATCAG
AAGATCAAAGGGCTACGCGGTGCACACCAGCATGTGTTGATCACCAGCTGTCTGCACCTCACATGTAG
GCTATCAGCTATATAAGGTTTTGCTATTCATTGAAAGCAGTAGTGACTGATTTGTATATA
>MinSyn_154|Strength:0.007683984
GCGTGTCGTTTTAGTGAGGCTGACGTAAGGGATGACGCACAACTATGCCCAATTAGGTTGTCATCGAA
AGGACAGTACTCACTGCTTTGTCAAAAGCTAAAAAAGATGATGCAACTGGTACTTCTCTCTGCCGACA
GTGGTCCCAAATGTGTACAGGGCTCACTGCTAGGACAGCCACTTGTGTCTTGTGTACAGGGCTCACTG
CTAGGCTATATAAGGTTTTGCTATTCATTGAAAGCAGTAGTGACTGATTTGTATATA
>MinSyn_1863|Strength:0.007889615
GCGTGTCGTTTTAGTGAGGTGAAGATAAGATAATAATGTTGAAGATAAGACTATCAGCTTAGCAAGAC
CTCTACAAAATGACGTAAGCCATGACGTCTATAGCTTTGTCAAAAGCTAAAAAAGATGATGCACTGTG
GGAGCCACCACTTTATGACCCCCGCCGATGACGCGGGAAGGGCTCACTGCTAGGAAATTTCGGGAAAC

CTCCTCGGGCTATCAAGCAAGTGGATACAAAACTGGTACTTGTCTATATAAGGTTTTGCTATTCATTG
AAAGCAGTAGTGACTGATTTGTATATA
>MinSyn_112|Strength:0.007940764
GCGTGTCGTTTTAGTGAGGATCGAAAGGACAGTATACAAAACTGGTGGAAAAAGAAGAGGTAGGACCG
ATGCTGATCTTTCCATCAACAAATAATCCAAGTAAGATTAGGTTGTCTGCACCTCACTGACGTAAGGG
ATGACGCACATATCAGCTTAATTTCGGGAAACCTCCTCGGACCACTATGCCCAATTAGAGGTGGCTCC
TACTGTGTACAGGGCTCACTGCTTGAAGATAAGATAATAATGTTGAAGATAAGATGTCTGCACCTCAC
ATGTCTATATAAGGTTTTGCTATTCATTGAAAGCAGTAGTGACTGATTTGTATATA
>MinSyn_1439|Strength:0.007973784
GCGTGTCGTTTTAGTGAGGATGACGTAAGCCATGACGTCTACAAGACCTCTACAAAACTGGTACTTGT
GAGCAAGTGGATTGGTACTTGTGTACAGGCAGCCACTTGTGTAAGACCTCTACAAAACAAATATTTCT
TGTACTATGCAGTGGTCCCTCCACATTAGGTAATTTCGGGAAACCTCCTCGGTTGTCTGCACCTCACA
TGCTATATAAGGTTTTGCTATTCATTGAAAGCAGTAGTGACTGATTTGTATATA
>MinSyn_1616|Strength:0.008112032
GCGTGTCGTTTTAGTGAGGATTGCGATAAAGGAAAGGAGACCTCTATGACGTAAGCCATGACGTCTAG
CCCAATTAGGTTGTCTGCACCTCACATGCTTTGTCAAAAGCTAAAAAAGATGATGCTTAGCAAGACCT
CTACTCTCTCTGCCGACAGTGGTCCCAAATCAGCTTAGCAAGACCTCTACAAAACTATCGAAAGGACA
GTAACGACCACTATGCCCTATATAAGGTTTTGCTATTCATTGAAAGCAGTAGTGACTGATTTGTATAT
A
>MinSyn_1717|Strength:0.008168529
GCGTGTCGTTTTAGTGAGGTCCATCAACAAATAATCCAAGTAAGTACTTGTGTACAGGGCTCACTGCT
AGCTGACGTAAGGGATGACGCACATGCCTGCCTTGATGATCGAAAGGACAGTAACGACCACTATGCTT
TTCAACAATGCCTGCCTTGATGATAACCACGTCTACAACTAACCATTATTGCGGGCTATCAGCTTAGC
GCACACCAGCATGTGTTGATCACCAGCTCAGCTTCTATATAAGGTTTTGCTATTCATTGAAAGCAGTA
GTGACTGATTTGTATATA
>MinSyn_1547|Strength:0.008275314
GCGTGTCGTTTTAGTGAGGTCACTATCAGCTTGCCTGCCTTCTGACGTAAGGGATGACGCACATTGTG
TACAGGGCTCACTGCTAGGAGAAGATTGATGAAAAGTCAAAAACAAAAATCAATTATAACTGGTACTT
GTCCATCAACAAATAATCCAAGTAAGCTACAAAATGAAGATAAGATAATAATGTTGAAGATAAGAAGC
TTAGCAAGACCTCTCTATATAAGGTTTTGCTATTCATTGAAAGCAGTAGTGACTGATTTGTATATA
>MinSyn_1348|Strength:0.008308529
GCGTGTCGTTTTAGTGAGGCAAATATTTCTTGTTCTGACGTAAGGGATGACGCACAAGCTTAGCAAGA
CCTCTACAAAACTGTCCATCAACAAATAATCCAAGTAAGTCACATGTAGGCTATTGGTGGAGCACGAC
AAGATGAAGCATCTTCCGGGCTCACATCCTTACCGCTATGGGTAAGATTGACCTCTACAAAACTGGTA
CTTGTGTACTATATAAGGTTTTGCTATTCATTGAAAGCAGTAGTGACTGATTTGTATATA
>MinSyn_1140|Strength:0.008405147
GCGTGTCGTTTTAGTGAGGTGAAGATAAGATAATAATGTTGAAGATAAGACTGCTAGGAGGACCGCAG
TGGTCCCTCCACAGCTTAGCAAGACCTCTACAAAACTGGTAATGAGCTAAGCACATACGTCAGTGCTG
ATCTGGAAAAAGAAGAGGTGCTGATCTAGGTGGCTCCTACTCACATGTAGGGCACACCAGCATGTGTT
GATCACCAGCTGGTAGGTCACGACCACTATGCCCAATTAGGAACCACGTCTACAATGCCCAATTAGGT
TGTCTGCACCTCAAAGATTGATGAAAAGTCAAAAACAAAAATCAATTATGGAGGACCGATGCATTGCG
ATAAAGGAAAGGGCCCAATTAGGTTGTCTATATAAGGTTTTGCTATTCATTGAAAGCAGTAGTGACTG
ATTTGTATATA
>MinSyn_1716|Strength:0.00854693
GCGTGTCGTTTTAGTGAGGAAGATTGATGAAAAGTCAAAAACAAAAATCAATTATGGCTCACTGCTAG
GAGGACCAGCCACTTGTGTCTTGTGTACAGGGCTCACTGCTAGGAGGTCCATCAACAAATAATCCAAG
TAAGCCTCACATATGAGCTAAGCACATACGTCAGCTCTACAAAACTGGTACTTGTGTACAGGGAAAAA
TGTCAAAGATACTGGTACTTGTGTACAGGGCTCACTGCTAGTCACTATCAGCTAGGCATTGCGATAAA
GGAAAGGTCACTGCTAGGAGGACCGACAAATATTTCTTGTAAACTGGTACTTGTGTGCACACCAGCAT
GTGTTGATCACCAGCTGATCTTGCCTGCCTTGATCTATATAAGGTTTTGCTATTCATTGAAAGCAGTA
GTGACTGATTTGTATATA
>MinSyn_1826|Strength:0.008584496
GCGTGTCGTTTTAGTGAGGATGACGTAAGCCATGACGTCTAGGTACTTGTGTACAGGGCTAAGATTGA
TGAAAAGTCAAAAACAAAAATCAATTATCAAAACTGGTACTTGTGAAAAATGTCAAAGATAAGCAAGA

CCTCTACAAACAGCCACTTGTGTGCTCACTAACCACGTCTACAAGACCGATGCTGATCTTGCCCTATA
TAAGGTTTTGCTATTCATTGAAAGCAGTAGTGACTGATTTGTATATA
>MinSyn_1627|Strength:0.008703569
GCGTGTCGTTTTAGTGAGGAAAAATGTCAAAGATAGCTATCAGCTTAGCAAGACCTCTACAAAACTGG
TGGAGCACGACAGGACCGATCCATCAACAAATAATCCAAGTAAGCTGCTTTTCAACAATGTAGGCTAT
GACGTAAGCCATGACGTCTAGATCTTGGTGGGAGCCACCAATGCTGATCTTGCCTGCCTTGTTATGAC
CCCCGCCGATGACGCGGGACCAATTAGGTTGTCTGCACCTCACATGTATTGCGATAAAGGAAAGGGTT
CTATATAAGGTTTTGCTATTCATTGAAAGCAGTAGTGACTGATTTGTATATA
>MinSyn_1686|Strength:0.008718987
GCGTGTCGTTTTAGTGAGGAAGATTGATGAAAAGTCAAAAACAAAAATCAATTATTCGCGGTCTGACG
TAAGGGATGACGCACAGGACCGATGCTTTTCAACAAGGCTATCAGCTTAGCAAGCACACCAGCATGTG
TTGATCACCAGCTCACATGTATCCATCAACAAATAATCCAAGTAAGACAAAACTGGTACTTGTGTACA
GGAACCATTATTGCGTACAGGGCTATATAAGGTTTTGCTATTCATTGAAAGCAGTAGTGACTGATTTG
TATATA
>MinSyn_1551|Strength:0.008727066
GCGTGTCGTTTTAGTGAGGAACCACGTCTACAAACAGGGCTCACTGCTAGTGAAGATAAGATAATAAT
GTTGAAGATAAGACATGTAGGCTATCAGCAGCAAGTGGATGGCTATCAGCTTAGCAAGACCTCTACAA
AAAAAATGTCAAAGATAATCAGCTTAGCAAGACCTCTACACAGCCACTTGTGTCACATGTAGGCTATC
AGCTTCTGACGTAAGGGATGACGCACATCACATGTATCGAAAGGACAGTATATTTTCAACAACAGGGC
TCACAAATATTTCTTGTCTAGGAGGACCGATGCTGATCTTGCCTGCCTATATAAGGTTTTGCTATTCA
TTGAAAGCAGTAGTGACTGATTTGTATATA
>MinSyn_1791|Strength:0.008789707
GCGTGTCGTTTTAGTGAGGATCCTTACCGCTATGGGTAAGATTACCTCACATGTAGGCTATCAGCTCA
GTGGTCCCTCCACGTTGTCTGCACCTCACAATGACGTAAGCCATGACGTCTACCGATGCTGCAAATAT
TTCTTGTGGTTGTGGAAAAAGAAGAGGTTTTATGACCCCCGCCGATGACGCGGGACCTCTACAATCTC
TCTGCCGACAGTGGTCCCAAAATCAGCTTAGCAAGACCTCTACACTATATAAGGTTTTGCTATTCATT
GAAAGCAGTAGTGACTGATTTGTATATA
>MinSyn_1946|Strength:0.008805981
GCGTGTCGTTTTAGTGAGGATTGCGATAAAGGAAAGGGTACAGGGCTCACTGCTAGGAGGACCGATGA
TGACGTAAGCCATGACGTCTACAAGACCTCTACAAAACTGGTACATCGAAAGGACAGTACTAGGAGGA
CCGATGCTGATCTTTATGACCCCCGCCGATGACGCGGGACTTGGTGGAGCACGACACTTTTCAACAAC
GGTAGGTCACGACCACTATGCCCCTATATAAGGTTTTGCTATTCATTGAAAGCAGTAGTGACTGATTT
GTATATA
>MinSyn_1105|Strength:0.008837367
GCGTGTCGTTTTAGTGAGGTCAGAAGATCAAAGGGCTACTGCCTTGATTTTCAACAAGCCCAATTAGG
TTGTCTGCACCTCACATGTTCTCTCTGCCGACAGTGGTCCCAAATTGTGTACAGCAAATATTTCTTGT
TATGCCCAATTAGGTTGTCATGACGTAAGCCATGACGTCTAATTCCATCAACAAATAATCCAAGTAAG
AGCAAGACCTCTACAAAAATCGAAAGGACAGTACTAGGAGGACCGATGCTGATCTTGAAAAATGTCAA
AGATATGTACACTATATAAGGTTTTGCTATTCATTGAAAGCAGTAGTGACTGATTTGTATATA
>MinSyn_1571|Strength:0.008849122
GCGTGTCGTTTTAGTGAGGTCACTATCAGCCTGGTACTTGTGTACAGGGCTGGTGGAGCACGACATCA
CGACCACTATGCCTGACGTAAGGGATGACGCACAGTGTACAGGGCTCACTGCTAGGAATTTCGGGAAA
CCTCCTCGCAAAACTGGTACTTGTGTACAGGGCTCGTGGGAGCCACCAATGTAGGCTATCAATTGCGA
TAAAGGAAAGGCTTGCCTGCCTTGATGATACTATATAAGGTTTTGCTATTCATTGAAAGCAGTAGTGA
CTGATTTGTATATA
>MinSyn_1684|Strength:0.008993169
GCGTGTCGTTTTAGTGAGGTCACTATCAGCCACCTCACATGTAGGCTATCAGCGGAAAAAGAAGAGGT
ACTATGCCCAATTCAAATATTTCTTGTCCTCGCTTTGTCAAAAGCTAAAAAAGATGATGCATCTTGCC
TGCCTTGATTCTCTCTGCCGACAGTGGTCCCAAATAAACCATTATTGCGTGCCCAATTAGGTTGTCTA
TGACGTAAGCCATGACGTCTATCACGACCATCCTTACCGCTATGGGTAAGATTTGTAGGCTATCAGCT
TAGCAAGACCTCCAGCCACTTGTGTAATTAGGTTGTCTGCTATATAAGGTTTTGCTATTCATTGAAAG
CAGTAGTGACTGATTTGTATATA
>MinSyn_1407|Strength:0.009010382
GCGTGTCGTTTTAGTGAGGTCAGAAGATCAAAGGGCTAATGCCCAATTAGGTTGTCTATGACGTAAGC

CATGACGTCTACCTGTTATGACCCCCGCCGATGACGCGGGAGTGTACAGGGCTCACTGCTAAACCACG
TCTACAATACAAAACTGGTACTTGTGTAGCACACCAGCATGTGTTGATCACCAGCTCAAGACCTCTAC
AAAACTGGTACTTGCTATATAAGGTTTTGCTATTCATTGAAAGCAGTAGTGACTGATTTGTATATA
>MinSyn_1469|Strength:0.009013918
GCGTGTCGTTTTAGTGAGGATTGCGATAAAGGAAAGGACAGTGGTCCCTCCACCTCTAATGAGCTAAG
CACATACGTCAGGCCTTGAAATTTCGGGAAACCTCCTCGTTGTGTACAGGGCTCACTGCTAGGAGGAG
CACACCAGCATGTGTTGATCACCAGCTTCTTGCCTGCCTTGATGATATCTCTCTGCCGACAGTGGTCC
CAAACGACCACTATGATCGAAAGGACAGTATGTCACTATCAGCATGTAGGCTATCAGCTTAGCAAGAC
AAGATTGATGAAAAGTCAAAAACAAAAATCAATTATCTGATCTTGCCTGCCTTGATGACTATATAAGG
TTTTGCTATTCATTGAAAGCAGTAGTGACTGATTTGTATATA
>MinSyn_1874|Strength:0.009025062
GCGTGTCGTTTTAGTGAGGAGCAAGTGGATGTAGGCTATCAGCTTAGCAAGACTGAAGCATCTTCCGC
CCAGCCACTTGTGTGCGGTAGGTCACGACCACTATGCAAAAATGTCAAAGATACTTGTGTACAGGGCT
CACTGCTAGGAATGACGTAAGCCATGACGTCTAGACCTCTACAAAACTGGTACTTGTGTACATCTCTC
TGCCGACAGTGGTCCCAAAGCTATCAGCTTAGCAAGACCTTCAGAAGATCAAAGGGCTATACTTGTGT
ACCTATATAAGGTTTTGCTATTCATTGAAAGCAGTAGTGACTGATTTGTATATA
>MinSyn_1228|Strength:0.009025352
GCGTGTCGTTTTAGTGAGGAATTTCGGGAAACCTCCTCGATTAGGTTGTCTGCACCTATGAGCTAAGC
ACATACGTCAGGCCTTGCAAATATTTCTTGTTTAGGTTGTCTGCACCTCACATCCTTACCGCTATGGG
TAAGATTCAAGATTGATGAAAAGTCAAAAACAAAAATCAATTATGTACAGGGCTCACTGCTAGGAGGA
CCCAGCCACTTGTGTTCTACAAAACTGGTACTAACCATTATTGCGGTCTGCACCTCACATGTAGGTCA
CTATCAGCCACGACCATCAGAAGATCAAAGGGCTATGATCTTGCCTGCCTTGATGACTATATAAGGTT
TTGCTATTCATTGAAAGCAGTAGTGACTGATTTGTATATA
>MinSyn_1196|Strength:0.009078681
GCGTGTCGTTTTAGTGAGGTCTCTCTGCCGACAGTGGTCCCAAAATGATATGGTGGAGCACGACACCT
CTACAAAACTATCCTTACCGCTATGGGTAAGATTTGCTAGGAGGACCGATGCTGATCTGACGTAAGGG
ATGACGCACACAGAGCAAGTGGATAGGTTATTGCGATAAAGGAAAGGATGCCCAATTAGGTTGTCTGC
CAGTGGTCCCTCCACGTCACGACAAATATTTCTTGTGAGGACCGATGCTGATCTTGCTATATAAGGTT
TTGCTATTCATTGAAAGCAGTAGTGACTGATTTGTATATA
>MinSyn_1536|Strength:0.009092187
GCGTGTCGTTTTAGTGAGGAACCACGTCTACAAACGACCACTATGCCCAATTATGAGCTAAGCACATA
CGTCAGGCGGTAGGTCACGACCACTATGCTCACTATCAGCGTATCGAAAGGACAGTAACATGTAGGCT
ATCAGGCACACCAGCATGTGTTGATCACCAGCTTGCCTGCTCTCTCTGCCGACAGTGGTCCCAAACAA
TTAGGTTGTCTGCACTGGTGGAGCACGACAACCTCACATGTAGGCTATCAGCTTATCCATCAACAAAT
AATCCAAGTAAGGCTTAGGTGGCTCCTACCTCACATGTAGGCTATCAGCTTAGCTATATAAGGTTTTG
CTATTCATTGAAAGCAGTAGTGACTGATTTGTATATA
>MinSyn_1278|Strength:0.00909437
GCGTGTCGTTTTAGTGAGGAACCATTATTGCGCCAATTAGGTTGTCTGCAGGAAAAAGAAGAGGTTAG
GCTATCAGCTTAGCAAGACCTAATTTCGGGAAACCTCCTCGGACCACTATGCCCAATTAGGTCAAATA
TTTCTTGTTAGGTCATGAAGATAAGATAATAATGTTGAAGATAAGATATGCCCAAGATTGATGAAAAG
TCAAAAACAAAAATCAATTATGATGCTGATCTTGCCTGCCTCTGACGTAAGGGATGACGCACAGTAGG
TCACGACCACTATGCCCAATCCTTACCGCTATGGGTAAGATTGCGGTAGTGAAGCATCTTCCATGCTA
TATAAGGTTTTGCTATTCATTGAAAGCAGTAGTGACTGATTTGTATATA
>MinSyn_1256|Strength:0.009175925
GCGTGTCGTTTTAGTGAGGCTGACGTAAGGGATGACGCACAGACGTGGGAGCCACCACAAGACCTCTA
CAAAACTGGCACACCAGCATGTGTTGATCACCAGCTACTGCTAGGAGGCTTTGTCAAAAGCTAAAAAA
GATGATGCGGTTGTTTTCAACAAGCCTGCCTTGATGAGGTGGCTCCTACATGCCCAATCTATATAAGG
TTTTGCTATTCATTGAAAGCAGTAGTGACTGATTTGTATATA
>MinSyn_1779|Strength:0.009177695
GCGTGTCGTTTTAGTGAGGATCCTTACCGCTATGGGTAAGATTGCTGACGTAAGGGATGACGCACACT
GCTTGAAGCATCTTCCGGTAGGTCACGACCAAATATTTCTTGTGGCTTTGTCAAAAGCTAAAAAAGAT
GATGCTCTGCACCTCACATGGCACACCAGCATGTGTTGATCACCAGCTGCACCTCACATGTAGGCTAT
CAGCTCTATATAAGGTTTTGCTATTCATTGAAAGCAGTAGTGACTGATTTGTATATA
>MinSyn_1291|Strength:0.009258186

GCGTGTCGTTTTAGTGAGGGTGGGAGCCACCAATGCTGGTGGAGCACGACACCACTATGCCCAATTAG
GTTGTCTGAGCAAGTGGATCTGGTAAGATTGATGAAAAGTCAAAAACAAAAATCAATTATGGTACTTG
CTGACGTAAGGGATGACGCACACTGCACCTCACATGTAGGCTATATTGCGATAAAGGAAAGGCTATCA
GCTTAGCAAGACCTCTACAAAACGCACACCAGCATGTGTTGATCACCAGCTGACCGATGCTCTATATA
AGGTTTTGCTATTCATTGAAAGCAGTAGTGACTGATTTGTATATA
>MinSyn_1267|Strength:0.009381857
GCGTGTCGTTTTAGTGAGGATCCTTACCGCTATGGGTAAGATTTGGTACTTGATGACGTAAGCCATGA
CGTCTACCAATTAGGTTGTCTGTCCATCAACAAATAATCCAAGTAAGGATCTTGCCTGCCAAGATTGA
TGAAAAGTCAAAAACAAAAATCAATTATGACCGATGCGCACACCAGCATGTGTTGATCACCAGCTCTG
CACCTCACACTATATAAGGTTTTGCTATTCATTGAAAGCAGTAGTGACTGATTTGTATATA
>MinSyn_1970|Strength:0.009438651
GCGTGTCGTTTTAGTGAGGCAAATATTTCTTGTACCTCTACAAAACTTCACTATCAGCGTGTACATGA
AGCATCTTCCACCGATGCTGAAGCAAGTGGATCACATGTAGGCTATCAGCTTAGCAAAAAAATGTCAA
AGATATAGCAAGACCTCTACAAAATTTCGGGAAACCTCCTCGATGCTGATCTTGATGACGTAAGCCAT
GACGTCTACTTAGCAAGACCTCTACAAAACTGGTACGTGGGAGCCACCATCTACAAAACTGGTCAGTG
GTCCCTCCACATCTTGCCTGCCTCTATATAAGGTTTTGCTATTCATTGAAAGCAGTAGTGACTGATTT
GTATATA
>MinSyn_1457|Strength:0.009439569
GCGTGTCGTTTTAGTGAGGTCAGAAGATCAAAGGGCTAGTCTGCACCTCAGCACACCAGCATGTGTTG
ATCACCAGCTCTCACTGCTAGGAGGACCGATGCTGATCGCTTTGTCAAAAGCTAAAAAAGATGATGCT
GCACCTCACATGTAGGCTAAAAAATGTCAAAGATATTAGGTTGTCTGCACCTCACATGTAGAAGATTG
ATGAAAAGTCAAAAACAAAAATCAATTATCATGTAGGCTATCAAATTTCGGGAAACCTCCTCGAGGCC
TGACGTAAGGGATGACGCACAATCTTGCCTGTCACTATCAGCCCCAATTCAGCCACTTGTGTAACTGG
TACTTGTCTATATAAGGTTTTGCTATTCATTGAAAGCAGTAGTGACTGATTTGTATATA
>MinSyn_1823|Strength:0.00944882
GCGTGTCGTTTTAGTGAGGATGAGCTAAGCACATACGTCAGGACCACTATGCCCTTATGACCCCCGCC
GATGACGCGGGAGGTATCAGAAGATCAAAGGGCTAGGAGGACCGATGCTGATCTTGCCTATTGCGATA
AAGGAAAGGAATTAGGTTGTCTGCACCTCACATGTATTTTCAACAATCAGCTTAGCAAGACCTCTACA
AAAAATGTCAAAGATAGGTCACGACCCAGTGGTCCCTCCACCTGCACCTCACATGTAGGCTATCAGTG
AAGCATCTTCCGGTAGGTCACGACCACTACTATATAAGGTTTTGCTATTCATTGAAAGCAGTAGTGAC
TGATTTGTATATA
>MinSyn_1795|Strength:0.009495236
GCGTGTCGTTTTAGTGAGGCAGCCACTTGTGTCTATGCCCAATTAGGTTGTCTGCACCTCAATGAGCT
AAGCACATACGTCAGAGGCTATCAGCTTAGAATTTCGGGAAACCTCCTCGCACATATTGCGATAAAGG
AAAGGAATTAGGTTGTCTTTTTCAACAAGTCACGACCACTATGCCCAATTAGGAGCAAGTGGATGACC
GATGCTGAATCGAAAGGACAGTAGCAAGACCTCTACAAAACTGGTACTTGGGAAAAAGAAGAGGTCGA
TGCTGAAAGATTGATGAAAAGTCAAAAACAAAAATCAATTATAGGCTATATAAGGTTTTGCTATTCAT
TGAAAGCAGTAGTGACTGATTTGTATATA
>MinSyn_1180|Strength:0.009507787
GCGTGTCGTTTTAGTGAGGATGAGCTAAGCACATACGTCAGGCTGATCTTGCCTGCCTTGATGATTTT
TCAACAATGCTAGGAGGACCGATGCTGATCTTTCTCTCTGCCGACAGTGGTCCCAAAGTACAGGGCTC
ACTGCTAGGAGGACCGATGAGCAAGTGGATACCTCACATGTAGGCTATCAATTTCGGGAAACCTCCTC
GCTGATATCGAAAGGACAGTACCTCTTTATGACCCCCGCCGATGACGCGGGATATCAGAAGATTGATG
AAAAGTCAAAAACAAAAATCAATTATCTGATCTCTATATAAGGTTTTGCTATTCATTGAAAGCAGTAG
TGACTGATTTGTATATA
>MinSyn_1851|Strength:0.009508953
GCGTGTCGTTTTAGTGAGGAGCAAGTGGATACAGTGGTCCCTCCACGGCTATCAGCTTAGCAAGACCT
CTACAAAATGAAGCATCTTCCCTTGTGTACAGGGCTCACTGCTAGGCTGACGTAAGGGATGACGCACA
TAGGAGGACCGATCCTTACCGCTATGGGTAAGATTAGCAAGACCTCTACAACAGCCACTTGTGTACCG
ATGCTGATCCATCAACAAATAATCCAAGTAAGGCCTTGATGATACTATATAAGGTTTTGCTATTCATT
GAAAGCAGTAGTGACTGATTTGTATATA
>MinSyn_1515|Strength:0.009538386
GCGTGTCGTTTTAGTGAGGCAGCCACTTGTGTTGATCTTGCCTGCCTTGAGCACACCAGCATGTGTTG
ATCACCAGCTACCTCTCAAATATTTCTTGTTGTAGGCTATCAAGGTGGCTCCTACCCGATGCTGATCT

TGGGAAAAAGAAGAGGTATCTTGCCTGCCTTGATGAAGATTGATGAAAAGTCAAAAACAAAAATCAAT
TATGTTGTCTGCACCTCACATGTAGGCTGACGTAAGGGATGACGCACAAAACTGGTACTTGTGTACAG
GGCAACCATTATTGCGGTAGGCTATCAGCTTAGCAAGACCTCTACACTATATAAGGTTTTGCTATTCA
TTGAAAGCAGTAGTGACTGATTTGTATATA
>MinSyn_1412|Strength:0.009585007
GCGTGTCGTTTTAGTGAGGCAGCCACTTGTGTTAGGAGGACCGAAACCACGTCTACAACTAGGAGGAC
CGATCCTTACCGCTATGGGTAAGATTCACGACCACTATCAGAAGATCAAAGGGCTACAGCTTAGCAAG
ACCTCTACAAATTTCGGGAAACCTCCTCGTTGTGTACAGGGCTCACTGGGAAAAAGAAGAGGTGCTCA
CTGCTAGGAGGAGTGGGAGCCACCAACCGATGCTGATCTATGACGTAAGCCATGACGTCTATAGCAAG
ACCTCTACAAATCCATCAACAAATAATCCAAGTAAGCTGCTAGGAGGACCGATGCTATATAAGGTTTT
GCTATTCATTGAAAGCAGTAGTGACTGATTTGTATATA
>MinSyn_1516|Strength:0.009587954
GCGTGTCGTTTTAGTGAGGGGAAAAAGAAGAGGTGGTACTTGTGTACAGGGCTCACTGCTAGGATTTT
CAACAAGACCACTATGCCCAATTAGGTTAGCAAGTGGATAGCAAGATGACGTAAGCCATGACGTCTAC
ACGACCACTATGCCCAATCCTTACCGCTATGGGTAAGATTTAGCAAAAAAATGTCAAAGATATCACAT
GTAGGCTATCAGCTTAGCAAGACATCGAAAGGACAGTACAATCTATATAAGGTTTTGCTATTCATTGA
AAGCAGTAGTGACTGATTTGTATATA
>MinSyn_1678|Strength:0.009589111
GCGTGTCGTTTTAGTGAGGGCTTTGTCAAAAGCTAAAAAAGATGATGCTTATGAGCTAAGCACATACG
TCAGGTACTTGTGTACAGGGCTCACTGATCCTTACCGCTATGGGTAAGATTGCTAGGAGGACCGATGC
TGATCATCGAAAGGACAGTAATGTAGGCTATCAGCTTAGCAAGACAACCACGTCTACAACTTGCCTGC
CTTGTGGTGGAGCACGACAACTTGTGTACAGGGCTCACTGCTAGGAGGAGGAAAAAGAAGAGGTCACT
ATGCCCAATAGGTGGCTCCTACAGGGCTCACTGCTACTATATAAGGTTTTGCTATTCATTGAAAGCAG
TAGTGACTGATTTGTATATA
>MinSyn_1656|Strength:0.009593182
GCGTGTCGTTTTAGTGAGGGTGGGAGCCACCAAAAACTGGTACTTGTGTACAGGGCTGACGTAAGGGA
TGACGCACATCTGCACCTTCAGAAGATCAAAGGGCTAGCTAGGAATCCTTACCGCTATGGGTAAGATT
GGAGGACCGATGCTGATCTTGCCTGCCTTAGCAAGTGGATGTAGGTCTCTCTGCCGACAGTGGTCCCA
AAACTATCTATATAAGGTTTTGCTATTCATTGAAAGCAGTAGTGACTGATTTGTATATA
>MinSyn_1702|Strength:0.009598821
GCGTGTCGTTTTAGTGAGGTGGTGGAGCACGACATCTACATGAGCTAAGCACATACGTCAGGCAAGAC
CTCTACAAAACTGGTACTTGTGAGCAAGTGGATAGCAAGACCTCTGCACACCAGCATGTGTTGATCAC
CAGCTCTATGGGAAAAAGAAGAGGTTCACATGTAGCTTTGTCAAAAGCTAAAAAAGATGATGCCTCAC
TGTGGGAGCCACCACTAAGATTGATGAAAAGTCAAAAACAAAAATCAATTATTGTACAGGGCTCACTG
CTAGGAGGACCGACAGCCACTTGTGTAGCAAGACCTCTATATAAGGTTTTGCTATTCATTGAAAGCAG
TAGTGACTGATTTGTATATA
>MinSyn_1154|Strength:0.009604391
GCGTGTCGTTTTAGTGAGGATGAGCTAAGCACATACGTCAGTGTACAGGGCTCACTGCTAGGAAGATT
GATGAAAAGTCAAAAACAAAAATCAATTATACAAAACTAACCATTATTGCGCCGATGCTGATCTTGCC
AAAAATGTCAAAGATACCACTATGCCCAATTAGGTTGTCTGCACCAATTTCGGGAAACCTCCTCGCAA
TTAGGTTGTCTGCACCTCACATGTAGTTTTCAACAAGCCTGCCTTGTTATGACCCCCGCCGATGACGC
GGGATGTACAGGGCTCACTGCTAGGACTATATAAGGTTTTGCTATTCATTGAAAGCAGTAGTGACTGA
TTTGTATATA
>MinSyn_1386|Strength:0.00961981
GCGTGTCGTTTTAGTGAGGATGAGCTAAGCACATACGTCAGGATGCTGATCTTGCCCAGCCACTTGTG
TAAGCAAGTGGATACTTGTGTACAGGGCTCATCTCTCTGCCGACAGTGGTCCCAAATAATTTCGGGAA
ACCTCCTCGGCACCTCACATGTAGGCTATTTTTCAACAAACCTCACATGTAGGCTATCATTGCGATAA
AGGAAAGGAAAACTGGTACTTGTGTAAACCACGTCTACAACCCAATTAGGTTTTATGACCCCCGCCGA
TGACGCGGGACAGCTTAGCAAGACCTCCTATATAAGGTTTTGCTATTCATTGAAAGCAGTAGTGACTG
ATTTGTATATA
>MinSyn_1721|Strength:0.009622854
GCGTGTCGTTTTAGTGAGGTCAGAAGATCAAAGGGCTAGACCGATGCTGATCTTGCCTATCCTTACCG
CTATGGGTAAGATTCCTCTACAAAACTGGTACTTGTGTACACTGACGTAAGGGATGACGCACATACAG
GGCTCACTGCTAGGAGAACCACGTCTACAAGCTCACTGTCTCTCTGCCGACAGTGGTCCCAAACCTCA

CATGTATCACTATCAGCGTGTACAGGGCTCACTGCTAGGCTATATAAGGTTTTGCTATTCATTGAAAG
CAGTAGTGACTGATTTGTATATA
>MinSyn_1577|Strength:0.009626294
GCGTGTCGTTTTAGTGAGGATGACGTAAGCCATGACGTCTAGTTGTCTGCACCTCGCACACCAGCATG
TGTTGATCACCAGCTCTACAAAACTGGTACTTGTGTACAGGTCTCTCTGCCGACAGTGGTCCCAAAGG
TCACGACCACTATGCCCAATTAGGTCCATCAACAAATAATCCAAGTAAGTGACTATATAAGGTTTTGC
TATTCATTGAAAGCAGTAGTGACTGATTTGTATATA
>MinSyn_1428|Strength:0.009645923
GCGTGTCGTTTTAGTGAGGCAGCCACTTGTGTCGGAAAAAGAAGAGGTTACTTGTGTACATCCTTACC
GCTATGGGTAAGATTGATGACGTAAGCCATGACGTCTATCACATGTAGGCTATCATCCATCAACAAAT
AATCCAAGTAAGTGTAGGCTATCAGCTTATGACCCCCGCCGATGACGCGGGACTGATCTTGCCTGCCT
CAGAAGATCAAAGGGCTAGTGTACACTATATAAGGTTTTGCTATTCATTGAAAGCAGTAGTGACTGAT
TTGTATATA
>MinSyn_1219|Strength:0.009667784
GCGTGTCGTTTTAGTGAGGAACCACGTCTACAATCAGCTTAGCAAGACATGACGTAAGCCATGACGTC
TAGCCCAATTAGGTTGTCTGCACCTCACAGGAAAAAGAAGAGGTGATGCTGATCTTGCATTGCGATAA
AGGAAAGGAATTAGGTTGTCTGCACCAAAAATGTCAAAGATAGTCACGACCACTATGCCCAATTAGGT
CTATATAAGGTTTTGCTATTCATTGAAAGCAGTAGTGACTGATTTGTATATA
>MinSyn_1668|Strength:0.009717493
GCGTGTCGTTTTAGTGAGGATGACGTAAGCCATGACGTCTATGCCCAATTAGGTTGTTCACTATCAGC
CACCTCACATGTAGGCTATCAGCTTAGCAAATTTCGGGAAACCTCCTCGTCACGACCACTATGCAAAT
ATTTCTTGTCACATGTAGGCTATTTTCAACAAACCGATGCTGATCTTGCCCTATATAAGGTTTTGCTA
TTCATTGAAAGCAGTAGTGACTGATTTGTATATA
>MinSyn_1833|Strength:0.009735716
GCGTGTCGTTTTAGTGAGGTCTCTCTGCCGACAGTGGTCCCAAACAAGAAGCAAGTGGATTGTAGGCT
ATAACCATTATTGCGATGTAGGCTATCAGCTTAGCAAGACCTAAGATTGATGAAAAGTCAAAAACAAA
AATCAATTATCACTATGCCCAATTAGGTTGTCTGCGCACACCAGCATGTGTTGATCACCAGCTGCAAG
ACCTCTACAAAACTGGCTGACGTAAGGGATGACGCACAACTGGTACTTGTGTACAGGATTGCGATAAA
GGAAAGGCATGTAGGCTATCAGCTTAGCAAGACCTCTCTATATAAGGTTTTGCTATTCATTGAAAGCA
GTAGTGACTGATTTGTATATA
>MinSyn_1546|Strength:0.009810641
GCGTGTCGTTTTAGTGAGGCAGCCACTTGTGTTATGCCCAATGACGTAAGCCATGACGTCTAACGACC
ACTATTCTCTCTGCCGACAGTGGTCCCAAACACTGCTAGGAGGACCGATGCTGATCTTGCTTTGTCAA
AAGCTAAAAAAGATGATGCGCAAGACCTCTACATGAAGCATCTTCCGGTCACGACCACTATGCCTATA
TAAGGTTTTGCTATTCATTGAAAGCAGTAGTGACTGATTTGTATATA
>MinSyn_1784|Strength:0.009879511
GCGTGTCGTTTTAGTGAGGGCACACCAGCATGTGTTGATCACCAGCTTAATGAGCTAAGCACATACGT
CAGTGCCTGCCTTGATGAAACCATTATTGCGTCTACAAAACTGGTACTTGTGTACAGGTCAGAAGATC
AAAGGGCTATCACTGCTAGGAGGACCGAATCGAAAGGACAGTACACTATGCCATCCTTACCGCTATGG
GTAAGATTCACCTCACATGTAGGCTATCAGCTTAGCAAGCAAGTGGATAAGACAATTTCGGGAAACCT
CCTCGAAAACTGGTACTTGTGTACAGGGCCTATATAAGGTTTTGCTATTCATTGAAAGCAGTAGTGAC
TGATTTGTATATA
>MinSyn_1677|Strength:0.009914117
GCGTGTCGTTTTAGTGAGGTCAGAAGATCAAAGGGCTATAGGTCACGACCACATGAGCTAAGCACATA
CGTCAGCTGCTAGGAGGACCGATGCTGCTTTGTCAAAAGCTAAAAAAGATGATGCCAAAACTGGTACT
TGTGTACAGTGGTGGAGCACGACAGCCTGGTGGGAGCCACCATAGCAAGACCTCTAAGATTGATGAAA
AGTCAAAAACAAAAATCAATTATTGCCATCCTTACCGCTATGGGTAAGATTGTACAGGGCTCACTGCA
AAAATGTCAAAGATAACATGTAGGCTATCAGCCTATATAAGGTTTTGCTATTCATTGAAAGCAGTAGT
GACTGATTTGTATATA
>MinSyn_119|Strength:0.009950325
GCGTGTCGTTTTAGTGAGGTGAAGCATCTTCCCTGCTAGGAGGACCGATGCTGATCTTGCTGGTGGAG
CACGACACACTATGCCCACTGACGTAAGGGATGACGCACACTGCTAGGAGTCCATCAACAAATAATCC
AAGTAAGACCTCTACAAAACTGGTACTTGGCACACCAGCATGTGTTGATCACCAGCTCTGGTACTTGT
AAAAATGTCAAAGATACACTGCCTATATAAGGTTTTGCTATTCATTGAAAGCAGTAGTGACTGATTTG

TATATA
>MinSyn_1911|Strength:0.010018527
GCGTGTCGTTTTAGTGAGGTGAAGCATCTTCCAGGGCTCACTGCTAGGAGGACCGATGCTGAGGAAAA
AGAAGAGGTAGGACCAATTTCGGGAAACCTCCTCGAGCTTTGTCAAAAGCTAAAAAAGATGATGCATT
AGGTTGTCTGCACCTCACATGTAGGCTATGACGTAAGCCATGACGTCTAGACCGATGCTGATCTTGCC
TGCCTTGATGTTTTCAACAACATGTAGGCTATCAGCTTAGCAAGACCTCAGGTGGCTCCTACCAGGCT
ATATAAGGTTTTGCTATTCATTGAAAGCAGTAGTGACTGATTTGTATATA
>MinSyn_1142|Strength:0.010018768
GCGTGTCGTTTTAGTGAGGATGAGCTAAGCACATACGTCAGCTAGGATGGTGGAGCACGACACTACAA
AACTGGTACTTGTGTTCCATCAACAAATAATCCAAGTAAGGACCACTATGCCTCTCTCTGCCGACAGT
GGTCCCAAACTTGTGTACAGGGCTCACTGCTAGAGCAAGTGGATTCAATTTCGGGAAACCTCCTCGTT
AGGTTGTCTGCACCTCACATGTCAGAAGATCAAAGGGCTAAGGACCGATGCTGATCAAAAATGTCAAA
GATACTGGTACTTGTGTCTATATAAGGTTTTGCTATTCATTGAAAGCAGTAGTGACTGATTTGTATAT
A
>MinSyn_1816|Strength:0.010076077
GCGTGTCGTTTTAGTGAGGTTATGACCCCCGCCGATGACGCGGGAGCTATCAGCTTAGCCAAATATTT
CTTGTGTACTTGTGTACAGGGCTCACTGCTAGGATGAGCTAAGCACATACGTCAGATTAGGTTGTCTG
CACCTCACATTGAAGATAAGATAATAATGTTGAAGATAAGAGGTAGGTCATCCTTACCGCTATGGGTA
AGATTAACTGGTACTTGTGTACAGGGCTCACTGCTTTTTCAACAAGCAGGTGGCTCCTACGTACAGGG
CTCACTGCTAGGAGGACCGATTGCGATAAAGGAAAGGTCTACTATATAAGGTTTTGCTATTCATTGAA
AGCAGTAGTGACTGATTTGTATATA
>MinSyn_1935|Strength:0.010083032
GCGTGTCGTTTTAGTGAGGGGAAAAAGAAGAGGTCCACTAATTGCGATAAAGGAAAGGACTATGCCCA
ATTAGGTTGTCTGCCTGACGTAAGGGATGACGCACACTTAGCAAGACCTCTAGGTGGCTCCTACTATC
AGCTTAGCAAGATCACTATCAGCCGCGGTAGGTCGCTTTGTCAAAAGCTAAAAAAGATGATGCAGTGG
GAGCCACCATCACATGTACTATATAAGGTTTTGCTATTCATTGAAAGCAGTAGTGACTGATTTGTATA
TA
>MinSyn_1375|Strength:0.010168007
GCGTGTCGTTTTAGTGAGGATCGAAAGGACAGTATTAGCAAATGACGTAAGCCATGACGTCTACCACT
ATGCCCAATTAAACCATTATTGCGTGAGCTTTGTCAAAAGCTAAAAAAGATGATGCTCTACATGAAGA
TAAGATAATAATGTTGAAGATAAGATTGCTCTCTCTGCCGACAGTGGTCCCAAATCTACAAACTATAT
AAGGTTTTGCTATTCATTGAAAGCAGTAGTGACTGATTTGTATATA
>MinSyn_1155|Strength:0.010243331
GCGTGTCGTTTTAGTGAGGATGACGTAAGCCATGACGTCTACCCAATTAGGTTGTCTGCACCTCACAG
TGGTCCCTCCACGCTTAGCATTGCGATAAAGGAAAGGAGCAAGACCTCTACAAAACTGGTACTTTATG
ACCCCCGCCGATGACGCGGGAGGCTCACTGCTAGGAGGACCGATGCTATATAAGGTTTTGCTATTCAT
TGAAAGCAGTAGTGACTGATTTGTATATA
>MinSyn_1580|Strength:0.010275649
GCGTGTCGTTTTAGTGAGGGCACACCAGCATGTGTTGATCACCAGCTTAGGTTGTCTGCACCTCACAT
GTATTGCGATAAAGGAAAGGAGGACCGATGCTGATCTTGCCTGCCTTGAACCATTATTGCGGATCCAT
CAACAAATAATCCAAGTAAGTGCACCTCACATGTATGAAGCATCTTCCTCACATGTAGATCCTTACCG
CTATGGGTAAGATTTCAGCTTAGCAAGACCTCCTGACGTAAGGGATGACGCACAGGCAACCACGTCTA
CAAAGCTTAGCAAGACCTCTCAAATATTTCTTGTAGCTTACTATATAAGGTTTTGCTATTCATTGAAA
GCAGTAGTGACTGATTTGTATATA
>MinSyn_1483|Strength:0.010343937
GCGTGTCGTTTTAGTGAGGATCCTTACCGCTATGGGTAAGATTCTTGTGTACAGGGCTCACTGCTAGG
AGATCGAAAGGACAGTACACATGTAGGCTCTGACGTAAGGGATGACGCACAACCGATGCTGATCTTGC
CTGGGAAAAAGAAGAGGTGTTGTCTGCACCTCACATGTAGGCTAAATTTCGGGAAACCTCCTCGAGGA
CCGATGCTGATCTTGCCTGCCTCTATATAAGGTTTTGCTATTCATTGAAAGCAGTAGTGACTGATTTG
TATATA
>MinSyn_1307|Strength:0.0103845
GCGTGTCGTTTTAGTGAGGTGGTGGAGCACGACAACTATGCCCAATTATGAGCTAAGCACATACGTCA
GTCACTGCTAGGAGATTGCGATAAAGGAAAGGAGAGGTGGCTCCTACCTATGCCCAATTAGTTTTCAA
CAATCAGCTTAGCAAGACCTCCAGCCACTTGTGTTGATGATCCATCAACAAATAATCCAAGTAAGCGG

TAGGTCACGACCACAGCAAGTGGATATTAGGTTGTCTGCATGAAGCATCTTCCAGACCTCTACAAAAC
TGGTACTTGTGTACACTATATAAGGTTTTGCTATTCATTGAAAGCAGTAGTGACTGATTTGTATATA
>MinSyn_1231|Strength:0.01039448
GCGTGTCGTTTTAGTGAGGTGAAGCATCTTCCAGCTTAGCAAGACCTCTATCACTATCAGCGTGTACA
GGGAAAAATGTCAAAGATATCACAACCATTATTGCGGCTAGGAGGACCGATGCTGAAACCACGTCTAC
AATGTCTGCACCTCACATGACGTAAGCCATGACGTCTATTGTCTGCACCTCACATGTAGGTTTTCAAC
AAGTTTCCATCAACAAATAATCCAAGTAAGCTACAAAACTGGTACTTGTGTACAGCTATATAAGGTTT
TGCTATTCATTGAAAGCAGTAGTGACTGATTTGTATATA
>MinSyn_1392|Strength:0.010422733
GCGTGTCGTTTTAGTGAGGTGAAGCATCTTCCGATGCTGATCTTGCTGACGTAAGGGATGACGCACAA
GGTTGTCTGCACCTCACATAGGTGGCTCCTACAGGTCACGACCACTATGCCCCAGTGGTCCCTCCACA
CGACCTCACTATCAGCCCTGCAAAAATGTCAAAGATATAGGAGGACCGATGCTGATCTTCTATATAAG
GTTTTGCTATTCATTGAAAGCAGTAGTGACTGATTTGTATATA
>MinSyn_1249|Strength:0.010454956
GCGTGTCGTTTTAGTGAGGGGAAAAAGAAGAGGTACTGGTACTTCAGCCACTTGTGTTTGTGTACAGG
GCTCACTGACGTAAGGGATGACGCACACATGTAGCAAGTGGATAGGTCACGACCACTATGCCCAATTA
GTGAAGATAAGATAATAATGTTGAAGATAAGAAGGTTGTCTGCACCTCACATGTAGGCTTGGTGGAGC
ACGACAGGCTATATAAGGTTTTGCTATTCATTGAAAGCAGTAGTGACTGATTTGTATATA
>MinSyn_1308|Strength:0.010560995
GCGTGTCGTTTTAGTGAGGAACCATTATTGCGTCTACAAAACTGGTACTTGTGTACAGGGCTTTATGA
CCCCCGCCGATGACGCGGGAGTAGGCTATCAGCTTAGAAAAATGTCAAAGATAATGCCCAATTAGGTT
GTCTGCACCGCTTTGTCAAAAGCTAAAAAAGATGATGCTCTACAAAACTGGAGGTGGCTCCTACTTAG
GTTGTCTGCACCTCACATGTACAAATATTTCTTGTAACTGGTACTTGTGTACAGGGCTCTCTCTCTGC
CGACAGTGGTCCCAAATGCCCAATTAGGTTGTCTGGTGGAGCACGACACCTCTACAAAACTGGTACTT
GTGTACAATGACGTAAGCCATGACGTCTACAAAACTCTATATAAGGTTTTGCTATTCATTGAAAGCAG
TAGTGACTGATTTGTATATA
>MinSyn_1177|Strength:0.010583141
GCGTGTCGTTTTAGTGAGGAGCAAGTGGATCGGCTTTGTCAAAAGCTAAAAAAGATGATGCAGGGCTC
ACTGTGGTGGAGCACGACACAAGACCTCTACATGACGTAAGCCATGACGTCTAAGGACCGATGCTGAT
CTTGCCTGAAGCATCTTCCAGCAAGACCTCTACAAAACTGGTAAAGATTGATGAAAAGTCAAAAACAA
AAATCAATTATTTGCCTGCCTTCTATATAAGGTTTTGCTATTCATTGAAAGCAGTAGTGACTGATTTG
TATATA
>MinSyn_1809|Strength:0.01058544
GCGTGTCGTTTTAGTGAGGCAAATATTTCTTGTCTAGGAGGACCGATGTGAAGCATCTTCCACCTCTA
CAAAACTGGTACTTGTGTACAGTGGTGGAGCACGACAAATTAGGTTGTCTGCACCTCACATGAAGATT
GATGAAAAGTCAAAAACAAAAATCAATTATATGTAGGCTATTCACTATCAGCTCTACAAAACTGGTAC
TTGTGTACTTATGACCCCCGCCGATGACGCGGGATCACGACCACTATGCCCAATTAGATCCTTACCGC
TATGGGTAAGATTTATGCCTCAGAAGATCAAAGGGCTACGCGGTAGGTCACGACCACTAATGAGCTAA
GCACATACGTCAGCGATGCTGATCTTGCCTGCCTTGCTATATAAGGTTTTGCTATTCATTGAAAGCAG
TAGTGACTGATTTGTATATA
>MinSyn_1433|Strength:0.01065618
GCGTGTCGTTTTAGTGAGGAATTTCGGGAAACCTCCTCGTACAAAACTGGTACAGCCACTTGTGTTGC
CATGAGCTAAGCACATACGTCAGTGCCTGCCTTGATATCCTTACCGCTATGGGTAAGATTAGGAGGAC
CGATGCTGATCTTGCCTCTCTCTGCCGACAGTGGTCCCAAAACCACTATGCCCAATTAGGTTGTCTGC
GGAAAAAGAAGAGGTGATGCTGATCTTGCCTGCCTTGATGATAAAGATTGATGAAAAGTCAAAAACAA
AAATCAATTATAGGTTGCTATATAAGGTTTTGCTATTCATTGAAAGCAGTAGTGACTGATTTGTATAT
A
>MinSyn_146|Strength:0.010693943
GCGTGTCGTTTTAGTGAGGTCTCTCTGCCGACAGTGGTCCCAAACTCACTGCTAGGAGGACCGATGCA
TGAGCTAAGCACATACGTCAGGGTAGGATCCTTACCGCTATGGGTAAGATTAGGGCTCACTGCTAACC
ACGTCTACAAGCAAGACCTCTACAAAACTGGTACCAGTGGTCCCTCCACTAGCAAGTGGATTGCTAGC
ACACCAGCATGTGTTGATCACCAGCTCTGATCTTGCCTGCCAGCCACTTGTGTCTGCACCTCACATGT
AGGCTATCAGCTTAGCTATATAAGGTTTTGCTATTCATTGAAAGCAGTAGTGACTGATTTGTATATA
>MinSyn_1293|Strength:0.010761936

GCGTGTCGTTTTAGTGAGGATGACGTAAGCCATGACGTCTAGGCTCACTGCTAGGAGGAGCAAGTGGA
TAGGGCTCACTGCTATGAAGATAAGATAATAATGTTGAAGATAAGACTGCTAGGAGGACATTGCGATA
AAGGAAAGGCAGGGCTCACTATCAGCGCTAGGAGGACCGATCTATATAAGGTTTTGCTATTCATTGAA
AGCAGTAGTGACTGATTTGTATATA
>MinSyn_1209|Strength:0.010995678
GCGTGTCGTTTTAGTGAGGTGGTGGAGCACGACAGGTACGCACACCAGCATGTGTTGATCACCAGCTC
TGCTAGGAGGAGGAAAAAGAAGAGGTTAGGAGGACCGATGCTGATCTTGCATGACGTAAGCCATGACG
TCTAATCTCAGAAGATCAAAGGGCTAACTGGTACTTGTGTATCCATCAACAAATAATCCAAGTAAGCT
ACAAAACTGGTACTTGTGTACAGGGCTCCTATATAAGGTTTTGCTATTCATTGAAAGCAGTAGTGACT
GATTTGTATATA
>MinSyn_1230|Strength:0.011020269
GCGTGTCGTTTTAGTGAGGTGAAGATAAGATAATAATGTTGAAGATAAGAGCGGTAGGTCACGACCAC
TATGACGTAAGCCATGACGTCTAGTAGTCACTATCAGCACCTCACATGTATTATGACCCCCGCCGATG
ACGCGGGAACGCTTTGTCAAAAGCTAAAAAAGATGATGCAGCAAGACCTCTACAAAACTGGTACTTGT
CTATATAAGGTTTTGCTATTCATTGAAAGCAGTAGTGACTGATTTGTATATA
>MinSyn_1847|Strength:0.01103375
GCGTGTCGTTTTAGTGAGGTCACTATCAGCGGAGGACCGATGCTGATCTTGAAGATAAGATAATAATG
TTGAAGATAAGACCGATGCTGATCTTGCCTGCCTTGATGATCGAAAGGACAGTATCTACAAAACTGGT
ACTAACCATTATTGCGCTCACTGCTAGGAGGACCGATGCTATCCTTACCGCTATGGGTAAGATTGGGC
TCACTGCTAGATGACGTAAGCCATGACGTCTAGGTTGTCTGAGGTGGCTCCTACTCACATAATTTCGG
GAAACCTCCTCGTAGGCTATATAAGGTTTTGCTATTCATTGAAAGCAGTAGTGACTGATTTGTATATA
>MinSyn_1879|Strength:0.011153636
GCGTGTCGTTTTAGTGAGGGCACACCAGCATGTGTTGATCACCAGCTGGGCTCACTGCTAGGAGGACC
GATGCCAGTGGTCCCTCCACGAGGACCGATGCTGATCTTGCCTGCAGCCACTTGTGTGCACCTCACAT
GTAGGCTATCAGCATCCTTACCGCTATGGGTAAGATTGACCACTATGCCCAATTAGGTTGTCTATTGC
GATAAAGGAAAGGAATGAGCTAAGCACATACGTCAGATCTTGCCTGCCTTGATGATTGAAGCATCTTC
CAGAAGATTGATGAAAAGTCAAAAACAAAAATCAATTATCCGATGCTGATCTTGCAGCAAGTGGATAA
GACCTCTACAAAACTCTATATAAGGTTTTGCTATTCATTGAAAGCAGTAGTGACTGATTTGTATATA
>MinSyn_1768|Strength:0.011156075
GCGTGTCGTTTTAGTGAGGAATTTCGGGAAACCTCCTCGTATGCCCAATTATCAGAAGATCAAAGGGC
TAACAAAACTGGTACTTGTGTACAGGGCATGAGCTAAGCACATACGTCAGGTTGTCTGCACCTCACAT
GTAGGCTATAAGATTGATGAAAAGTCAAAAACAAAAATCAATTATGTACTTGTGTAGGAAAAAGAAGA
GGTAGGTTGTCTGCACCTCACATGTAGGCTATATTGCGATAAAGGAAAGGTCGCGGTCAGCCACTTGT
GTCCCAATTAGGTCTATATAAGGTTTTGCTATTCATTGAAAGCAGTAGTGACTGATTTGTATATA
>MinSyn_1960|Strength:0.011185653
GCGTGTCGTTTTAGTGAGGTCCATCAACAAATAATCCAAGTAAGTATGCCCAATTAGGTTGTCTGCAT
CGAAAGGACAGTAATGCCCAATTAGGTTGTCTGCACTGAAGCATCTTCCACTATGCCCAATTAGGTTG
TTCAGAAGATCAAAGGGCTAACCGATGCTGACTGACGTAAGGGATGACGCACACTGCTAGGAGGACCG
ATCAGTGGTCCCTCCACTTAGCAAGACCTCTGGTGGAGCACGACAAGGGCTCACCTATATAAGGTTTT
GCTATTCATTGAAAGCAGTAGTGACTGATTTGTATATA
>MinSyn_1338|Strength:0.011186922
GCGTGTCGTTTTAGTGAGGATTGCGATAAAGGAAAGGACAAAACTGGTACTGGTGGAGCACGACATGA
TCTTGCCTGCCTCAGAAGATCAAAGGGCTATTGTCTTTATGACCCCCGCCGATGACGCGGGATATGCC
CAATTAGGTTGTCTGCACCTCAACCACGTCTACAAGCCCAATTAGGTTGTCTGCACCTCGTGGGAGCC
ACCACGGTAGGTCACGACCACTATGAGCTAAGCACATACGTCAGAATTAGGTTGTCTAATTTCGGGAA
ACCTCCTCGTTGTCTGCACCTCACATGTAGGCTATCTGAAGCATCTTCCAGGACCGATGCTGATCTTG
CCTGCCTTGCTATATAAGGTTTTGCTATTCATTGAAAGCAGTAGTGACTGATTTGTATATA
>MinSyn_174|Strength:0.011253946
GCGTGTCGTTTTAGTGAGGCTGACGTAAGGGATGACGCACACGACCACAATTTCGGGAAACCTCCTCG
TAGGTCACGACCACTATGCAAAAATGTCAAAGATACGCGGTAGGTCACGACCAGCTTTGTCAAAAGCT
AAAAAAGATGATGCAGACCTCTACAAAACTGGTACTCTATATAAGGTTTTGCTATTCATTGAAAGCAG
TAGTGACTGATTTGTATATA
>MinSyn_1114|Strength:0.011274609
GCGTGTCGTTTTAGTGAGGATGACGTAAGCCATGACGTCTACACATGCAGCCACTTGTGTGGACCGGG

AAAAAGAAGAGGTAACTGGTACTTGTTGAAGCATCTTCCAGGAGGACCGATGCTGATCTTGATCGAAA
GGACAGTATTAGCAAGACCTCTACAAAACTGGTACCTATATAAGGTTTTGCTATTCATTGAAAGCAGT
AGTGACTGATTTGTATATA
>MinSyn_1888|Strength:0.011320303
GCGTGTCGTTTTAGTGAGGCAGCCACTTGTGTTACAGGGCTCACTGCTAGGAGGACCGATTTATGACC
CCCGCCGATGACGCGGGAACTATGCCAACCATTATTGCGAGGGCACACCAGCATGTGTTGATCACCAG
CTCTGGTACTTGTGTACAGGGAATTTCGGGAAACCTCCTCGCCACTATGCCCAATTAGGTTGTCTGCA
CATGACGTAAGCCATGACGTCTAGTACTTGTGTACTGAAGATAAGATAATAATGTTGAAGATAAGACA
TGTAGGCCTATATAAGGTTTTGCTATTCATTGAAAGCAGTAGTGACTGATTTGTATATA
>MinSyn_1552|Strength:0.011321949
GCGTGTCGTTTTAGTGAGGGGAAAAAGAAGAGGTCAGGGCTCACTGCTAGGAGAGCAAGTGGATCCTC
ACATGTAGGCTATTGAAGATAAGATAATAATGTTGAAGATAAGAAGGTCACGACCACTATGCCCAATT
AAAGATTGATGAAAAGTCAAAAACAAAAATCAATTATACTGACGTAAGGGATGACGCACACACGACCA
CTATGCCCAATTAGGTCACTATCAGCATTAGGTTGTCTGCACCTCACATGTAGGCTCTATATAAGGTT
TTGCTATTCATTGAAAGCAGTAGTGACTGATTTGTATATA
>MinSyn_1576|Strength:0.011332464
GCGTGTCGTTTTAGTGAGGAGCAAGTGGATATGATCGAAAGGACAGTAGTACAGGGCTCACTCACTAT
CAGCGCAAGACCTCTACAAAACTGGTACTTGTGCTTTGTCAAAAGCTAAAAAAGATGATGCGATCTTG
CCTGCCTTGATGATAAAGATTGATGAAAAGTCAAAAACAAAAATCAATTATCTATGCCCAATTAGGTT
TTTTCAACAAGTAGGCTATCAGCTTAGCAAGACCTCAGCCACTTGTGTTCACTGCTAGGAGGACCGAT
GTCCATCAACAAATAATCCAAGTAAGCTCTGACGTAAGGGATGACGCACATGCTGATCTTGCCTGCCT
TGATGATCTATATAAGGTTTTGCTATTCATTGAAAGCAGTAGTGACTGATTTGTATATA
>MinSyn_1843|Strength:0.011421247
GCGTGTCGTTTTAGTGAGGTCACTATCAGCGATTTTTCAACAAATGCCCAATTAGGTTGTCTGCATCT
CTCTGCCGACAGTGGTCCCAAAAGGAGGACCGATGAAGATAAGATAATAATGTTGAAGATAAGACTCT
ACAAAACTGGTACTTGTGTTTATGACCCCCGCCGATGACGCGGGAACATGTAGCACACCAGCATGTGT
TGATCACCAGCTTGTGTACAGGGCTCACTGCTAGGAATCCTTACCGCTATGGGTAAGATTATTAGGTT
GTCTGATCGAAAGGACAGTACTGCCTTGATGATACTGACGTAAGGGATGACGCACATGTCTGCACCTC
ACATGCTATATAAGGTTTTGCTATTCATTGAAAGCAGTAGTGACTGATTTGTATATA
>MinSyn_1882|Strength:0.011432108
GCGTGTCGTTTTAGTGAGGTGGTGGAGCACGACATGGTACTTGTGTACAATGACGTAAGCCATGACGT
CTATCACATGTAGGCTATCAGCTTAGCAAGACAGCAAGTGGATCCAATTAGGTTGTCTGCACCTCACT
GAAGATAAGATAATAATGTTGAAGATAAGACACATGTAGGCTATCAGCTTACTATATAAGGTTTTGCT
ATTCATTGAAAGCAGTAGTGACTGATTTGTATATA
>MinSyn_1783|Strength:0.01143482
GCGTGTCGTTTTAGTGAGGGCTTTGTCAAAAGCTAAAAAAGATGATGCCCTCTACAAAACTGGTACTT
GTGTACAATGAGCTAAGCACATACGTCAGCTCACATGTAGGCTATCAGCTTCAGTGGTCCCTCCACTG
TACAGGGCTCACTGCTAGGAGTTATGACCCCCGCCGATGACGCGGGAGCTAGGAGGACCGATGATCGA
AAGGACAGTAGACCACTATGCCCAATTAGGTTGTCTAACCACGTCTACAATATCAGCTTAGCAAGACC
TATATAAGGTTTTGCTATTCATTGAAAGCAGTAGTGACTGATTTGTATATA
>MinSyn_1146|Strength:0.011474438
GCGTGTCGTTTTAGTGAGGATCCTTACCGCTATGGGTAAGATTAGCTTAGCAAGACCTCTACAAAACT
GGTACAATTTCGGGAAACCTCCTCGGACCGATGCTGATCTTGCCTGGCACACCAGCATGTGTTGATCA
CCAGCTTGCCCAATTAGGTTGTCTGCACCTCACATGTTATGACCCCCGCCGATGACGCGGGACACTAT
GCCCAATTATGAAGCATCTTCCCTCTACAAAACTGGTACTTGTGTACAGATGAGCTAAGCACATACGT
CAGTTAGGTTGTCTGCAGGTGGCTCCTACCTGATCTTGCCTGCCTCACTATCAGCGGGCTCACTGCTA
CTATATAAGGTTTTGCTATTCATTGAAAGCAGTAGTGACTGATTTGTATATA
>MinSyn_1411|Strength:0.011519272
GCGTGTCGTTTTAGTGAGGAAAAATGTCAAAGATATTAGGTTGTCTGCACCTCACATGTAGGCCAGTG
GTCCCTCCACCAGGGCTCACTGCTAGGAGGACGTGGGAGCCACCAGGTCACGACCAAATTTCGGGAAA
CCTCCTCGGGACCGATGCAGCAAGTGGATTGTGTACAGGGCTCACTGATGACGTAAGCCATGACGTCT
AATGCTGATCTTGCCTGCCTTGATGATGAAGCATCTTCCTGCACCTCACATGTAGGCTCTATATAAGG
TTTTGCTATTCATTGAAAGCAGTAGTGACTGATTTGTATATA
>MinSyn_1798|Strength:0.011532113

GCGTGTCGTTTTAGTGAGGGCACACCAGCATGTGTTGATCACCAGCTTGCACCTCACCAGCCACTTGT
GTGGAATGAGCTAAGCACATACGTCAGCCGTCACTATCAGCTCTACAAATTTCGGGAAACCTCCTCGC
TAAAAATGTCAAAGATACAAGACCTCTACAAAACTGCTTTGTCAAAAGCTAAAAAAGATGATGCACTG
GTACTTGTGTACAGGGCTCACTGCTATCGAAAGGACAGTAGCACCTCACATGTAGGCTATCAGCTTCT
ATATAAGGTTTTGCTATTCATTGAAAGCAGTAGTGACTGATTTGTATATA
>MinSyn_1715|Strength:0.011553762
GCGTGTCGTTTTAGTGAGGTCCATCAACAAATAATCCAAGTAAGGCACCTCACATGTAGGCTATCAGC
TTAGCAAATATTTCTTGTAATTAGGTTGTCTGCACCTCACATATTGCGATAAAGGAAAGGTTGTGTAC
AGGGCTCACTGCTAGGAGTCACTATCAGCTCACGACCACTATGCCCAAAACCATTATTGCGGGACCGA
TGCTGATCTTGCCTGCTTATGACCCCCGCCGATGACGCGGGACAATTAGGTTGTCTGCACCAGGTGGC
TCCTACAGACCTCTACAAAACTGGTACTTGTGTACATGAGCTAAGCACATACGTCAGTCAGCTTCTAT
ATAAGGTTTTGCTATTCATTGAAAGCAGTAGTGACTGATTTGTATATA
>MinSyn_1918|Strength:0.011703536
GCGTGTCGTTTTAGTGAGGTCTCTCTGCCGACAGTGGTCCCAAAGACCTCTACAAAACTGGAAGATTG
ATGAAAAGTCAAAAACAAAAATCAATTATCCTCACATGTAGGCTATCAGCTTAAAAATGTCAAAGATA
TTAGGTTGTCTGCACCTCAAGGTGGCTCCTACTAGGCTATCAGCTTAGCAGCACACCAGCATGTGTTG
ATCACCAGCTGGTTGTCTGCACCTCACATGTCACTATCAGCACTTGTGTACATTATGACCCCCGCCGA
TGACGCGGGAACAGGGCTCACTGCTAGGAGGACCATGAGCTAAGCACATACGTCAGTGCCTGCTATAT
AAGGTTTTGCTATTCATTGAAAGCAGTAGTGACTGATTTGTATATA
>MinSyn_1698|Strength:0.011706835
GCGTGTCGTTTTAGTGAGGATCGAAAGGACAGTAGTACAGGGCTCACTGCTAGGAGGATTTTCAACAA
AGGTCACGACCAAACCATTATTGCGTTGTGTACAGGGCTCACTGCAAGATTGATGAAAAGTCAAAAAC
AAAAATCAATTATAAACTGGTACTTGTGTACAGATGACGTAAGCCATGACGTCTACCTCTACAAAACT
GGTACTTGAACCACGTCTACAACTCACTGCTAGGAGGACCGATGCTGACTATATAAGGTTTTGCTATT
CATTGAAAGCAGTAGTGACTGATTTGTATATA
>MinSyn_1590|Strength:0.011713668
GCGTGTCGTTTTAGTGAGGTCAGAAGATCAAAGGGCTATCTTGCCTGCCTTTCCATCAACAAATAATC
CAAGTAAGATTAGGTTGTCTGCACCTCACATGTAGGAAGATTGATGAAAAGTCAAAAACAAAAATCAA
TTATGTGGTGGAGCACGACAGTGTACAGGGCTCACTGCTAGGAGGGCACACCAGCATGTGTTGATCAC
CAGCTCTGGTACTTGTGTACAGGGCTCTTATGACCCCCGCCGATGACGCGGGAGGCTATCAACCACGT
CTACAAGACCGAAAAAATGTCAAAGATATGCTAATGAGCTAAGCACATACGTCAGTCTGCACCTCCTA
TATAAGGTTTTGCTATTCATTGAAAGCAGTAGTGACTGATTTGTATATA
>MinSyn_1953|Strength:0.011775806
GCGTGTCGTTTTAGTGAGGAATTTCGGGAAACCTCCTCGCTATGCCCAATTAGGTTGTCTGCGCTTTG
TCAAAAGCTAAAAAAGATGATGCCACGACCACTATGCCCAATTAATGACGTAAGCCATGACGTCTATA
CTTGTGTACAGGGCTCACTGCTAGGAATCCTTACCGCTATGGGTAAGATTTCGAACCATTATTGCGAC
TGCTAGGAGGACCGATCTATATAAGGTTTTGCTATTCATTGAAAGCAGTAGTGACTGATTTGTATATA
>MinSyn_1413|Strength:0.011776908
GCGTGTCGTTTTAGTGAGGCAGCCACTTGTGTCACATGTAGGCTATCGGAAAAAGAAGAGGTCGCGGT
AGGTCACGACCACTATGCCCAATTGCGATAAAGGAAAGGCAATTAGGTTGTCTGCATGAAGCATCTTC
CTCACTGCTAGTGAAGATAAGATAATAATGTTGAAGATAAGAACTTGTGTACAGGGCTCACTGCTAGT
GGGAGCCACCAGGAAGGTGGCTCCTACATCTTGCCTGCCTTGATGATTATGACCCCCGCCGATGACGC
GGGACTGGTATGAGCTAAGCACATACGTCAGCTCACATGTAGGCTATCAGCTTAGCAAGCTATATAAG
GTTTTGCTATTCATTGAAAGCAGTAGTGACTGATTTGTATATA
>MinSyn_1331|Strength:0.011778241
GCGTGTCGTTTTAGTGAGGTGAAGATAAGATAATAATGTTGAAGATAAGAGGGCTCACTGCTAGGAGG
ACCGAAAAAATGTCAAAGATAGCGGTAGGTCACGACCACTATGTCAGAAGATCAAAGGGCTAGCCCAA
TTAGGTTGTCTGCACACCAGCATGTGTTGATCACCAGCTGCACCTCACCAAATATTTCTTGTATGAGG
TGGCTCCTACAGCAAGACCTCTACAAAACTGGTACTTGTTGGTGGAGCACGACATAGGAGGACCGATG
CTTCACTATCAGCAAATGAGCTAAGCACATACGTCAGAGGTCACGACCACTATGCCCAACTATATAAG
GTTTTGCTATTCATTGAAAGCAGTAGTGACTGATTTGTATATA
>MinSyn_1474|Strength:0.011823565
GCGTGTCGTTTTAGTGAGGGCTTTGTCAAAAGCTAAAAAAGATGATGCGGTTCCATCAACAAATAATC
CAAGTAAGGTTTTTCAACAATAGGCTACTGACGTAAGGGATGACGCACAGAGGACCGTTATGACCCCC

GCCGATGACGCGGGACGACCACTATGCCCAATTAGGTTGTTGAAGCATCTTCCTCTACAAAACTGGTA
CTTGTGCTATATAAGGTTTTGCTATTCATTGAAAGCAGTAGTGACTGATTTGTATATA
>MinSyn_1409|Strength:0.011863318
GCGTGTCGTTTTAGTGAGGAATTTCGGGAAACCTCCTCGCAAGAATGACGTAAGCCATGACGTCTATT
AGCAAGACCTCTACAAAACTGGTAGGTGGCTCCTACATTAGGTTGTAACCACGTCTACAAACCAGCAC
ACCAGCATGTGTTGATCACCAGCTGAGGACCGATGCTGATCTTGCCTATATAAGGTTTTGCTATTCAT
TGAAAGCAGTAGTGACTGATTTGTATATA
>MinSyn_1936|Strength:0.011868161
GCGTGTCGTTTTAGTGAGGATGACGTAAGCCATGACGTCTAGAGGACCGATGCTGATCAGCAAGTGGA
TAACTGGTACTTAATTTCGGGAAACCTCCTCGGCCTGCATTGCGATAAAGGAAAGGGTCTGCACCTCA
CATGTCAGTGGTCCCTCCACTGTCTGCACCTCTATATAAGGTTTTGCTATTCATTGAAAGCAGTAGTG
ACTGATTTGTATATA
>MinSyn_1799|Strength:0.011870137
GCGTGTCGTTTTAGTGAGGAAGATTGATGAAAAGTCAAAAACAAAAATCAATTATGGACCGATGCTGA
TCTTGCCTGCCTTGACAGTGGTCCCTCCACGCTAGGAGGACCGATGCTGATCTTGCCGCACACCAGCA
TGTGTTGATCACCAGCTGCTAGGAGGACCGATGCTGATCTGACGTAAGGGATGACGCACACAGGGCTC
ACTGCTAGGAAAAAATGTCAAAGATAGTACAGGGCTCACTGCTAGGAGGCTATATAAGGTTTTGCTAT
TCATTGAAAGCAGTAGTGACTGATTTGTATATA
>MinSyn_1236|Strength:0.011890587
GCGTGTCGTTTTAGTGAGGTCTCTCTGCCGACAGTGGTCCCAAATAGGTCACGACCACTATGCGGAAA
AAGAAGAGGTAGGGCTCACTGCTAGGAGGTCCATCAACAAATAATCCAAGTAAGCATGTAGGCTATCA
GCTTAGCAAGACCTTCACTATCAGCCTGCACACCAGCATGTGTTGATCACCAGCTCAGGGCTCACTGC
TATGAGCTAAGCACATACGTCAGTGTCTGCACCTCACATGTAGGAAGATTGATGAAAAGTCAAAAACA
AAAATCAATTATTTGTCTGCACCTCAGTGGTCCCTCCACCGCGGTAGGTCACGACCACTATATAAGGT
TTTGCTATTCATTGAAAGCAGTAGTGACTGATTTGTATATA
>MinSyn_1898|Strength:0.011906125
GCGTGTCGTTTTAGTGAGGAGGTGGCTCCTACTATCCTTACCGCTATGGGTAAGATTCTGCTAGGAGA
ATTTCGGGAAACCTCCTCGGTAGGTCACGACCACTATCAGTGGTCCCTCCACTGCCCAATTAGGTTGT
CTGCACCTCACATGGCACACCAGCATGTGTTGATCACCAGCTGTAAACCACGTCTACAATGCTGATCT
TGCCTGCCTTATGAGCTAAGCACATACGTCAGTAGGAGGACCGATGCTGATCTTGCGGAAAAAGAAGA
GGTTATTATGACCCCCGCCGATGACGCGGGAGGGCTCACTGCTAGGAGGACCGATCTATATAAGGTTT
TGCTATTCATTGAAAGCAGTAGTGACTGATTTGTATATA
>MinSyn_1711|Strength:0.01194997
GCGTGTCGTTTTAGTGAGGAACCATTATTGCGCGACCACTATGCCCAATTAGGTTGTCTGCATCGAAA
GGACAGTACTGGTAACCACGTCTACAATAGGCTATCAGCTTAGCAAGACCTCTCAGCCACTTGTGTGG
CTATCAGCTTAGCAAGACCTCTACATCACTATCAGCTGGTACTTGATGAGCTAAGCACATACGTCAGT
AGGTCACGATCCATCAACAAATAATCCAAGTAAGACCTCTACAAAACTGGTACTTGTGTATCAGAAGA
TCAAAGGGCTATATCAGCTTAGCAAGACTGAAGCATCTTCCCTAGGAGGACCCTATATAAGGTTTTGC
TATTCATTGAAAGCAGTAGTGACTGATTTGTATATA
>MinSyn_19|Strength:0.011980219
GCGTGTCGTTTTAGTGAGGCTGACGTAAGGGATGACGCACATAGGTTGTCTGCACCTCACATGTAGGC
AGTGGTCCCTCCACGTAGGTCACGACCATCCATCAACAAATAATCCAAGTAAGCTTATGAAGATAAGA
TAATAATGTTGAAGATAAGAACCTCTACAAACTATATAAGGTTTTGCTATTCATTGAAAGCAGTAGTG
ACTGATTTGTATATA
>MinSyn_1601|Strength:0.012119397
GCGTGTCGTTTTAGTGAGGTCAGAAGATCAAAGGGCTACGACCACTATGCCCAATTAGGTTGTATGAG
CTAAGCACATACGTCAGGTACAGGGCTCACTGCTAGGAGGAAGATTGATGAAAAGTCAAAAACAAAAA
TCAATTATCGGTAGGTCACGACCACTATTTATGACCCCCGCCGATGACGCGGGAGTGTACAGGGCTCA
CTATCCTTACCGCTATGGGTAAGATTGTCACGACCACTAACCACGTCTACAATCTATATAAGGTTTTG
CTATTCATTGAAAGCAGTAGTGACTGATTTGTATATA
>MinSyn_1224|Strength:0.012168939
GCGTGTCGTTTTAGTGAGGATGAGCTAAGCACATACGTCAGTGCCTGCCTTGATCGAAAGGACAGTAT
CTGCACCTCATCCATCAACAAATAATCCAAGTAAGGGTACTTGTGTAAGATTGATGAAAAGTCAAAAA
CAAAAATCAATTATACAACCACGTCTACAACAAGACCTCTACAAAACTGGTAGCAAGTGGATACCACT

ATGCCCATGAAGATAAGATAATAATGTTGAAGATAAGAACACTATATAAGGTTTTGCTATTCATTGAA
AGCAGTAGTGACTGATTTGTATATA
>MinSyn_1239|Strength:0.012363831
GCGTGTCGTTTTAGTGAGGTGAAGCATCTTCCGTACTTGTGTACAGGGCTCACTGCTAGGATCCTTAC
CGCTATGGGTAAGATTTCTTTCAGAAGATCAAAGGGCTAGCACCGCTTTGTCAAAAGCTAAAAAAGAT
GATGCTCACATGTAGGCTATCAGCTTAGCAAGACCTCCATCAACAAATAATCCAAGTAAGTGTGTACA
GGGCTCATTATGACCCCCGCCGATGACGCGGGAACTTGTGTACCTGACGTAAGGGATGACGCACAATC
TTGCCTGCCTCACTATCAGCTGTGTACAGGGCTCACTGCTACTATATAAGGTTTTGCTATTCATTGAA
AGCAGTAGTGACTGATTTGTATATA
>MinSyn_1449|Strength:0.012369835
GCGTGTCGTTTTAGTGAGGTCAGAAGATCAAAGGGCTAACGACCACTATGCCCAATTAGGTATGACGT
AAGCCATGACGTCTACAGGGCTCAAAAATGTCAAAGATATACAGGGCTCACTGCTAGGAGGACCGATC
CATCAACAAATAATCCAAGTAAGTCGCGGTAGGTCACGACCACTATGCCCTATATAAGGTTTTGCTAT
TCATTGAAAGCAGTAGTGACTGATTTGTATATA
>MinSyn_1328|Strength:0.012434893
GCGTGTCGTTTTAGTGAGGTTTTCAACAATAGGCTATCAGCTTAGCGCACACCAGCATGTGTTGATCA
CCAGCTAACTGGTACTTTTATGACCCCCGCCGATGACGCGGGATGCTGATCTTGCCTGCCTTGATAAA
AATGTCAAAGATAAGGACCGATGCTGAGTGGGAGCCACCATGCCCAATTAGGTTGTCTGCACCTCATG
AGCTAAGCACATACGTCAGGGTAGGTCACGACCACTATCAAATATTTCTTGTCATCCTTACCGCTATG
GGTAAGATTCACTGCTAGGAGGACCGATGCTGATCTTCTATATAAGGTTTTGCTATTCATTGAAAGCA
GTAGTGACTGATTTGTATATA
>MinSyn_1357|Strength:0.012452321
GCGTGTCGTTTTAGTGAGGGCTTTGTCAAAAGCTAAAAAAGATGATGCGTGTACAGGGCTCACTGCTA
TGGTGGAGCACGACATTAGCAAGACCTCTACAAAACTGGTACTTTTTTCAACAAGACCGATGCTGAAA
TTTCGGGAAACCTCCTCGAACTGGTACTTGTGTACAGTCACTATCAGCGTTGAAGCATCTTCCGGACC
GATGCTGATCTTGCCTGATGAGCTAAGCACATACGTCAGTCACTGCTAGGAGGACAGGTGGCTCCTAC
TTCTCTCTGCCGACAGTGGTCCCAAACTCTACAAAACTCTATATAAGGTTTTGCTATTCATTGAAAGC
AGTAGTGACTGATTTGTATATA
>MinSyn_1185|Strength:0.01248165
GCGTGTCGTTTTAGTGAGGTTATGACCCCCGCCGATGACGCGGGATCTACAAAACTCAGTGGTCCCTC
CACTTGCCTGCCTTGCAAATATTTCTTGTAGGCTATCAGCTTAGCAAGACCTCTACAATCACTATCAG
CACGACCACTATGCCCAATTAGGTTGTCTGCTGACGTAAGGGATGACGCACAGGCTCACTGCTAGGAG
GACCGATATCGAAAGGACAGTATGCCCAATTAGGTTGCTATATAAGGTTTTGCTATTCATTGAAAGCA
GTAGTGACTGATTTGTATATA
>MinSyn_1537|Strength:0.01250223
GCGTGTCGTTTTAGTGAGGGCACACCAGCATGTGTTGATCACCAGCTCCCAATTAGGTTGTCTGCACC
TCACATTCAGAAGATCAAAGGGCTAATTAGGTTGTCTGCACCTCACATTGAAGCATCTTCCAGCAAGA
CCTGACGTAAGGGATGACGCACACTGCACCTCACATGTAGGCTTTGTCAAAAGCTAAAAAAGATGATG
CTCTGCACCTCACATGTAGGCTCTATATAAGGTTTTGCTATTCATTGAAAGCAGTAGTGACTGATTTG
TATATA
>MinSyn_1363|Strength:0.01252127
GCGTGTCGTTTTAGTGAGGTTATGACCCCCGCCGATGACGCGGGATTAGGTTCTGACGTAAGGGATGA
CGCACATATCGAAAGGACAGTATCGCGGTAGAACCATTATTGCGGGACCGATGCTGAGGTGGCTCCTA
CTCACATGTAGTCAGAAGATCAAAGGGCTAATGCTGATCTTGCCCTATATAAGGTTTTGCTATTCATT
GAAAGCAGTAGTGACTGATTTGTATATA
>MinSyn_1286|Strength:0.012570366
GCGTGTCGTTTTAGTGAGGATTGCGATAAAGGAAAGGGGTACTTCAGTGGTCCCTCCACGAGGACCGA
TGCTGATCTTGCGCACACCAGCATGTGTTGATCACCAGCTGACCTCTACAAAACTGGTGGTGGAGCAC
GACAACATGCTGACGTAAGGGATGACGCACAAAGACCTCTAAACCACGTCTACAATCGCGGTAGGAGC
AAGTGGATTACAGGGCTCACTGCTAGCTATATAAGGTTTTGCTATTCATTGAAAGCAGTAGTGACTGA
TTTGTATATA
>MinSyn_1335|Strength:0.012594406
GCGTGTCGTTTTAGTGAGGATCGAAAGGACAGTAGTAGGTCACGACCACTATGCACACCAGCATGTGT
TGATCACCAGCTTGTCTGCACCTCACACTGACGTAAGGGATGACGCACACTATCAGAAGATTGATGAA

AAGTCAAAAACAAAAATCAATTATACAAAACTGGTACTTGTGTACAGGGCTCATTTTCAACAAACTAC
TATATAAGGTTTTGCTATTCATTGAAAGCAGTAGTGACTGATTTGTATATA
>MinSyn_149|Strength:0.012631436
GCGTGTCGTTTTAGTGAGGAATTTCGGGAAACCTCCTCGCGAAACCATTATTGCGATGCTGATCTTGC
AAGATTGATGAAAAGTCAAAAACAAAAATCAATTATTGTACAGGTGGCTCCTACGAGGACCGATGCTG
ATCATGAGCTAAGCACATACGTCAGTGCTGATCTTGCCTGCCTATCGAAAGGACAGTAACAGCAAGTG
GATGGAGGACCGATGCTGATCTTGCGTGGGAGCCACCATTGTCTGCACCTCACATTTATGACCCCCGC
CGATGACGCGGGATGCACCTCACATGTAGGCTCTATATAAGGTTTTGCTATTCATTGAAAGCAGTAGT
GACTGATTTGTATATA
>MinSyn_1611|Strength:0.012666956
GCGTGTCGTTTTAGTGAGGTGAAGCATCTTCCACTGCTAGGAGGACCGATGCTGATCTAGCAAGTGGA
TGTACAGGGCTCACTGCTTTATGACCCCCGCCGATGACGCGGGAACAGGGCTCACTGCTAGGAGGAGC
ACACCAGCATGTGTTGATCACCAGCTGACTGAAGATAAGATAATAATGTTGAAGATAAGAGATTGGTG
GAGCACGACATTAGGTTGTCTGCTCCATCAACAAATAATCCAAGTAAGCTCACTGCTAGGAGGACCGA
TGCTGATCTTATGAGCTAAGCACATACGTCAGACTATATAAGGTTTTGCTATTCATTGAAAGCAGTAG
TGACTGATTTGTATATA
>MinSyn_1408|Strength:0.012671851
GCGTGTCGTTTTAGTGAGGAGCAAGTGGATTTGTTTTCAACAATCTGCACCTCACATGTAGGCTATCA
GCTCCATCAACAAATAATCCAAGTAAGGACCTCTACAAAACATTGCGATAAAGGAAAGGCTGGTACTT
GTGTACAGGGCTCACTGCTGACGTAAGGGATGACGCACAATAACCACGTCTACAATGATCTTGCGCAC
ACCAGCATGTGTTGATCACCAGCTTCACTGCTATATAAGGTTTTGCTATTCATTGAAAGCAGTAGTGA
CTGATTTGTATATA
>MinSyn_1447|Strength:0.012721913
GCGTGTCGTTTTAGTGAGGAATTTCGGGAAACCTCCTCGCACTATGCCCAATTAGGGTGGGAGCCACC
AAAAACTGGTAAAAAATGTCAAAGATAGTCTGCACCTCACATGTAGGCTATCAAGCAAGTGGATATTA
GGTTGTCTGCTGAAGCATCTTCCGCCTGCCTTGATGGCACACCAGCATGTGTTGATCACCAGCTCATG
TAGGCTATCAGCTTAGCAAGACCTCAAGATTGATGAAAAGTCAAAAACAAAAATCAATTATATGTAGC
TGACGTAAGGGATGACGCACACCTTGATGATCTATATAAGGTTTTGCTATTCATTGAAAGCAGTAGTG
ACTGATTTGTATATA
>MinSyn_1268|Strength:0.012803815
GCGTGTCGTTTTAGTGAGGAAGATTGATGAAAAGTCAAAAACAAAAATCAATTATAGGGGGAAAAAGA
AGAGGTCACTAATTGCGATAAAGGAAAGGGCGGTAGGTCACAGCAAGTGGATCAGCTTAGCTCACTAT
CAGCTGCACCTCACATGTAGGCCAAATATTTCTTGTAGGTCACGACCACTATGCCCAATTAGGTTCAG
TGGTCCCTCCACTCACGACCAAAAAATGTCAAAGATATACAAAACTGGTACTTGTGTACAGGATGACG
TAAGCCATGACGTCTAAGGACCGATGCTATATAAGGTTTTGCTATTCATTGAAAGCAGTAGTGACTGA
TTTGTATATA
>MinSyn_1163|Strength:0.012820105
GCGTGTCGTTTTAGTGAGGGGAAAAAGAAGAGGTGTTGTCTGGCTTTGTCAAAAGCTAAAAAAGATGA
TGCACTGCTAGAACCATTATTGCGAGGCTATCAGCTTAGCAAGACCTCTAATTGCGATAAAGGAAAGG
CAGGGCTCACTTCACTATCAGCAGGGCTCGCACACCAGCATGTGTTGATCACCAGCTACCTCTACAAA
ACTGGTAAGGTGGCTCCTACCCGATGCTGATCTTGCCTGCCTTGATGATGACGTAAGCCATGACGTCT
ATGGTACTTGTGTACAGGGCTCACTCTATATAAGGTTTTGCTATTCATTGAAAGCAGTAGTGACTGAT
TTGTATATA
>MinSyn_1145|Strength:0.012823351
GCGTGTCGTTTTAGTGAGGATGACGTAAGCCATGACGTCTAACCGATGCTGATCTTGCCTGCCAAGAT
TGATGAAAAGTCAAAAACAAAAATCAATTATACTATGCCCAATTATCTCTCTGCCGACAGTGGTCCCA
AATGTAGGCTATCAGCTTAGCAACTATATAAGGTTTTGCTATTCATTGAAAGCAGTAGTGACTGATTT
GTATATA
>MinSyn_1722|Strength:0.012832586
GCGTGTCGTTTTAGTGAGGGCTTTGTCAAAAGCTAAAAAAGATGATGCTGGTACTTGTGTAATTGCGA
TAAAGGAAAGGTACAGGGCTCACTGCTAGGATGAAGATAAGATAATAATGTTGAAGATAAGAAGGTTG
TCTGCACCTCAATGAGCTAAGCACATACGTCAGCACATGTAGGCTATCAGCTTAGCAAGATTATGACC
CCCGCCGATGACGCGGGACTGCACCTCACATGTAGGCTTCTCTCTGCCGACAGTGGTCCCAAACAATT
AGGTTGTCTGCACCTCACATGTAGGCTATATAAGGTTTTGCTATTCATTGAAAGCAGTAGTGACTGAT

TTGTATATA
>MinSyn_1446|Strength:0.012967068
GCGTGTCGTTTTAGTGAGGGCACACCAGCATGTGTTGATCACCAGCTGTTGTCTGCACAAGATTGATG
AAAAGTCAAAAACAAAAATCAATTATCCCAATTAGGTTGTCAACCACGTCTACAACTTGCCTTTATGA
CCCCCGCCGATGACGCGGGAACTGGTACTTGTGTACAGGGCTCAGCCACTTGTGTGCCCAATTAGGTT
GTCTCTCTCTGCCGACAGTGGTCCCAAACTATGACGTAAGCCATGACGTCTATCAGAGCAAGTGGATC
CTCACATGTAGGCTATCAGCTTAGCTATATAAGGTTTTGCTATTCATTGAAAGCAGTAGTGACTGATT
TGTATATA
>MinSyn_1466|Strength:0.012999657
GCGTGTCGTTTTAGTGAGGTTTTCAACAAGTGGTGGGAGCCACCACTGCACCTCACATGTAGGCTATC
AGCTTCAGAAGATCAAAGGGCTATACAAGGAAAAAGAAGAGGTCACTATTCCATCAACAAATAATCCA
AGTAAGTATGAGCTAAGCACATACGTCAGTCACATGTAGGCTATTTATGACCCCCGCCGATGACGCGG
GAGACCAAGGTGGCTCCTACGGTTGTCTGCACCTCACATGTAGGCTATGGTGGAGCACGACATTAGGT
TGTCTGCACCTCACATGTAGGCTATATAAGGTTTTGCTATTCATTGAAAGCAGTAGTGACTGATTTGT
ATATA
>MinSyn_1242|Strength:0.013060508
GCGTGTCGTTTTAGTGAGGTTTTCAACAATGGTACTTGGCTTTGTCAAAAGCTAAAAAAGATGATGCT
GGTACTTGTGTACTCCATCAACAAATAATCCAAGTAAGACAACCACGTCTACAAGGTCACGACCACTA
TGCCCAATTAGGTGCACACCAGCATGTGTTGATCACCAGCTAATTAGGTTGTCTGCAAACCATTATTG
CGTGATCTTGCCTGCCATGAGCTAAGCACATACGTCAGAGCTTAGCAAGACCTCTACAAAACTGGAAA
AATGTCAAAGATAGACCGATCTATATAAGGTTTTGCTATTCATTGAAAGCAGTAGTGACTGATTTGTA
TATA
>MinSyn_173|Strength:0.013075334
GCGTGTCGTTTTAGTGAGGAGCAAGTGGATTGCTAGGAGGACCGATGCTGATCTTGTGGTGGAGCACG
ACAGATCTTGTCACTATCAGCATCAGCTTAGCAAGGCTTTGTCAAAAGCTAAAAAAGATGATGCCCCA
ATTAGGTTTGAAGATAAGATAATAATGTTGAAGATAAGATATGAAAAATGTCAAAGATAACTGGTACT
TGAATTTCGGGAAACCTCCTCGCTGGGAAAAAGAAGAGGTTGTCTGCACTGACGTAAGGGATGACGCA
CATATCAGCTTAGCAAGACCTCCTATATAAGGTTTTGCTATTCATTGAAAGCAGTAGTGACTGATTTG
TATATA
>MinSyn_1763|Strength:0.013083461
GCGTGTCGTTTTAGTGAGGAACCACGTCTACAACTCACATGTAGGCTATCAGCTTAGCATGACGTAAG
CCATGACGTCTAGGCTCACTGCTAGGAGGACCGATGCTGATCGCACACCAGCATGTGTTGATCACCAG
CTAGGTCAGAAGATCAAAGGGCTATCAGCTTAGCAAGACCTCCTATATAAGGTTTTGCTATTCATTGA
AAGCAGTAGTGACTGATTTGTATATA
>MinSyn_1148|Strength:0.013099742
GCGTGTCGTTTTAGTGAGGATTGCGATAAAGGAAAGGGCTATCAGCTTAGCAAGACCTCTACAAAAAA
AAATGTCAAAGATACGGTAGGTCACGACCACTATGCCCATCCTTACCGCTATGGGTAAGATTGAGGAC
CGATGCTGATCTTGCCTGCCATGACGTAAGCCATGACGTCTACTACAAAACTGGTACAGGTGGCTCCT
ACCTGATCTTGCCTGCCTTGATGATCTATATAAGGTTTTGCTATTCATTGAAAGCAGTAGTGACTGAT
TTGTATATA
>MinSyn_1572|Strength:0.013101805
GCGTGTCGTTTTAGTGAGGGCTTTGTCAAAAGCTAAAAAAGATGATGCGTAGGCTATCAGCTAAAAAT
GTCAAAGATAGGTCACGACCACCAGCCACTTGTGTGGTTGTCTGCACCTCACATGTAGAGGTGGCTCC
TACTGTACAGGGCTCACTGCTAGGAGATGAGCTAAGCACATACGTCAGAGGACCGATGCTGATCTTTG
AAGCATCTTCCAGGACCGATTTTTCAACAATCTTGCCTGCATCGAAAGGACAGTACTGCTAGGAGGAA
TTTCGGGAAACCTCCTCGCCTATATAAGGTTTTGCTATTCATTGAAAGCAGTAGTGACTGATTTGTAT
ATA
>MinSyn_190|Strength:0.0131312
GCGTGTCGTTTTAGTGAGGAAGATTGATGAAAAGTCAAAAACAAAAATCAATTATTGTACAGGGCTCA
CTGCTAGGAGGGAAAAAGAAGAGGTAAAACTGTTTTCAACAACAGGGCTCACTGGCTTTGTCAAAAGC
TAAAAAAGATGATGCACCACTATGCCCAATTAGGTTGTCTGCACCAGCCACTTGTGTGTAGGTCACGA
CCACTATGCCCATGAGCTAAGCACATACGTCAGAGGACCGATGCTGATCTTGCCTGCCTTGTGGTGGA
GCACGACAGAGGACCGCTATATAAGGTTTTGCTATTCATTGAAAGCAGTAGTGACTGATTTGTATATA
>MinSyn_1897|Strength:0.013137379

GCGTGTCGTTTTAGTGAGGTCTCTCTGCCGACAGTGGTCCCAAAGTACAGGGCTCATCCATCAACAAA
TAATCCAAGTAAGTGCCTGCCTTGATGAACCATTATTGCGCTGATCTTGCCTGCCTTGATTTTCAACA
ATTAGGTTGTCTGCACCTCAAGCAAGTGGATGTCACGACCACTATGCCCAATTAGTCACTATCAGCGA
TGATATGAGCTAAGCACATACGTCAGTCGCGGTAGGTCACGACCACTATGCCCAAAACCACGTCTACA
AGTAGGCTATCAGCTCTATATAAGGTTTTGCTATTCATTGAAAGCAGTAGTGACTGATTTGTATATA
>MinSyn_1853|Strength:0.013171396
GCGTGTCGTTTTAGTGAGGCAGTGGTCCCTCCACAGACCTCTACAAAACTGGTACTTGTCAAATATTT
CTTGTCAAAACTGGTAAAAATGTCAAAGATAGACCACTATGCCCATCGAAAGGACAGTACTGATCTTG
CCTGCCTTTTTCAACAAGCTTAGCAAGACCTCAGCAAGTGGATACCTCTACAAGCACACCAGCATGTG
TTGATCACCAGCTTGTACAGGGCTCACTGCTAGGAGGACCGATGAGCTAAGCACATACGTCAGCTATG
CCCAATTAGGTTGTCCTATATAAGGTTTTGCTATTCATTGAAAGCAGTAGTGACTGATTTGTATATA
>MinSyn_177|Strength:0.013213225
GCGTGTCGTTTTAGTGAGGTGAAGATAAGATAATAATGTTGAAGATAAGAGAGGACCGATGCTGATCT
TGCCTTTTTCAACAAGCTGATCTTGCCTGCCTTGATGATATGACGTAAGCCATGACGTCTAGTGTACA
GGGCTGAAGCATCTTCCCTTGCCTGCTCACTATCAGCCTATCAGCTTAGCAAGACCTCTACAAAACCT
ATATAAGGTTTTGCTATTCATTGAAAGCAGTAGTGACTGATTTGTATATA
>MinSyn_1553|Strength:0.013221764
GCGTGTCGTTTTAGTGAGGAACCATTATTGCGCCACTTCCATCAACAAATAATCCAAGTAAGGATCTT
GCCTGCCTTGATCAAATATTTCTTGTTAGGCTATCAGCTTAATTTCGGGAAACCTCCTCGTATCAGCT
TATTTTCAACAATAAAAAATGTCAAAGATAGGTAGGTCACGACCACTATGCCCATGAGCTAAGCACAT
ACGTCAGTTGTCTGCACCTCACATGTAGGCTATTGAAGATAAGATAATAATGTTGAAGATAAGATGCA
CCTCACATGTAGGCTCTATATAAGGTTTTGCTATTCATTGAAAGCAGTAGTGACTGATTTGTATATA
>MinSyn_1765|Strength:0.013222297
GCGTGTCGTTTTAGTGAGGCAAATATTTCTTGTTCTTGCCTGCCTTGAAGCAAGTGGATCAGCTTCAG
CCACTTGTGTGTTGTTATGACCCCCGCCGATGACGCGGGATAGCAAGACCTCTACAAAACTGGTACTT
AAGATTGATGAAAAGTCAAAAACAAAAATCAATTATAGACCTCTACAAAAAAAAATGTCAAAGATAAT
CTTGCCTGCCTTGATGATATCCTTACCGCTATGGGTAAGATTCGGTAATGACGTAAGCCATGACGTCT
AATCAGCTTAGCAAGACCTATATAAGGTTTTGCTATTCATTGAAAGCAGTAGTGACTGATTTGTATAT
A
>MinSyn_14|Strength:0.013232946
GCGTGTCGTTTTAGTGAGGCTGACGTAAGGGATGACGCACACTAGGAGGACCGATGCTGATCTGTGGG
AGCCACCACAAAACTGGTACTTGTGTACAGGGCAGTGGTCCCTCCACTTGTTGAAGCATCTTCCTGCC
CAATTAGGTTGTCTGCACCCTATATAAGGTTTTGCTATTCATTGAAAGCAGTAGTGACTGATTTGTAT
ATA
>MinSyn_157|Strength:0.013282556
GCGTGTCGTTTTAGTGAGGCAAATATTTCTTGTTGCCTGCCTTGTCCATCAACAAATAATCCAAGTAA
GGTAGGCTATCAGCTTAGATCCTTACCGCTATGGGTAAGATTCACCTCACATGAAGCATCTTCCCTAT
CAGCTTAGCAAGACCTCTACAATTATGACCCCCGCCGATGACGCGGGACTCTACAAAACTGGTACTTG
TGTACAGGGCAGGTGGCTCCTACTATGCCCAATTAGCTGACGTAAGGGATGACGCACAGTAGGCTATC
AGCTTAGCAAGACTATATAAGGTTTTGCTATTCATTGAAAGCAGTAGTGACTGATTTGTATATA
>MinSyn_1679|Strength:0.013300194
GCGTGTCGTTTTAGTGAGGAACCATTATTGCGGGACCGATGAAGATTGATGAAAAGTCAAAAACAAAA
ATCAATTATTAGCAAGTGGATCACTGCTAGGAGGACCGATTGCGATAAAGGAAAGGGTGTACAGGGCT
CACTGCTAGGAAACCACGTCTACAATCACGACCACTATGCCCAATTAGGTTGTGAAGCATCTTCCAGA
CCTCTACAAAACATGACGTAAGCCATGACGTCTAGATCTTAGGTGGCTCCTACCACCTCACATGTAGG
CTATCAGCTTAGCTATATAAGGTTTTGCTATTCATTGAAAGCAGTAGTGACTGATTTGTATATA
>MinSyn_1204|Strength:0.013323328
GCGTGTCGTTTTAGTGAGGAAAAATGTCAAAGATAACTATGCCCATTTTCAACAATTGATGATTGGTG
GAGCACGACATCTGCACCTCACATGTAGGCTATCACTATCAGCTGCCCAATTAGGTTGGCTTTGTCAA
AAGCTAAAAAAGATGATGCCCGATGCTGAAACCATTATTGCGTATCAGCTTAGCAAGACCTCTACAAG
CAAGTGGATTACAGGGCTCACTGCTAGGATCAGAAGATCAAAGGGCTACTGCCTGACGTAAGGGATGA
CGCACACTAGGAGCTATATAAGGTTTTGCTATTCATTGAAAGCAGTAGTGACTGATTTGTATATA
>MinSyn_1604|Strength:0.013350119
GCGTGTCGTTTTAGTGAGGAACCACGTCTACAAACCTCACATGTAGCTTTGTCAAAAGCTAAAAAAGA

TGATGCGTCACGACCACTTGGTGGAGCACGACAATGAAGATAAGATAATAATGTTGAAGATAAGACGC
GGTAGGTCACGACCACTAAACCATTATTGCGTAGGTTGATCCTTACCGCTATGGGTAAGATTTAGGTC
ACGACCACTATGCCCAATTTTTCAACAAACCTGACGTAAGGGATGACGCACACTCACTGCTAGGAGGA
CCGATGCTGATCCTATATAAGGTTTTGCTATTCATTGAAAGCAGTAGTGACTGATTTGTATATA
>MinSyn_1810|Strength:0.013352419
GCGTGTCGTTTTAGTGAGGTTATGACCCCCGCCGATGACGCGGGACACCTCACATGTAGGCTGCTTTG
TCAAAAGCTAAAAAAGATGATGCACTTGTGTACAGGGCTCACTGCTAGATGACGTAAGCCATGACGTC
TACTGCCTTGATTGGTGGAGCACGACACACGACCACTATGCTCTCTCTGCCGACAGTGGTCCCAAATA
TGCCTATATAAGGTTTTGCTATTCATTGAAAGCAGTAGTGACTGATTTGTATATA
>MinSyn_1751|Strength:0.013381577
GCGTGTCGTTTTAGTGAGGCAGCCACTTGTGTCTCACATGTAGGCTATCAGCTTAGCACTGACGTAAG
GGATGACGCACAAAAAATCCTTACCGCTATGGGTAAGATTGTAGGCTATCAGCTCTCTCTGCCGACAG
TGGTCCCAAACTATCAGCTTAGCAAGACCTCTACAAAACCTATATAAGGTTTTGCTATTCATTGAAAG
CAGTAGTGACTGATTTGTATATA
>MinSyn_1885|Strength:0.013436162
GCGTGTCGTTTTAGTGAGGTGGTGGAGCACGACAACTGCTAGGAGGACCGATGCTGATCTTGCCAAAA
ATGTCAAAGATAACTATGCCCAAATTTCGGGAAACCTCCTCGGTACTTGTGTACAGGGCTCACTGCTA
GGAGAAGATTGATGAAAAGTCAAAAACAAAAATCAATTATGACCACTATGCCCAATTAGGTTGTATGA
GCTAAGCACATACGTCAGTGTACAGGGTCTCTCTGCCGACAGTGGTCCCAAAGTACTTGTGTACAGGG
CTCACTGCTCTATATAAGGTTTTGCTATTCATTGAAAGCAGTAGTGACTGATTTGTATATA
>MinSyn_1183|Strength:0.013450511
GCGTGTCGTTTTAGTGAGGATGACGTAAGCCATGACGTCTATCTGCACCTCACATGTATTTTCAACAA
CGACCACTATGCCCATTGCGATAAAGGAAAGGGCCCAATTAGGTTGTCTGCACCAGCCACTTGTGTAC
ATGTAGGCTATCAGCTTCTATATAAGGTTTTGCTATTCATTGAAAGCAGTAGTGACTGATTTGTATAT
A
>MinSyn_1544|Strength:0.013501055
GCGTGTCGTTTTAGTGAGGCAGTGGTCCCTCCACCCCAATTAGGTTGTCTGCACAAATATTTCTTGTG
CTAGGAGAATTTCGGGAAACCTCCTCGGCTGACGTAAGGGATGACGCACACTCACTGCTAGGAGGACC
GTCACTATCAGCACTTGTGTACAGGGCTCACTGCTAGGTGGCTCCTACCTTGTGTACACTATATAAGG
TTTTGCTATTCATTGAAAGCAGTAGTGACTGATTTGTATATA
>MinSyn_1925|Strength:0.013505895
GCGTGTCGTTTTAGTGAGGATTGCGATAAAGGAAAGGGCTCACTGCTTCAGAAGATCAAAGGGCTATA
CCTGACGTAAGGGATGACGCACACTGGTACTTGTGTACAAACCACGTCTACAAACAGGGCTCACTGAA
GCATCTTCCCGCAGCAAGTGGATTCAGCTTAGCAAGACCTCTACCTATATAAGGTTTTGCTATTCATT
GAAAGCAGTAGTGACTGATTTGTATATA
>MinSyn_1738|Strength:0.013548349
GCGTGTCGTTTTAGTGAGGATGAGCTAAGCACATACGTCAGTAGGAGGACCGATAGGTGGCTCCTACT
AGGCTATCAATCGAAAGGACAGTACAATTTCGGGAAACCTCCTCGGGCTATCAGCTTAGCAAGACCTC
TACATCACTATCAGCCACCTCACATGTAACCACGTCTACAATGTCAGAAGATCAAAGGGCTAAGCTTA
GCAAGACCTCTACAACTATATAAGGTTTTGCTATTCATTGAAAGCAGTAGTGACTGATTTGTATATA
>MinSyn_1301|Strength:0.013550622
GCGTGTCGTTTTAGTGAGGAAGATTGATGAAAAGTCAAAAACAAAAATCAATTATCTTGCCTGCCGGA
AAAAGAAGAGGTACAAAACTGGTACTTGTGTACAGGGCGTGGGAGCCACCAAAACTGGTACTTGTGTT
TTTCAACAACAGGGCTCACTGCTAGGAGGTCAGAAGATCAAAGGGCTAGGTAGGTCACGACCACTATC
CATCAACAAATAATCCAAGTAAGTATCAGCTTAGCAAGACATGAGCTAAGCACATACGTCAGAGCAAG
ACCTCTCTATATAAGGTTTTGCTATTCATTGAAAGCAGTAGTGACTGATTTGTATATA
>MinSyn_1835|Strength:0.013597325
GCGTGTCGTTTTAGTGAGGCAGCCACTTGTGTGCCTGCCTTGATGAATTTCGGGAAACCTCCTCGCAA
GACCTCTAACCATTATTGCGGCTTAGCAAGACCTCTACAAAACTGGGTGGGAGCCACCAAAGACCTCT
ACATGGTGGAGCACGACAACAAAACTGGTACTTGTGTACAGGGAGGTGGCTCCTACACCTCTACAAAA
CTCTGACGTAAGGGATGACGCACATTAGCAAATCGAAAGGACAGTACACGACCACTATGCCCAATTAG
GTTGCTATATAAGGTTTTGCTATTCATTGAAAGCAGTAGTGACTGATTTGTATATA
>MinSyn_1913|Strength:0.013633877
GCGTGTCGTTTTAGTGAGGTGAAGCATCTTCCACTGCTAGGAGGACCGATGCTGATCTTGGTGGAGCA

CGACATACAAAACTGGTACTTGTGTACAGGGCTCAGCCACTTGTGTGCAATTTCGGGAAACCTCCTCG
GGAGGACCGATGCTGAACCATTATTGCGCGGTAGGTCACGACCACTATGCCCAATTAATGAGCTAAGC
ACATACGTCAGCCGATGCTGATCTTGCCTGCCTCCATCAACAAATAATCCAAGTAAGGCAAGACCTCT
ACACTATATAAGGTTTTGCTATTCATTGAAAGCAGTAGTGACTGATTTGTATATA
>MinSyn_1951|Strength:0.013683728
GCGTGTCGTTTTAGTGAGGTCTCTCTGCCGACAGTGGTCCCAAAGTAGGCTATCAGCTTAGCAAGACC
TCTATCCATCAACAAATAATCCAAGTAAGCAAGACCTTTTCAACAAGTACTAAGATTGATGAAAAGTC
AAAAACAAAAATCAATTATTGCTGCTTTGTCAAAAGCTAAAAAAGATGATGCTGGTGGGAGCCACCAT
GTGATGACGTAAGCCATGACGTCTATATGCCCAATTAGGTTGTCTGTCAGAAGATCAAAGGGCTAAGG
TCACGACCTATATAAGGTTTTGCTATTCATTGAAAGCAGTAGTGACTGATTTGTATATA
>MinSyn_1365|Strength:0.013687681
GCGTGTCGTTTTAGTGAGGTCCATCAACAAATAATCCAAGTAAGGAGGACCGATGCTGATCTTGCCTG
CCTTGTCAGAAGATCAAAGGGCTAGTACTGGTGGAGCACGACAACAAAACTGGTACTTGTGTACAGGG
CTCAAAAATGTCAAAGATACAGGGCTCACTGCTAGGAGGACCGATGCTCACTATCAGCGCAGCCACTT
GTGTACCTCACATGTAGGCTATCAGCTTATGACGTAAGCCATGACGTCTAACAGGGCTCACTGCTAGG
ACTATATAAGGTTTTGCTATTCATTGAAAGCAGTAGTGACTGATTTGTATATA
>MinSyn_1305|Strength:0.01368889
GCGTGTCGTTTTAGTGAGGTCTCTCTGCCGACAGTGGTCCCAAAATTAAAGATTGATGAAAAGTCAAA
AACAAAAATCAATTATTGTGTACAGGGCTCACTGCTAGGAATTGCGATAAAGGAAAGGGTACTTGTGT
ACAGGGCTCACTATGAGCTAAGCACATACGTCAGCCACTATGCCCAATTAGGTTGTCTGAACCATTAT
TGCGCAATTAGGTTGTCTGCACTCAGAAGATCAAAGGGCTATGCCCAATTAGGTTGTCTGCACCTCAC
ACTATATAAGGTTTTGCTATTCATTGAAAGCAGTAGTGACTGATTTGTATATA
>MinSyn_1630|Strength:0.013703006
GCGTGTCGTTTTAGTGAGGAGCAAGTGGATACGACCACTATGCCCATGAAGATAAGATAATAATGTTG
AAGATAAGACTTAGCAAGACCTCAGCCACTTGTGTTATGCCCAATTAGGTTGTCTGCACCCAGTGGTC
CCTCCACTAGCAAGACCTCTACAAAAATCCTTACCGCTATGGGTAAGATTCGACCACTATGCCCATGA
GCTAAGCACATACGTCAGCACATGTAGGCTATCAGCTTAGCATCACTATCAGCGGCAAATATTTCTTG
TCGCTATATAAGGTTTTGCTATTCATTGAAAGCAGTAGTGACTGATTTGTATATA
>MinSyn_1234|Strength:0.013713806
GCGTGTCGTTTTAGTGAGGAAAAATGTCAAAGATACTTAGCAAGACCTCTACAAAACTGGTACTCAAA
TATTTCTTGTCAATTAGGTAACCATTATTGCGCTGATCTTGCCTGATGACGTAAGCCATGACGTCTAC
CCAATTAGGTTGTCTGCACCTCCAGCCACTTGTGTGCCCAATTAGGTTGTCTGCACCTCACATGCTAT
ATAAGGTTTTGCTATTCATTGAAAGCAGTAGTGACTGATTTGTATATA
>MinSyn_1973|Strength:0.013719673
GCGTGTCGTTTTAGTGAGGTTTTCAACAATTAGGTTGTGTGGGAGCCACCAACCAGTGGTCCCTCCAC
CAGCTTAGCAAGACCTCTACAAAATGACGTAAGCCATGACGTCTATTAGGTTGTCTGCACCTCACATG
TAGGTCCATCAACAAATAATCCAAGTAAGCACATGTAGGCTATCAGCTTAGCACTATATAAGGTTTTG
CTATTCATTGAAAGCAGTAGTGACTGATTTGTATATA
>MinSyn_1480|Strength:0.013776719
GCGTGTCGTTTTAGTGAGGTGAAGATAAGATAATAATGTTGAAGATAAGAAGCAAGACCTCTACAAAA
CTGAATTTCGGGAAACCTCCTCGACAGGGCTCACTGGCACACCAGCATGTGTTGATCACCAGCTTATG
CCCAATTAGGTTGTCTGCACCAAGATTGATGAAAAGTCAAAAACAAAAATCAATTATGCGGTAGGTCA
CGACCACTACTGACGTAAGGGATGACGCACAGTCACGACCACTATGGTGGAGCACGACACACCTCACA
TGCTATATAAGGTTTTGCTATTCATTGAAAGCAGTAGTGACTGATTTGTATATA
>MinSyn_1277|Strength:0.013777226
GCGTGTCGTTTTAGTGAGGATTGCGATAAAGGAAAGGTATGCCCAATTAGGTTGAATTTCGGGAAACC
TCCTCGAGGACCGATGCTGATCTTTATGACCCCCGCCGATGACGCGGGAACCACTATGCCCAGCAAGT
GGATCCCAATTTGAAGATAAGATAATAATGTTGAAGATAAGATACTTGTGTACATGAGCTAAGCACAT
ACGTCAGAAGACCTCTACAAAACTGGTCAGAAGATCAAAGGGCTAACCTCTACAAAACTGGTACTTGT
GCTATATAAGGTTTTGCTATTCATTGAAAGCAGTAGTGACTGATTTGTATATA
>MinSyn_1210|Strength:0.013786316
GCGTGTCGTTTTAGTGAGGGCTTTGTCAAAAGCTAAAAAAGATGATGCGCTATCAGCACACCAGCATG
TGTTGATCACCAGCTATCATGACGTAAGCCATGACGTCTACAATTAGGTTGTCTGCACCTCACAACCA
TTATTGCGACTTGTGTACAGGGCTTCAGAAGATCAAAGGGCTAGACCACTACTATATAAGGTTTTGCT

ATTCATTGAAAGCAGTAGTGACTGATTTGTATATA
>MinSyn_1388|Strength:0.01380264
GCGTGTCGTTTTAGTGAGGAAAAATGTCAAAGATACATGTAGGCTATCAGCTTAGCAAGACCTCTGCA
CACCAGCATGTGTTGATCACCAGCTACATGTAGGCTATCAGCAACCACGTCTACAAGATGCTGAATGA
GCTAAGCACATACGTCAGGGACCGATGCTGATCTTGCCTGCCTTGGTGGAGCACGACATGCCTGCCTT
GATGATCGAAAGGACAGTATGTACAGGGCTTTGTCAAAAGCTAAAAAAGATGATGCTGTCTGCACCTC
ACTATATAAGGTTTTGCTATTCATTGAAAGCAGTAGTGACTGATTTGTATATA
>MinSyn_1119|Strength:0.013824972
GCGTGTCGTTTTAGTGAGGAGCAAGTGGATTGCTAGGAGGACCGATGCTGATCTTGCCTGCAAATATT
TCTTGTCTTAGCAAGACCTCTACAAAACTGGTATTTTCAACAACCGATGCTGATCTTGCCTGCCTTAA
GATTGATGAAAAGTCAAAAACAAAAATCAATTATTACAAAACTGGTACTTGTGTACAGGGCTCAATGA
CGTAAGCCATGACGTCTAGCCGCTTTGTCAAAAGCTAAAAAAGATGATGCGTAGGCTATCAGCTTAGC
CTATATAAGGTTTTGCTATTCATTGAAAGCAGTAGTGACTGATTTGTATATA
>MinSyn_1255|Strength:0.013882962
GCGTGTCGTTTTAGTGAGGAAAAATGTCAAAGATAAAGACCTCTACAAAACTGGTACTTGGGAAAAAG
AAGAGGTGTGTACTCCATCAACAAATAATCCAAGTAAGTATATGAGCTAAGCACATACGTCAGGGTCA
CGACCTGAAGATAAGATAATAATGTTGAAGATAAGATTAGGTTGTCTGCACCTCATCACTATCAGCGG
GCAAATATTTCTTGTCTAGGAGGACCGATGCTGATCTTGCCTGATTGCGATAAAGGAAAGGATGATAC
TATATAAGGTTTTGCTATTCATTGAAAGCAGTAGTGACTGATTTGTATATA
>MinSyn_1132|Strength:0.013905043
GCGTGTCGTTTTAGTGAGGGCTTTGTCAAAAGCTAAAAAAGATGATGCAGGTTGTCTGCAATTGCGAT
AAAGGAAAGGTAGGTCACGACCACTCACTATCAGCAGGAGGACCGATGCTGATCTTGCCATCGAAAGG
ACAGTATAGGAGGACCGATCAGAAGATCAAAGGGCTAATTTTCAACAACGCGGTAGGTCCAGCCACTT
GTGTGGAAGATTGATGAAAAGTCAAAAACAAAAATCAATTATGCTGACGTAAGGGATGACGCACAATC
AGCTATATAAGGTTTTGCTATTCATTGAAAGCAGTAGTGACTGATTTGTATATA
>MinSyn_140|Strength:0.013906028
GCGTGTCGTTTTAGTGAGGAAAAATGTCAAAGATACATTGCGATAAAGGAAAGGGACCACTATGCCCA
ATTAGGTTGTCTAACCATTATTGCGCTATGCCCAATTACAGTGGTCCCTCCACTAGGTTGATGACGTA
AGCCATGACGTCTAATGTAGGCTATCAGGGAAAAAGAAGAGGTCTCAGAAGATCAAAGGGCTAAAACT
GGTACTATATAAGGTTTTGCTATTCATTGAAAGCAGTAGTGACTGATTTGTATATA
>MinSyn_1533|Strength:0.013946867
GCGTGTCGTTTTAGTGAGGTCACTATCAGCCGCGGTAGGTCACGACTTATGACCCCCGCCGATGACGC
GGGATTAGGTTGTCTGCACCTCACATGTAAGCAAGTGGATCTCACATGTATGAGCTAAGCACATACGT
CAGATGCTGATCTTGCCAGCCACTTGTGTTAGGTTGTCTGCACCTCACATGTAGGCTGCTTTGTCAAA
AGCTAAAAAAGATGATGCCTGGAATTTCGGGAAACCTCCTCGGTACTTGTGTACAGGGCTCACTGCTA
TATAAGGTTTTGCTATTCATTGAAAGCAGTAGTGACTGATTTGTATATA
>MinSyn_1818|Strength:0.013952648
GCGTGTCGTTTTAGTGAGGCAGCCACTTGTGTGACCTCTACAAAACTGGTACTTGTGTACAGGGAAAA
AGAAGAGGTAAGCTGACGTAAGGGATGACGCACATGATGATAGGTGGCTCCTACTAGCAAGTGGATGT
GTACAGGCAGTGGTCCCTCCACCCAATTAATCGAAAGGACAGTATGACTATATAAGGTTTTGCTATTC
ATTGAAAGCAGTAGTGACTGATTTGTATATA
>MinSyn_1792|Strength:0.013975329
GCGTGTCGTTTTAGTGAGGGCTTTGTCAAAAGCTAAAAAAGATGATGCGTGTACAGGGCTCACTGCTA
GGAGGTGGTGGAGCACGACAGGTCACGACCACTATGCCCACTGACGTAAGGGATGACGCACAGACCGA
ACCATTATTGCGGAGGAACCACGTCTACAATGCTAGGAGGACCGATGCTGATCTTGCCTGCTATATAA
GGTTTTGCTATTCATTGAAAGCAGTAGTGACTGATTTGTATATA
>MinSyn_1380|Strength:0.014018396
GCGTGTCGTTTTAGTGAGGATGAGCTAAGCACATACGTCAGTCACTCACTATCAGCAGGCTATCAGCT
TAGCAAGAAACCACGTCTACAAAAGACCTCTACAAAACTGGTACGCTTTGTCAAAAGCTAAAAAAGAT
GATGCAGACCTCTACAAAACTGGTACTTGTGTCCATCAACAAATAATCCAAGTAAGTCTGCACCTCAC
ATGTAGGCTATATAAGGTTTTGCTATTCATTGAAAGCAGTAGTGACTGATTTGTATATA
>MinSyn_1931|Strength:0.014037436
GCGTGTCGTTTTAGTGAGGAGGTGGCTCCTACTAGGTCACGACCACTAGCAAGTGGATACGACCACTA
TGCCCAATTAGGCAAATATTTCTTGTGGTCACGACCACTATGCCCAATTAATGACGTAAGCCATGACG

TCTAATGTAGGCTATCAGCTTAGCAAGAAGATTGATGAAAAGTCAAAAACAAAAATCAATTATACCTA
TATAAGGTTTTGCTATTCATTGAAAGCAGTAGTGACTGATTTGTATATA
>MinSyn_1525|Strength:0.014088714
GCGTGTCGTTTTAGTGAGGCTGACGTAAGGGATGACGCACATACTTGTGTACAGGGCTCACTGCTAGT
GAAGATAAGATAATAATGTTGAAGATAAGAGTACAGGGCTCACTTCAGAAGATCAAAGGGCTACAGCT
TAGCAAGACCTCTCTATATAAGGTTTTGCTATTCATTGAAAGCAGTAGTGACTGATTTGTATATA
>MinSyn_1519|Strength:0.014095611
GCGTGTCGTTTTAGTGAGGGTGGGAGCCACCATGCTGATTGAAGCATCTTCCGGCTATCAGCTTAGCA
AGACCTCAGGTGGCTCCTACGACCACTATGCCCTCACTATCAGCTAGGAGGACAGCCACTTGTGTTAG
CAAGACCTCTTCCATCAACAAATAATCCAAGTAAGCGCTTTTCAACAAACAGGGCTCACTAACCACGT
CTACAACTATGCCCAATTAGGTTGTCTGCACCTCATGACGTAAGCCATGACGTCTACTACACTATATA
AGGTTTTGCTATTCATTGAAAGCAGTAGTGACTGATTTGTATATA
>MinSyn_1272|Strength:0.014104787
GCGTGTCGTTTTAGTGAGGCAGCCACTTGTGTAATTAGGTTGTCTGCACCAAATATTTCTTGTGGTAC
TTGTGTACAGGGCTCATTTTCAACAAGCTACAGTGGTCCCTCCACTCTGCACCTCATGACGTAAGCCA
TGACGTCTACTCACTGCTAGGAGGACCGATGAACCATTATTGCGCAAGACCTCTACAAAACTGGTACT
ATATAAGGTTTTGCTATTCATTGAAAGCAGTAGTGACTGATTTGTATATA
>MinSyn_1203|Strength:0.014135392
GCGTGTCGTTTTAGTGAGGGGAAAAAGAAGAGGTTGATCTTGCCTGCCTTGATGATATGACGTAAGCC
ATGACGTCTATGCTATCCTTACCGCTATGGGTAAGATTCAGCTTAGCAAGACCTCTCACTATCAGCCA
CGACCACTATGCCCAAGGTGGCTCCTACCTCTCTATATAAGGTTTTGCTATTCATTGAAAGCAGTAGT
GACTGATTTGTATATA
>MinSyn_1336|Strength:0.014141353
GCGTGTCGTTTTAGTGAGGATTGCGATAAAGGAAAGGGGTAGGTCACGACCACTATGCCCAATTAGAA
CCACGTCTACAAGTACTTGTGTACAGGGCTCACTGCTATGAGCTAAGCACATACGTCAGTTGTCTGCA
CCTCACATGTAGGCTATCATCTCTCTGCCGACAGTGGTCCCAAATACAAAACTGGTACTTGTGTACAG
CCACTTGTGTTGTGGGAGCCACCATCAGCTTAGCAAAAAATGTCAAAGATAGTAGGTCACCTATATAA
GGTTTTGCTATTCATTGAAAGCAGTAGTGACTGATTTGTATATA
>MinSyn_1141|Strength:0.014145385
GCGTGTCGTTTTAGTGAGGTGAAGATAAGATAATAATGTTGAAGATAAGATGTCTGCACCTCACATGT
AGAGGTGGCTCCTACTTGATAGCAAGTGGATGCCTGCCTTGATGATCCTTACCGCTATGGGTAAGATT
CGACCACTATGCCCAATCGAAAGGACAGTATATGCCCAATTAGGTTGTCTCTCTGCCGACAGTGGTCC
CAAAGCTCACATGAGCTAAGCACATACGTCAGACTTGTGTACAGGGCTCACTGCTAGGAGCTATATAA
GGTTTTGCTATTCATTGAAAGCAGTAGTGACTGATTTGTATATA
>MinSyn_1920|Strength:0.014159408
GCGTGTCGTTTTAGTGAGGCAGTGGTCCCTCCACGGAGGATTATGACCCCCGCCGATGACGCGGGATA
CAGGGCTCACTGCTAGGAGGACCAACCACGTCTACAACAATTAGGTTGTCCTGACGTAAGGGATGACG
CACATGTAGGCTATCAGCTTAGCAAGATCGAAAGGACAGTAGATCTTGCCTGCCTTGATGATACTATA
TAAGGTTTTGCTATTCATTGAAAGCAGTAGTGACTGATTTGTATATA
>MinSyn_1830|Strength:0.014219067
GCGTGTCGTTTTAGTGAGGATGAGCTAAGCACATACGTCAGACAAAACTGGTACTTGTGATCGAAAGG
ACAGTACGACCAAGATTGATGAAAAGTCAAAAACAAAAATCAATTATAGGTCACGACCATTGCGATAA
AGGAAAGGCACTGCTAGGAGGACCGATGCTGATCTTGCAACCACGTCTACAAAGGGCTCACTGCTAGG
AGGACTATATAAGGTTTTGCTATTCATTGAAAGCAGTAGTGACTGATTTGTATATA
>MinSyn_1287|Strength:0.014227403
GCGTGTCGTTTTAGTGAGGAATTTCGGGAAACCTCCTCGGGTCACGACCATGGTGGAGCACGACAGGT
CACGACCACTATGCCCAATTAGGCAAATATTTCTTGTGAGGACCGATGCTGATCATCCTTACCGCTAT
GGGTAAGATTCAATTAGGTTGTCTGCACATGAGCTAAGCACATACGTCAGCTCTACTCTCTCTGCCGA
CAGTGGTCCCAAATCACTGCTAGGAGGACCGATTTTCAACAAGGACCGATGCTGATCTCTATATAAGG
TTTTGCTATTCATTGAAAGCAGTAGTGACTGATTTGTATATA
>MinSyn_138|Strength:0.014340143
GCGTGTCGTTTTAGTGAGGGTGGGAGCCACCATACAGGGCTCACTGCTAGGAGAAAAATGTCAAAGAT
ATCTGCACCTCACATGTAGGCTATCATCTCTCTGCCGACAGTGGTCCCAAATGTCTGCTGAAGATAAG
ATAATAATGTTGAAGATAAGACAAGACCTCTACAAAACTATGAGCTAAGCACATACGTCAGCGGTAGG

TCACGACCACTATGCCCAAGATTGATGAAAAGTCAAAAACAAAAATCAATTATAGGTCTATATAAGGT
TTTGCTATTCATTGAAAGCAGTAGTGACTGATTTGTATATA
>MinSyn_1728|Strength:0.014349293
GCGTGTCGTTTTAGTGAGGATGAGCTAAGCACATACGTCAGCCTGCCTTGATGATAACCATTATTGCG
TTGTGTACAGGGCTCATGAAGCATCTTCCCACTGCTAGGGGAAAAAGAAGAGGTGGCTCAGCAAGTGG
ATTACTTGTGTACAGGGCTCACTGCTCAGCCACTTGTGTGCGGTAGGTCACGACCACTATGCCCAATT
ACTATATAAGGTTTTGCTATTCATTGAAAGCAGTAGTGACTGATTTGTATATA
>MinSyn_1633|Strength:0.014367986
GCGTGTCGTTTTAGTGAGGATCGAAAGGACAGTAATGCCCAATTAGGTTAAAAATGTCAAAGATACAC
CTCACATGTAGGCTATGAAGATAAGATAATAATGTTGAAGATAAGATGGTACTTCTCTCTGCCGACAG
TGGTCCCAAATAGGTTGTGTGGGAGCCACCACTATGCCCAATTAGGTTGTCTGCACCTCACAGCCACT
TGTGTGATGCTGATCTTGCCTGCCATGACGTAAGCCATGACGTCTAGTACTTGTGCTATATAAGGTTT
TGCTATTCATTGAAAGCAGTAGTGACTGATTTGTATATA
>MinSyn_1417|Strength:0.014377533
GCGTGTCGTTTTAGTGAGGATCGAAAGGACAGTACTCATGAGCTAAGCACATACGTCAGTGCACCTCA
CATGTAGGCTATCAGGCTTTGTCAAAAGCTAAAAAAGATGATGCGGCTTGAAGCATCTTCCTAGGATT
GCGATAAAGGAAAGGGCTTAGCAAGACCTCTACAAAACTGAAGATAAGATAATAATGTTGAAGATAAG
AGCACCTCCTATATAAGGTTTTGCTATTCATTGAAAGCAGTAGTGACTGATTTGTATATA
>MinSyn_1184|Strength:0.014386012
GCGTGTCGTTTTAGTGAGGAACCATTATTGCGACATGTTCCATCAACAAATAATCCAAGTAAGAACAA
ATATTTCTTGTTTGGGAAAAAGAAGAGGTGACCTCTCAGAAGATCAAAGGGCTAACCACTATGCCCAA
AAAAATGTCAAAGATACCACTATGCCAGCCACTTGTGTCGATGCTGATCGCACACCAGCATGTGTTGA
TCACCAGCTGAGGACCGATGCTGATCTTGCCTGCATGACGTAAGCCATGACGTCTATCTATATAAGGT
TTTGCTATTCATTGAAAGCAGTAGTGACTGATTTGTATATA
>MinSyn_1615|Strength:0.014397024
GCGTGTCGTTTTAGTGAGGATCCTTACCGCTATGGGTAAGATTCCAATTAGGTTGTCTGCACCTCACA
TGTGAAGATAAGATAATAATGTTGAAGATAAGAACTTGTGTACAGGGGGAAAAAGAAGAGGTGCCCAA
TTAGGTTGTCTGCAGGTGGCTCCTACCCTGCCTTGATAACCACGTCTACAATATCAGCTTAGCAATTT
TCAACAACTGCACCTCACATGATGAGCTAAGCACATACGTCAGAGCTTAGCAAGCTATATAAGGTTTT
GCTATTCATTGAAAGCAGTAGTGACTGATTTGTATATA
>MinSyn_1991|Strength:0.014442195
GCGTGTCGTTTTAGTGAGGCTGACGTAAGGGATGACGCACATCGCGGTAGGTCTGAAGCATCTTCCGG
AGGACCGATGCTGATCTTGCCTGCCTTTATGACCCCCGCCGATGACGCGGGAACCACTATGCCCAATT
AGGTTGTCTGCTATATAAGGTTTTGCTATTCATTGAAAGCAGTAGTGACTGATTTGTATATA
>MinSyn_1316|Strength:0.014527935
GCGTGTCGTTTTAGTGAGGTGAAGATAAGATAATAATGTTGAAGATAAGAGGTCACGACCACTATGTG
AAGCATCTTCCATTAGGTTGTCTGCACCTCAATTTCGGGAAACCTCCTCGTAGGCTATCCTGACGTAA
GGGATGACGCACATGCACCTCACATGTATCAGAAGATCAAAGGGCTATAGGCTATCAGCTTAGCACTA
TATAAGGTTTTGCTATTCATTGAAAGCAGTAGTGACTGATTTGTATATA
>MinSyn_1506|Strength:0.014569141
GCGTGTCGTTTTAGTGAGGGTGGGAGCCACCATCAGCTTAGCAAGACCTCTACAAAACTGGATCCTTA
CCGCTATGGGTAAGATTTATGAGCTAAGCACATACGTCAGGGAGGACCGATGCTGATCTTGCCTGCCT
TGAGCAAGTGGATTTAGGTTGTCTGCACCTCAACCACGTCTACAAATTAGGTTGTCTGCACCTTTTTC
AACAATTAGGTTCACTATCAGCTGGTACTTGTGTACAGGGCTCACTGCCTATATAAGGTTTTGCTATT
CATTGAAAGCAGTAGTGACTGATTTGTATATA
>MinSyn_196|Strength:0.014602695
GCGTGTCGTTTTAGTGAGGATGAGCTAAGCACATACGTCAGCAATTAGGTTGTCTGTGGGAGCCACCA
TAGGTCACGACCACGCACACCAGCATGTGTTGATCACCAGCTGTACTTGTGTACAGGGCTCAATTTCG
GGAAACCTCCTCGTAGGCTATCAGCTTAGCATGAAGATAAGATAATAATGTTGAAGATAAGAGGTCAC
CTATATAAGGTTTTGCTATTCATTGAAAGCAGTAGTGACTGATTTGTATATA
>MinSyn_1957|Strength:0.014705508
GCGTGTCGTTTTAGTGAGGATGACGTAAGCCATGACGTCTACACCTCACATGTAGGCTATCAGCTTAG
CACAGTGGTCCCTCCACTGATCTTGCCTGCAATTTCGGGAAACCTCCTCGCAAGACCTCTACAAAACT
GGTACTTGCTATATAAGGTTTTGCTATTCATTGAAAGCAGTAGTGACTGATTTGTATATA

>MinSyn_1265|Strength:0.01472211
GCGTGTCGTTTTAGTGAGGGTGGGAGCCACCAGACCACTATGCCCAATTAGGTTAGGTGGCTCCTACA
GGTCACTGACGTAAGGGATGACGCACATGTAGGCTATCAGCTTAGCAAGACCTCTCAGAAGATCAAAG
GGCTAAGGAGGACCGATGCTGATCTTGCCTGCCTTCTATATAAGGTTTTGCTATTCATTGAAAGCAGT
AGTGACTGATTTGTATATA
>MinSyn_1539|Strength:0.014761855
GCGTGTCGTTTTAGTGAGGGTGGGAGCCACCAGCACCTCACATGTAAATTTCGGGAAACCTCCTCGAA
TGGTGGAGCACGACATATCAGCTTAGCAAGACAACCACGTCTACAATACTTGTGTACAGGGCTCACTG
CTAGGAGGAGCAAGTGGATATGCCCAATTAGGTTGTCTGCACCTAACCATTATTGCGCTGCACCTCAC
ATGTACTGACGTAAGGGATGACGCACACACGACCACTATGCCCACTATATAAGGTTTTGCTATTCATT
GAAAGCAGTAGTGACTGATTTGTATATA
>MinSyn_1699|Strength:0.014773544
GCGTGTCGTTTTAGTGAGGTTATGACCCCCGCCGATGACGCGGGATACAAAACTGGTACCTGACGTAA
GGGATGACGCACATACAGGGCTCACTGCTAGGAGGACCGACAGCCACTTGTGTGCTATTGAAGCATCT
TCCGAACCACGTCTACAAGCTAGGAGGACCTATATAAGGTTTTGCTATTCATTGAAAGCAGTAGTGAC
TGATTTGTATATA
>MinSyn_160|Strength:0.014783509
GCGTGTCGTTTTAGTGAGGCAGCCACTTGTGTGTGTACAGGGCTCACTGCTAGGAAAAAGAAGAGGTA
CTGCTAGGAGGACATTGCGATAAAGGAAAGGGCTTAGCAAGACCTCTACAAGGTGGCTCCTACTATGC
CCAATTAGGTTGTCTGCACATGAGCTAAGCACATACGTCAGACGTCAGAAGATCAAAGGGCTACCTCT
ACAAAACAAATATTTCTTGTGGTAGGTCACGACCACTATGCCCAACTATATAAGGTTTTGCTATTCAT
TGAAAGCAGTAGTGACTGATTTGTATATA
>MinSyn_1448|Strength:0.014817218
GCGTGTCGTTTTAGTGAGGAACCATTATTGCGGCTATCAGCTTAGCAAGTGAAGCATCTTCCTGTACA
GGGCTCACTGCTAGGAGGACAACCACGTCTACAACTTAGCAAGACCTCTACAAAATCTCTCTGCCGAC
AGTGGTCCCAAAACCGATGTGGTGGAGCACGACAACCGATGCTGATCTATGACGTAAGCCATGACGTC
TACTAGGAGGACCGATAAAAATGTCAAAGATAATGCTGATCTTGCCTATATAAGGTTTTGCTATTCAT
TGAAAGCAGTAGTGACTGATTTGTATATA
>MinSyn_1862|Strength:0.014823299
GCGTGTCGTTTTAGTGAGGTTATGACCCCCGCCGATGACGCGGGAGCTATTCTCTCTGCCGACAGTGG
TCCCAAACGACCACTATGCCCAATTAGGTTGTGGGAGCCACCAGTCTGCACCTCACATGTAGGCTACA
GCCACTTGTGTACCTCTACAAAATTTCGGGAAACCTCCTCGACCAAAAATGTCAAAGATAGACCGACT
GACGTAAGGGATGACGCACATGTACAGGGCTCACTGCTAGGAGGACTATATAAGGTTTTGCTATTCAT
TGAAAGCAGTAGTGACTGATTTGTATATA
>MinSyn_1186|Strength:0.014991629
GCGTGTCGTTTTAGTGAGGTCACTATCAGCGAGGACCGATGCTGTGAAGATAAGATAATAATGTTGAA
GATAAGAACAAAACTGGTACTTGTGTACAGGAACCACGTCTACAATCAGCTTAGCAATGAGCTAAGCA
CATACGTCAGAGGACCGATGCTGATCTTGCCTGCCTTGATTTTCAACAAGGCTCTCTCTCTGCCGACA
GTGGTCCCAAACATGTAGGGGAAAAAGAAGAGGTTCACTGCTCTATATAAGGTTTTGCTATTCATTGA
AAGCAGTAGTGACTGATTTGTATATA
>MinSyn_131|Strength:0.015013369
GCGTGTCGTTTTAGTGAGGTCACTATCAGCCTGGTACTTAAGATTGATGAAAAGTCAAAAACAAAAAT
CAATTATCTATGCCCAATTAGGTTGTCTGCACCTTATGACCCCCGCCGATGACGCGGGAGTCTGATGA
GCTAAGCACATACGTCAGGTGTACAGGGCTCTCTCTGCCGACAGTGGTCCCAAACGCGGTAGGTCACG
ACCACTATGCCCAATTGGAAAAAGAAGAGGTTCACGACCACCTATATAAGGTTTTGCTATTCATTGAA
AGCAGTAGTGACTGATTTGTATATA
>MinSyn_1961|Strength:0.015175171
GCGTGTCGTTTTAGTGAGGATCCTTACCGCTATGGGTAAGATTCGACCACTATGCCCAAAGGTGGCTC
CTACGCCTGCCTTGATATTGCGATAAAGGAAAGGTAGGAGGACCGATGCTGATCTTGCCTGAACCACG
TCTACAAATGCCCAAATTTCGGGAAACCTCCTCGACTGGTACTTGTGTGGTGGAGCACGACAGTACTA
TGAGCTAAGCACATACGTCAGCCAATTAGGTTGTCTGCCTATATAAGGTTTTGCTATTCATTGAAAGC
AGTAGTGACTGATTTGTATATA
>MinSyn_1585|Strength:0.015187179
GCGTGTCGTTTTAGTGAGGAACCACGTCTACAAATGTAGGCTTCTCTCTGCCGACAGTGGTCCCAAAA

CGACCACTATGCCCAATTAGGTTGTCTGCATGAGCTAAGCACATACGTCAGGTAGGCTATCAGCTTGC
ACACCAGCATGTGTTGATCACCAGCTACCTCACATGTAGGCTATCAGCAATTTCGGGAAACCTCCTCG
CTAGGAGGACCGATGCTAGCAAGTGGATAATTAGGTTCTATATAAGGTTTTGCTATTCATTGAAAGCA
GTAGTGACTGATTTGTATATA
>MinSyn_1262|Strength:0.015385827
GCGTGTCGTTTTAGTGAGGATCGAAAGGACAGTACGCGGTAGGTCACGACCACAACCATTATTGCGTC
TTGCCTGCCTATGACGTAAGCCATGACGTCTAGGGCTCACTCAGTGGTCCCTCCACCACGACCATCAC
TATCAGCCTTGTGTACAGGGCTCACTGCTAGGACTATATAAGGTTTTGCTATTCATTGAAAGCAGTAG
TGACTGATTTGTATATA
>MinSyn_1368|Strength:0.015405703
GCGTGTCGTTTTAGTGAGGAACCACGTCTACAAGACCTATGACGTAAGCCATGACGTCTAACGACCAC
TATGCCCAATTAGGTTGTAAGATTGATGAAAAGTCAAAAACAAAAATCAATTATTTCTCTCTGCCGAC
AGTGGTCCCAAACTGCTATATAAGGTTTTGCTATTCATTGAAAGCAGTAGTGACTGATTTGTATATA
>MinSyn_1887|Strength:0.015410324
GCGTGTCGTTTTAGTGAGGAAAAATGTCAAAGATAATGCCCAATTAGGTTGTCTGCACCTCATGACGT
AAGCCATGACGTCTAGTACAGGGCTCACTGCTAGGTTTTCAACAACTACAAACCACGTCTACAATGTA
CAGGGCTCACTGCTAGGAGGACCGCTATATAAGGTTTTGCTATTCATTGAAAGCAGTAGTGACTGATT
TGTATATA
>MinSyn_1599|Strength:0.015477867
GCGTGTCGTTTTAGTGAGGTGAAGATAAGATAATAATGTTGAAGATAAGAGGTAGGTCACGACCATGA
CGTAAGCCATGACGTCTAGGTCAAATATTTCTTGTCTTGCCTGCCTTCTCTCTGCCGACAGTGGTCCC
AAAACATGTAGGCTATCAGCTTAGCACTATATAAGGTTTTGCTATTCATTGAAAGCAGTAGTGACTGA
TTTGTATATA
>MinSyn_1673|Strength:0.015492488
GCGTGTCGTTTTAGTGAGGTGAAGCATCTTCCTGCTAGGAGGACCGATGCTGATCTTGGCACACCAGC
ATGTGTTGATCACCAGCTCCTCTACAAAACTGGTAAGCAAGTGGATCCACTATGCCCTGGTGGAGCAC
GACATCGCGGTAGGTCACGACCACTAAAAATGTCAAAGATAGACCACTATGCCCAATTAGGTTGTCTG
CACCTGACGTAAGGGATGACGCACAGCCTTCTATATAAGGTTTTGCTATTCATTGAAAGCAGTAGTGA
CTGATTTGTATATA
>MinSyn_1545|Strength:0.015569373
GCGTGTCGTTTTAGTGAGGTGGTGGAGCACGACACGACCACTATGCCCAAACCATTATTGCGGTAGGC
TATCAGCTTAGCAAGACCTCTACGTGGGAGCCACCAGCTAATGACGTAAGCCATGACGTCTACTGCTA
GGAGGCAGTGGTCCCTCCACACATGTAGGCTATCAGCTTAGCAAGACTATATAAGGTTTTGCTATTCA
TTGAAAGCAGTAGTGACTGATTTGTATATA
>MinSyn_1529|Strength:0.015641746
GCGTGTCGTTTTAGTGAGGAGCAAGTGGATTTGTGTACAAAGATTGATGAAAAGTCAAAAACAAAAAT
CAATTATCATCCTTACCGCTATGGGTAAGATTCAGCTTAGCAAGACCTCTTGGTGGAGCACGACAAGG
CTAATTTCGGGAAACCTCCTCGAGGCTATCAGCTTAGCAAGACCTCTAATGACGTAAGCCATGACGTC
TACAGGGCTCACCAGTGGTCCCTCCACGGTCTATATAAGGTTTTGCTATTCATTGAAAGCAGTAGTGA
CTGATTTGTATATA
>MinSyn_1509|Strength:0.015732682
GCGTGTCGTTTTAGTGAGGTTATGACCCCCGCCGATGACGCGGGAGGTAGGTCACGACATTGCGATAA
AGGAAAGGATGCCCAATTAGGTTGTCTGCACCTCACATATGACGTAAGCCATGACGTCTAGTAGGCTA
TAAGATTGATGAAAAGTCAAAAACAAAAATCAATTATACCACTATCTATATAAGGTTTTGCTATTCAT
TGAAAGCAGTAGTGACTGATTTGTATATA
>MinSyn_1362|Strength:0.015743414
GCGTGTCGTTTTAGTGAGGAAGATTGATGAAAAGTCAAAAACAAAAATCAATTATACGAACCACGTCT
ACAAGAGGACCGAGCAAGTGGATGTAGGTCACGACCACTAATGACGTAAGCCATGACGTCTACTACAA
AACTGGTACTTGTGTACACAGCCACTTGTGTCACATGTAGGCTATCTATATAAGGTTTTGCTATTCAT
TGAAAGCAGTAGTGACTGATTTGTATATA
>MinSyn_1990|Strength:0.015832882
GCGTGTCGTTTTAGTGAGGGGAAAAAGAAGAGGTGAGGACCGATGCTGAAGGTGGCTCCTACGGTATT
GCGATAAAGGAAAGGCTACATCCATCAACAAATAATCCAAGTAAGTCAGCTTAGCAAGAAGCAAGTGG
ATTTGAAGCATCTTCCTCGCGGTAGGTCACGACCACTATGATGAGCTAAGCACATACGTCAGTATGCC

CAATTAGGTTGTCTGCACCTCACACTATATAAGGTTTTGCTATTCATTGAAAGCAGTAGTGACTGATT
TGTATATA
>MinSyn_1100|Strength:0.015856441
GCGTGTCGTTTTAGTGAGGAGCAAGTGGATTGATCCATCAACAAATAATCCAAGTAAGGGCTCACTGC
TAGGAGGAGGAAAAAGAAGAGGTATGTAGGCTATCAGCTTAGCAAGACCTCTGAAGATAAGATAATAA
TGTTGAAGATAAGAGGCTATCAGGTGGGAGCCACCAGCACCTCACATGTAGGCTACTGACGTAAGGGA
TGACGCACACCTCACATGTAGGCTCTATATAAGGTTTTGCTATTCATTGAAAGCAGTAGTGACTGATT
TGTATATA
>MinSyn_1299|Strength:0.015996178
GCGTGTCGTTTTAGTGAGGAACCATTATTGCGTGCTGATCTTGCCAGGTGGCTCCTACCAGCTTAGCA
AGACCTCTACAAAACAAAAATGTCAAAGATAAAACTGGTCAGTGGTCCCTCCACAACTGGTGAAGCAT
CTTCCGCTATCAGCTTAGCAAGACCTCAGAAGATCAAAGGGCTAAGGAATGACGTAAGCCATGACGTC
TAAACTGGTACTTGTGTACAGGCTATATAAGGTTTTGCTATTCATTGAAAGCAGTAGTGACTGATTTG
TATATA
>MinSyn_1742|Strength:0.01604858
GCGTGTCGTTTTAGTGAGGGGAAAAAGAAGAGGTCACATGTAGGTGGGAGCCACCACACCTCACATGT
AGGCTATCAGCTTAGCATGACGTAAGCCATGACGTCTATACTTGTGTACAGGGCTCACTGCTAGGAGA
GGTGGCTCCTACGTCATGAAGCATCTTCCGTCTATATAAGGTTTTGCTATTCATTGAAAGCAGTAGTG
ACTGATTTGTATATA
>MinSyn_1416|Strength:0.01612076
GCGTGTCGTTTTAGTGAGGCTGACGTAAGGGATGACGCACAGTGTACAGGGCACACCAGCATGTGTTG
ATCACCAGCTGATTGCGATAAAGGAAAGGGGTTGTCTGCACCTCACATGTCAGCCACTTGTGTCAATT
CTATATAAGGTTTTGCTATTCATTGAAAGCAGTAGTGACTGATTTGTATATA
>MinSyn_1524|Strength:0.016263314
GCGTGTCGTTTTAGTGAGGGCACACCAGCATGTGTTGATCACCAGCTGTAGGTCACGACCACTATGCC
CAATTAGGTGGAAAAAGAAGAGGTGTACTTGTGTACAGGGCTTCAGAAGATCAAAGGGCTAAAGACCT
CTACAAAACTGGTACTAACCATTATTGCGTCACATATGAGCTAAGCACATACGTCAGAGGTTGTCAAT
TTCGGGAAACCTCCTCGCCTATATAAGGTTTTGCTATTCATTGAAAGCAGTAGTGACTGATTTGTATA
TA
>MinSyn_1202|Strength:0.016280051
GCGTGTCGTTTTAGTGAGGGCACACCAGCATGTGTTGATCACCAGCTCTCACTGCTCTGACGTAAGGG
ATGACGCACAAGGTTGTCTGCACCTCACATGTAGGCTATCACTATCAGCAATTAGGTTGTCTGCGGAA
AAAGAAGAGGTCACGACTATATAAGGTTTTGCTATTCATTGAAAGCAGTAGTGACTGATTTGTATATA
>MinSyn_1652|Strength:0.01636562
GCGTGTCGTTTTAGTGAGGAAAAATGTCAAAGATACACGACCCAGTGGTCCCTCCACGATCCATCAAC
AAATAATCCAAGTAAGTCTTGCCTGCCTTGATGGCTTTGTCAAAAGCTAAAAAAGATGATGCAGACCA
TGAGCTAAGCACATACGTCAGACTATGCCCAATTAGGTTGGAAAAAGAAGAGGTAAACCACGTCTACA
ATGATCACTATCAGCACCCTATATAAGGTTTTGCTATTCATTGAAAGCAGTAGTGACTGATTTGTATA
TA
>MinSyn_1398|Strength:0.01638099
GCGTGTCGTTTTAGTGAGGAATTTCGGGAAACCTCCTCGGTAGGCTATCAGCTTAGCAAGACCTTCTC
TCTGCCGACAGTGGTCCCAAAGGGCTCACTGCTAGGAGGACCGATGCTGATTTATGACCCCCGCCGAT
GACGCGGGACGCGATGACGTAAGCCATGACGTCTATCGCGGTAGGTCACGACCCAAATATTTCTTGTT
AGGTCACGACCACTACTATATAAGGTTTTGCTATTCATTGAAAGCAGTAGTGACTGATTTGTATATA
>MinSyn_1394|Strength:0.016388453
GCGTGTCGTTTTAGTGAGGTTATGACCCCCGCCGATGACGCGGGACCAATTAGCAGCCACTTGTGTAC
CACTATCGAAAGGACAGTACAAAACTGGTACTTGTGTACAGGGAACCACGTCTACAACTGCCTTGTTT
TCAACAATGCTAGGAGGACCATCCTTACCGCTATGGGTAAGATTCTCACATGACGTAAGCCATGACGT
CTAGACCTCTACAAAACTATATAAGGTTTTGCTATTCATTGAAAGCAGTAGTGACTGATTTGTATATA
>MinSyn_1777|Strength:0.016472304
GCGTGTCGTTTTAGTGAGGGCACACCAGCATGTGTTGATCACCAGCTACATGTAGGCTATCAGCTAAC
CACGTCTACAATTAGGTTGTCTGCACCTCACATGAGGTGGCTCCTACGTCACGACCACTATGCCCAAT
TAGGGTGGGAGCCACCACTTGTGCTGACGTAAGGGATGACGCACACCGATGCTGATCTCAGCCACTTG
TGTGGTCACGACCCTATATAAGGTTTTGCTATTCATTGAAAGCAGTAGTGACTGATTTGTATATA

>MinSyn_1171|Strength:0.016488914
GCGTGTCGTTTTAGTGAGGTTTTCAACAATAGGTTGTCTGCACCTAAAAATGTCAAAGATAGGAGGAC
CGATGCTGATCTTGCAGCAAGTGGATGGTTGTCTGCACTCTCTCTGCCGACAGTGGTCCCAAAATAAC
CATTATTGCGTACTTGTGTACAGGGCTCACTCTGACGTAAGGGATGACGCACAGGACCGATGCTGATC
TTGCCTGCCTTGCTATATAAGGTTTTGCTATTCATTGAAAGCAGTAGTGACTGATTTGTATATA
>MinSyn_1252|Strength:0.016507638
GCGTGTCGTTTTAGTGAGGAAAAATGTCAAAGATACAAGACCTCTACAGTGGTCCCTCCACTCTACAA
AACTGGTACTTGTGTACAGGGCATCGAAAGGACAGTATGCTGATCTTGCACACCAGCATGTGTTGATC
ACCAGCTCAAGACCTCTACAAAACTGGTACTTCTGACGTAAGGGATGACGCACATAAACCATTATTGC
GCTTGCCTGCCTTCTATATAAGGTTTTGCTATTCATTGAAAGCAGTAGTGACTGATTTGTATATA
>MinSyn_1749|Strength:0.016581818
GCGTGTCGTTTTAGTGAGGAGGTGGCTCCTACAAGACCTGGTGGAGCACGACAGAGGACCGATGCTGA
TCTTGCCTGCCTTAACCATTATTGCGGGTCACGACCACTATGAACCACGTCTACAAACTTGTGGTGGG
AGCCACCAGCATCAGAAGATCAAAGGGCTAGTACTTGTGTACAGGGCTCAATGACGTAAGCCATGACG
TCTAGCTGATCTCTATATAAGGTTTTGCTATTCATTGAAAGCAGTAGTGACTGATTTGTATATA
>MinSyn_1312|Strength:0.016800029
GCGTGTCGTTTTAGTGAGGATGACGTAAGCCATGACGTCTATGCCCAATTAGGTTAAAAATGTCAAAG
ATAATTATGACCCCCGCCGATGACGCGGGAAGGCTATCAGCTTAGCAAATCGAAAGGACAGTAGCTAT
ATAAGGTTTTGCTATTCATTGAAAGCAGTAGTGACTGATTTGTATATA
>MinSyn_1503|Strength:0.017197562
GCGTGTCGTTTTAGTGAGGTCACTATCAGCTTGCCATCCTTACCGCTATGGGTAAGATTAGGTTGTCT
GCACCTCACATGTAGATCGAAAGGACAGTACGACCACCAAATATTTCTTGTCGACCACTATGCATGAG
CTAAGCACATACGTCAGTGCTGATCTTGCCTGCCTTGATATTGCGATAAAGGAAAGGCACTGCTAGGA
GCTATATAAGGTTTTGCTATTCATTGAAAGCAGTAGTGACTGATTTGTATATA
>MinSyn_1579|Strength:0.017243608
GCGTGTCGTTTTAGTGAGGGTGGGAGCCACCAGTATCCATCAACAAATAATCCAAGTAAGCTAGGAGG
ACCGATGCTGAGGAAAAAGAAGAGGTTCAGCTTAGCAAGACCTCAGCCACTTGTGTAGACCTCATGAG
CTAAGCACATACGTCAGCCACTATGCCCAATTAATCCTTACCGCTATGGGTAAGATTTGCTGATCTTG
CCTATATAAGGTTTTGCTATTCATTGAAAGCAGTAGTGACTGATTTGTATATA
>MinSyn_1454|Strength:0.017245451
GCGTGTCGTTTTAGTGAGGTGAAGATAAGATAATAATGTTGAAGATAAGACGGTAGGTCTCAGAAGAT
CAAAGGGCTACACTGCTAGGAGGACCGATGATGAGCTAAGCACATACGTCAGGCTATCAGCTTAGCAA
GACCTCTACAGCTTTGTCAAAAGCTAAAAAAGATGATGCTCGCGGTAGGTCACGACCACTATGCCCAA
CTATATAAGGTTTTGCTATTCATTGAAAGCAGTAGTGACTGATTTGTATATA
>MinSyn_1458|Strength:0.017342237
GCGTGTCGTTTTAGTGAGGTTATGACCCCCGCCGATGACGCGGGATCACATGTAGGCTATCAGCTTAT
CACTATCAGCAGGAGGACCGATGCTGATCTTGCCTCAAATATTTCTTGTTGGTACTTGTGTACAGGGG
AAAAAGAAGAGGTCCTGACGTAAGGGATGACGCACAGGTAGGTCACGACCACTATGCCCAATTAGGCT
ATATAAGGTTTTGCTATTCATTGAAAGCAGTAGTGACTGATTTGTATATA
>MinSyn_1128|Strength:0.017386695
GCGTGTCGTTTTAGTGAGGCTGACGTAAGGGATGACGCACAATGTAGGCTATCACTATCAGCCACATG
TAGGCTATCAGCTTAGCAAGACCTTCAGAAGATCAAAGGGCTATCTACAAAACTGGTACCTATATAAG
GTTTTGCTATTCATTGAAAGCAGTAGTGACTGATTTGTATATA
>MinSyn_1341|Strength:0.017432471
GCGTGTCGTTTTAGTGAGGAGCAAGTGGATTGCCCAATGACGTAAGCCATGACGTCTATAGGTTGTCT
CAGCCACTTGTGTCCAATTAATTTCGGGAAACCTCCTCGTGCCCAATTAGGTTGTCTGCACCTCACAC
TATATAAGGTTTTGCTATTCATTGAAAGCAGTAGTGACTGATTTGTATATA
>MinSyn_1106|Strength:0.017532053
GCGTGTCGTTTTAGTGAGGATGACGTAAGCCATGACGTCTAGTACAGGGCTCACTGCTTATGACCCCC
GCCGATGACGCGGGACCTGCCTTGATGGCTTTGTCAAAAGCTAAAAAAGATGATGCGACCCTATATAA
GGTTTTGCTATTCATTGAAAGCAGTAGTGACTGATTTGTATATA
>MinSyn_1200|Strength:0.017562848
GCGTGTCGTTTTAGTGAGGATCGAAAGGACAGTATGGTACTTGTGTACAGGTGAAGCATCTTCCAGGA
CCGATGATGAGCTAAGCACATACGTCAGGCGGTAGGTCACGACCACTATGCCCAATCAAATATTTCTT

GTCCGATGCTGATCTTGCCTGCCATCCTTACCGCTATGGGTAAGATTGTCACTATCAGCTAGGTCTAT
ATAAGGTTTTGCTATTCATTGAAAGCAGTAGTGACTGATTTGTATATA
>MinSyn_1173|Strength:0.017570611
GCGTGTCGTTTTAGTGAGGTGGTGGAGCACGACAGCTATCAGCCAAATATTTCTTGTCATGACGTAAG
CCATGACGTCTAGGCTCACTAACCATTATTGCGATGCCAAGATTGATGAAAAGTCAAAAACAAAAATC
AATTATAAGACTATATAAGGTTTTGCTATTCATTGAAAGCAGTAGTGACTGATTTGTATATA
>MinSyn_1425|Strength:0.017630818
GCGTGTCGTTTTAGTGAGGTCCATCAACAAATAATCCAAGTAAGGTCACGACCACTATGCCCAATTTC
GGGAAACCTCCTCGGGCTATCAGCTGGTGGAGCACGACAGACCGAGTGGGAGCCACCATAGCAAGACC
AGCCACTTGTGTGGCTCACTGCTAGGAGGACATGACGTAAGCCATGACGTCTATACTTGTGTACTATA
TAAGGTTTTGCTATTCATTGAAAGCAGTAGTGACTGATTTGTATATA
>MinSyn_1261|Strength:0.017729452
GCGTGTCGTTTTAGTGAGGCTGACGTAAGGGATGACGCACAGCTGATCTTGCCTGCCTTGATGATTAT
GACCCCCGCCGATGACGCGGGAAGGGCTCACAAATATTTCTTGTATGTAGGCTATCAGCTATATAAGG
TTTTGCTATTCATTGAAAGCAGTAGTGACTGATTTGTATATA
>MinSyn_1421|Strength:0.017919029
GCGTGTCGTTTTAGTGAGGCTGACGTAAGGGATGACGCACAGAGGACCGATGCTGATCTTGCCTTTAT
GACCCCCGCCGATGACGCGGGAAGCATGGTGGAGCACGACAGATGCTGATCTTGCCTCTATATAAGGT
TTTGCTATTCATTGAAAGCAGTAGTGACTGATTTGTATATA
>MinSyn_1978|Strength:0.01798332
GCGTGTCGTTTTAGTGAGGATCCTTACCGCTATGGGTAAGATTGCCTGCCTTGATGATAACCACGTCT
ACAACAAGACCTCTACAAAACTGGTACTAAAAATGTCAAAGATAGCTCACTGCTAGGAGGACCGATGC
TGATCTCACTATCAGCCAATTAGCTGACGTAAGGGATGACGCACAACTTGTGTACAGCTATATAAGGT
TTTGCTATTCATTGAAAGCAGTAGTGACTGATTTGTATATA
>MinSyn_1739|Strength:0.018000983
GCGTGTCGTTTTAGTGAGGGCTTTGTCAAAAGCTAAAAAAGATGATGCTATGCCATGAGCTAAGCACA
TACGTCAGGGGCTAACCACGTCTACAACTTAGCAAGACCTCTACAAAACCAAATATTTCTTGTCTGCA
CCTCACATGTAGGCTATCAGCTTTCACTATCAGCGACCTCTACAAGCAAGTGGATTTCTATATAAGGT
TTTGCTATTCATTGAAAGCAGTAGTGACTGATTTGTATATA
>MinSyn_1598|Strength:0.01826208
GCGTGTCGTTTTAGTGAGGAATTTCGGGAAACCTCCTCGAAAACTGGTACTTGTGTACAGGGCAAAAA
TGTCAAAGATAACTTGTGTACAGGGCTCACTGCTAGGATGAGCTAAGCACATACGTCAGAAAGATTGA
TGAAAAGTCAAAAACAAAAATCAATTATCATGTAGGCTATCAGCTTAGCAAGACCTATATAAGGTTTT
GCTATTCATTGAAAGCAGTAGTGACTGATTTGTATATA
>MinSyn_199|Strength:0.018377414
GCGTGTCGTTTTAGTGAGGGCTTTGTCAAAAGCTAAAAAAGATGATGCAACTGGTACTTGTATGAGCT
AAGCACATACGTCAGCAATCACTATCAGCCTCACATGTAGGCAGCCACTTGTGTTCGCGGTAGGTCAC
GACCACTATGCTCCATCAACAAATAATCCAAGTAAGACCTCACATGTAGGCTCTATATAAGGTTTTGC
TATTCATTGAAAGCAGTAGTGACTGATTTGTATATA
>MinSyn_1127|Strength:0.018418006
GCGTGTCGTTTTAGTGAGGATGAGCTAAGCACATACGTCAGCCACTATGCCCAAGTGGGAGCCACCAG
TATCGAAAGGACAGTAGTAGGCTATCAGCTTAGCAAGAAATTTCGGGAAACCTCCTCGTGAAAAATGT
CAAAGATAAGGACCGATGCTGATCTTTCCATCAACAAATAATCCAAGTAAGAGCTATATAAGGTTTTG
CTATTCATTGAAAGCAGTAGTGACTGATTTGTATATA
>MinSyn_1619|Strength:0.018463155
GCGTGTCGTTTTAGTGAGGTCCATCAACAAATAATCCAAGTAAGCGACCACTATGCCCAATTAGGTTG
TCTGCAGCTTTGTCAAAAGCTAAAAAAGATGATGCGCCCAATTAGGTTGTCTGCATGAGCTAAGCACA
TACGTCAGCAAAACTGGTACTTGCAAATATTTCTTGTCCTGCCTTGATGATCTATATAAGGTTTTGCT
ATTCATTGAAAGCAGTAGTGACTGATTTGTATATA
>MinSyn_1707|Strength:0.018710217
GCGTGTCGTTTTAGTGAGGAACCACGTCTACAACTGCACCTCACATGTAGGCCTGACGTAAGGGATGA
CGCACATCACTGCTAGGAGGACCGTGGGAGCCACCAGTAGGCTATCAGCTTAGCAAGACCTCTACACT
ATATAAGGTTTTGCTATTCATTGAAAGCAGTAGTGACTGATTTGTATATA
>MinSyn_198|Strength:0.018717309

GCGTGTCGTTTTAGTGAGGGTGGGAGCCACCAAAACTGGTACTTGTATGAGCTAAGCACATACGTCAG
ACGACAGGTGGCTCCTACAAAACTGGTAAATTTCGGGAAACCTCCTCGCCAAATATTTCTTGTGTGAA
GATAAGATAATAATGTTGAAGATAAGATGCACCTCACATGTAGGCTATCCTATATAAGGTTTTGCTAT
TCATTGAAAGCAGTAGTGACTGATTTGTATATA
>MinSyn_1359|Strength:0.01872148
GCGTGTCGTTTTAGTGAGGTGAAGATAAGATAATAATGTTGAAGATAAGATCAGCTTTTCAACAAACA
GGGCTCAAAGATTGATGAAAAGTCAAAAACAAAAATCAATTATACTGGTACTTGTGTAGGAAAAAGAA
GAGGTGTCAATGACGTAAGCCATGACGTCTAACAGGGCTCACTGCTAGGCTATATAAGGTTTTGCTAT
TCATTGAAAGCAGTAGTGACTGATTTGTATATA
>MinSyn_1856|Strength:0.018856866
GCGTGTCGTTTTAGTGAGGATCGAAAGGACAGTAACTATGCCCAATTAGGTTTTCAACAAGGGCTCAC
TGCTAGGAGGAATGAGCTAAGCACATACGTCAGGACCACTATGCCCAATTAGGTTGTCTGAAGATAAG
ATAATAATGTTGAAGATAAGAGCTGATCTTGCCTGCCTTGATGATCTATATAAGGTTTTGCTATTCAT
TGAAAGCAGTAGTGACTGATTTGTATATA
>MinSyn_1426|Strength:0.018889394
GCGTGTCGTTTTAGTGAGGTGGTGGAGCACGACAGCTCACTGCTAGGAGGACCGATGCTGACTGACGT
AAGGGATGACGCACAGACCGATGCCAAATATTTCTTGTAGACCTCTACAAAACAAAAATGTCAAAGAT
AACTATATAAGGTTTTGCTATTCATTGAAAGCAGTAGTGACTGATTTGTATATA
>MinSyn_1176|Strength:0.019021971
GCGTGTCGTTTTAGTGAGGAATTTCGGGAAACCTCCTCGAGGTTGTCTGAAGATAAGATAATAATGTT
GAAGATAAGATGTCTGCACCATCCTTACCGCTATGGGTAAGATTATATGAGCTAAGCACATACGTCAG
CACTCAAATATTTCTTGTGCTATCAGCTTAGCAAGACCTCTACAACTATATAAGGTTTTGCTATTCAT
TGAAAGCAGTAGTGACTGATTTGTATATA
>MinSyn_1682|Strength:0.01909241
GCGTGTCGTTTTAGTGAGGTGGTGGAGCACGACAGCCATGAGCTAAGCACATACGTCAGGTGAAGCAT
CTTCCGGCTTTTTCAACAACACTATGCCCAATTAGGTTGTCTGTGAAGATAAGATAATAATGTTGAAG
ATAAGACACGACCAAAAAATGTCAAAGATATTGTGTACAGGGCTCTATATAAGGTTTTGCTATTCATT
GAAAGCAGTAGTGACTGATTTGTATATA
>MinSyn_1435|Strength:0.019177112
GCGTGTCGTTTTAGTGAGGTCTCTCTGCCGACAGTGGTCCCAAAATCTGACGTAAGGGATGACGCACA
TCACTGCTAGGAGGACCGATGCTGTGGTGGAGCACGACAGCCTGCCCAAATATTTCTTGTTGCTATAT
AAGGTTTTGCTATTCATTGAAAGCAGTAGTGACTGATTTGTATATA
>MinSyn_1460|Strength:0.019201602
GCGTGTCGTTTTAGTGAGGATCCTTACCGCTATGGGTAAGATTGCTATCAGCTTAGCAAGACCTCTTG
AAGCATCTTCCGTGTACAGGGCTCACGTGGGAGCCACCAGCTCACTGCTAGGAGGACCGATGATGACG
TAAGCCATGACGTCTAGACCTCTACAAAACTGGTACTTGTCTATATAAGGTTTTGCTATTCATTGAAA
GCAGTAGTGACTGATTTGTATATA
>MinSyn_1382|Strength:0.019204899
GCGTGTCGTTTTAGTGAGGAGCAAGTGGATCAGCTTAGCAAGACCTCTACAAACAGTGGTCCCTCCAC
ACGACCAAAGATTGATGAAAAGTCAAAAACAAAAATCAATTATCTGGTACTTGTGTACCTGACGTAAG
GGATGACGCACAGCTAGGAGGACCGATGCTGATCTTGCCTGCTATATAAGGTTTTGCTATTCATTGAA
AGCAGTAGTGACTGATTTGTATATA
>MinSyn_193|Strength:0.01928955
GCGTGTCGTTTTAGTGAGGGCACACCAGCATGTGTTGATCACCAGCTAAACTGTCAGAAGATCAAAGG
GCTAACCTCACATGTAGATCCTTACCGCTATGGGTAAGATTCCTCACATGTAGGCTATCAGCTTAGAT
GACGTAAGCCATGACGTCTACACTATGCCCAATTAGGTTGCTATATAAGGTTTTGCTATTCATTGAAA
GCAGTAGTGACTGATTTGTATATA
>MinSyn_1423|Strength:0.019379955
GCGTGTCGTTTTAGTGAGGATCCTTACCGCTATGGGTAAGATTACAAAACTGGTACTTGTGTTCACTA
TCAGCATCTCAGTGGTCCCTCCACTAGCAAGTGGATTCAATGAGCTAAGCACATACGTCAGCTTAGCA
AGATGGTGGAGCACGACATCACTGCTAGGAGGACCGATGCCTATATAAGGTTTTGCTATTCATTGAAA
GCAGTAGTGACTGATTTGTATATA
>MinSyn_1964|Strength:0.019455789
GCGTGTCGTTTTAGTGAGGAACCATTATTGCGCAAAACTGGTACTTGTGTACACTGACGTAAGGGATG

ACGCACATACAGGGCTCACTGCTATTATGACCCCCGCCGATGACGCGGGATTAGGTTGTCTGCCTATA
TAAGGTTTTGCTATTCATTGAAAGCAGTAGTGACTGATTTGTATATA
>MinSyn_1638|Strength:0.019478852
GCGTGTCGTTTTAGTGAGGTGAAGATAAGATAATAATGTTGAAGATAAGAACTGGTGGAGCACGACAC
TGGTACTTGTGGCTTTGTCAAAAGCTAAAAAAGATGATGCCATGTAGGCTATCAGCTTAGCAAGACCT
GACGTAAGGGATGACGCACATGTACAGGGCTCACTGCTCTATATAAGGTTTTGCTATTCATTGAAAGC
AGTAGTGACTGATTTGTATATA
>MinSyn_1523|Strength:0.01950201
GCGTGTCGTTTTAGTGAGGTTATGACCCCCGCCGATGACGCGGGATGTGTACAGGGCTGCTTTGTCAA
AAGCTAAAAAAGATGATGCTAGGTTGTCTGCACCTCACATGTCAAATATTTCTTGTCGACCACTATGC
CCAATTAGGTTGATGACGTAAGCCATGACGTCTATTAGCTATATAAGGTTTTGCTATTCATTGAAAGC
AGTAGTGACTGATTTGTATATA
>MinSyn_1952|Strength:0.01957652
GCGTGTCGTTTTAGTGAGGATTGCGATAAAGGAAAGGTCTGCACCTCACATGTAGGCTATCAGCTTAA
TGAGCTAAGCACATACGTCAGTGTGTACAGGGCTCACTGCTAGGATGGTGGAGCACGACAACTGGTAC
TTGTGTACAGTTATGACCCCCGCCGATGACGCGGGATCTATATAAGGTTTTGCTATTCATTGAAAGCA
GTAGTGACTGATTTGTATATA
>MinSyn_1859|Strength:0.019577704
GCGTGTCGTTTTAGTGAGGTGAAGCATCTTCCTGTGTACAGGGCTCACTGCTAGGAGGACATCGAAAG
GACAGTAACTGGTACTTGTGTACAGGGCTCAAAGATTGATGAAAAGTCAAAAACAAAAATCAATTATT
ACAAAACTGGTCTGACGTAAGGGATGACGCACACTGATCTATATAAGGTTTTGCTATTCATTGAAAGC
AGTAGTGACTGATTTGTATATA
>MinSyn_1182|Strength:0.019781909
GCGTGTCGTTTTAGTGAGGATTGCGATAAAGGAAAGGAGGAGGACCGATGCTGACTGACGTAAGGGAT
GACGCACACAAGACCTCTACTCAGAAGATCAAAGGGCTATTTTTCAACAACTATGCCCAATTCTATAT
AAGGTTTTGCTATTCATTGAAAGCAGTAGTGACTGATTTGTATATA
>MinSyn_1495|Strength:0.019805874
GCGTGTCGTTTTAGTGAGGCTGACGTAAGGGATGACGCACAGGAGGACCGATGCTGATCTTGCTTTTC
AACAAAGGGCTCACTGCTATCACTATCAGCATCAGCTTAGCAAGACCTATATAAGGTTTTGCTATTCA
TTGAAAGCAGTAGTGACTGATTTGTATATA
>MinSyn_1451|Strength:0.019861756
GCGTGTCGTTTTAGTGAGGCTGACGTAAGGGATGACGCACACTTGCCTGCCTTGATGATCACTATCAG
CTACATCGAAAGGACAGTAGGCTCACTGCTAGGAGGACCGATGCTGCTATATAAGGTTTTGCTATTCA
TTGAAAGCAGTAGTGACTGATTTGTATATA
>MinSyn_1444|Strength:0.019867379
GCGTGTCGTTTTAGTGAGGGTGGGAGCCACCATACAGGGCTCACTGCTAGGAGGACCGATTGCGATAA
AGGAAAGGACAAAACTATGAGCTAAGCACATACGTCAGACCACTATGCCCAATTAGGTTGTCTGAAGA
TTGATGAAAAGTCAAAAACAAAAATCAATTATTTCTATATAAGGTTTTGCTATTCATTGAAAGCAGTA
GTGACTGATTTGTATATA
>MinSyn_1697|Strength:0.020079372
GCGTGTCGTTTTAGTGAGGAACCATTATTGCGTTGTGTACAGGGCTCACTTCAGAAGATCAAAGGGCT
AATGCCCAATTAGGTTGTCATGAGCTAAGCACATACGTCAGTAGCAAGACCTCTACAGCTTTGTCAAA
AGCTAAAAAAGATGATGCGTACTTGTGTACACTATATAAGGTTTTGCTATTCATTGAAAGCAGTAGTG
ACTGATTTGTATATA
>MinSyn_1229|Strength:0.020088803
GCGTGTCGTTTTAGTGAGGCTGACGTAAGGGATGACGCACAGCTCACTGCTAGGAGGACCGATGCTGC
AGTGGTCCCTCCACTCTACAAAAGTGGGAGCCACCACTTGCCTGCCTATATAAGGTTTTGCTATTCAT
TGAAAGCAGTAGTGACTGATTTGTATATA
>MinSyn_1405|Strength:0.020189986
GCGTGTCGTTTTAGTGAGGATGACGTAAGCCATGACGTCTATGTACAGGGGTGGGAGCCACCACATCC
TTACCGCTATGGGTAAGATTGTGTACAGGGCTCACTGCTAGGAGGCTATATAAGGTTTTGCTATTCAT
TGAAAGCAGTAGTGACTGATTTGTATATA
>MinSyn_1647|Strength:0.020283857
GCGTGTCGTTTTAGTGAGGAATTTCGGGAAACCTCCTCGTTAGCAAGACCAGGTGGCTCCTACTAGGA

GGACCGATGCTGATCTTGCGGAAAAAGAAGAGGTGATGAGCTAAGCACATACGTCAGTCTGAAGATTG
ATGAAAAGTCAAAAACAAAAATCAATTATTTCTATATAAGGTTTTGCTATTCATTGAAAGCAGTAGTG
ACTGATTTGTATATA
>MinSyn_1772|Strength:0.020378984
GCGTGTCGTTTTAGTGAGGGGAAAAAGAAGAGGTCAAGACCTCTACAAGGTGGCTCCTACGGCTAACC
ATTATTGCGCTGGTACTTGTGTACAGGAACCACGTCTACAAAGCTTAGCAAGACCTCTACAAAACATG
AGCTAAGCACATACGTCAGCACGACCACTATATAAGGTTTTGCTATTCATTGAAAGCAGTAGTGACTG
ATTTGTATATA
>MinSyn_1314|Strength:0.020477556
GCGTGTCGTTTTAGTGAGGTCACTATCAGCCAATTAGGTTGTTTATGACCCCCGCCGATGACGCGGGA
TACAGGGCTCACTGCTAGATTGCGATAAAGGAAAGGGCTATCAGCTTAGCAAGACCTCTACTGACGTA
AGGGATGACGCACATTGTCTGCACCTCTATATAAGGTTTTGCTATTCATTGAAAGCAGTAGTGACTGA
TTTGTATATA
>MinSyn_165|Strength:0.020485095
GCGTGTCGTTTTAGTGAGGAATTTCGGGAAACCTCCTCGATGCTGATCTTGCCTGCCTTGATAGGTGG
CTCCTACAGCTTAGCAAGAAAGATTGATGAAAAGTCAAAAACAAAAATCAATTATTCACATGTAGGAT
GACGTAAGCCATGACGTCTAAGCAAGACCTATATAAGGTTTTGCTATTCATTGAAAGCAGTAGTGACT
GATTTGTATATA
>MinSyn_1841|Strength:0.020530189
GCGTGTCGTTTTAGTGAGGATTGCGATAAAGGAAAGGGTCTGCACCTCACAACCACGTCTACAAGCTA
TCAGCTTAGCAAGACCTCTACAAAATGGTGGAGCACGACATACAAAACTGGTACTTGATGAGCTAAGC
ACATACGTCAGTAGGTCACGACCACCTATATAAGGTTTTGCTATTCATTGAAAGCAGTAGTGACTGAT
TTGTATATA
>MinSyn_1377|Strength:0.020803487
GCGTGTCGTTTTAGTGAGGTGAAGCATCTTCCAGAAAAATGTCAAAGATACTGCCTTGATGATGAGCT
AAGCACATACGTCAGTGTGTACCAAATATTTCTTGTACCTCTACAAAACTTCACTATCAGCTATCAGC
TTAGCAAGACCTCTACAAAACTCTATATAAGGTTTTGCTATTCATTGAAAGCAGTAGTGACTGATTTG
TATATA
>MinSyn_1932|Strength:0.020810557
GCGTGTCGTTTTAGTGAGGGCACACCAGCATGTGTTGATCACCAGCTCTCACATGAGCTAAGCACATA
CGTCAGATTAGGTTGTCTGAACCACGTCTACAAAGACCTCTACAAAACTGGTACTTGTGTATGAAGCA
TCTTCCAAGACCTCTACAAAACCTATATAAGGTTTTGCTATTCATTGAAAGCAGTAGTGACTGATTTG
TATATA
>MinSyn_1110|Strength:0.020929979
GCGTGTCGTTTTAGTGAGGATGAGCTAAGCACATACGTCAGTGTAGGCTATCAGCTTAGCAAGACCTC
AGAAGATCAAAGGGCTATGCTGATCTTGCCTGGCACACCAGCATGTGTTGATCACCAGCTCTATGCCC
AATTAGGTTGTTTTCAACAAGCCTATATAAGGTTTTGCTATTCATTGAAAGCAGTAGTGACTGATTTG
TATATA
>MinSyn_1479|Strength:0.020942219
GCGTGTCGTTTTAGTGAGGTCAGAAGATCAAAGGGCTACCTCACATGTAGGCTATCAGCTCACTATCA
GCTTAGGTTGTCTGCACCTCACATGTAGGCTAACCATTATTGCGTGTAGGCTATCAGCTATGACGTAA
GCCATGACGTCTATGCCTTGACTATATAAGGTTTTGCTATTCATTGAAAGCAGTAGTGACTGATTTGT
ATATA
>MinSyn_1754|Strength:0.021112742
GCGTGTCGTTTTAGTGAGGAATTTCGGGAAACCTCCTCGTATCATCCTTACCGCTATGGGTAAGATTT
AGCAAGACCTCTACAAAACTGGATGAGCTAAGCACATACGTCAGCTCACTGCTAGGAGGATCGAAAGG
ACAGTAGTAGGCTATCAGCTCTATATAAGGTTTTGCTATTCATTGAAAGCAGTAGTGACTGATTTGTA
TATA
>MinSyn_155|Strength:0.021300695
GCGTGTCGTTTTAGTGAGGCAGTGGTCCCTCCACAAGACCTCTACAAAACTGGTAAGGTGGCTCCTAC
CTTAGCAAGATCGAAAGGACAGTATTAGCAAGACCATGAGCTAAGCACATACGTCAGTGCTAGGAGGA
CCGATGCTGATCTTGCCCTATATAAGGTTTTGCTATTCATTGAAAGCAGTAGTGACTGATTTGTATAT
A
>MinSyn_1150|Strength:0.021440992

GCGTGTCGTTTTAGTGAGGGCACACCAGCATGTGTTGATCACCAGCTTTGCCTGCCTTGATGATATCC
TTACCGCTATGGGTAAGATTTGCTGATCTTGCCTATGAGCTAAGCACATACGTCAGTCACTGCTAGGA
GGACCGATGCTGATCTCTATATAAGGTTTTGCTATTCATTGAAAGCAGTAGTGACTGATTTGTATATA
>MinSyn_1573|Strength:0.021448408
GCGTGTCGTTTTAGTGAGGATCCTTACCGCTATGGGTAAGATTCCTCTATGAGCTAAGCACATACGTC
AGTTGTGTACAGGGCTCACTGCTAGGAAGGTGGCTCCTACCCTCTTATGACCCCCGCCGATGACGCGG
GAAGCAAGACCTCTACACTATATAAGGTTTTGCTATTCATTGAAAGCAGTAGTGACTGATTTGTATAT
A
>MinSyn_1431|Strength:0.021452326
GCGTGTCGTTTTAGTGAGGATTGCGATAAAGGAAAGGCGATATGACGTAAGCCATGACGTCTAGGCTG
CACACCAGCATGTGTTGATCACCAGCTTGCCCAATTAGGTTGTCTGCACTATATAAGGTTTTGCTATT
CATTGAAAGCAGTAGTGACTGATTTGTATATA
>MinSyn_1845|Strength:0.021573241
GCGTGTCGTTTTAGTGAGGTTTTCAACAAAAGACCTCTACAAAACTGGTACTTGTGATCCTTACCGCT
ATGGGTAAGATTTAGGCTATCAGCTTAGCAAGACCTCATGACGTAAGCCATGACGTCTATGCTGATCT
TGCCTGCCTTGATGCTATATAAGGTTTTGCTATTCATTGAAAGCAGTAGTGACTGATTTGTATATA
>MinSyn_1725|Strength:0.022325321
GCGTGTCGTTTTAGTGAGGTCCATCAACAAATAATCCAAGTAAGACCGATGCTTTTTCAACAACCACT
ATGCAATTTCGGGAAACCTCCTCGGTGTACAGGGCTCACTGCTAGGAATGACGTAAGCCATGACGTCT
ATACAGGGCTATATAAGGTTTTGCTATTCATTGAAAGCAGTAGTGACTGATTTGTATATA
>MinSyn_1827|Strength:0.022713563
GCGTGTCGTTTTAGTGAGGAGGTGGCTCCTACCCATGAGCTAAGCACATACGTCAGAGGCTATCAGCT
TAGCAAGACCTCTTGAAGCATCTTCCTGGTACTTGTGTACAGGGCTCACATCGAAAGGACAGTACTGC
CTTGCTATATAAGGTTTTGCTATTCATTGAAAGCAGTAGTGACTGATTTGTATATA
>MinSyn_1103|Strength:0.022795355
GCGTGTCGTTTTAGTGAGGCTGACGTAAGGGATGACGCACAGACCGATGCTGATCTTGCCTGCCTTGA
TGAAGCATCTTCCACATGTAGGCTATCAGCTTCTATATAAGGTTTTGCTATTCATTGAAAGCAGTAGT
GACTGATTTGTATATA
>MinSyn_1522|Strength:0.023264462
GCGTGTCGTTTTAGTGAGGCAGCCACTTGTGTCAGGGCTCACTGCTAGGAGGACCGATGATGAGCTAA
GCACATACGTCAGAGGAGGACCGATGCTGATCTTGCCTGCCAATTTCGGGAAACCTCCTCGCACCTCC
TATATAAGGTTTTGCTATTCATTGAAAGCAGTAGTGACTGATTTGTATATA
>MinSyn_1120|Strength:0.023463712
GCGTGTCGTTTTAGTGAGGAAAAATGTCAAAGATAGGGCTCACTGCTAGGAGGACCGATGCTGTTTTC
AACAACTATGCCCAATTAGGTTGTCTGCACCTCACATGAGCTAAGCACATACGTCAGAAACTGGTCTA
TATAAGGTTTTGCTATTCATTGAAAGCAGTAGTGACTGATTTGTATATA
>MinSyn_1956|Strength:0.023685096
GCGTGTCGTTTTAGTGAGGATGAGCTAAGCACATACGTCAGTAACCATTATTGCGTGTGGGAGCCACC
AAAACTGTGAAGCATCTTCCAGGTCACGACCAAATTTCGGGAAACCTCCTCGGCTTAGCAAGACCCTA
TATAAGGTTTTGCTATTCATTGAAAGCAGTAGTGACTGATTTGTATATA
>MinSyn_1395|Strength:0.023692929
GCGTGTCGTTTTAGTGAGGTCACTATCAGCGGACCGATGCTGCAGTGGTCCCTCCACCATTGCGATAA
AGGAAAGGTGCCTGCCTTGATGATGAGCTAAGCACATACGTCAGTGTACAGGGCTCACTGCTAGCTAT
ATAAGGTTTTGCTATTCATTGAAAGCAGTAGTGACTGATTTGTATATA
>MinSyn_189|Strength:0.023844492
GCGTGTCGTTTTAGTGAGGTTATGACCCCCGCCGATGACGCGGGACACTGCTAGGAGGACCGATGCTG
ATCTTGATGAGCTAAGCACATACGTCAGCTGGTACAGCCACTTGTGTGTTGTCTGCACCTCACCTATA
TAAGGTTTTGCTATTCATTGAAAGCAGTAGTGACTGATTTGTATATA
>MinSyn_1890|Strength:0.023993332
GCGTGTCGTTTTAGTGAGGCAGTGGTCCCTCCACGTCTGCACATCGAAAGGACAGTACTGCTAGGAGG
ACCGATGCTGATCGTGGGAGCCACCATGATGAGCTAAGCACATACGTCAGCGACCACTATGCCTATAT
AAGGTTTTGCTATTCATTGAAAGCAGTAGTGACTGATTTGTATATA
>MinSyn_1934|Strength:0.024559775
GCGTGTCGTTTTAGTGAGGCTGACGTAAGGGATGACGCACACCTCTACAAAACTGGTATGAAGCATCT

TCCAGGCTATCAGCTTAGCAAGACCCTATATAAGGTTTTGCTATTCATTGAAAGCAGTAGTGACTGAT
TTGTATATA
>MinSyn_15|Strength:0.024589983
GCGTGTCGTTTTAGTGAGGTCAGAAGATCAAAGGGCTAAAAACTGGTACTTGTGTACAGGCTGACGTA
AGGGATGACGCACACTGCTAGTCTCTCTGCCGACAGTGGTCCCAAAGGCTATCAGCTTCTATATAAGG
TTTTGCTATTCATTGAAAGCAGTAGTGACTGATTTGTATATA
>MinSyn_1260|Strength:0.024750139
GCGTGTCGTTTTAGTGAGGCAGTGGTCCCTCCACCACTATGCCCAATTAGGTTGTCCTGACGTAAGGG
ATGACGCACAATGTAGGCTATCAGCTTAGCAAGAAAAATGTCAAAGATAACCGATGCTATATAAGGTT
TTGCTATTCATTGAAAGCAGTAGTGACTGATTTGTATATA
>MinSyn_1137|Strength:0.024775628
GCGTGTCGTTTTAGTGAGGTTTTCAACAAGACCTCTACAAAACTGGTACAGGTGGCTCCTACTGCCCA
ATTAGGTTGTCTGCACCTCAAACCATTATTGCGAGGTTGTCTGCACCTCACATGCTGACGTAAGGGAT
GACGCACAAAATGAGCTAAGCACATACGTCAGCATGTAGGCTATGAAGATAAGATAATAATGTTGAAG
ATAAGACCTCTACAAAACTTCACTATCAGCACTTGAACCACGTCTACAATTAAGATTGATGAAAAGTC
AAAAACAAAAATCAATTATGGTCACGACCACTATGCCCAATCTATATAAGGTTTTGCTATTCATTGAA
AGCAGTAGTGACTGATTTGTATATA
>MinSyn_1655|Strength:0.024785352
GCGTGTCGTTTTAGTGAGGAAAAATGTCAAAGATAAACAGCCACTTGTGTCAGCTTAGCAAGTGAAGC
ATCTTCCTATGCCCAATTAGGTTGTCTGCACCTCTGACGTAAGGGATGACGCACAGCTATATAAGGTT
TTGCTATTCATTGAAAGCAGTAGTGACTGATTTGTATATA
>MinSyn_1264|Strength:0.024914246
GCGTGTCGTTTTAGTGAGGCAAATATTTCTTGTCTGCTAGGAGAACCATTATTGCGTCACATGTAGGC
TATCAGCTTAGATGAGCTAAGCACATACGTCAGAGCAAGACCTCTACAAAACTGCTATATAAGGTTTT
GCTATTCATTGAAAGCAGTAGTGACTGATTTGTATATA
>MinSyn_1351|Strength:0.02498151
GCGTGTCGTTTTAGTGAGGTCAGAAGATCAAAGGGCTATACAAAACTGGATTGCGATAAAGGAAAGGG
GGCTCACTGCTAGGAGGACCGAATGAGCTAAGCACATACGTCAGTACTTGTGTACTATATAAGGTTTT
GCTATTCATTGAAAGCAGTAGTGACTGATTTGTATATA
>MinSyn_1610|Strength:0.025426811
GCGTGTCGTTTTAGTGAGGCAGTGGTCCCTCCACCACATGTAGGCTATCAGCTTAGCAAGACAGGTGG
CTCCTACCATGTAGGCTATGACGTAAGCCATGACGTCTATAGGTCACGACCCTATATAAGGTTTTGCT
ATTCATTGAAAGCAGTAGTGACTGATTTGTATATA
>MinSyn_1149|Strength:0.025764592
GCGTGTCGTTTTAGTGAGGTTATGACCCCCGCCGATGACGCGGGAGTTGTCTGCACCTCACATGATGA
GCTAAGCACATACGTCAGGGTACTTGTAGCAAGTGGATGTACAGGGCTCCTATATAAGGTTTTGCTAT
TCATTGAAAGCAGTAGTGACTGATTTGTATATA
>MinSyn_1437|Strength:0.02605098
GCGTGTCGTTTTAGTGAGGAATTTCGGGAAACCTCCTCGGACCTCTACAAAACTGGTACTTATGACGT
AAGCCATGACGTCTAGCGGTGAAGCATCTTCCAATTAGGTTGTCTGCCTATATAAGGTTTTGCTATTC
ATTGAAAGCAGTAGTGACTGATTTGTATATA
>MinSyn_1907|Strength:0.026053205
GCGTGTCGTTTTAGTGAGGATGAGCTAAGCACATACGTCAGCAAAACTCACTATCAGCTATCAGCTTA
GTGGTGGAGCACGACACTGGTACTTGTGTACAGGGCTCACTGCTAGCTATATAAGGTTTTGCTATTCA
TTGAAAGCAGTAGTGACTGATTTGTATATA
>MinSyn_136|Strength:0.026105415
GCGTGTCGTTTTAGTGAGGTCCATCAACAAATAATCCAAGTAAGCGCATGACGTAAGCCATGACGTCT
ATATGCCCAATTAGGTTGTCTGCTGGTGGAGCACGACAGATGCTGATCTATATAAGGTTTTGCTATTC
ATTGAAAGCAGTAGTGACTGATTTGTATATA
>MinSyn_1594|Strength:0.02650641
GCGTGTCGTTTTAGTGAGGATTGCGATAAAGGAAAGGTGCCCAATTAGGTTGTCTGCACCTATGACGT
AAGCCATGACGTCTAGGTTGTCTGCACCTCACATGTAGGCTATCTATATAAGGTTTTGCTATTCATTG
AAAGCAGTAGTGACTGATTTGTATATA
>MinSyn_1602|Strength:0.026664157

GCGTGTCGTTTTAGTGAGGATGAGCTAAGCACATACGTCAGACCTCAAGCAAGTGGATAGGAGGACCG
ATGCTGATCTTGCCTGCCTTAGGTGGCTCCTACGCCTGCCTTCTATATAAGGTTTTGCTATTCATTGA
AAGCAGTAGTGACTGATTTGTATATA
>MinSyn_1595|Strength:0.026712149
GCGTGTCGTTTTAGTGAGGTTTTCAACAACAAGACCTCTACAAAACAGCCACTTGTGTTATCAGCTTA
GCACTGACGTAAGGGATGACGCACAGTCACGACCACTATGCCCTATATAAGGTTTTGCTATTCATTGA
AAGCAGTAGTGACTGATTTGTATATA
>MinSyn_1724|Strength:0.026964157
GCGTGTCGTTTTAGTGAGGAAAAATGTCAAAGATAACCTCACATGTAGGCTATCAGTTATGACCCCCG
CCGATGACGCGGGAACAATGACGTAAGCCATGACGTCTACTCCTATATAAGGTTTTGCTATTCATTGA
AAGCAGTAGTGACTGATTTGTATATA
>MinSyn_1666|Strength:0.027074567
GCGTGTCGTTTTAGTGAGGTTTTCAACAAGACCGATGCTGATCATGACGTAAGCCATGACGTCTAATG
TAGGCTATCAGCTTAGATCCTTACCGCTATGGGTAAGATTACTATATAAGGTTTTGCTATTCATTGAA
AGCAGTAGTGACTGATTTGTATATA
>MinSyn_1369|Strength:0.027097141
GCGTGTCGTTTTAGTGAGGTTATGACCCCCGCCGATGACGCGGGACTCTACAAAAACCACGTCTACAA
ACAGGGCAACCATTATTGCGCAAGACCTCTACAAAACTGGTACTTCAGAAGATCAAAGGGCTAAGCAG
CCACTTGTGTGATGCTGATCTTGCCTGAGGTGGCTCCTACAATTAGGTGGAAAAAGAAGAGGTTACAG
GATGAGCTAAGCACATACGTCAGAGATGACGTAAGCCATGACGTCTAGGTCACGACCACTATGCCCAA
TTAGGTTGCTATATAAGGTTTTGCTATTCATTGAAAGCAGTAGTGACTGATTTGTATATA
>MinSyn_135|Strength:0.027147305
GCGTGTCGTTTTAGTGAGGATGAGCTAAGCACATACGTCAGGCCCAATTAGGTTGTATTGCGATAAAG
GAAAGGGTACAGAACCACGTCTACAATACTTGTGTACAGGCTATATAAGGTTTTGCTATTCATTGAAA
GCAGTAGTGACTGATTTGTATATA
>MinSyn_1233|Strength:0.027756945
GCGTGTCGTTTTAGTGAGGTCACTATCAGCGACCGATGCTGATCTTGCCTGCCTTATGACCCCCGCCG
ATGACGCGGGAGCCTGCCTAGCAAGTGGATGGTCACGACCACTATGCAAAAATGTCAAAGATAAGACC
TCTACAAAACTGGTACTATGAGCTAAGCACATACGTCAGCTATCAGCTTAGCATGACGTAAGCCATGA
CGTCTAGCAAGACCTCTTGAAGATAAGATAATAATGTTGAAGATAAGAGACCACTATGCCCAATTAGG
TTGATCGAAAGGACAGTATCACTAAGATTGATGAAAAGTCAAAAACAAAAATCAATTATAAAACTGGT
ACTTGTGTACTATATAAGGTTTTGCTATTCATTGAAAGCAGTAGTGACTGATTTGTATATA
>MinSyn_1894|Strength:0.027838859
GCGTGTCGTTTTAGTGAGGATGAGCTAAGCACATACGTCAGTACAAAACTGGTACTTGTGTACAGGGT
CAGAAGATCAAAGGGCTATGCTAGGAGGACCGATGCTATATAAGGTTTTGCTATTCATTGAAAGCAGT
AGTGACTGATTTGTATATA
>MinSyn_151|Strength:0.028037313
GCGTGTCGTTTTAGTGAGGATTGCGATAAAGGAAAGGGATCTTGCCTGCTGACGTAAGGGATGACGCA
CACTGGTATCGAAAGGACAGTAATGCTGATCTTGCCTATATAAGGTTTTGCTATTCATTGAAAGCAGT
AGTGACTGATTTGTATATA
>MinSyn_1608|Strength:0.028142077
GCGTGTCGTTTTAGTGAGGAACCACGTCTACAAACAAAACTGGAGCAAGTGGATTGTAATGAGCTAAG
CACATACGTCAGAGGTTGTCTGCACCTCACATGTCTATATAAGGTTTTGCTATTCATTGAAAGCAGTA
GTGACTGATTTGTATATA
>MinSyn_1557|Strength:0.028721352
GCGTGTCGTTTTAGTGAGGGTGGGAGCCACCAAAAACTGGTACTTGTGTAATGACGTAAGCCATGACG
TCTAACTGCTAGGAGGACCGATGCTGATCTCTATATAAGGTTTTGCTATTCATTGAAAGCAGTAGTGA
CTGATTTGTATATA
>MinSyn_1660|Strength:0.028867742
GCGTGTCGTTTTAGTGAGGTGAAGATAAGATAATAATGTTGAAGATAAGAACCTCTACAAAACCTGAC
GTAAGGGATGACGCACATCTTGCCTGCCTTGCTATATAAGGTTTTGCTATTCATTGAAAGCAGTAGTG
ACTGATTTGTATATA
>MinSyn_1187|Strength:0.029307077
GCGTGTCGTTTTAGTGAGGATCGAAAGGACAGTAGCTATCAGCATGACGTAAGCCATGACGTCTATGC

ACCTCACATGTAGGCTATCAGCTTAGCCTATATAAGGTTTTGCTATTCATTGAAAGCAGTAGTGACTG
ATTTGTATATA
>MinSyn_1635|Strength:0.029953691
GCGTGTCGTTTTAGTGAGGTCCATCAACAAATAATCCAAGTAAGGGTACTTGTGTACAGGGCTCAATG
AGCTAAGCACATACGTCAGAACTGGCTATATAAGGTTTTGCTATTCATTGAAAGCAGTAGTGACTGAT
TTGTATATA
>MinSyn_1689|Strength:0.030270748
GCGTGTCGTTTTAGTGAGGCTGACGTAAGGGATGACGCACACTAGGAGGACCGATGATTGCGATAAAG
GAAAGGTTGCCTGCCTTGATGATCTATATAAGGTTTTGCTATTCATTGAAAGCAGTAGTGACTGATTT
GTATATA
>MinSyn_1418|Strength:0.030300597
GCGTGTCGTTTTAGTGAGGAACCATTATTGCGGCCTTGATGAATGACGTAAGCCATGACGTCTAACAG
GGCTCACTGCTAGGAGGACCGACTATATAAGGTTTTGCTATTCATTGAAAGCAGTAGTGACTGATTTG
TATATA
>MinSyn_1814|Strength:0.030938623
GCGTGTCGTTTTAGTGAGGAGGTGGCTCCTACCTTGCCTGCCTATGACGTAAGCCATGACGTCTAAGG
GCTCACTGCTAGGAGGACCCTATATAAGGTTTTGCTATTCATTGAAAGCAGTAGTGACTGATTTGTAT
ATA
>MinSyn_1896|Strength:0.031020139
GCGTGTCGTTTTAGTGAGGATGAGCTAAGCACATACGTCAGCGTCACTATCAGCACGGAAAAAGAAGA
GGTAGGAGGACCGATGCTGACTATATAAGGTTTTGCTATTCATTGAAAGCAGTAGTGACTGATTTGTA
TATA
>MinSyn_17|Strength:0.031716429
GCGTGTCGTTTTAGTGAGGCTGACGTAAGGGATGACGCACAAGGCTATCAGTGAAGCATCTTCCTCTT
GCCTGCCTTGATGATACTATATAAGGTTTTGCTATTCATTGAAAGCAGTAGTGACTGATTTGTATATA
>MinSyn_1811|Strength:0.031865714
GCGTGTCGTTTTAGTGAGGATCCTTACCGCTATGGGTAAGATTCAAAACTGGTACTTATGACGTAAGC
CATGACGTCTAGTGTACTATATAAGGTTTTGCTATTCATTGAAAGCAGTAGTGACTGATTTGTATATA
>MinSyn_115|Strength:0.031871281
GCGTGTCGTTTTAGTGAGGTCAGAAGATCAAAGGGCTACACTATGCCCAATTAGGTTGTTTATGACCC
CCGCCGATGACGCGGGACACCTCACATGATGAGCTAAGCACATACGTCAGGGACCGATGCTGATCTCT
GACGTAAGGGATGACGCACATCACTGCTAGGAGGACCGATGCTGATTGGTGGAGCACGACATCACGAC
CACTATGCCCAATTATTTTCAACAATGCAAATATTTCTTGTTGTACAGGGCTCACTGCCAGTGGTCCC
TCCACGATGAGCTTTGTCAAAAGCTAAAAAAGATGATGCCGGCTATATAAGGTTTTGCTATTCATTGA
AAGCAGTAGTGACTGATTTGTATATA
>MinSyn_1471|Strength:0.032122302
GCGTGTCGTTTTAGTGAGGCTGACGTAAGGGATGACGCACAGCTTAGCAAGACCTTATGACCCCCGCC
GATGACGCGGGATTGCTATATAAGGTTTTGCTATTCATTGAAAGCAGTAGTGACTGATTTGTATATA
>MinSyn_1681|Strength:0.032341304
GCGTGTCGTTTTAGTGAGGATGACGTAAGCCATGACGTCTATCGCGGTAGGTCATCCATCAACAAATA
ATCCAAGTAAGGGTCTATATAAGGTTTTGCTATTCATTGAAAGCAGTAGTGACTGATTTGTATATA
>MinSyn_1329|Strength:0.032919259
GCGTGTCGTTTTAGTGAGGATCGAAAGGACAGTAACCGATGCTGATCATGACGTAAGCCATGACGTCT
AGCCCAATTAGCTATATAAGGTTTTGCTATTCATTGAAAGCAGTAGTGACTGATTTGTATATA
>MinSyn_1478|Strength:0.033065233
GCGTGTCGTTTTAGTGAGGTCAGAAGATCAAAGGGCTACCCAATTAGGTTGTTTATGACCCCCGCCGA
TGACGCGGGATTAGGTTGTCTGCACCTCACATGTAGGCTAATGAGCTAAGCACATACGTCAGTAGGAG
GACCGATGCTGATCTTGCCTATGACGTAAGCCATGACGTCTATCTGCACCTCACATGCACACCAGCAT
GTGTTGATCACCAGCTCTAGGAGGACCGCAGTGGTCCCTCCACGGAGGACCGATGCTGATCTTCTCTC
TGCCGACAGTGGTCCCAAATTAGGTTGTCTGCACCTCACTCCATCAACAAATAATCCAAGTAAGAAAC
TGGTACTTGTGTACAGGGCTATATAAGGTTTTGCTATTCATTGAAAGCAGTAGTGACTGATTTGTATA
TA
>MinSyn_1747|Strength:0.033414286
GCGTGTCGTTTTAGTGAGGCTGACGTAAGGGATGACGCACATAGGTCACGACCACATCGAAAGGACAG

TACTCTACACTATATAAGGTTTTGCTATTCATTGAAAGCAGTAGTGACTGATTTGTATATA
>MinSyn_1441|Strength:0.033563403
GCGTGTCGTTTTAGTGAGGCAGCCACTTGTGTGGAGGACCGATGCTGAGCACACCAGCATGTGTTGAT
CACCAGCTATCAGCTTAGCAAGACCTCTTTTTCAACAAACATATCGAAAGGACAGTATTAGGTTGTCT
GCACCTCATCAGAAGATCAAAGGGCTAACAGGGCTCACTGCTAGGAGGACTCTCTCTGCCGACAGTGG
TCCCAAATACAGGGCTAACCATTATTGCGATGCCCAATTAGGTTGTCTGCACCTCTGACGTAAGGGAT
GACGCACATACAGGGCTCACTGCTAGGAGGACATGACGTAAGCCATGACGTCTAGTTGTCTGCACCTC
CTATATAAGGTTTTGCTATTCATTGAAAGCAGTAGTGACTGATTTGTATATA
>MinSyn_142|Strength:0.033609848
GCGTGTCGTTTTAGTGAGGATGAGCTAAGCACATACGTCAGCTACAAAACTGGTACTTGTGTACAGCA
AGTGGATGCTATATAAGGTTTTGCTATTCATTGAAAGCAGTAGTGACTGATTTGTATATA
>MinSyn_1700|Strength:0.033685895
GCGTGTCGTTTTAGTGAGGAGCAAGTGGATGTAGGTCACGACCACTTGAAGATAAGATAATAATGTTG
AAGATAAGACACGACCACTATGCCCAATTAGGAACCACGTCTACAAGGCTCACTGCTAGGAGGACCGA
TGCTGATCCTGACGTAAGGGATGACGCACATCTGCACCTCACATGTAGGCTATGACGTAAGCCATGAC
GTCTATCACGACCACTATGCCCAATATCGAAAGGACAGTACTGCACCTCACATGTAGGCTAGCTTTGT
CAAAAGCTAAAAAAGATGATGCAGCTTAGCAAGACCTCTACAAAACTGGCTATATAAGGTTTTGCTAT
TCATTGAAAGCAGTAGTGACTGATTTGTATATA
>MinSyn_192|Strength:0.033866412
GCGTGTCGTTTTAGTGAGGAGCAAGTGGATTCAGCTTAGCTGACGTAAGGGATGACGCACATCACATG
TAGGCTACTATATAAGGTTTTGCTATTCATTGAAAGCAGTAGTGACTGATTTGTATATA
>MinSyn_1680|Strength:0.034331037
GCGTGTCGTTTTAGTGAGGATGACGTAAGCCATGACGTCTAAGGTCACGACTGACGTAAGGGATGACG
CACAAGGGCTCACTGCTGGTGGAGCACGACATAGGAGGACCGATGCTGATCTTGCCAGCAAGTGGATG
TCTGCACCTCACATGCAGCCACTTGTGTAATTAGGTTTCACTATCAGCAATTAGGTTGTCTGCACCTC
ACAGTGGGAGCCACCACTCTACAAAACTGGTACTTGTGTACACTATATAAGGTTTTGCTATTCATTGA
AAGCAGTAGTGACTGATTTGTATATA
>MinSyn_1790|Strength:0.034391473
GCGTGTCGTTTTAGTGAGGAGCAAGTGGATTGCTAGGAGGACCGATGACGTAAGCCATGACGTCTACA
CGACCCTATATAAGGTTTTGCTATTCATTGAAAGCAGTAGTGACTGATTTGTATATA
>MinSyn_1271|Strength:0.034676772
GCGTGTCGTTTTAGTGAGGCTGACGTAAGGGATGACGCACAGAGGTGGCTCCTACCTATGACGTAAGC
CATGACGTCTATAGGAGGACCGATGCTGATCTTGAAGCATCTTCCCATTGCGATAAAGGAAAGGAGGG
CTCACTGCTAGGAGGACCGATAGCAAGTGGATGTAGGTCACGACCACTATGCACACCAGCATGTGTTG
ATCACCAGCTAGACCTCTACAAAACTGGCTTTGTCAAAAGCTAAAAAAGATGATGCCGATGCTGATCT
TGCCTGCCTTGCTATATAAGGTTTTGCTATTCATTGAAAGCAGTAGTGACTGATTTGTATATA
>MinSyn_1531|Strength:0.034719531
GCGTGTCGTTTTAGTGAGGCTGACGTAAGGGATGACGCACACTACATCGAAAGGACAGTAACTATGCC
CAATCTATATAAGGTTTTGCTATTCATTGAAAGCAGTAGTGACTGATTTGTATATA
>MinSyn_1346|Strength:0.035195238
GCGTGTCGTTTTAGTGAGGTTTTCAACAACCTCTACAAACTGACGTAAGGGATGACGCACAGCTTAGC
AACTATATAAGGTTTTGCTATTCATTGAAAGCAGTAGTGACTGATTTGTATATA
>MinSyn_1532|Strength:0.035240476
GCGTGTCGTTTTAGTGAGGAGGTGGCTCCTACCGGTAGGTCACGATGACGTAAGCCATGACGTCTACT
TGCTATATAAGGTTTTGCTATTCATTGAAAGCAGTAGTGACTGATTTGTATATA
>MinSyn_1159|Strength:0.036054424
GCGTGTCGTTTTAGTGAGGATCGAAAGGACAGTATTGTCTGCACCTCACATGGAAAAAGAAGAGGTGG
TTGTCTGCTTTGTCAAAAGCTAAAAAAGATGATGCTGCCCAATTAGGTTGTCTTCAGAAGATCAAAGG
GCTATAGGTCACGACCACTATGCCCTGAAGCATCTTCCCTAATTTCGGGAAACCTCCTCGAGGTTGTC
TGCACCTCACATGTAGGCTATCCTGACGTAAGGGATGACGCACAAACTGGTACTTGTGTACAGGGCTA
TGAGCTAAGCACATACGTCAGCAGGGCTCACTGCTCTATATAAGGTTTTGCTATTCATTGAAAGCAGT
AGTGACTGATTTGTATATA
>MinSyn_1661|Strength:0.036673993
GCGTGTCGTTTTAGTGAGGGTGGAGCCACCAGATCTTGCCTGCAACCATTATTGCGCACATGTAGGC

TATCAGATGACGTAAGCCATGACGTCTATCACATGTCTGACGTAAGGGATGACGCACACGATGCTGAT
CTTGCCTGCCTTAAGATTGATGAAAAGTCAAAAACAAAAATCAATTATGAGATTGCGATAAAGGAAAG
GGGTTGTCTGCACCTCACACTATATAAGGTTTTGCTATTCATTGAAAGCAGTAGTGACTGATTTGTAT
ATA
>MinSyn_1339|Strength:0.03689043
GCGTGTCGTTTTAGTGAGGTGAAGATAAGATAATAATGTTGAAGATAAGAACATGTAGGCTATCAGCT
TAGCAAGAGCTTTGTCAAAAGCTAAAAAAGATGATGCGCAAGACCTCTACAAAACTGGTACTTGTCTG
ACGTAAGGGATGACGCACATGATATGACGTAAGCCATGACGTCTATACAAAACTGGTACTTGTGCTAT
ATAAGGTTTTGCTATTCATTGAAAGCAGTAGTGACTGATTTGTATATA
>MinSyn_1950|Strength:0.037131645
GCGTGTCGTTTTAGTGAGGGGAAAAAGAAGAGGTGGTACTTGTGTACAGGGCTCTTTTCAACAACAAA
ACTGGTACTTGTGTACAGCAGCCACTTGTGTCATGTAGGCTATCAGCTTAGCAAGACCTAAAAATGTC
AAAGATATAGGTTGTATGAGCTAAGCACATACGTCAGTGCTGATCTTGCCTGCCATGACGTAAGCCAT
GACGTCTATCTACAAAACTGTCAGAAGATCAAAGGGCTAGCAAGACCTCTACAAAACTGGTCTATATA
AGGTTTTGCTATTCATTGAAAGCAGTAGTGACTGATTTGTATATA
>MinSyn_1366|Strength:0.037265546
GCGTGTCGTTTTAGTGAGGATGAGCTAAGCACATACGTCAGCTTTTCAACAATGCTGATCTTGCTATA
TAAGGTTTTGCTATTCATTGAAAGCAGTAGTGACTGATTTGTATATA
>MinSyn_1384|Strength:0.040124557
GCGTGTCGTTTTAGTGAGGAGCAAGTGGATAGGACCGATGCTGATCTTGCCTGCCTATGAGCTAAGCA
CATACGTCAGTTGTCTGCACCTCACCTGACGTAAGGGATGACGCACAACTTGTGTACAGGGCTGAAGC
ATCTTCCACTGGTAAAGATTGATGAAAAGTCAAAAACAAAAATCAATTATTCACGACCACTATGCCCA
ATTAGGTTGTCTAACCACGTCTACAAGCCTATATAAGGTTTTGCTATTCATTGAAAGCAGTAGTGACT
GATTTGTATATA
>MinSyn_1947|Strength:0.040510839
GCGTGTCGTTTTAGTGAGGCAAATATTTCTTGTAGGTTGTCTGCTGACGTAAGGGATGACGCACACAC
CTCACATGTAGGCAGCAAGTGGATTTAGCAAATGAGCTAAGCACATACGTCAGTAGCAAGACCTCTAC
AAAACTGGTACTAGGTGGCTCCTACGAGGACCGATGCTGATCTCTCTCTGCCGACAGTGGTCCCAAAA
GGAGGACCGATGCTTATGACCCCCGCCGATGACGCGGGAGTCTGCACCTCACATGTAGGAACCACGTC
TACAATAAAAATGTCAAAGATAGTTGTCTGCACCTCACATGTAGGCTCTATATAAGGTTTTGCTATTC
ATTGAAAGCAGTAGTGACTGATTTGTATATA
>MinSyn_1541|Strength:0.041325136
GCGTGTCGTTTTAGTGAGGCTGACGTAAGGGATGACGCACAACAGCCACTTGTGTTAGCAAGACCTCT
ACAAAACTGGATGAGCTAAGCACATACGTCAGTCTGCACCTCACATGTAGGCTATCAGCTAAGATTGA
TGAAAAGTCAAAAACAAAAATCAATTATGGGCTCACTGCTAGGAGGAACCACGTCTACAAGTCACGAC
ATTGCGATAAAGGAAAGGGGTAGGTCACGACCACTATGCCCAATTTTCAACAACCACTATGCCCAATA
AAAATGTCAAAGATACCCAAGCACACCAGCATGTGTTGATCACCAGCTGTCTGCCTATATAAGGTTTT
GCTATTCATTGAAAGCAGTAGTGACTGATTTGTATATA
>MinSyn_1162|Strength:0.041810331
GCGTGTCGTTTTAGTGAGGTCCATCAACAAATAATCCAAGTAAGGCTCACTGCTAGGAGGACCGATGC
TGAATTTCGGGAAACCTCCTCGTGATCTTGCCATGACGTAAGCCATGACGTCTACGATGCTGCAGCCA
CTTGTGTTCAGCTTAGCAAGACCTCATGAGCTAAGCACATACGTCAGATGCTGATCTTGCCTGCCTTG
ATGCAGTGGTCCCTCCACGGTCACGAAAGATTGATGAAAAGTCAAAAACAAAAATCAATTATCATGTA
GGCTATCAGCTTAGCAAGATGGTGGAGCACGACATCACGACCACTATGCCCAATTAGGCTATATAAGG
TTTTGCTATTCATTGAAAGCAGTAGTGACTGATTTGTATATA
>MinSyn_1658|Strength:0.041972455
GCGTGTCGTTTTAGTGAGGGGAAAAAGAAGAGGTCGATGACGTAAGCCATGACGTCTAGGGCTATGAG
CTAAGCACATACGTCAGACCACTATGCCCAATTAGGTTGTGGGAGCCACCACTTAGCATCCATCAACA
AATAATCCAAGTAAGCTAGGAGGACCGATGCTGATCTCTATATAAGGTTTTGCTATTCATTGAAAGCA
GTAGTGACTGATTTGTATATA
>MinSyn_10|Strength:0.041973963
GCGTGTCGTTTTAGTGAGGAACCACGTCTACAAAATTAGGTTGTCTGATGAGCTAAGCACATACGTCA
GAAAACTGGTACTTGTGTACACAGCCACTTGTGTCTCATGACGTAAGCCATGACGTCTAGCAAGACCT
CTACAAAACTGGTACAATTTCGGGAAACCTCCTCGTATCAGCTTAGCAAGATCCTTACCGCTATGGGT

AAGATTAGCTTTGTCAAAAGCTAAAAAAGATGATGCAGACCTCTAACCATTATTGCGGGTAGGTCACG
ACCACTATGCCCAATCACTATCAGCTAGGTCACGACCACTATGCTATATAAGGTTTTGCTATTCATTG
AAAGCAGTAGTGACTGATTTGTATATA
>MinSyn_1943|Strength:0.041979111
GCGTGTCGTTTTAGTGAGGTTATGACCCCCGCCGATGACGCGGGACAGCTTAGCAAGACCTCTAAGAT
TGATGAAAAGTCAAAAACAAAAATCAATTATGCTAGGAGGACCGATGCTGAGGTGGCTCCTACTGTGT
ACAGGGCTCTGACGTAAGGGATGACGCACAGAGGACAGCAAGTGGATGACCGATGCATGACGTAAGCC
ATGACGTCTAACTGCTAGGAGGACCGATGTGAAGCATCTTCCCAGCTTAGCAAGACCTCTACAAACTA
TATAAGGTTTTGCTATTCATTGAAAGCAGTAGTGACTGATTTGTATATA
>MinSyn_1455|Strength:0.04253744
GCGTGTCGTTTTAGTGAGGATCGAAAGGACAGTAGAACCATTATTGCGTGCACCTCACATGTAGGCTA
TCACTGACGTAAGGGATGACGCACACATGTAGGCTATCAGCTTAGCAAGACCATGAGCTAAGCACATA
CGTCAGCTATGCCCAATTATCCTTACCGCTATGGGTAAGATTTAGGGCACACCAGCATGTGTTGATCA
CCAGCTTGCTGATCTTTCAGAAGATCAAAGGGCTAGGTCACGACCACTATGAAGCATCTTCCTAGCCT
ATATAAGGTTTTGCTATTCATTGAAAGCAGTAGTGACTGATTTGTATATA
>MinSyn_1502|Strength:0.043725803
GCGTGTCGTTTTAGTGAGGATGAGCTAAGCACATACGTCAGCTCTACAAAACTGCTGACGTAAGGGAT
GACGCACAGCTGATCGTGGGAGCCACCACTAGATTGCGATAAAGGAAAGGCTGCACCTCACATGTAGG
CAGCAAGTGGATTTGTTTATGACCCCCGCCGATGACGCGGGAACCTCTACAAAACTGGTACTTCTATA
TAAGGTTTTGCTATTCATTGAAAGCAGTAGTGACTGATTTGTATATA
>MinSyn_1396|Strength:0.044485253
GCGTGTCGTTTTAGTGAGGATGACGTAAGCCATGACGTCTACTCACATGTAGGCTCTGACGTAAGGGA
TGACGCACACTGGTACTTGTGTCCATCAACAAATAATCCAAGTAAGCCAATTAGGTTGTCTGCACCTC
ACATCTCTCTGCCGACAGTGGTCCCAAAGGTACTTGTGTACACAAATATTTCTTGTGTGTACACTATA
TAAGGTTTTGCTATTCATTGAAAGCAGTAGTGACTGATTTGTATATA
>MinSyn_1746|Strength:0.045862884
GCGTGTCGTTTTAGTGAGGTCTCTCTGCCGACAGTGGTCCCAAAACATGTAGGCTATCAGCTTAGCAA
GACCGCTTTGTCAAAAGCTAAAAAAGATGATGCCTACGTGGGAGCCACCATCTGCACCTCACATGGGA
AAAAGAAGAGGTTAGCAAGACCTCTACAATGAGCTAAGCACATACGTCAGGCTGCACACCAGCATGTG
TTGATCACCAGCTTGATGACGTAAGCCATGACGTCTATCACATGTAGGCTATCAGCTTAGCAAGACCC
TATATAAGGTTTTGCTATTCATTGAAAGCAGTAGTGACTGATTTGTATATA
>MinSyn_1963|Strength:0.049207601
GCGTGTCGTTTTAGTGAGGTGAAGATAAGATAATAATGTTGAAGATAAGAGGCTATCAGCTTAGCAAG
ACCTCTACATGAGCTAAGCACATACGTCAGCAGGGCTATGACGTAAGCCATGACGTCTAGAGGACCGA
TGCTGATCTTGCTATATAAGGTTTTGCTATTCATTGAAAGCAGTAGTGACTGATTTGTATATA
>MinSyn_1172|Strength:0.049452849
GCGTGTCGTTTTAGTGAGGGCACACCAGCATGTGTTGATCACCAGCTTAGGCTATCAGCTTAGATGAC
GTAAGCCATGACGTCTAAAACTGGTACTAACCACGTCTACAATGATATGAGCTAAGCACATACGTCAG
CCAATTAGGTTGTCTGCACCTCACATGTTCAGAAGATCAAAGGGCTATTAGCAAGACCTCTACAAAAC
AGGTGGCTCCTACGTACTTGTGTACAGCTATATAAGGTTTTGCTATTCATTGAAAGCAGTAGTGACTG
ATTTGTATATA
>MinSyn_1326|Strength:0.051305632
GCGTGTCGTTTTAGTGAGGTTTTCAACAAAGCAAGACCTTATGACCCCCGCCGATGACGCGGGACTGC
TATCCTTACCGCTATGGGTAAGATTTGTGTACAGGGCTCACTGCTAGGAGGAGCAAGTGGATCACCTC
AATTGCGATAAAGGAAAGGGCTAGGAGGACCGATGCTGATCTTGCCTTCACTATCAGCCTGGTAATGA
GCTAAGCACATACGTCAGTGCTTTGTCAAAAGCTAAAAAAGATGATGCCAGGGCTCACTGCTAGGAGG
ACCGACTGACGTAAGGGATGACGCACAACCACTATGCCCAATTAGGCTATATAAGGTTTTGCTATTCA
TTGAAAGCAGTAGTGACTGATTTGTATATA
>MinSyn_1484|Strength:0.051479447
GCGTGTCGTTTTAGTGAGGGCTTTGTCAAAAGCTAAAAAAGATGATGCCCTCACATGTAGGCTATCAG
CTTATCACTATCAGCTGTCTATGAGCTAAGCACATACGTCAGACCACTATGCCCAATTAGGTTGTCTG
ATGACGTAAGCCATGACGTCTAAAGAAGCAAGTGGATGACCTCTATCTCTCTGCCGACAGTGGTCCCA
AAGCCCTATATAAGGTTTTGCTATTCATTGAAAGCAGTAGTGACTGATTTGTATATA
>MinSyn_1643|Strength:0.054842667

GCGTGTCGTTTTAGTGAGGTCAGAAGATCAAAGGGCTAAACTGGTACTTGTGTACAGGTTATGACCCC
CGCCGATGACGCGGGATGTCTGCTGAAGCATCTTCCGGACCGATATGAGCTAAGCACATACGTCAGCA
CCTCACATGTAGGCTATCAGCTGGAAAAAGAAGAGGTTCACATGTCTGACGTAAGGGATGACGCACAC
ATGTAGGCTATCAGCTTAGCAAAAGATTGATGAAAAGTCAAAAACAAAAATCAATTATTACTCTATAT
AAGGTTTTGCTATTCATTGAAAGCAGTAGTGACTGATTTGTATATA
>MinSyn_1300|Strength:0.057345832
GCGTGTCGTTTTAGTGAGGATCGAAAGGACAGTACACCTCACATGTAGGCTATCAGCTTAGCAAATTG
CGATAAAGGAAAGGTGCCCAATTAGGTTGTCTGCACCAGCCACTTGTGTAGGAGGACCGATGCTGATC
TTGCCTGCCTTATGAGCTAAGCACATACGTCAGTTGTGTACAGGGGCTTTGTCAAAAGCTAAAAAAGA
TGATGCATGACTGACGTAAGGGATGACGCACAGCCCAATTACTATATAAGGTTTTGCTATTCATTGAA
AGCAGTAGTGACTGATTTGTATATA
>MinSyn_1948|Strength:0.05755919
GCGTGTCGTTTTAGTGAGGTGGTGGAGCACGACACTATGAGCTAAGCACATACGTCAGTAGGAGGACC
GTCCATCAACAAATAATCCAAGTAAGCCTCTACAAAACTGGTACTTGTATGACGTAAGCCATGACGTC
TAGTCACGACCACTATAGCAAGTGGATTGCCTGCCTTGATGAAGATAAGATAATAATGTTGAAGATAA
GACTCACATGTAGGCTATCAGCTTAGTGGGAGCCACCATCAGCTTAGTCAGAAGATCAAAGGGCTAAC
CACTATGCCCAATTACTATATAAGGTTTTGCTATTCATTGAAAGCAGTAGTGACTGATTTGTATATA
>MinSyn_1170|Strength:0.058370223
GCGTGTCGTTTTAGTGAGGATGAGCTAAGCACATACGTCAGTATCAGCTCTGACGTAAGGGATGACGC
ACATCTGCACCTCACATGTAGGCTATCAGCCACTTGTGTGATGCTGATCTCTATATAAGGTTTTGCTA
TTCATTGAAAGCAGTAGTGACTGATTTGTATATA
>MinSyn_1194|Strength:0.05863679
GCGTGTCGTTTTAGTGAGGAACCACGTCTACAAGGCTCACTGCTAGGAGGACCGATGCTGATGAAGAT
AAGATAATAATGTTGAAGATAAGACGACCACTGGAAAAAGAAGAGGTCAAGACCTCTACAAAACTCTG
ACGTAAGGGATGACGCACATTGTCTGCACCTCACATGTAATTGCGATAAAGGAAAGGGTAGGTCACGA
CCACTATGCCATGAGCTAAGCACATACGTCAGCTATGCCCAATTAGGATCCTTACCGCTATGGGTAAG
ATTTTAGCAAGACTATATAAGGTTTTGCTATTCATTGAAAGCAGTAGTGACTGATTTGTATATA
>MinSyn_1445|Strength:0.059317638
GCGTGTCGTTTTAGTGAGGAAAAATGTCAAAGATACTCTAACCATTATTGCGACTATGCCCAATTAGG
TTGTCTGCAAGCAAGTGGATTGTCTGCACTTATGACCCCCGCCGATGACGCGGGAAATGAGCTAAGCA
CATACGTCAGCGACCACTATGCCCAATTAGGTTGATTGCGATAAAGGAAAGGCACGACCACTATGCCC
AATTAGGTTATGACGTAAGCCATGACGTCTATGGTACTGAAGCATCTTCCACCTCTACAAAACTGGCT
TTGTCAAAAGCTAAAAAAGATGATGCAGACCTCTCTATATAAGGTTTTGCTATTCATTGAAAGCAGTA
GTGACTGATTTGTATATA
>MinSyn_1903|Strength:0.060483837
GCGTGTCGTTTTAGTGAGGTGAAGATAAGATAATAATGTTGAAGATAAGACAATTAGGTTGTCTGCAC
CTCACATGATGACGTAAGCCATGACGTCTACAAAACTGGTACTTGTGTACAGGGTGAAGCATCTTCCG
GTTGTCTGCTGACGTAAGGGATGACGCACACACCTCACATGTAGGCTATCAGCTTCCATCAACAAATA
ATCCAAGTAAGGGACCGATGCTGATCTTCTATATAAGGTTTTGCTATTCATTGAAAGCAGTAGTGACT
GATTTGTATATA
>MinSyn_1275|Strength:0.06066881
GCGTGTCGTTTTAGTGAGGATGACGTAAGCCATGACGTCTACTATATTGCGATAAAGGAAAGGGCCCA
ATTAGGTTGTCTGCTGACGTAAGGGATGACGCACAAGGACCGATGCTGATAAAAATGTCAAAGATAAC
TATTCACTATCAGCAGGTTGTCTGCACCTCACATGTAGTGGTGGAGCACGACAAGGACCGATGCTGAT
CTTGCCTGCTATATAAGGTTTTGCTATTCATTGAAAGCAGTAGTGACTGATTTGTATATA
>MinSyn_1415|Strength:0.061343697
GCGTGTCGTTTTAGTGAGGATGAGCTAAGCACATACGTCAGCAATTAGGTTGTCTGCACCTCACATTT
ATGACCCCCGCCGATGACGCGGGAGCTAGGAGGACCGATGCTCTGACGTAAGGGATGACGCACACGGT
AGGTCACGTGAAGATAAGATAATAATGTTGAAGATAAGATGCCCAATTAGGTGCTTTGTCAAAAGCTA
AAAAAGATGATGCCCCAATTAGGTTGTCTGCACCTCACATGTATCGAAAGGACAGTAGACCACTATGC
CCAATCAAATATTTCTTGTAGGTTGTCTCTATATAAGGTTTTGCTATTCATTGAAAGCAGTAGTGACT
GATTTGTATATA
>MinSyn_1726|Strength:0.062428681
GCGTGTCGTTTTAGTGAGGCTGACGTAAGGGATGACGCACAATGTAGGCTATCAGCTTAGCAAGACCT

CTGAAGCATCTTCCAGGAGGACCGATGCTGATCTTGCATGAGCTAAGCACATACGTCAGGGCTATCAG
CTTAGCAAGACCTCTACAAATTTCGGGAAACCTCCTCGGCTCACTGCTAGGAGGACCGATGCTGCAGT
GGTCCCTCCACATTAGGTTGTCTGCGCTTTGTCAAAAGCTAAAAAAGATGATGCCACCTCACATGTAG
GCTATCCTATATAAGGTTTTGCTATTCATTGAAAGCAGTAGTGACTGATTTGTATATA
>MinSyn_1238|Strength:0.062548544
GCGTGTCGTTTTAGTGAGGATGAGCTAAGCACATACGTCAGTGCCCAATTAGGTTGTCATTGCGATAA
AGGAAAGGCCACTATGCCCAATTCTGACGTAAGGGATGACGCACACTGCCTTGATGATAAAGATTGAT
GAAAAGTCAAAAACAAAAATCAATTATATGCTGATCTTGCCTGCCTTGATGATAATCGAAAGGACAGT
ATTAGGTTGTCTGCACAGTGGTCCCTCCACGCTATATAAGGTTTTGCTATTCATTGAAAGCAGTAGTG
ACTGATTTGTATATA
>MinSyn_1153|Strength:0.062877652
GCGTGTCGTTTTAGTGAGGTCCATCAACAAATAATCCAAGTAAGACCACTATGCCCAATTAGGTTGTC
TGTCAGAAGATCAAAGGGCTAGCTCACTGCTAGGAGGACCCTGACGTAAGGGATGACGCACAACTGGT
ACTTGTGTACAGGGCTCACTATCAGCTAGGAGGACCGATGCTGATCTTGCAACCATTATTGCGCTGCT
AGGAGGACCGATGCTGATCTTATGACGTAAGCCATGACGTCTATGCACCTCACATGTTTATGACCCCC
GCCGATGACGCGGGAGATGCTGAAATTTCGGGAAACCTCCTCGGGTTGTCTGCACCTCACATGTTGAA
GCATCTTCCGACCACTATGCCCAATTAGGTCTATATAAGGTTTTGCTATTCATTGAAAGCAGTAGTGA
CTGATTTGTATATA
>MinSyn_1158|Strength:0.064265005
GCGTGTCGTTTTAGTGAGGTGAAGATAAGATAATAATGTTGAAGATAAGACCTGACGTAAGGGATGAC
GCACAGACCGATGCTGATCTTGCCTGCCTTGCAAATATTTCTTGTCTGATCATGACGTAAGCCATGAC
GTCTATCTACAAAACTGGTACTTGTGTACAGGGCTTATGACCCCCGCCGATGACGCGGGACACATGTA
GGCTATCAGCTCTATATAAGGTTTTGCTATTCATTGAAAGCAGTAGTGACTGATTTGTATATA
>MinSyn_1385|Strength:0.065361181
GCGTGTCGTTTTAGTGAGGCTGACGTAAGGGATGACGCACATGGTACTTGTGTACAGGGCTCACTGCA
AATATTTCTTGTTAGGTTGTCTGCACCTCACATGTATGAGCTAAGCACATACGTCAGAGGACCGATGC
TGATCTTGTCCATCAACAAATAATCCAAGTAAGTAGGAGGACCGATGCTGATCTTGCCTGGGAAAAAG
AAGAGGTCAATTAGGTTGTCTGCACCTCACATGTGTGGGAGCCACCAAAGACCCTATATAAGGTTTTG
CTATTCATTGAAAGCAGTAGTGACTGATTTGTATATA
>MinSyn_1704|Strength:0.06567382
GCGTGTCGTTTTAGTGAGGAGGTGGCTCCTACTCGTCAGAAGATCAAAGGGCTAGCGGTATGAGCTAA
GCACATACGTCAGTGTCTAAGATTGATGAAAAGTCAAAAACAAAAATCAATTATGCTCACTGCTAGGA
CTGACGTAAGGGATGACGCACACTGCTAGGAGCACACCAGCATGTGTTGATCACCAGCTGAACCATTA
TTGCGACTTGTGTACAGGGCTCACTGCTAGGACTATATAAGGTTTTGCTATTCATTGAAAGCAGTAGT
GACTGATTTGTATATA
>MinSyn_1285|Strength:0.067091247
GCGTGTCGTTTTAGTGAGGAGCAAGTGGATGGTCACTGACGTAAGGGATGACGCACAACCTCTACAAA
ACTGGTACTTGTGTATGACGTAAGCCATGACGTCTATCTGTTTTCAACAAGAGGACCCAGTGGTCCCT
CCACGACCCTATATAAGGTTTTGCTATTCATTGAAAGCAGTAGTGACTGATTTGTATATA
>MinSyn_1263|Strength:0.067441069
GCGTGTCGTTTTAGTGAGGCAAATATTTCTTGTACTGGTACTTGTGTACAGGGCTGAAGCATCTTCCT
CACTGCTAGGAGGACCTTTTCAACAAACCGATGCTGATCTTGCCTGCCTTATGAGCTAAGCACATACG
TCAGCACTGCTAGGAGGACCGATGCTCACTATCAGCGTACAGGGCTCACTGCTAGGAGGACCGATGCT
GACGTAAGGGATGACGCACACTGGTACTTGTGTACAGGGCTCACTATATAAGGTTTTGCTATTCATTG
AAAGCAGTAGTGACTGATTTGTATATA
>MinSyn_1518|Strength:0.0689391
GCGTGTCGTTTTAGTGAGGCAAATATTTCTTGTGCTCACTGCTAGGAGGACCGAGCAAGTGGATCACT
GCATGACGTAAGCCATGACGTCTAGTTGTCTGCACCTCACATGTAGGCTATCAATCGAAAGGACAGTA
GCCTGCTCAGAAGATCAAAGGGCTACTTAGCAAGACCTCTACAAAACTGGTACCTGACGTAAGGGATG
ACGCACAGGTAGGTCACGACCAAGATTGATGAAAAGTCAAAAACAAAAATCAATTATCGACCACTATG
CCCAATTAGGTTGTTCTCTCTGCCGACAGTGGTCCCAAAATCAGCTTAGCAAGACCTCTACAAAACCT
ATATAAGGTTTTGCTATTCATTGAAAGCAGTAGTGACTGATTTGTATATA
>MinSyn_1247|Strength:0.070280416
GCGTGTCGTTTTAGTGAGGAGCAAGTGGATGCACCTCACATGTAGGCTATCAGCGCACACCAGCATGT

GTTGATCACCAGCTTATGCCCAATTAGGTTGTCTGCACCTCAAGATTGATGAAAAGTCAAAAACAAAA
ATCAATTATACTATGCATGAGCTAAGCACATACGTCAGCCACTATGCCCAATTAGGTTGTCTGCACCC
AGTGGTCCCTCCACGACCTCTACAAAACTGGTACTTGTGTACATCTCTCTGCCGACAGTGGTCCCAAA
ATCAGCTTAGCATGACGTAAGCCATGACGTCTAACAGGGCTCACTGCTAGTGGTGGAGCACGACATAG
GAGGACCGATGCTGATCTTGCCTCTATATAAGGTTTTGCTATTCATTGAAAGCAGTAGTGACTGATTT
GTATATA
>MinSyn_1817|Strength:0.070778551
GCGTGTCGTTTTAGTGAGGCAGTGGTCCCTCCACGACCTCTACAAAAGCACACCAGCATGTGTTGATC
ACCAGCTAGGAGGACCGAGTGGGAGCCACCAGCTATCAGCTTAGCAAGACCTCTGACGTAAGGGATGA
CGCACACTTAGCAAGACCTCTACAAAACTGGTATCACTATCAGCGGGCTCACTGCTAGGAGGATCTCT
CTGCCGACAGTGGTCCCAAAACAAAAAAAATGTCAAAGATATAGGATGAGCTAAGCACATACGTCAGT
GCACCTCACATGTAGGCTATCAGAAGATTGATGAAAAGTCAAAAACAAAAATCAATTATCAAAACTGG
TACTTGTGTACAGGGCTATATAAGGTTTTGCTATTCATTGAAAGCAGTAGTGACTGATTTGTATATA
>MinSyn_1459|Strength:0.071048987
GCGTGTCGTTTTAGTGAGGTCACTATCAGCCCAATTAGGTTGTCTGCACCTGAAGATAAGATAATAAT
GTTGAAGATAAGATACAGGGCTCACAGCCACTTGTGTCGACCACTATGCCCAATTAGGTTGTCTTTTC
AACAATCTGCACCTCACATGTAGGCTATCAGATGACGTAAGCCATGACGTCTAGGGCTCACTGCTAGG
AAAAAATGTCAAAGATATATGCCCAATTAGGTTGTCTGCACTGACGTAAGGGATGACGCACACTATGC
CCAATTAGGTTGTCTGCACCATGAGCTAAGCACATACGTCAGACCACTATGCCCAATTAGGTTGTCTG
CACTATATAAGGTTTTGCTATTCATTGAAAGCAGTAGTGACTGATTTGTATATA
>MinSyn_1834|Strength:0.074312714
GCGTGTCGTTTTAGTGAGGATTGCGATAAAGGAAAGGGTACAGGTTATGACCCCCGCCGATGACGCGG
GAGCTATCAGCACACCAGCATGTGTTGATCACCAGCTAGGCTATCAGCTTAGCGTGGGAGCCACCACA
TGTAGGCTATCAGCTTAGCAAGACCATGAGCTAAGCACATACGTCAGTGCCCAATTAGGTTGTCTGCA
CAAATATTTCTTGTTAGGTTGTCTTCCATCAACAAATAATCCAAGTAAGATCAGCTTAGCAAGACCTC
TACAAAACTCTGACGTAAGGGATGACGCACATCACTGCTACTATATAAGGTTTTGCTATTCATTGAAA
GCAGTAGTGACTGATTTGTATATA
>MinSyn_170|Strength:0.07519134
GCGTGTCGTTTTAGTGAGGATGAGCTAAGCACATACGTCAGGTAGGTCACGACCACTATGCCCAAATT
TCGGGAAACCTCCTCGCTGCACCTATGACGTAAGCCATGACGTCTAACCACTATGCCCAATTAGGTTG
TCTGCATTATGACCCCCGCCGATGACGCGGGATCGCGGTAGGTCACGACCACCTATATAAGGTTTTGC
TATTCATTGAAAGCAGTAGTGACTGATTTGTATATA
>MinSyn_1670|Strength:0.077797735
GCGTGTCGTTTTAGTGAGGAATTTCGGGAAACCTCCTCGAGGGCTCACTGGGAAAAAGAAGAGGTCTC
ACATGTAGGCTATCAGCTTAGCATGAGCTAAGCACATACGTCAGGCTGATCTTGCCTGCCTTAAGATT
GATGAAAAGTCAAAAACAAAAATCAATTATACTGGTACTTCAGCCACTTGTGTCACATGTAGGCTATC
AGCTTCTGACGTAAGGGATGACGCACAACCAGCAAGTGGATCTTAGCAAGACCTCTACAACCATTATT
GCGAGGCTATCAGCTTAGCAACTATATAAGGTTTTGCTATTCATTGAAAGCAGTAGTGACTGATTTGT
ATATA
>MinSyn_1592|Strength:0.078365236
GCGTGTCGTTTTAGTGAGGAGCAAGTGGATACCTCACATGTAGGCTATTCCATCAACAAATAATCCAA
GTAAGATGATGTGGGAGCCACCAGTGTACAGGGCTCACTGCTAGGAGGACCGAAATTTCGGGAAACCT
CCTCGCGACCAATGAGCTAAGCACATACGTCAGGCAAGACCTCTACAAAACTATCCTTACCGCTATGG
GTAAGATTTAGGTCACGACCACTATGCCCAATAAAAATGTCAAAGATATGTGTACAGGGCTCACTGCT
CTGACGTAAGGGATGACGCACAACTGGTACTTGCTATATAAGGTTTTGCTATTCATTGAAAGCAGTAG
TGACTGATTTGTATATA
>MinSyn_1251|Strength:0.079008272
GCGTGTCGTTTTAGTGAGGATCCTTACCGCTATGGGTAAGATTGGATGAGCTAAGCACATACGTCAGA
ATTAGGTTGGAAAAAGAAGAGGTCTTGTGTACAGGGCTCACTGCTATCAGAAGATCAAAGGGCTATCA
CATGTACTGACGTAAGGGATGACGCACAAGCAAGACCTTATGACCCCCGCCGATGACGCGGGAGTGAA
GCATCTTCCGTCGTGGGAGCCACCAGCACCTCTATATAAGGTTTTGCTATTCATTGAAAGCAGTAGTG
ACTGATTTGTATATA
>MinSyn_181|Strength:0.079606806
GCGTGTCGTTTTAGTGAGGAAAAATGTCAAAGATATAGGTCACGACCACTATGCCCAACTGACGTAAG

GGATGACGCACACACTATGCCCGTGGGAGCCACCAAAGACCTCTACAAAACTGGTATTGCGATAAAGG
AAAGGGACCTATGAGCTAAGCACATACGTCAGCTGCTGAAGATAAGATAATAATGTTGAAGATAAGAA
CCGATGCCTATATAAGGTTTTGCTATTCATTGAAAGCAGTAGTGACTGATTTGTATATA
>MinSyn_1614|Strength:0.079684025
GCGTGTCGTTTTAGTGAGGAACCACGTCTACAATGCCCAATTAGGTTGTCTGCACCTCACATAGCAAG
TGGATCAATTAGGTTGTCTGATGACGTAAGCCATGACGTCTAAGGAGGACCGATGCTGATCTTGAGGT
GGCTCCTACTCATCACTATCAGCCACTATGCCCAATTAGGTTGTCATGAGCTAAGCACATACGTCAGG
GTACTTGTGTACAGGGCTCTATATAAGGTTTTGCTATTCATTGAAAGCAGTAGTGACTGATTTGTATA
TA
>MinSyn_1254|Strength:0.079729326
GCGTGTCGTTTTAGTGAGGATGACGTAAGCCATGACGTCTAAGCAAGACCTCTACAAAACTGGTATTA
TGACCCCCGCCGATGACGCGGGACTTGTGTACAGGGCTCACTGCCTGACGTAAGGGATGACGCACATT
AGCACACCAGCATGTGTTGATCACCAGCTTCAGCTTAGCAAGACCTCTACAAAACTGAAGCATCTTCC
CACTGCTAGGAGGACCGACTATATAAGGTTTTGCTATTCATTGAAAGCAGTAGTGACTGATTTGTATA
TA
>MinSyn_1169|Strength:0.080347598
GCGTGTCGTTTTAGTGAGGATCCTTACCGCTATGGGTAAGATTACTATGCCCAATTAGGTTGATGAGC
TAAGCACATACGTCAGATGTAGGCTATCAACCACGTCTACAAGCCGGAAAAAGAAGAGGTCTTGTGTA
CAGGGCTCACTGCTAGGATTGCGATAAAGGAAAGGGCAAGCTGACGTAAGGGATGACGCACACTTAGC
AGCCACTTGTGTGTAGGCTATCAGCTTAGCAAGACCTCTACTCAGAAGATCAAAGGGCTATATCAGCT
ATATAAGGTTTTGCTATTCATTGAAAGCAGTAGTGACTGATTTGTATATA
>MinSyn_1912|Strength:0.080438168
GCGTGTCGTTTTAGTGAGGATGAGCTAAGCACATACGTCAGTCTGCACCTCCTGACGTAAGGGATGAC
GCACATTCTATATAAGGTTTTGCTATTCATTGAAAGCAGTAGTGACTGATTTGTATATA
>MinSyn_1735|Strength:0.082703058
GCGTGTCGTTTTAGTGAGGCTGACGTAAGGGATGACGCACATTAGGTTGTCTGCACCTCACATGTAGG
CTATTATGACCCCCGCCGATGACGCGGGACTACAAAACTGGTACTTGTGTACAACCATTATTGCGAAT
GAGCTAAGCACATACGTCAGACTGGTACTTGTGTACAGGGCTCATCTCTCTGCCGACAGTGGTCCCAA
AATGCTGATCTTGCCTGCCTTGATGGAAAAAGAAGAGGTACATGTGAAGCATCTTCCCTCTATATAAG
GTTTTGCTATTCATTGAAAGCAGTAGTGACTGATTTGTATATA
>MinSyn_1780|Strength:0.084241172
GCGTGTCGTTTTAGTGAGGCTGACGTAAGGGATGACGCACAACTTGTGTACAGGATGAGCTAAGCACA
TACGTCAGGCGGTAGGTATGACGTAAGCCATGACGTCTAGGTGAAGCATCTTCCCAAGACCTCTACAA
AACTGGTACTCTATATAAGGTTTTGCTATTCATTGAAAGCAGTAGTGACTGATTTGTATATA
>MinSyn_1133|Strength:0.091253226
GCGTGTCGTTTTAGTGAGGAGGTGGCTCCTACTACAGGGCTCAATTTCGGGAAACCTCCTCGGCTCAC
TGCTAGGAGGACCGATGCTGATCTCTGACGTAAGGGATGACGCACAATCAGCTTATCACTATCAGCGG
TTGTCTGCACAAATATTTCTTGTTCTGCACCTATCGAAAGGACAGTACGATGCTGATCTAGCAAGTGG
ATCACGACCACTATGCATGACGTAAGCCATGACGTCTAGCACCTCACATGTAGGCTCTATATAAGGTT
TTGCTATTCATTGAAAGCAGTAGTGACTGATTTGTATATA
>MinSyn_1486|Strength:0.091991677
GCGTGTCGTTTTAGTGAGGATGACGTAAGCCATGACGTCTAGATGCTGATCTTGCCTGCCTTGTTATG
ACCCCCGCCGATGACGCGGGAACAAAACTGGTACTTGTGTACAGGGATGAGCTAAGCACATACGTCAG
GCCCAATTAGGTTCTCTCTGCCGACAGTGGTCCCAAAGCTCTGACGTAAGGGATGACGCACATGCACC
TCACATGTAGGCTATTGCGATAAAGGAAAGGCGGTAGGTCACGACCACGCTTTGTCAAAAGCTAAAAA
AGATGATGCCTGGTACTTGTAGGTGGCTCCTACCTTGCCTGCCTTGCTATATAAGGTTTTGCTATTCA
TTGAAAGCAGTAGTGACTGATTTGTATATA
>MinSyn_1360|Strength:0.09216997
GCGTGTCGTTTTAGTGAGGATGACGTAAGCCATGACGTCTAGCTCACTGCTAGGAGGACCATCCTTAC
CGCTATGGGTAAGATTCCTGCCTTGTCACTATCAGCGTAGGCTATCAGCTTAGATGAGCTAAGCACAT
ACGTCAGCGATGCTGATCTTGCCTGCCTTGATGTCCATCAACAAATAATCCAAGTAAGTATCAGCTTC
TATATAAGGTTTTGCTATTCATTGAAAGCAGTAGTGACTGATTTGTATATA
>MinSyn_1641|Strength:0.092326662
GCGTGTCGTTTTAGTGAGGAACCACGTCTACAAGGTACTTGTGTACAGGGCTCACGCACACCAGCATG

TGTTGATCACCAGCTTCTTGCCTTTATGACCCCCGCCGATGACGCGGGATACAGGGCTCACTGCATGA
CGTAAGCCATGACGTCTACGGAAAAAGAAGAGGTTACTTGTGTACAGGGCTCACTGCAAGATTGATGA
AAAGTCAAAAACAAAAATCAATTATCACTGCTAGGAGGACCGATGCTGATCTTGCCAGCCACTTGTGT
CTATCAGCTTAGCAAGACCTCTACAAACTGACGTAAGGGATGACGCACAGTACTTGTGTACAGGGCTA
TATAAGGTTTTGCTATTCATTGAAAGCAGTAGTGACTGATTTGTATATA
>MinSyn_1829|Strength:0.092455952
GCGTGTCGTTTTAGTGAGGATGACGTAAGCCATGACGTCTACACGACTCCATCAACAAATAATCCAAG
TAAGCTTAGCAAGACCTCTACAACAGCCACTTGTGTAGGTCACGACCACTACTGACGTAAGGGATGAC
GCACACTGCACCTCACATGTAGGCTATCAGAATTTCGGGAAACCTCCTCGACTGCTAGGAGGCTATAT
AAGGTTTTGCTATTCATTGAAAGCAGTAGTGACTGATTTGTATATA
>MinSyn_1434|Strength:0.093750719
GCGTGTCGTTTTAGTGAGGAGCAAGTGGATACGACCACTATGCCCAATTAGGTTTCCATCAACAAATA
ATCCAAGTAAGCAGCTTAGCAAGACCTCTACACTGACGTAAGGGATGACGCACAGTCACGACCACTAT
GAATTTCGGGAAACCTCCTCGCGCGGTAGGTCACGAAAAATGTCAAAGATAGACCGATGCTGATCTCA
AATATTTCTTGTGTAGGCTATCAGCTTAGCAAGACCTCTAATGAGCTAAGCACATACGTCAGACCTCT
ACAAAACTGGTACTATATAAGGTTTTGCTATTCATTGAAAGCAGTAGTGACTGATTTGTATATA
>MinSyn_1383|Strength:0.095013158
GCGTGTCGTTTTAGTGAGGCTGACGTAAGGGATGACGCACAACTATGCCCAATTAGGTTGTCTGCACC
TCAAACCACGTCTACAACTTGATGAGTGGGAGCCACCAGGTACTTGTGTACAATCCTTACCGCTATGG
GTAAGATTGCGGTAGGTCACGACCACTATGCCCAATTAGGAAAAAGAAGAGGTTTATGACGTAAGCCA
TGACGTCTAGGCTATCAGCTTAGCAAGACCTCTACATCTCTCTGCCGACAGTGGTCCCAAAGGTACTT
GGTGGAGCACGACAGTTGTCTGCACCTCACATGTAGGTCAGAAGATCAAAGGGCTACACTGCTACTAT
ATAAGGTTTTGCTATTCATTGAAAGCAGTAGTGACTGATTTGTATATA
>MinSyn_1609|Strength:0.096985134
GCGTGTCGTTTTAGTGAGGTTTTCAACAAAGGCCTGACGTAAGGGATGACGCACAATGCTGATCTTGC
CTGCGTGGGAGCCACCACACCTCACATGTAGAGCAAGTGGATCTCTACAAAACTGGTACTTGTGTACA
ATGACGTAAGCCATGACGTCTACAAGACCTCTGAAGCATCTTCCCCGATGCTGATCCTATATAAGGTT
TTGCTATTCATTGAAAGCAGTAGTGACTGATTTGTATATA
>MinSyn_1870|Strength:0.097195721
GCGTGTCGTTTTAGTGAGGATGACGTAAGCCATGACGTCTATCACATAATTTCGGGAAACCTCCTCGT
GCCCAATTATCCTTACCGCTATGGGTAAGATTAGGGCTCACTGCTCTGACGTAAGGGATGACGCACAG
TAGGTGAAGCATCTTCCCTGGTACTTGTGTACAGGGCTCTATATAAGGTTTTGCTATTCATTGAAAGC
AGTAGTGACTGATTTGTATATA
>MinSyn_1325|Strength:0.100001438
GCGTGTCGTTTTAGTGAGGAATTTCGGGAAACCTCCTCGTCAGCTTAGCAAGACCTGACGTAAGGGAT
GACGCACACACTATGCCCAATTAGGTTGTCTGCACAGCAAGTGGATGTAGGTCACGACCACTATGCCC
ATTGCGATAAAGGAAAGGATGTAGGCTATCAGCTTAGCAAGACCTCTGGAAAAAGAAGAGGTCTGATC
TTGATGACGTAAGCCATGACGTCTACTGGTACTTGTGTACAGGGCTCGCACACCAGCATGTGTTGATC
ACCAGCTACTATATAAGGTTTTGCTATTCATTGAAAGCAGTAGTGACTGATTTGTATATA
>MinSyn_1819|Strength:0.100413018
GCGTGTCGTTTTAGTGAGGTCAGAAGATCAAAGGGCTATTAGCAAGACCTCTACAAAACTCTGACGTA
AGGGATGACGCACACCACTATGCCCAATTATCGAAAGGACAGTAACAAAACTGGTACTATTGCGATAA
AGGAAAGGATCTTATGACGTAAGCCATGACGTCTATGCTAGGAGGATCTCTCTGCCGACAGTGGTCCC
AAATAGGCTATCAGCTTAGCAAGACCTATGAGCTAAGCACATACGTCAGCTTAGCAAGACCTCGCACA
CCAGCATGTGTTGATCACCAGCTCTCACATGTAGGCTATCACTATATAAGGTTTTGCTATTCATTGAA
AGCAGTAGTGACTGATTTGTATATA
>MinSyn_1701|Strength:0.101038208
GCGTGTCGTTTTAGTGAGGTCAGAAGATCAAAGGGCTAGTCACGACCACTAATGAGCTAAGCACATAC
GTCAGTACTTGTGTACAGGGCTCACTGCTAGGAGGAACCACGTCTACAATGCTGATCTTGCCTGCCTT
GATGATCGAAAGGACAGTATATCACTATCAGCTCTGCACCTCACATATGACGTAAGCCATGACGTCTA
TTAGGTTGTTGAAGCATCTTCCGCCTGCCTTCTATATAAGGTTTTGCTATTCATTGAAAGCAGTAGTG
ACTGATTTGTATATA
>MinSyn_1373|Strength:0.104475847
GCGTGTCGTTTTAGTGAGGATGAGCTAAGCACATACGTCAGACATGTAGGCTATCAGCTTAGCAAAAC

CACGTCTACAACTTGTCAGCCACTTGTGTAAGACCTCTACAAAACTGGTACTTGTGTACGCACACCAG
CATGTGTTGATCACCAGCTACCGATGCTGATCTTGCCTGCCTTGATGATCTGACGTAAGGGATGACGC
ACATCGCGGTAGGTCACGAGCAAGTGGATCCCAATTAGGTTGTCTGCACCTCACATGTGGAAAAAGAA
GAGGTGTGTACAGGGCTCTATATAAGGTTTTGCTATTCATTGAAAGCAGTAGTGACTGATTTGTATAT
A
>MinSyn_1191|Strength:0.105653793
GCGTGTCGTTTTAGTGAGGATGAGCTAAGCACATACGTCAGGTGAAGATAAGATAATAATGTTGAAGA
TAAGACTCTACAAAACTCAGTGGTCCCTCCACTAGGTTGTCTGCACCGCACACCAGCATGTGTTGATC
ACCAGCTCATTCAGAAGATCAAAGGGCTAAGGACCGATGCTGATCTTGCCTGCCTAGCAAGTGGATTC
ACTGCTAGGAATGACGTAAGCCATGACGTCTATCTGCACAACCACGTCTACAATCTTGCCTGCCTTGA
TAAGATTGATGAAAAGTCAAAAACAAAAATCAATTATCTCACATGTAGGCTATCAGCTTAGCAAGACT
ATATAAGGTTTTGCTATTCATTGAAAGCAGTAGTGACTGATTTGTATATA
>MinSyn_1804|Strength:0.105672962
GCGTGTCGTTTTAGTGAGGCTGACGTAAGGGATGACGCACAATCATCACTATCAGCACGACCACTATG
CCAAGATTGATGAAAAGTCAAAAACAAAAATCAATTATGCTATCAGCTTAGCAAGACCTCTTATGACC
CCCGCCGATGACGCGGGATGTAGGCTATCAGCTTAGCAAGACCTCTACAAATATTTCTTGTGATGACG
TAAGCCATGACGTCTAGTAGGCTATCAGCTTAGCAAGACCTCTACAATCGAAAGGACAGTACAATTAG
GTTGTCTGCACTCTCTCTGCCGACAGTGGTCCCAAAACTATATAAGGTTTTGCTATTCATTGAAAGCA
GTAGTGACTGATTTGTATATA
>MinSyn_1900|Strength:0.1097612
GCGTGTCGTTTTAGTGAGGTGAAGCATCTTCCCCACTATGCCCAATTAGGTTGTCTGCACCTATGAGC
TAAGCACATACGTCAGAAACTGGTACTTGTGATCCTTACCGCTATGGGTAAGATTATGCCTTATGACC
CCCGCCGATGACGCGGGAGGTAGGTCACGACCACTATGCAACCATTATTGCGCAAAACTGCAGCCACT
TGTGTCCACTATGCCTGGTGGAGCACGACACGGTAGGTCACGACCACTATGCCCTGACGTAAGGGATG
ACGCACAGTAGGTCACGACCACTAAGCAAGTGGATCTCACATGTAGGCTATCTATATAAGGTTTTGCT
ATTCATTGAAAGCAGTAGTGACTGATTTGTATATA
>MinSyn_1968|Strength:0.110213888
GCGTGTCGTTTTAGTGAGGAGGTGGCTCCTACTTAGCAAGACCTCTACAAAACTGGTACTTGTGGTGG
AGCACGACAGATGACGTAAGCCATGACGTCTACCCAATTAGGTTGTCTGCACCTCACATGCAGTGGTC
CCTCCACTGATCTTGCCGGAAAAAGAAGAGGTGCAAGAATCCTTACCGCTATGGGTAAGATTGCTTAG
CTCCATCAACAAATAATCCAAGTAAGTGTGTACAGGGCTCACCTGACGTAAGGGATGACGCACATGCC
CAACTATATAAGGTTTTGCTATTCATTGAAAGCAGTAGTGACTGATTTGTATATA
>MinSyn_139|Strength:0.11021439
GCGTGTCGTTTTAGTGAGGGTGGGAGCCACCATTAGGTTGTCTGCACCTCACAATGACGTAAGCCATG
ACGTCTATGCTGATCTTGCCTGCCTTGATGAATTTCGGGAAACCTCCTCGTCACTGCTAAGATTGATG
AAAAGTCAAAAACAAAAATCAATTATAAGACCTCTACAAAACTGGTCACTATCAGCCTGCCTTGATGA
TATGAGCTAAGCACATACGTCAGAAGACCTCTACAAAACTGGTCTATATAAGGTTTTGCTATTCATTG
AAAGCAGTAGTGACTGATTTGTATATA
>MinSyn_1151|Strength:0.112497872
GCGTGTCGTTTTAGTGAGGATGACGTAAGCCATGACGTCTAACATGTAGGCTATCAGCTTAGCATCAC
TATCAGCTCTGCACCTCAACCATTATTGCGACTATGCCCAATTAGGTTAACCACGTCTACAAACCACT
ATGCCCAATTAGGTTGTCTGCAATTGCGATAAAGGAAAGGGATGCTGTGGGAGCCACCATCTGCACCC
TGACGTAAGGGATGACGCACATAGCAAGACCTCTACAAAACTGGTACTCTCTCTGCCGACAGTGGTCC
CAAAGGTTGTCTGCACCTCACATCTATATAAGGTTTTGCTATTCATTGAAAGCAGTAGTGACTGATTT
GTATATA
>MinSyn_1526|Strength:0.117509529
GCGTGTCGTTTTAGTGAGGATGACGTAAGCCATGACGTCTAGGTTGTCTGCACAATTTCGGGAAACCT
CCTCGCTGATCTTGCCTGCCTTGATGCTTTGTCAAAAGCTAAAAAAGATGATGCTGCACCTCACATGT
AGGCTAATGAGCTAAGCACATACGTCAGGAGGACCGATGCTGATCCTATATAAGGTTTTGCTATTCAT
TGAAAGCAGTAGTGACTGATTTGTATATA
>MinSyn_1563|Strength:0.119493104
GCGTGTCGTTTTAGTGAGGATGACGTAAGCCATGACGTCTAAGGACCAGCAAGTGGATCATCCTTACC
GCTATGGGTAAGATTTTGCCTGCCTTGATGATTCACTATCAGCACCACTATGCCCAATTAGATTGCGA
TAAAGGAAAGGACTGGTACTTATGAGCTAAGCACATACGTCAGGTCACGACCACTATGCCCAATTCTA

```
TATAAGGTTTTGCTATTCATTGAAAGCAGTAGTGACTGATTTGTATATA
>MinSyn_1500|Strength:0.119523619
GCGTGTCGTTTTAGTGAGGATGAGCTAAGCACATACGTCAGAAAACTGGTACTTGTGTTTTTCAACAA
CACTATGCCCAATTAGGTTGTCAACCATTATTGCGAGACCTCTACAAAACGGAAAAAGAAGAGGTCTT
AGCAAGATGACGTAAGCCATGACGTCTAAGCAAGACCTCTACTATATAAGGTTTTGCTATTCATTGAA
AGCAGTAGTGACTGATTTGTATATA
>MinSyn_1462|Strength:0.123242925
GCGTGTCGTTTTAGTGAGGAACCACGTCTACAAACCACTATGCCCAATTAGGTTGATGAGCTAAGCAC
ATACGTCAGCTATGCCCAATTAGGTCAGTGGTCCCTCCACACCGATGCTGATCTTGCCTGCCATCGAA
AGGACAGTACGCGGTACAGCCACTTGTGTGTTTTTCAACAATACTTGTGTACAGGGCAAATATTTCTT
GTAAACTGGTACTTGTGTACAGGGCTAAAAATGTCAAAGATACGGTAATGACGTAAGCCATGACGTCT
ACTGATCTTGCCTATATAAGGTTTTGCTATTCATTGAAAGCAGTAGTGACTGATTTGTATATA
>MinSyn_1637|Strength:0.126837233
GCGTGTCGTTTTAGTGAGGCTGACGTAAGGGATGACGCACACACTATGCCGCTTTGTCAAAAGCTAAA
AAAGATGATGCACTATGCCCAGTGGGAGCCACCAACCACTATGCCCAATTAGGTTGTTTTCAACAATG
TACAGGGCTCACTCACTATCAGCGCTAGGAGGACCGATGCTGATCTTGCCTATGAGCTAAGCACATAC
GTCAGGCTATCAGCTTAGCAAGACCTCCTATATAAGGTTTTGCTATTCATTGAAAGCAGTAGTGACTG
ATTTGTATATA
>MinSyn_1556|Strength:0.127145529
GCGTGTCGTTTTAGTGAGGATGAGCTAAGCACATACGTCAGGCAAGACCTCTACAAAACTGGTGTGGG
AGCCACCAATCTTGCCTGCCTTGATGAGGTGGCTCCTACACTTGTTGAAGATAAGATAATAATGTTGA
AGATAAGAGGCCAGCCACTTGTGTAGGAGGACCGATGCTGATCTTGCTGACGTAAGGGATGACGCACA
GTCTGCACCTCACATGTAGCTATATAAGGTTTTGCTATTCATTGAAAGCAGTAGTGACTGATTTGTAT
ATA
>MinSyn_1569|Strength:0.13425283
GCGTGTCGTTTTAGTGAGGCTGACGTAAGGGATGACGCACAAGGCTATCAGCTTAGCAAGACCTCTAC
AAAAATTTCGGGAAACCTCCTCGAGGGCTCACTGCTAGGAGGACCGATGAACCATTATTGCGTGTAGG
CTATCAGCTTAGTCCATCAACAAATAATCCAAGTAAGTATCAAATATTTCTTGTGGACCGATGCTGAT
CTTGCCTGCCTTGATGATGAGCTAAGCACATACGTCAGTACTTGTGTACAGGGCTCACCTATATAAGG
TTTTGCTATTCATTGAAAGCAGTAGTGACTGATTTGTATATA
>MinSyn_159|Strength:0.136308251
GCGTGTCGTTTTAGTGAGGATGACGTAAGCCATGACGTCTAGCTAGGAGGACCGATGCTGATCTTGCC
TATCGAAAGGACAGTAACTTGTGTACAGGTGGTGGAGCACGACAGGTACTTGTGTACAGGGCTCGTGG
GAGCCACCACGATGCTGACAGCCACTTGTGTATTAGGTTGTCTGCACCTCACATGTAGGCTTGAAGCA
TCTTCCTCAGCTTAGCAAGACCTCTACAAAACTGGTCTGACGTAAGGGATGACGCACATACAGGGCTC
ACTGCTAGGAGGACTATATAAGGTTTTGCTATTCATTGAAAGCAGTAGTGACTGATTTGTATATA
>MinSyn_1904|Strength:0.139816956
GCGTGTCGTTTTAGTGAGGATGACGTAAGCCATGACGTCTAGGCTATCAGCTTAGCGGAAAAAGAAGA
GGTACTATGCCCAATTAGGTTGTCTATCGAAAGGACAGTACGCGGTAGGTCACGCAAATATTTCTTGT
TTTATGACCCCCGCCGATGACGCGGGAGTACAGTGGTGGAGCACGACAGGCTATCAAATTTCGGGAAA
CCTCCTCGGATCTTGCCTGCCTTGATTCTCTCTGCCGACAGTGGTCCCAAATGCCTGCCTTGATATGA
GCTAAGCACATACGTCAGCACCTCACATGTAGGCTATCAGCTTCTATATAAGGTTTTGCTATTCATTG
AAAGCAGTAGTGACTGATTTGTATATA
>MinSyn_1824|Strength:0.142253509
GCGTGTCGTTTTAGTGAGGCTGACGTAAGGGATGACGCACATGTAGGCTATCAGCTTAGCAAAGATTG
ATGAAAAGTCAAAAACAAAAATCAATTATATCTTGCCTGCCTTGATGAGGAAAAAGAAGAGGTCAAGA
CCTCTACAAAACTGGTACTTCACTATCAGCAGGCTATCAGCTTTCTCTCTGCCGACAGTGGTCCCAAA
TACAAAACTGGTACTTGTGTACAGGGCCAGTGGTCCCTCCACATCCATCAACAAATAATCCAAGTAAG
GAGGACCGAATGAGCTAAGCACATACGTCAGCGCGGTAGGTCACGACCACTATGCTATATAAGGTTTT
GCTATTCATTGAAAGCAGTAGTGACTGATTTGTATATA
```
